# Evolution of Menopause and the Mama’s Boy Hypothesis

**DOI:** 10.64898/2026.06.17.732386

**Authors:** Tami Yosef, Liran Samuni, Yoav Ram

## Abstract

Post-reproductive lifespan is an evolutionary puzzle. In most mammals female fertility tracks survival, yet humans and a few toothed whales show survival after reproduction ends. Explaining when and why post-reproductive lifespan evolves is central to understanding the evolution of ageing, social structure, and intergenerational helping across species. Kinship-dynamics theory predicts that when males are philopatric, a female’s local relatedness—especially to male descendants—increases with age, potentially favoring late-life helping over continued reproduction. We develop an age-sex-structured kin-selection model to test whether a rare menopause-inducing modifier allele can invade an initially non-menopausal population through its direct effects on survival and fecundity and its indirect effects on relatives. We consider two evolutionary pathways: stop early, where reproduction ceases earlier with little change in lifespan, and live long, where lifespan extends beyond reproduction under disposable-soma trade-offs. Parameterized with demographic, dispersal, and helping-effect estimates from eight mammalian taxa, the model predicts empirically plausible ages of reproductive cessation and post-reproductive representation in humans and killer whales, but no invasion across plausible cessation ages in non-menopausal taxa. Global sensitivity analyses identify male dispersal and the effect of post-reproductive help on male survival as determinants of whether menopause evolves, motivating the “mama’s boy hypothesis”: menopause is most strongly favoured by selection when late-life care increases the survival and lifetime fitness of philopatric sons and grandsons.

## INTRODUCTION

Menopause is the irreversible cessation of female reproduction followed by a substantial post-reproductive lifespan (PRLS). Menopause is rare in mammals: besides humans, it has been documented in five toothed-whale species^1^, and possibly chimpanzees^2^. Menopause is evolutionarily puzzling because it ends female reproduction and thus incurs a direct fitness cost. The mother, grandmother, and reproductive conflict hypotheses propose that menopause can be favored when indirect fitness benefits and/or reduced reproductive overlap outweigh the direct loss of late-life reproduction^3^.

The conditions under which these hypotheses are met depend critically on kinship dynamics—how a female’s local relatedness to group members changes with age—which are shaped by dispersal and mating^4^. Under male-biased dispersal (common in mammals including baboons and elephants)^5^, female local relatedness does not increase with age as male kin leave and unrelated males enter (Figure 1EFGH), limiting selection for late-life helping. Under male philopatry (e.g., killer whales, humans and chimpanzees; Figure 1ABCD), a female’s relatedness can increase with age as male descendants remain in the group, strengthening kin selection for late-life helping^4^. Female-female relatedness, however, is expected to remain constant throughout a female’s life. Under female philopatry, females are born into groups with close female kin, so relatedness is high from the start; as generations progress, older female kin are effectively replaced by daughters and granddaughters, keeping relatedness similar. Under female dispersal, they enter groups without close female kin, and dispersing daughters prevent increase in relatedness to females, so relatedness remains low.

**Figure 1.**
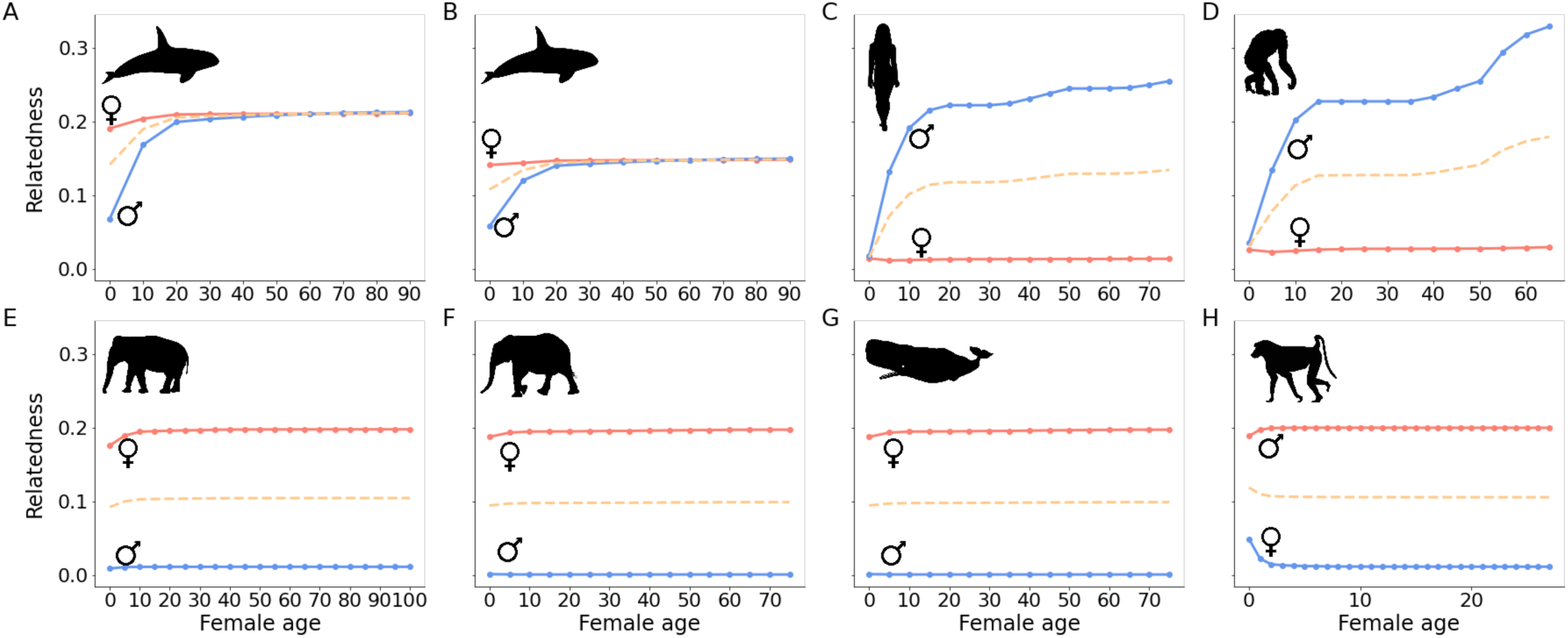
Local relatedness increases with female age under male philopatry. Parameters. *d*_*f*_, *d*_*m*_ denote female and male dispersal rates (0=philopatry, 1=dispersal), and *m* denotes the extent of mating outside the local group (0=outbreeding, 1=inbreeding). When males remain in the natal group (A-B: killer whales, C: humans, D: chimpanzees), female-male relatedness (blue line) increases with female age, and so does the average local relatedness (yellow line), while female-female relatedness (pink line) does not significantly change. **A)** Resident killer whales (*Orcinus orca,* resident ecotype), *d*_*f*_ = *d*_*m*_ = 0.1, *m* = 0. **B)** Transient killer whales (*Orcinus orca,* transient ecotype), *d*_*f*_ = *d*_*m*_ = 0.35, *m* = 0.2. **C)** Humans (*Homo sapiens*), *d*_*f*_ = 0.95, *d*_*m*_ = 0.05, *m* = 1. **D)** Chimpanzees (*Pan troglodytes*), *d*_*f*_ = 0.9, *d*_*m*_ = 0, *m* = 1. **E)** Asian elephants (*Elephas maximus*), *d*_*f*_ = 0.15, *d*_*m*_ = 0.95, *m* = 0.05. **F)** African elephants (*Loxodonta africana*), *d*_*f*_ = 0.15, *d*_*m*_ = 0.995, *m* = 0.01. **G)** Sperm whales (*Physeter macrocephalus*), *d*_*f*_ = 0.15, *d*_*m*_ = 0.995, *m* = 0.01. **H)** Amboseli baboons (*Papio cynocephalus*), *d*_*f*_ = 0.15, *d*_*m*_ = 0.95, *m* = 0.334.

In killer whales, dispersal is rare and mating typically occurs outside the social group, so a female’s local relatedness increases with age (Figure 1AB), especially through sons and grandsons that remain in the group^6,7^. This strengthens inclusive-fitness benefits of late-life helping and amplifies intergenerational reproductive conflict. Indeed, post-reproductive females are associated with increased survival of close kin, particularly adult sons and grandoffspring^8–11^. Menopause has also been reported in short-finned pilot whales, false killer whales, narwhals, and belugas—matrilineal toothed whales that live in long-term but fluid social units where sons and grandsons likewise tend to remain in the maternal group^12–17^—providing a rare comparative system for testing kin-selection explanations.

Ancestral human social organization is often reconstructed as female-biased dispersal with male philopatry. This view is supported by evidence of female dispersal in the genus *Pan*^18–23^ and by population-genetic signals frequently interpreted as stronger female than male gene flow, i.e., mtDNA vs Y-chromosome structure^24–27^. Ethnographic evidence also documents frequent female transfer at marriage in many foragers^28–30^. Under this structure, female’s local relatedness to males can increase with female age as sons and grandsons remain (Figure 1C and 1D), consistent with widespread grandmothering in humans^31,32^ and positive association between older female support and offspring fitness in chimpanzees^33–36^.

However, major gaps remain in understanding how these kinship dynamics translate into the evolution and rarity of menopause across mammals. Reconstructions of ancestral social structure and helping are indirect and cannot, by themselves, determine whether a menopause-inducing allele would be favored by selection. Existing theory rarely combines an explicit modifier-invasion framework with full age and sex structure, and a decomposition of fitness into survival and fecundity components, limiting its ability to predict when post-reproductive lifespan should evolve. Models are also seldom parameterized with comparable demographic data across species or evaluated against empirical post-reproductive representation. Finally, most studies ignore sex-biased helping—even when data suggest strong biases, as in killer whales^6,8–11,37^, leaving open how male-versus female-directed help shapes selection for menopause.

Here, we develop an age- and sex-structured kin-selection model^38,39^ and derive a closed-form invasion criterion for a rare menopause-inducing modifier allele. The modifier sets the age of reproductive cessation and reallocates late-life effort to helping, with kinship dynamics generated by sex-biased dispersal and mating^4^. We represent grandmothering as group-directed helping, mother effects as reduced maternal mortality costs, and reproductive conflict as reduced realized late-life fecundity under intergenerational competition. We analyse two evolutionary pathways, *stop early* and *live long*^1^, parameterize the model across menopausal and non-menopausal taxa, compare predicted versus observed post-reproductive lifespan, and quantify the effect of sex-biased helping on selection for menopause. We synthesize our results with existing male philopatry accounts^4^ as the “mama’s boy” hypothesis: menopause is favored when stopping late-life reproduction frees effort to support philopatric, high-cost males.

## RESULTS

We study a rare, menopause-inducing modifier allele introduced into a non-menopausal, age- and sex-structured population, where females reproduce through late life. Carriers (“donors”) stop reproducing from an allele-specified age and instead help the survival and fecundity of younger group members (“recipients”). We assume homozygotes and matings among carriers are negligible (as the allele is rare) and restrict recipients to younger age classes than the donor.

Selection on the modifier allele is evaluated via inclusive fitness, separating direct and indirect fitness components^38^. The direct component quantifies alleles lost when a donor ceases reproduction. The indirect component sums alleles gained through donor helping to recipients’ survival and fecundity, weighted by donor’s age-specific relatedness, which is generated by sex-biased dispersal and mating using the kinship dynamics framework^4^ (Figure 1). The total inclusive-fitness effect gives the invasion criterion λ: the menopause modifier is expected to invade when λ>0 (Figure 2).

**Figure 2.**
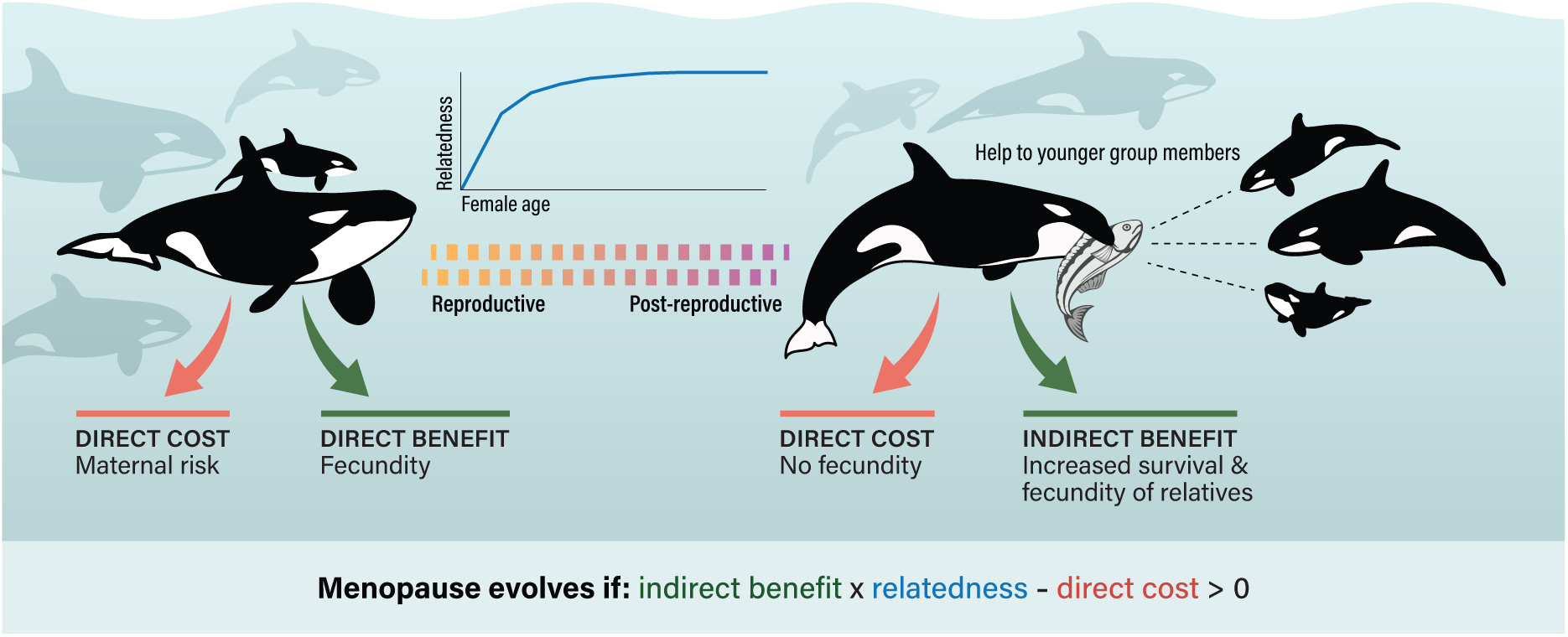
Model overview. A menopause-inducing modifier allele trades late-life fecundity (direct benefit) for post-reproductive helping that increases survival and fecundity of group members (indirect benefit), with benefits weighted by age-specific relatedness. Menopause is favored by selection when λ = indirect benefit x relatedness – direct cost > 0.

Our model incorporates three hypotheses for the evolution of menopause through explicit model assumptions: i) grandmothering as help from post-reproductive females to younger group members; ii) mother effects as reduced late-life maternal risk in post-reproductive females; and iii) reproductive conflict as reduced realized late-life fecundity of older females under intergenerational reproductive overlap. We analyze two life-history pathways to post-reproductive lifespan^1^: *stop early*, where the menopause modifier shortens the reproductive span by zeroing female fecundity after the cessation age; and *live long*, where the modifier extends lifespan beyond the end of reproduction under disposable-soma trade-offs^40,41^, by increasing post-reproductive survival and adjusting early-life survival and fecundity accordingly.

We parameterize the model using published empirical data, compiling life tables (age-and sex-specific survival and fecundity), sex-specific dispersal rates, inbreeding rates, and effects of post-reproductive help on both donor and recipient survival and fecundity across eight taxa with different kinship dynamics (Figure 1). Full details of the model and its parameterization are provided in the Online Methods.

### Model predictions agree with empirical post-reproductive representation values

We used the model to compute λ, the invasion criterion for the menopause-inducing modifier allele, across different ages of reproductive cessation for each species (Figure S1, S2). We then identify whether, and at what cessation age, λ first becomes positive, which marks the earliest age of reproductive cessation at which menopause is favored by selection. We then used this age as the predicted menopause-onset age (M) in the calculation of post-reproductive representation (PrR, the proportion of adult life spent after reproduction) for each species. We find that the predicted PrR values closely match empirical estimates, with minor deviations likely reflecting variation in life-history parameters (Figure 3, Figure S6).

**Figure 3.**
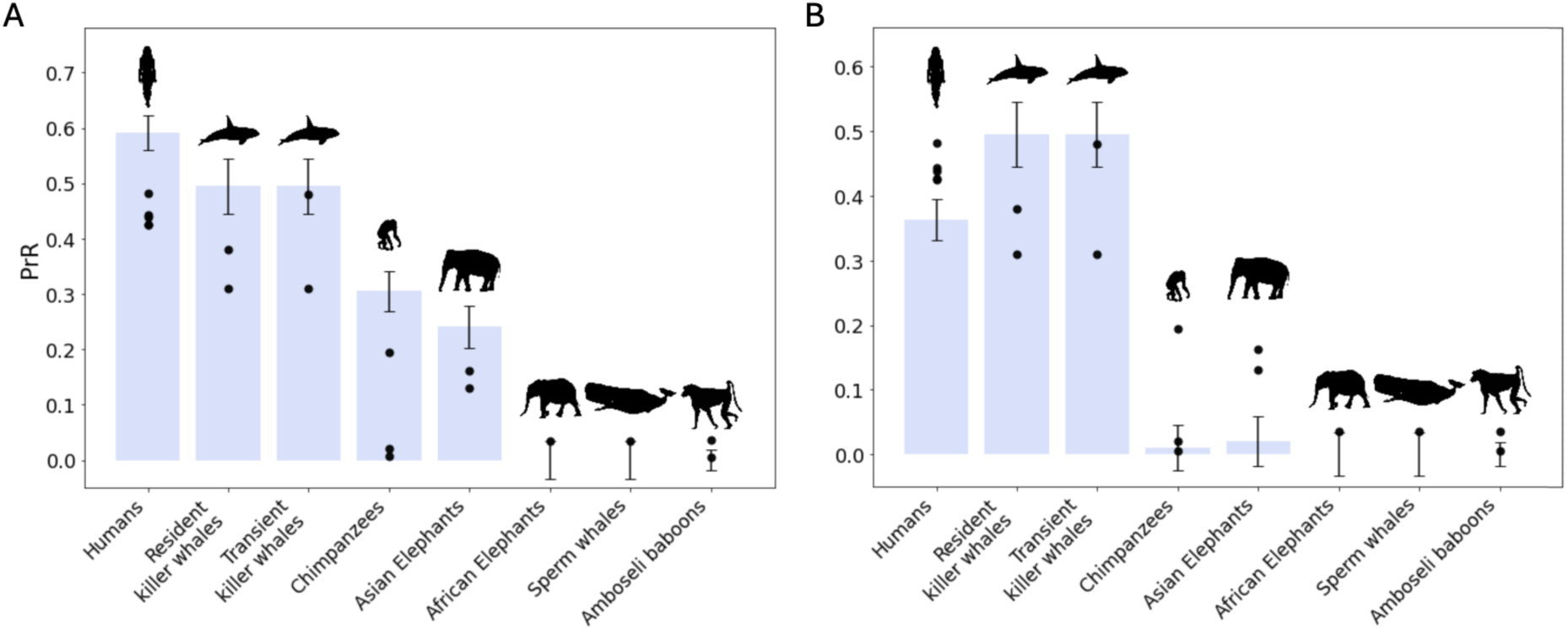
Model predictions agree with empirical post-reproductive representation (PrR) values across species. A) Predicted PrR values under the stop early and B) the live long pathways. Markers are empirical PrR values (Table 2). Bars show predicted PrR using model-predicted menopause onset age (Figures S1 and S2; minimal age at which *λ* > 0). Error bars reflect discretization uncertainty from species-specific age-class width, computed by 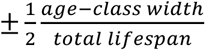.

In known menopausal species, both investigated pathways show strong support. In the stop early pathway, *λ* becomes positive in killer whales at ages 45–55 (Figure S1A-B), consistent with empirical evidence (Figure 3A)^7,42^. In humans, *λ* first becomes positive at ages 32.5-37.5 (Figure S1C), earlier than most empirical estimates of reproductive cessation (Figure 3A)^43^. This discrepancy may reflect uncertainty in the reconstruction of the hypothetical non-menopausal ancestral life table. Under the live long pathway, *λ* can become positive under modest trade-off costs, especially when only a few post-reproductive years are added (Figure S6). As additional years are added, early-life costs increase because larger trade-offs are required for longevity. Simultaneously, the longer helping period allows more indirect benefits to accumulate. Consequently, the parameter space favoring the modifier widens and tolerates higher trade-off costs as additional years are added, making extended post-reproductive lifespan adaptive in humans and two killer whale ecotypes, resident and transient (Figure 3B). This results in a positive *λ* with biologically plausible post-reproductive spans (Figure S2A–C)– approximately 30 years in humans and 50 years in killer whales–implying that inclusive-fitness benefits from prolonged life can offset early-life costs and allow menopause to evolve via increased longevity.

In contrast to humans and killer whales, neither pathway is supported for African elephants or sperm whales, which are both non-menopausal species with female-juvenile family units and males that live alone or in bachelor groups. In the stop early pathway, *λ* remains negative across all ages (Figure S1F-G). Under the live long pathway *λ* also remains negative across the tested range of trade-off parameters and post-reproductive years (Figure S2F–G, S6), even under favorable helping assumptions. These results are consistent with empirical evidence (Figure 3)^42,44^.

In Amboseli baboons, a non-menopausal species with multi-male and multi-female troops characterised by female philopatry and male dispersal^45^, the stop early pathway yields a positive *λ* at the age of 12, with an initial value of 7 × 10^−7^ and a maximum ≈ 10^−6^ (Figure S1H), but the values are 2-3 orders of magnitude lower than in menopausal species. At such levels, kin selection is likely weak in comparison to random genetic drift: the per-generation change in allele frequency due to drift is 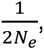 so kin selection requires 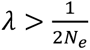. Matching the maximum *λ* implies *N*_*e*_ ≈ 3 × 10^6^. Census population size in Amboseli baboons is far smaller and *N*_*e*_is typically lower^46^. Thus, drift likely dominates selection, making the small positive λ effectively negligible and yielding very low PrR. Similarly, in the live long pathway, menopause evolves only when survival and fecundity trade-offs are vanishingly small (fecundity multiplier ∼1.004 and survival multiplier ∼1.001) if 20 post-reproductive years are added (Figure S2H). We initially focused on 20 added years as a plausible limit given that Amboseli baboons live up to 27 years^45^, but even extending the analysis to 30 added years resulted in similar values. Even with these additions, *λ* remains 2–4 orders of magnitude lower than in species with substantial post-reproductive lifespans such as killer whales, indicating very small inclusive-fitness benefits.

Asian elephants, in which females and juveniles form family units and males live alone or in bachelor groups, appear intermediate between menopausal and non-menopausal species. In the stop early pathway, λ becomes positive at ages 42.5–47.5 (Figure S1E). Their life-history data indicate a short post-reproductive lifespan of ∼11 years^47^. Although short, this post-reproductive stage exceeds that of clearly non-menopausal species, and some authors classify Asian elephants as menopausal^48^. However, unlike clearly menopausal species (i.e., humans and killer whales), where fertility declines sharply after ∼35 and ovulatory function ceases by ∼56^47^, elephants show a shorter, more gradual reduction in realized reproduction while physiological capacity can persist; some females reproduce past 60 years^49^. This pattern is consistent with our invasion criterion: behavioral or social factors can produce a positive and short PrR even when fertility has not fully ceased physiologically, which may explain why Asian elephants appear intermediate between menopausal and non-menopausal species. In the live long pathway, however, menopause evolves in Asian elephants only under specific trade-off regimes (Figure S6). Even then, *λ* remains close to zero and becomes only marginally positive (∼8 × 10^−6^) after ∼40 added years (Figure S2E). This indicates weak selection that may not overcome drift, and suggests that longevity-driven menopause is unlikely to evolve in this species.

Chimpanzees live in multi-male and multi-female communities characterised by male philopatry and female dispersal. In the stop early pathway, λ becomes positive at age 37.5-42.5 (Figure S1D). Earlier studies reported minimal post-reproductive lifespan in the Gombe community in Tanzania^50–52^. By contrast, Wood et al. (2023) recently reported high PrR values (∼0.195) in the Ngogo community in Uganda, indicating extended post-reproductive lifespan in at least some chimpanzee populations.^2^ Reports of old, nonreproductive females in other communities further support a post-reproductive stage (51–53; Samuni, unpublished data). Our results are consistent with a post-reproductive stage in chimpanzees; the higher PrR we predict relative to Wood et al. (2023) (Figure 3A) likely reflects our assumptions on the helping effects of older females^2^. However, the live long pathway results for chimpanzees are weaker; λ remains low and only becomes positive under narrow conditions (Figure S6), highlighting the need to consider multiple evolutionary pathways for this species.

### Evolution of menopause is driven by help towards male kin survival

In killer whales, post-reproductive help is sex-specific and often targeted toward males^9,10,37,56^. Indeed, the effect of post-reproductive help on the invasion criterion depends on the recipient sex (Figure S5). Therefore, we evaluated how these asymmetries shape the evolution of menopause across species using Sobol variance-based global sensitivity analysis^57–59^. The analysis decomposes variance in the invasion criterion λ into the contributions of the model parameters: first-order indices (Figure S3) measure the contributions by each parameter alone, and total-order indices (Figure 4) measure the contributions including all high-order interactions between model parameters.

**Figure 4.**
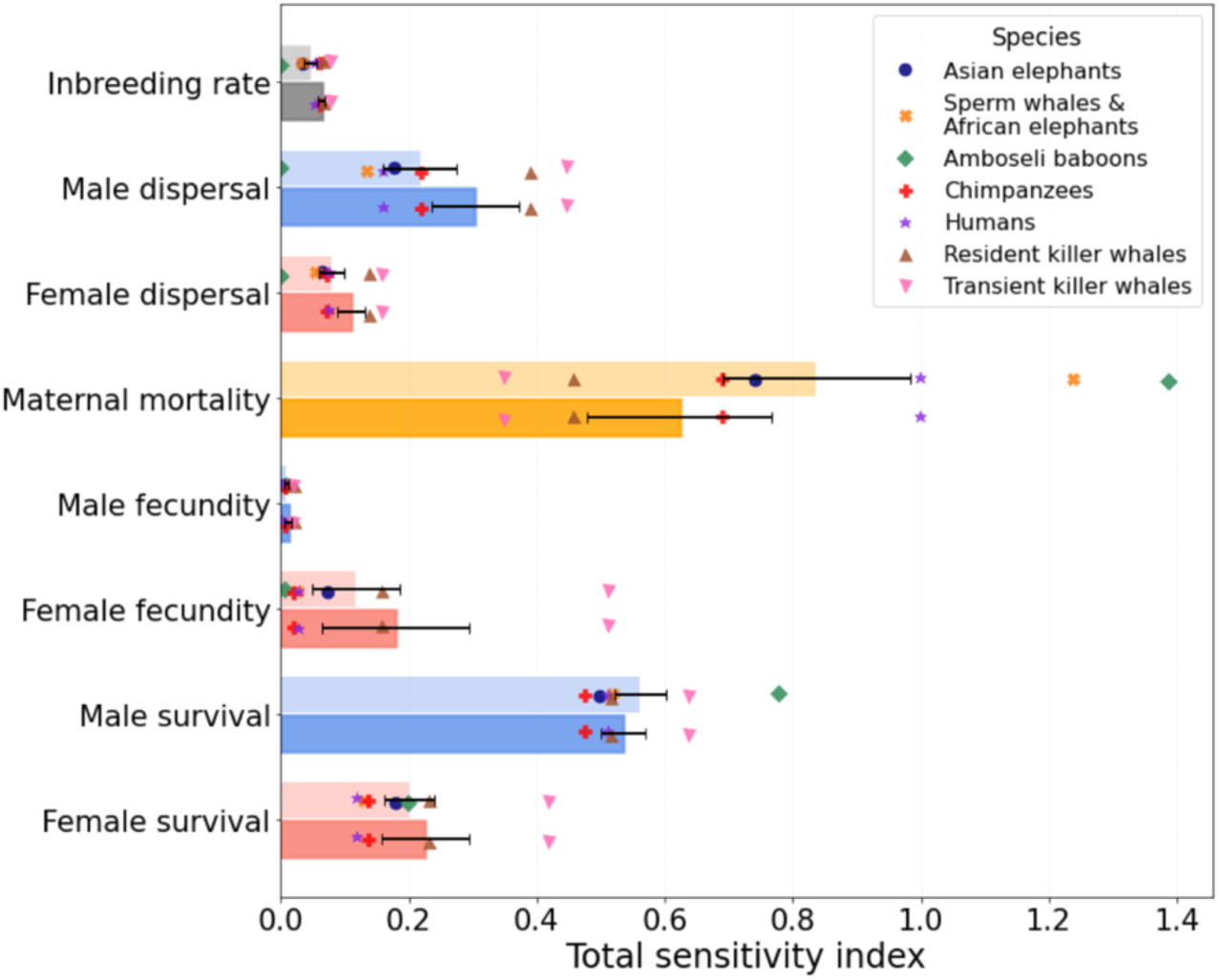
Maternal mortality and post-reproductive help to male survival are the dominant drivers of the evolution of menopause. The total-order sensitivity indices measure the contribution of each parameter (y-axis) to the variance in the invasion criterion λ, including effects through interactions with other parameters. Markers show indices in different species. Dark and light bars show the average indices in menopausal species and all species, respectively.

The sensitivity analysis indicates that the invasion criterion λ is most sensitive to maternal mortality, consistent with the mother hypothesis: cessation of reproduction removes maternal-risk effects, so higher late-life female survival increases the scope for post-reproductive help to relatives. The contribution of maternal mortality increases markedly in the total-order indices (Figure 4) relative to the first-order indices (Figure S3), suggesting amplification through interactions with other parameters. Total-order indices further show that λ is strongly influenced not only by the overall magnitude of post-reproductive help, but also by recipient sex. In particular, help that increases male survival accounts for a substantial share of the first-order (Figure S3) and total-order (Figure 4) sensitivity indices for λ, indicating that this sex-specific effect is robust to parameter interactions. By contrast, help to fecundity of either sex accounts for little variation in λ, plausibly because fecundity effects are confined to specific age classes, whereas survival benefits (and reductions in maternal mortality) can accumulate across multiple age classes.

Male dispersal also has a relatively large effect on λ, especially when it interacts with other parameters (Figure 4). This indicates that the benefits of helping males depend on their tendency to remain in the natal group where maternal help can be expressed. Together, these findings suggest a common evolutionary mechanism: the transition to a post-reproductive lifespan is favored when post-reproductive females provide survival advantages to male descendants that remain locally associated with their mothers and grandmothers.

## DISCUSSION

Here, we developed a sex- and age-structured kin-selection model (Figure 2), parameterized with demographic data and estimates of post-reproductive helping effects from eight mammalian species, to determine when a menopause-inducing modifier can invade a non-menopausal population and to predict the resulting duration of post-reproductive lifespan (PRLS). We analyzed two non-mutually exclusive evolutionary pathways across eight mammalian species: a stop early pathway, in which reproductive lifespan is extended while total lifespan is held constant, and a live long pathway, in which post-reproductive age classes are added to an ancestral life history under disposable-soma trade-offs^1,40,41^. Across species, we find that selection can favor substantial PRLS in humans and both resident and transient killer whales, consistent with their long empirical PRLS (Figure 3). By contrast, under the empirical life histories of African elephants, sperm whales, and Amboseli baboons, selection favors little or no PRLS, consistent with empirical estimates of minimal or no PRLS. Chimpanzees and Asian elephants show an intermediate pattern: given their demographic and helping parameters, selection favors only modest PRLS, and only over a narrower range of conditions compared to humans and killer whales. These results show that our model captures cross-species variation in PRLS.

In killer whales and humans, selection favored reproductive cessation at empirically realistic ages under both pathways (Figure 3). Under stop early, menopause evolved at midlife, matching observed ages of reproductive cessation. Under live long, realistic PRLS were also favored, as late-life helping benefits offset longevity costs under moderate trade-offs. For killer whales, the live long pathway aligns with comparative analyses suggesting that post-reproductive toothed whales have extended total lifespans (Figure 1b in ^1^). For humans, there is also evidence that reproductive senescence has not accelerated compared to our relative species, but rather, that adult survival rates have increased^2,50,60^.

In Asian elephants, the model suggests that menopause via the live long pathway is possible but only in a narrow region of parameter space and is likely to be difficult to realize evolutionarily because the predicted selection is very weak. The stop early pathway can generate selection for midlife reproductive cessation even in a species typically considered non-menopausal, consistent with Asian elephants’ gradual reproductive decline and modest post-reproductive period rather than an abrupt cessation^47^. Overall, these results point to a contingent, context-dependent route to PRLS in Asian elephants, with the live long pathway especially unlikely.

Similar patterns have been reported in long-lived primates not tested here. Gorillas and macaques can show menopause-like physiological changes, but these are most pronounced in captivity^61,62^. In the wild, female gorillas usually do not live long enough for complete reproductive cessation, yet recent long-term data indicate that some mountain gorillas experience extended post-reproductive survival, with several individuals living a decade or more after their last birth and a population-level post-reproductive representation of about 0.1^63^. Captive female western-lowland gorillas also exhibit prolonged reproductive inactivity and hormonal profiles characteristic of menopause, including reduced ovarian activity and low sex-steroid levels^61^. However, both male and female gorillas disperse from their natal groups and reproduce primarily by outbreeding^64^, resulting in low relatedness within groups and little opportunity for intergenerational helping. Although mothers provide prolonged support to their offspring, particularly to sons, this assistance generally ceases when offspring reach sexual maturity, after which mothers have no opportunity to help^65^, and grandmaternal assistance has not been observed. These demographic and social patterns suggest that, while gorillas may experience a limited post-reproductive lifespan, their high dispersal and weak kin structure likely prevent the evolution of adaptive, kin-selected menopause.

In contrast, selection does not favor PRLS in sperm whales and African elephants under both pathways, even with generous kin-helping effects under the live long pathway. Both species maintain high reproductive output late in life and lack the social or demographic structures (such as male philopatry) that might support a shift toward an extended non-reproductive lifespan. In Amboseli baboons, another non-menopausal species, selection for PRLS under the stop early pathway is far too weak to drive the evolution of PRLS. Under the live long pathway, selection is positive only under extreme and biologically implausible assumptions and remains too weak to overcome drift. These results reinforce the view that menopause is highly unlikely to evolve in this species.

In chimpanzees, we found menopause to be potentially favorable under the stop early pathway, with selection for PRLS around ages 40–45. While past studies based on the Gombe population reported minimal post-reproductive lifespan^50^, recently, Wood et al. (2023) documented high PrR values in the Ngogo population, and our model supports this latter view^2^. These differences likely reflect population-specific demography and disturbance: Gombe community is small, more affected by human activity, and experiences frequent disease outbreaks with high mortality and shorter life expectancy^66^, whereas Ngogo shows lower anthropogenic disturbance, larger and more stable demography with higher survival and longevity, and more favourable ecological conditions^67^. Ngogo may thus better approximate ancestral chimpanzee life-histories^68,69^. Importantly, old post-reproductive females have also been documented in other wild chimpanzee communities^53–55^, supporting the presence of a post-reproductive phase. Nevertheless, while selection does emerge under specific conditions under the live long pathway, it remains weak. This contrasts with the stronger selection for PRLS found under the stop early pathway. More generally, in many of these long-lived species we are constrained by the number of years a given population was studied—so only now we are starting to scrape the surface of what happens in late life.

Our model’s measure for selection, the invasion criterion *λ*, is highly sensitive to male-specific parameters (Figure 4), reinforcing the idea that the evolution of menopause is particularly influenced by indirect fitness gained through sons and grandsons. This aligns with observations in resident killer whales, where adult males exhibit a lifelong dependency on maternal support: males over 30 face an eight-fold increase in mortality risk following the death of their mother^9^, and the presence of adult sons, but not daughters, significantly reduces a mother’s subsequent reproductive success^37^. Complementary evidence from pre-industrial human populations points to a similar maternal cost: historical demographic records suggest that each additional son reduced a mother’s post-reproductive survival by ∼34 weeks, whereas no comparable effect was detected for daughters^70,71^, but such effects can be complicated due to cultural and social factors^72^. This suggests that across both humans and killer whales, the two species where our model predicts substantial PRLS, males represent a high-cost investment that requires females to prioritize offspring survival over further reproduction.

Investment in costly males may be a feature of menopausal lineages. A recent comparative analysis across toothed whales shows that menopausal species show pronounced sexual dimorphism, with males ∼20% larger than females on average, compared with ∼5% in non-menopausal toothed whale species^1^. This pattern is consistent with the idea that extended PRLS can help subsidize the energetic demands of producing and maintaining large, competitive males. In matrilineal toothed whales, males typically mate outside the natal group, so maternal support that improves male survival and competitive ability can raise inclusive fitness without intensifying local competition for mates or resources^1,4^.

Prosocial traits that determine the extent of post-reproductive help are central to our model. A sensitivity analysis identified male survival help as the dominant driver of menopause evolution, as well as male dispersal (Figure 4 and S3). Survival benefits accumulate across a recipient’s remaining lifespan and thus across multiple future reproductive opportunities, whereas fecundity benefits are restricted to specific age classes. When males remain in their natal group, increases in male survival generate long-lasting inclusive fitness benefits for mothers and grandmothers. Accordingly, in systems with bisexual philopatry and outbreeding (e.g., matrilineal whales) or with high female but low male dispersal and inbreeding (e.g., ancestral humans), relatedness to males rises with female age while relatedness to females changes little^4^ (Figure 1). This asymmetry magnifies the effect of male survival help on selection for PRLS, which our sensitivity analysis highlights as a particularly important determinant of menopause evolution.

Together with the strong observed effect of male philopatry (Figures 4, S3 and S4), and the fact that males are philopatric in all known menopausal species, our results point to a critical role for costly philopatric males. We therefore propose the *mama’s boy hypothesis*: selection for PRLS is strongest when post-reproductive females can increase the survival, and thus the lifetime reproductive success, or philopatric male descendants, because the inclusive-fitness gains of male survival help compound across a recipient’s remaining reproductive lifespan. In this sense, the mama’s boy hypothesis is a more specific version of the grandmother hypothesis^3,6,43^: rather than help to descendants in general, it predicts that help directed toward sons and grandsons should be disproportionately important.

Future empirical studies could test the mama’s boy hypothesis by quantifying the demographic and behavioral parameters most relevant to the evolution of menopause across a wider range of species, both menopausal and non-menopausal. Such work depends on rare lifetime datasets that follow individuals across decades, allowing researchers to document births, deaths, and reproductive histories in detail. With these kinds of long-term studies, such as the one presented by Weiss et al. (2023), it becomes possible to obtain detailed life-history data, including age-specific survival and fecundity curves, together with information on sex-specific dispersal and mating patterns that enable more precise predictions of kinship dynamics^37^. This would also make it possible to measure how older females, particularly post-reproductive ones, affect the survival and fecundity of male and female descendants—providing improved estimates of helping coefficients and further refining model predictions.

When discussing the evolution of menopause, it is important to distinguish between ultimate (evolutionary) and proximate (social and cultural) explanations. While our findings refine the grandmother hypothesis by showing that menopause likely evolved specifically because of the help females provide to their male descendants, this evolutionary legacy does not rigidly constrain its contemporary meanings or functions. The modern experience and understanding of menopause are shaped by cultural, social, and individual factors that diverge from its original evolutionary drivers. Thus, evolutionary explanations help us understand why menopause came to be, but they do not determine how it must be experienced or understood today.

## Acknowledgements

We thank Lilach Hadany, Darar Bega, Luke Rendell, Marc Feldman, Adi Stern, Lee Koren, Talia Borofsky, Daniel Weissman, Arnon Lotem, Yoav Livne, and Tal Simon for discussions and comments on the manuscript. This work was supported in part by the Israel Science Foundation (ISF 438/25, YR) and the Minerva Center for the study of Population Fragmentation (YR).

Figure 2 in the manuscript was Illustrated by Iris Shpitzer. Silhouettes were sourced from PhyloPic.org: killer whale (*Orcinus orca*) by Chris huh (CC BY 3.0, https://creativecommons.org/licenses/by/3.0/); African elephant (*Loxodonta africana*), Asian elephant (*Elephas maximus*), Amboseli baboon (*Papio cynocephalus*), chimpanzee (*Pan troglodytes*), and sperm whale (*Physeter macrocephalus*) by Steven Traver (CC BY 3.0, https://creativecommons.org/licenses/by/3.0/); and human (*Homo sapiens*) by T. Michael Keesey (CC0 1.0, https://creativecommons.org/publicdomain/zero/1.0/).

## STATEMENTS

We used ChatGPT (OpenAI) to assist with editing and proofreading of the manuscript text.

## CODE AVAILABILITY

All code used to generate the results and figures in this study is provided as Supplementary Code.

## ONLINE METHODS

A comprehensive overview of the model, as well as the selection of parameters, will be provided in this section, with a detailed description of its formulation in the Supplementary material.

### (i) Mathematical model

Our model is based on the work of Charlesworth & Charnov (1981), where they established conditions for the invasion of a rare altruistic allele into a diploid population homozygous for a non-altruistic allele^38^. In our model, the altruistic allele induces menopause, while the initial state of the population – homozygous for the non-altruistic allele – corresponds to a non-menopausal population where females continue reproducing later in life. We emphasize that the allele models reproductive cessation as a functional outcome but does not assume a specific biological mechanism; the cessation could be physiological (hormonal infertility) or behavioral (voluntary abstention from reproduction despite physiological capacity). In humans and killer whales, the non-menopausal populations are theoretical constructs, as these species are menopausal in reality, whereas in other species, the non-menopausal populations reflect actual biological conditions. Using this framework, we aim to determine whether a menopause-inducing allele can spread in such a population by influencing the survival and reproduction of the carrier and her relatives.

The menopause allele defines a "donor" (the female carrying the allele) and "recipients" (her group members). The relatedness between donors and recipients is represented as *r*_g_(*x*_*i*_), where *g* refers to the recipient’s sex and *x*_*i*_ refers to the donor’s age class. This relatedness is calculated using the model by Johnstone & Cant (2010)^4^. Their model presents a demographic-genetic model that analyses how identity-by-descent (IBD) probabilities evolve in a structured population with overlapping generations. The model tracks IBD between gene copies sampled from individuals of the same or different sex within the same patch (or group), incorporating key demographic parameters: sex-specific dispersal rates, birth rates, inbreeding probability, and group sizes. Using recursive equations, it calculates equilibrium IBD probabilities within and between sexes, and across age classes. These are then used to derive relatedness coefficients that vary with age and sex, which eventually quantifies how relatedness to group members changes over an individual’s lifespan.

We assume the menopause allele is initially rare. This enables us to neglect reproduction between carriers and thus disregard individuals homozygous for the menopause allele. Additionally, recipients are limited to age classes younger than the donor, preventing overlap between donor and recipient age classes. Finally, we assume the population grows exponentially at rate *ρ*, which simplifies the mathematical analysis^38^.

To evaluate whether the menopause allele will spread, the model consists of two inclusive fitness components. The direct component quantifies the cost a female carrier (donor) incurs by entering a post-reproductive phase at specific age classes. This cost reflects the reduction in number of offspring that would have inherited the menopause allele, as reproduction ceases earlier than in non-menopausal females. In the stop early parameterization, this is implemented by zeroing all female fecundity after the onset age in the empirical Leslie matrices. In the live long parameterization, the direct cost instead arises from extending lifespan beyond reproduction under disposable soma trade-offs, as described below. The indirect component measures the survival and fecundity benefits conferred to younger relatives (recipients) by the donor’s help, accounting for the proportion of these benefits attributable to the menopause allele. Implementation of these help effects is detailed in the following section.

The model ultimately assesses how many menopause alleles a donor loses directly by ceasing reproduction early, compared to how many alleles are indirectly gained by aiding her relatives. If the indirect benefits outweigh the direct reproductive costs, we expect the menopause allele to spread within the population. This condition corresponds to a positive *λ* in our model.

To describe an altruistic interaction between a donor from age class *x*_*i*_and a recipient from age class *x*_j_ and sex *g*, with relatedness *r*_g_(*x*_*i*_), we start by defining the cost for the donor. The cost comprises two factors: a change in the donor’s fecundity and in her survival probability. We first represent the reduction in her fecundity as δ_*D*_*m*_*f*_(*x*_*i*_), where *m*_*f*_(*x*_*i*_) is the fecundity of a female at age class *x*_*i*_ and δ_*D*_ denotes a change due to helping. Because this reduction occurs only if the donor reaches the relevant age class, it is weighted by the probability of her survival to that age, denoted *t*_*f*_(*x*_*i*_). Next, we normalize this expression by considering the probability that these potential offspring would have been carriers of the menopause allele (altruistic). The birth of an altruistic offspring when the donor’s age is *x*_*i*_ increases the allele frequency by 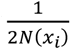, where *N*(*x*_*i*_) is the size of the population when the donor is of age *x*_*i*_. Thus, we must weight each birth 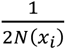. Under exponential growth, this is equivalent to weighting by 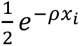.

Putting all these factors together, the expression for the weighted cost of menopause alleles in the population due to a reduction in the donor’s fecundity at age *x*_*i*_ is given by Equation 1.

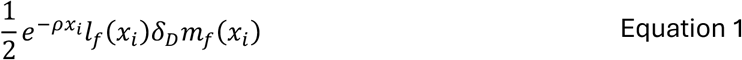

To quantify the change in the number of alleles produced by a donor due to a change in her survival, we first represent the change in survival to the next age class as δ_*D*_*P*_*f*_(*x*_*i*_), where *P*_*f*_(*x*_*i*_) is the probability a female survives from age class *x*_*i*_ to *x*_*i*_ + 1, and δ_*D*_ denotes a change due to helping. Since this occurs only if the donor reaches the relevant age class, it is weighted by the probability that she survives to that age class, denoted as *t*_*f*_(*x*_*i*_). To assess how this change in survival influences the dynamics of the menopause allele, we consider the reproductive value of the donor^73^, which represents the sum of her expected future offspring starting from the next age class (*x*_*i*_ + 1), while accounting for population growth and survival probabilities at each reproductive age. The general formula for reproductive value at age class *x*_*i*_, *v*_g_(*x*_*i*_), is shown in Equation 2.

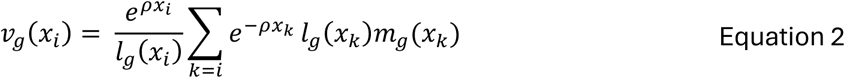

Here, *g* ∈ {*f*, *m*} denotes sex (*f* for female, *m* for male), *ρ* is the population growth rate, *t*_g_(*x*_*k*_) is the probability of surviving to age class *x*_*k*_, and *m*_g_(*x*_*k*_) is the fecundity at that age class.

Next, we normalize this expression by considering the probability that these potential offspring would have been carriers of the menopause allele. Thus, the expression for the weighted loss of altruist alleles in the population due to a change in the donor’s survival is given by Equation 3.

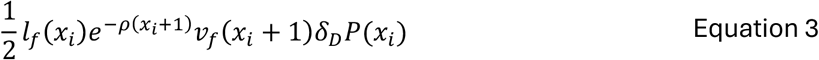

To account for the combined effects of changes in both survival and fecundity, we sum Equations 1 and 3. However, these expressions represent interactions between a single donor age class and a single recipient age class of a specific sex. We must consider interactions across all relevant donor age classes that are older than the menopause onset age (for every *k* ≥ *i*), as shown in Equation 4.

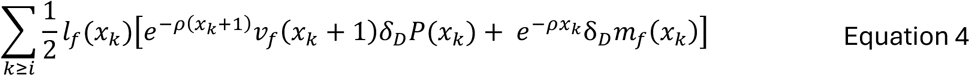

Next, we sum over interactions involving both male and female recipients, resulting in Equation 5.

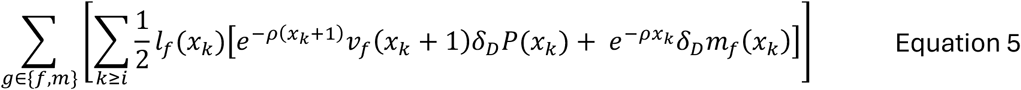

Since the factor ½ is a constant, we can simplify the equation to obtain the final expression for the total donor cost, shown as Equation 6. This represents the comprehensive cost to the donor across all relevant age classes and both recipient sexes, integrating the effects of changes in fecundity and survival.

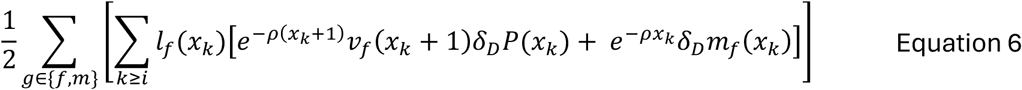

The benefit to recipients also consists of changes in their survival and fecundity due to the interaction. To quantify the number of alleles indirectly gained by a donor of age *x*_*i*_due to an increase in her relative’s fecundity, we first represent the increase in the recipient’s fecundity as δ_*R*_*m*_g_O*x*_j_P, where *m*_g_O*x*_j_P is the fecundity of a group member of sex *g* at age class *x*_j_, and δ_*R*_ denotes a change due to receiving help. This is weighted by the probability that the donor survives to her current age class, denoted as *t*_*f*_(*x*_*i*_), and normalized by the probability that the recipient’s offspring inherit the menopause allele, given by 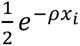. Since the model is sex-specific, we must also consider the proportion of interactions between females and each sex group, defined by the function 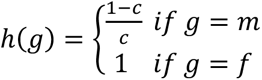, where *c* is the male fraction and 1 − *c* is the female fraction. Additionally, because the age distribution varies across species, we account for the fraction of interactions between donors of age *x*_*i*_ and recipients of a specific age and sex, denoted as *a*_g,j_O*x*_*i*_, *r*_g_(*x*_*i*_)P, where *r*_g_(*x*_*i*_) represents the relatedness between donor and recipient. The value of *a*_g,j_O*x*_*i*_, *r*_g_(*x*_*i*_)P is derived using the sex-specific age distribution formula from Otto C Day (2011)^74^.

Putting these factors together, the expression for the weighted gain of menopause alleles in the population due to an increase in a recipient’s fecundity is given by:

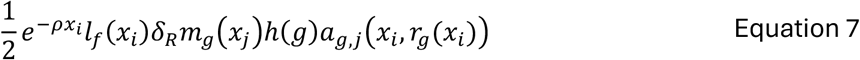

The expression for the donor’s indirect benefit from a change in a recipient’s survival is similar to the survival component of the donor, except we must account for age-specific interactions and sex-specific proportions:

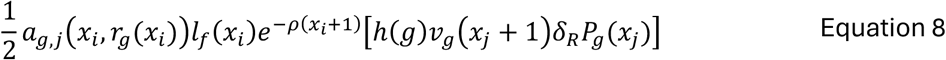

To capture the combined effects of changes in survival and fecundity, we sum Equations 7 and 8, and multiply by the relatedness of the donor to the recipient, *r*_g_(*x*_*i*_), as this gives the probability that the recipient carries the menopause allele. This results in:

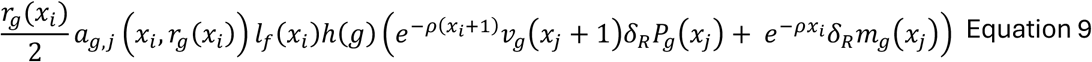

We then sum over all donor ages, all recipient ages, and both sexes to obtain the total number of alleles gained, given by Equation 10.

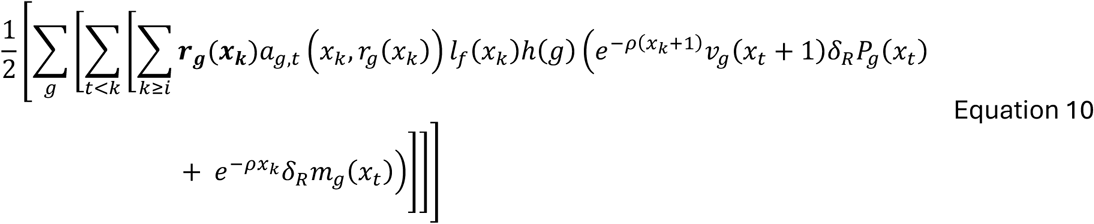

Finally, the total inclusive fitness effect of the menopause allele is obtained by summing Equations 10 and 6, resulting in:

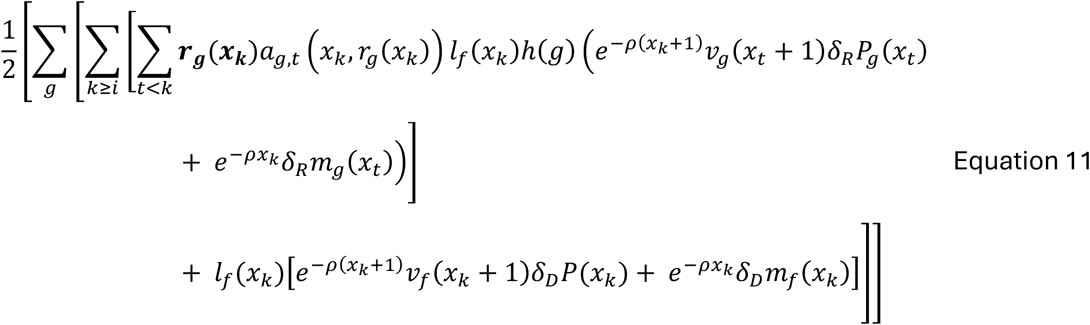

We denote this inclusive fitness effect as *λ*. If *λ* > 0, the rare menopause allele is expected to successfully invade the population^38^. An overview of all model parameters is in Table 1.

**Table 1:** Model Parameters.

| Expression | Value | Description |
| --- | --- | --- |
| $x_i, x_j$ | variable | Donor and recipient ages respectively. |
| $g$ | *female (f) or male (m)* | Recipient sex. |
| $r_g(x_i)$ | Determined by the model from Johnstone & Cant (2010) | Local relatedness coefficient of a female donor of age $x_i$ to her relatives of sex $g$. |
| $a_{g,j}(x_i, r_g(x_i))$ | $\frac{l_g(x_j)\rho^{-(x_j-1)}}{\sum_{k=1}^d l_g(x_k)\rho^{-(x_k-1)}}$ | Proportion of interactions of donor of age $x_i$ with recipient of age $x_j$ and sex $g$ (and thus relatedness $r_g(x_i)$ to the donor). Derived from sex-specific age distribution. |
| $l_g(x)$ | $\prod_{u=0}^{x-1} P_g(u)$ | Cumulative survival probability of sex $g$ from conception to age $x$. |
| $P_g(x)$ | Taken from a Leslie Matrix | Survival probability of sex $g$ from age $x$ to age $x + 1$. |
| $m_g(x)$ | Taken from a Leslie Matrix | Fecundity of sex $g$ at age $x$. |
| $e^{-\rho x}$ | — | Discount factor by growth rate and age $x$. The birth of a menopause allele carrier at time $x$ increases the frequency of the menopause allele by $\frac{1}{2N(x)}$, where $N(x)$ is the population size at time $y$. Following Rogers (1993), we weight each birth by $\frac{1}{2N(x)}$, equivalent to weighting by $e^{-\rho x}$ as we assume that the population is increasing exponentially at rate $\rho$. |
| $\rho$ | Leslie matrix leading eigenvalue | Population growth rate |
| $h(g)$ | $\begin{cases} \frac{1-c}{c} & \text{if } g = m \\ 1 & \text{if } g = f \end{cases}$ | Interaction weight of female donor with recipient of sex $g$, where $c$ is the fraction of males in the population (therefore females are $1 - c$). |
| $v_g(x)$ | $\frac{e^{\rho x}}{l_g(x)} \sum_{y=x} e^{-\rho y} k_g(y)$ | Reproductive value of sex $g$ at age $x$, relative to a value of unity for a new zygote. It's a measure for the contribution of an individual of a given |
| | $(x = 1, \dots, d)$ | age and sex to the future growth of the population. |
| $k_g(x)$ | $l_g(x)m_g(x)$ | Net offspring of sex $g$ in age $x$. |
| $\delta_A P_g(x)$ | $P_{g,mp}(x) - P_{g,no\ mp}(x)$ | Survival changes of recipient ($R$)/donor ($D$) from age $x$ to age $x + 1$ due to menopause.<br>$(A \in \{Donor\ (D), Recipient\ (R)\});$<br>no mp = no menopause, mp = menopause) |
| $\delta_A m_g(x)$ | $m_{g,mp}(x) - m_{g,no\ mp}(x)$ | Fecundity changes of recipient ($R$)/donor ($D$) in age $x$ due to menopause. |

We implemented this model under two alternative parameterizations (stop early and live long) to capture different life-history pathways to post-reproductive lifespan.

### (ii) Leslie matrices

The Leslie matrices show the fecundity (first row) and survival probabilities of individuals at different age classes (subdiagonal of each age class)^75^. Each matrix is sex-specific, but instead of using two separate Leslie matrices for males and females, we use one matrix that contains the values for both sexes. We write these matrices as 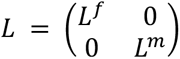, where *L*^*f*^ is the female Leslie *L^m^* matrix and *L*^*m*^ is the male Leslie matrix.

#### 1. Resident and Transient killer whales

The age classes for resident and transient killer whales are in 10-year intervals, from age 0 to 100. Values in the Leslie matrix are based on killer whales life-history data by Olesiuk et al., (2005) and Bigg et al. (1990) and thus portray a menopausal population with onset age 40-50 (Table S1)^76, 77^.

According to Ford et al. (2007), although few life history parameters are yet available for transient killer whales, values for resident killer whales presented in Olesiuk et al. (2005) may be generally representative^76,78^. This is also supported by Ford & Ellis (1999)^79^. Thus, survival and fecundity values of transient killer whales are assumed to be the same as those of resident killer whales.

#### 2. African elephants

The age classes in the matrix are in 5 year intervals, from age 0 to age 70.

The fecundity and survival values are sourced from the supplementary material provided by Chelliah et al. (2013) (Table S2)^80^.

#### 3. Sperm whales

The age classes in the matrix are in five year intervals, from age 0 to age 70.

Due to the lack of age-specific survivorship data for sperm whales, we use values from African elephants (*Loxodonta africana*) (Table S2), as the two species are reported to have comparable demographic parameters^81,82^.

#### 4. Asian elephants

The age classes in the matrix are in five-year intervals, from age 0 to age 65. Values are taken from Sukumar (1989) and Sukumar et al. (1998) (Table S3)^83,84^.

#### 5. Humans

The age classes in the matrix are in five year intervals, from age 0 to age 75, which was the available data age range (Table S4).

Survival and fecundity values for males and females were taken from Hill C Hurtado^85^, based on Aché hunter-gatherer data. The Aché provide a close analogy to ancestral human populations due to their demographics and social organization. Their hunting and foraging patterns reflect ancestral resource use^85^, while their fertility and mortality rates align with ancient populations^86^. Additionally, cooperative breeding and shared childcare resemble early human social behaviors^87^. Their relative isolation until recent contact further supports their relevance for studying ancient lifestyles.

#### 6. Amboseli baboons

Leslie matrices for baboons with one-year intervals between age classes (from age 0 to 27) were obtained from Kappeler & Pereira (2003) (Table S5)^45^.

#### 7. Chimpanzees

Chimpanzee life history patterns are exemplified by data from the Ngogo community, offering a setting with minimal human-induced mortality influences and incorporating several key factors essential for assessing the evolution of post-reproductive lifespans (Table S6).

Ngogo, located in Kibale National Park, Uganda, provides an exceptionally good habitat with favorable ecological conditions, resulting in higher population density and better overall health of the chimpanzees^67,69^. The community has also experienced relatively little anthropogenic disturbance and disease outbreaks compared to other chimpanzee populations^67^. As a result, Ngogo chimpanzees exhibit significantly higher survival rates than those observed in other wild populations, allowing researchers to observe individuals living to older ages and providing more comprehensive life history data^2^. Additionally, the Ngogo community is exceptionally large, offering a substantial sample size for demographic studies^69^.

Given these advantages, we used survival data of a total of 306 individuals from Wood et al. (2017), which is based on the Ngogo population, to construct Leslie matrices with five-year age intervals from birth to age 65^67^. For fecundity, we used values reported by Muller et al. (2020), which include data from three long-term study communities: Kanyawara in Kibale National Park, Uganda, as well as Mitumba and Kasekela in Gombe National Park, Tanzania^21^. While survival data at Ngogo differs from that of other chimpanzee populations, Wood et al. (2023) note that fertility patterns are consistent across populations, making the fecundity data from Muller et al. (2020) appropriate for use^2,21^.

### (iii) Modeling menopause pathways: stop early and live long

It is important to note that for non-menopausal species, empirical age-specific fecundity values inherently reflect male mating preferences. For menopausal species, empirical fecundity data are unavailable beyond reproductive cessation. In these cases, we extrapolated late-life fecundity values from comparable long-lived mammals such as African elephants and introduced an additional parameter allowing fecundity to decline at older ages, representing potential reproductive conflict in a hypothetical non-menopausal ancestor. The resulting competition factor was set to 1/1.67 for humans and killer whales (i.e. *m′_x_* = *m_x_*/1.67), following Croft et al. (2017), who reported that in killer whales, when mothers and daughters co-breed, the mortality hazard of calves from older-generation females is 1.7 times that of calves from younger females^6^. Model results remained qualitatively similar when this adjustment was omitted.

#### Stop early pathway

##### Menopausal species

To test the stop early hypothesis in menopausal species, we began with the empirical Leslie matrices described in the previous section. From these, we constructed a theoretical non-menopausal ancestral population by adding fecundity values at older ages, based on estimates from long-lived reproductive species such as elephants, to represent continued reproduction in the absence of menopause. We also removed all helping effects described in the “help provided by older females” sections below, resulting in decreased survival and fecundity of group members but extended reproduction for females. This ancestral population therefore represents a life-history strategy without menopause, optimized for prolonged reproduction but lacking the benefits of post-reproductive helping.

To simulate the introduction of the menopause allele, we iterated over each age class and designated it as the onset of menopause (*x*_*i*_). For the donor population, all fecundity values beyond *x*_*i*_ were set to zero. Because donors cannot simultaneously act as recipients, helping effects were excluded from this population. In some cases, later-life survival was adjusted upward to represent reduced risks associated with ceasing late reproduction^43^.

For the female recipient population, fecundity values were also set to zero after *x*_*i*_, because they carry the menopause-inducing allele with probability *r*_*f*_(*x*_*i*_). In contrast to donors, recipients do receive helping effects, with survival and fecundity values restored to their empirical levels to represent the assistance of post-reproductive females. Helping was restricted to younger age classes (*x* < *x*_*i*_), consistent with the assumption that older females aid only younger kin.

##### Non-menopausal species

For non-menopausal species, the empirical Leslie matrices already reflect the absence of menopause, and thus no ancestral reconstruction was required. Instead, we directly introduced the menopause allele by iterating over possible onset ages (*x*_*i*_) in the same manner as in the menopausal populations.

#### Live long pathway

##### Menopausal species

To test the live long hypothesis, we begin by modelling a theoretical non-menopausal ancestral population. In this baseline population, individuals do not survive beyond the last reproductive age class, *x*_*i*_. Post-reproductive survival is set to zero, and older females provide no help to kin, reducing the survival and fecundity of males and females by factors “lx_contribution” and “mx_contribution” (same parameters as in the stop early version), respectively, to represent the loss of assistance from post-reproductive females. According to the disposable soma theory^40,41^, in the absence of investment in somatic maintenance required for longevity, more resources can be allocated toward survival and fecundity during the reproductive years. Therefore, we increase the survival and fecundity of reproductive-age individuals by parameters “female_surv_tradeoff” and “female_fec_tradeoff” respectively, to reflect this resource allocation strategy.

This ancestral population represents a life history strategy optimized for early-life performance, with no investment in extended lifespan. To simulate the evolutionary emergence of longevity, we incrementally add post-reproductive age classes. For each added class, we proportionally cancel the early-life performance benefit, consistent with the trade-offs predicted by the disposable soma theory, so that energy devoted to longevity comes at the cost of reduced reproductive output or survival earlier in life. According to the disposable soma theory^40,41^, investing energy in somatic maintenance to support longevity comes at a cost to early-life survival and reproduction. Therefore, for each added post-reproductive age class, the early-life performance boost (i.e., the values of *female_surv_tradeoff* and *female_fec_tradeoff*) is proportionally reduced. This reflects the idea that when a longer post-reproductive lifespan is modelled, the cost of maintaining that lifespan can be distributed over more years. Specifically, as longevity increases, less energy remains available for early-life survival and reproduction.

Our model considers the evolution of a post-reproductive lifespan as a process involving two distinct roles, donors and recipients. We construct separate Leslie matrices for each, under the assumption that individuals cannot simultaneously act as both. For the recipient population, we restore survival and fecundity to the empirical values observed in the original Leslie matrix. This reflects both the return of assistance from post-reproductive donors and the cancellation of the earlier performance boost associated with shorter lifespan. This is because recipients carry the menopause-inducing allele with probability *r*_g_(*x*).

For the donor population (females only), survival and fecundity remain reduced by the lx_contribution and mx_contribution parameters, as donors are assumed not to assist each other. However, donors do survive into older age classes, and we cancel the reproductive and survival trade-offs from the ancestral model to represent the cost of increased longevity—consistent with the energy reallocation demands of extended somatic maintenance.

##### Non-menopausal species

To test the live long hypothesis in currently non-menopausal species, we followed a process similar to that used for menopausal species, with one major difference. In these populations, females reproduce only up to a species-specific age, *x*_*i*_, but afterwards they either die or live only briefly. As a result, empirical survival probabilities drop to zero beyond *x*_*i*_, and the original Leslie matrices include no non-zero survival entries after this point. This absence makes it impossible to directly add post-reproductive age classes using the observed data.

To simulate extended lifespan scenarios for comparison, we extrapolated survival beyond the peak reproductive period using an exponential decay model. For each species, we estimated a specific decay rate by fitting a log-linear regression to the declining portion of the empirical survival curve:

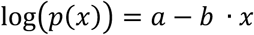

Where *p*(*x*) is the survival from age class *x* to age class *x* + 1, *a* is the intercept of the linear regression, representing the (log-transformed) survival probability at age zero within the range we’re fitting, and *b* is the estimated exponential decay rate. This regression was applied only to the segment of the survival curve where decline had already begun, omitting flat regions where survival remained constant due to data limitations.

We then used the estimated decay rate to extrapolate survival and relatedness to group members into added age classes beyond the observed range, simulating biologically plausible post-reproductive lifespans. These survival extrapolated values were incorporated into a new "extended" Leslie matrix for each species. Once the matrices included post-reproductive age classes, we proceeded with the same analytical steps as for the menopausal species.

#### Parameter Selection and Visualization

Following the exploration of the parameter space to determine the conditions under which a positive *λ* could be achieved for each species (Figure S6), we selected representative trade-off parameter sets to illustrate how *λ* varies with increasing post-reproductive lifespan. These parameter values are provided in Supplementary Table 7 (Table S7), where each value corresponds to an additional post-reproductive age class.

In Figure S2, 0 on the x-axis indicates no added post-reproductive years, representing a scenario where the donor lives only up to age, the last reproductive age based on empirical data. For instance, in killer whales, reproduction typically ends around age 40, so adding 10 years on the x-axis corresponds to a 50-year lifespan, 20 years to a 60-year lifespan, and so on.

### (iv) Life-history parameters

We apply the relatedness model developed by Johnstone and Cant (2010) (Figure 1)^4^. For all species, adult mortality parameters used in the calculation of kinship dynamics, *μ*_*f*_, *μ*_*m*_ for females and males, respectively, are derived from the corresponding entries in the Leslie matrix. Specifically, mortality is defined as: *μ*_*g*_(*x*) = 1 − *P_g_*(*x*), where *g* is the sex and *P*_g_(*x*) is the probability of surviving from age *x* to *x* + 1. We set the number of breeders per group to *n*_*f*_ = *n*_*m*_ = 3, where individuals are directly influenced by the reproductive behavior of a single female. The qualitative patterns seen in Figure 1, however, are unaffected by the number of breeders per group. Each column in the Leslie matrices represents an age class, with the length of the interval varying by species. Accordingly, we extract relatedness values at matching intervals for each species to ensure consistency between the demographic and kinship models.

#### 1. Resident killer whales

##### Dispersal, mating, kinship dynamics

Resident killer whales exhibit bisexual philopatry, meaning both sons and daughters tend to stay their entire lives within their natal social group, which is their mothers’ groups. Bisexual philopatry results in the formation of multi-generational matrilineal pods^77,88^. Therefore, we set the female and male dispersal parameters, *d*_*f*_, *d*_*m*_, respectively, to 0.1. That is, resident killer whales disperse to other pods with a probability of 0.1 ^4^. Moreover, because resident killer whales tend to mate by outbreeding^89^, we set the inbreeding rate, *m* (the fraction of young produced in a group and fathered by males within that group), to 0. We find that the relatedness of a donor to her group increases with her age, as in Croft et al. (2017) and Johnstone and Cant (2010)^4, 6^.

To examine how these kinship dynamics influence the evolution of menopause, we varied dispersal (*d*_*f*_, *d*_*m*_) and local mating (*m*) while holding the direct fitness costs and benefits of helping fixed (Figure S4). This allowed us to isolate how changes in relatedness alone affect the invasion criterion, λ. Our results confirm that the selective advantage of menopause is highly sensitive to the kinship environment. As expected from kin-selection theory, increasing the dispersal of either sex lowers a donor’s relatedness to her pod, which reduces λ and makes menopause less favored by selection (Figure S4A). In contrast, increasing the inbreeding rate raises group relatedness and increases λ, strongly favoring menopause (Figure S4B, S4C).

While high inbreeding with minimal dispersal creates the strongest selective pressure for menopause, we note that such conditions would also elevate inbreeding depression, which is likely maladaptive^90^. The resident killer whale parameters (indicated by the black star in Figure S4) fall within a "sweet spot" where λ remains positive, demonstrating that the observed bisexual philopatry and outcrossing patterns are sufficient to drive the evolution of menopause even without inbreeding.

##### Help provided by older females

Studies have shown that adult killer whales derive significant survival benefits from the presence of their mothers. According to Foster et al. (2012), the mortality risk for adult daughters increases by up to 2.7 times, and for adult sons by up to 8.7 times, in the year following their mother’s death^9^. However, since these values reflect the effects of maternal death rather than the reduced helping behavior of a living but reproductive mother, we adjusted the expected increase in offspring mortality in our non-menopausal population to more modest levels. Additionally, it is known that orca mothers face higher reproductive costs when supporting sons compared to daughters^37^. Given that resident killer whale males mate outside their natal group, preventing resource competition within the matriline, we assigned higher survival and fecundity benefits to male recipients compared to female recipients.

Furthermore, killer whale mothers and grandmothers have been observed to provide more assistance to their relatives’ survival than to their reproduction^10^. Therefore, we assume a stronger positive effect of menopause on the survival of recipients than their fecundity.

Based on these observations, we used the following life history parameter values. We assumed that female killer whales who continue reproducing beyond the onset age *x*_*i*_ experience a decrease in survival, which we set at 97% of the original survival rate. When reproduction cessation did not improve survival, similar results were observed. In the absence of menopause, we modelled female would be recipients as having reduced fecundity (95% of the original values) and reduced survival (80% of the original values). For male recipients, the absence of menopause reduced fecundity to 75% and survival to 50% of the original values. Due to the lack of fecundity estimates for post-menopausal female killer whales, we used values from a long-lived, continuously reproducing species – Asian elephants – which are similar to those of young female killer whales. However, Croft et al. (2017) showed that young females have a reproductive advantage over older females during overlap, indicating reproductive conflict^6^. To account for this, we adjusted the fecundity values by applying a competition factor of 1.67 ^6^.

From these changes in life history values due to the presence of the menopause allele, we constructed a new Leslie matrix reflecting the theoretical life history the non-menopausal population. By comparing this theoretical matrix with the original population matrix, we calculated the necessary differences for Equation 11.

To ensure our conclusions were not dependent on these specific point estimates, we systematically varied the strength of post-reproductive effects on the survival and fecundity of younger males and females (Figure S5). We found that while the absolute magnitude of λ varies with these parameters, the sign of λ remains consistently positive across a broad range of biologically plausible values, indicating that the selection for menopause is robust to specific parameterization of "help" in killer whales. Specifically, increasing help to male survival produces a marked increase in λ, whereas help to male fecundity has a more modest effect. This suggests that survival benefits, which accumulate across the recipient’s remaining lifespan, have a stronger impact on the evolution of menopause than age-specific fecundity gains. While the relationship between help and λ was generally monotonic, we observed that when help to female survival increased to extreme values, λ rose initially but eventually declined (Figure S5). This non-monotonicity arises from the quadratic term in the survival function, which feeds back on both donors and recipients. Despite these fluctuations in magnitude, the qualitative result, that post-reproductive helping behavior drives selection for the menopause allele, remains stable.

#### 2. Transient killer whales

##### Dispersal, mating, kinship dynamics

Dispersal and mating-wise, transient killer whales have higher dispersal rates than residents. Thus, we set *d*_*f*_ = *d*_*m*_ = 0.35, and *m* = 0.2, as in Nielsen et al.^7^.

##### Help provided by older females

Same as in resident killer whales.

#### 3. Sperm whales

##### Dispersal, mating, kinship dynamics

There is considerable variation in the age at which males disperse from their family units and begin their progressive relocation to cooler waters, but the mean seems to be about 6 years of age^82,91,92^. Males gradually become sexually mature in their teens, but do not seem to take an active role in breeding by entering the warm-water breeding grounds until their late twenties^82^.

Like elephants, females and their young live in matriarchal groups called pods, while bulls live apart. Bulls sometimes form loose bachelor groups with other males of similar age and size. As they grow older, they typically live solitary lives, only returning to the pod to socialize or to breed. Studies in various ocean basins have documented mature males returning to the same areas over the years^92–94^. Across a 2010–2017 Azores study, most mature males were seen in only a single year^38, 41^, two were seen in two years, and one male was re-encountered across eight years, making it a rare case of a persistently resighted male in this region^94^. Thus, we will use the following dispersal parameters for sperm whales: *d*_*f*_ = 0.15; *d*_*m*_ = 0.995. As for mating, it seems there is mainly outbreeding—in warm water, males rove between groups of females and associate with specific females over a few days to weeks^93,95,96^. The same males are rarely re-sighted with the same group of females or in the same breeding ground over different years^93,96,97^. Since there are no resources for the rate of inbreeding, we take *m* = 0.01, assuming a random encounter of a male with his natal female group is very rare.

##### Help provided by older females

The values applied to sperm whales are the same as those that will be presented to African elephants, due to their similar life histories and the lack of specific estimates in the literature^81,82^.

#### 4. African elephants

##### Dispersal, mating, kinship dynamics

African elephants have a matriarchal social structure, with adult females and their offspring living in herds^98,99^. Adult male elephants typically leave their natal family group at around 14 years of age while the females remain in their natal group^83,99^. This separation promotes reproduction by outbreeding^5,83^. Thus, we take demography parameters that are identical to sperm whales: *d*_*f*_ = 0.15, *d*_*m*_ = 0.995, *m* = 0.01. This results in a relatedness plot that is identical to that of sperm whales.

##### Help provided by older females

Parker et al. (2021) found that orphaned African elephants have lower survival rates compared to non-orphaned elephants^99^. Specifically, the survival probability for orphaned elephants aged 3-8 is 0.86, while for non-orphaned elephants it is 0.965. For elephants aged 9-18, the survival probability is 0.936 for orphans and 0.966 for non-orphans. Beyond this age, being orphaned does not appear to significantly affect survival. Based on these findings, we calculate the reduction in survival without parental care as 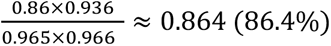. Since this figure reflects the complete absence of maternal care, we adjust it slightly higher, accounting for the benefits of social connections that mothers may still provide, regardless of menopause^91^. We therefore adjust the survival value to 0.864 × 1.12 ≈ 0.967, representing 96.7% survival in a theoretical menopausal African elephant population, compared to non-menopausal populations. As Parker et al. did not report sex-specific data, we assume this value applies to both sexes^99^.

Given that males typically leave the group between the ages of 4 and 21^98,100^, we assume the maximum age at which males can receive maternal care is 15. Additionally, as survival does not decline after age 18 in the absence of mothers, we assume 25 as the upper age limit for females for receiving maternal care.

In terms of fecundity, Parker et al. reported that orphaned elephants exhibit higher fertility than non-orphans between ages 9-18, with no effect beyond age 19^99^. This increased fertility may be due to reduced reproductive competition within the group. Since males leave the group before reaching reproductive maturity, we assume no change in male fecundity between menopausal and non-menopausal populations. However, since females remain in the group, we adjust their fecundity by dividing orphan fertility (0.093) by non-orphan fertility (0.082), yielding a change in fecundity due to theoretical menopause of 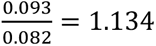. Given that this value represents conditions in the absence of maternal care, we retain it, as menopausal females would no longer reproduce.

#### 5. Asian elephants

##### Dispersal, mating, kinship dynamics

Males typically disperse between the ages of 11 and 20^101^, so we set *d*_*m*_ = 0.95. In contrast, females generally remain in their natal groups^102^, so we set *d*_*f*_ = 0.15. As males leave their natal groups and either live solitarily or join bachelor groups^103^, we assume non-local reproduction with *m* = 0.05.

##### Help provided by older females

Lahdenperä et al. found that Asian elephants whose grandmothers live outside the group, and were therefore unable to assist younger relatives, experienced reduced survival^49^. If the mothers of these elephants were younger than 20, their offspring’s survival up to age 5 was reduced by 32%. However, when the mothers were older than 20, the reduction in offspring survival was approximately 7%. To calculate the overall decrease in survival, we consider the age distribution and compute it as follows:

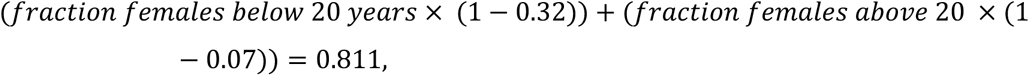

Where the fraction of females of specific age classes is taken from the age distribution calculated from the expression in row 4 in Table 1.

Since this value reflects the complete absence of maternal care, we adjust it slightly upwards to account for the remaining benefits that mothers might provide, even without menopause. Thus, we take 0.811 × 1.12 ≈ 0.91, so that in a theoretically menopausal Asian elephant population, survival is 91% of that in the real non-menopausal population. As Lahdenperä et al. did not provide sex-specific data, we assume this value applies to both sexes^49^. Given that males typically disperse between the ages of 11 and 20^101^, we assume the maximum age at which males can receive maternal care is 15.

In terms of fecundity, Lahdenperä et al. reported that mothers with the presence of grandmothers had shorter interbirth intervals (0.355 compared to 0.45), indicating higher fecundity^49^. We thus infer that a non-menopausal population has a fecundity that is 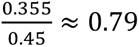 of a menopausal population. Since males leave the group before reaching reproductive maturity, we assume no change in male fecundity between menopausal and non-menopausal populations.

#### 6. Humans

##### Dispersal, mating, kinship dynamics

We assume ancestral female humans used to disperse from their natal group while males remained in them. Thus, we set *d*_*f*_ = 0.95 and *d*_*m*_ = 0.05. Also, because we assume mating is local after dispersal, *m* = 1^104,105^.

##### Help provided by older females

According to Lahdenperä et al., the length of a woman’s post-reproductive lifespan has a significant positive effect on the number of grandchildren she produces in both Finnish and Canadian pre-industrial populations^47^. In each population, this relationship is comparable, equating to an increase of approximately two additional grandchildren for every ten years that a woman survives beyond age 50. From their findings, we can infer that the fecundity of individuals with deceased mothers is about 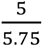 of those with living mothers. Additionally, they observed that mothers with dispersed offspring had, on average, 3.5 grandchildren, while mothers with local offspring had 4.7. This suggests that the absence of grandmothers results in 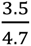 grandchildren, relative to when grandmothers are present. To obtain a general estimate, we take the average of these two findings: 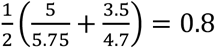

In terms of survival, their results indicate that only about 15% of offspring with deceased grandmothers reach age 15, while approximately 20% of individuals with grandmothers over age 60 survive to age 15, and around 30% with grandmothers under age 60 survive to this age. To calculate the survival advantage provided by post-reproductive versus reproductive grandmothers, we use the ratio: 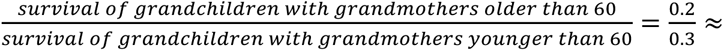 0.67.

Due to the lack of sex-specific values in the literature, we assume these estimates apply equally to male and female recipients, with the maximum age at which females can receive help set at 20-25 years, at which point they disperse.

#### 7. Amboseli baboons

##### Dispersal, mating, kinship dynamics

Males disperse while females are philopatric^106^. Male dispersal occurs when they reach the age of 7, meaning they can be assisted by older females only up to that age. Male baboons only start reproducing at age 7 so most males do not reproduce while they are around their mothers. However, there is a large fraction of males that do manage to reproduce in their natal group before dispersing. According to Alberts C Altmann^107^, 50% of males leave the natal group *after* mating, and almost all males leave the group after they reach age 13^108^. Thus, we set *m*, the inbreeding parameter, to be

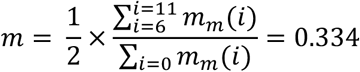

where *m*_*m*_(*x*) is the fecundity value for males of age class x. For dispersal, we use *d*_*f*_ = 0.15 and *d*_*m*_ = 0.95.

##### Help provided by older females

Zipple et al. demonstrated that early-life adversity experienced by female baboons leads to elevated mortality in their offspring, but this effect is both age and context dependent^109^. Since these effects are limited to early life and depend on the mother’s reduced ability to provide care, and based on other species parameters, we model only a modest reduction of 2% and 3% in female and male offspring survival respectively, in populations lacking menopause.

According to Alberts & Altmann (1995), maternal traits—particularly maternal rank—strongly influence the timing of male reproductive maturation, including earlier testicular enlargement and attainment of adult dominance rank^107^. These maternal effects prepare males for reproduction, but actual fecundity depends on current social-demographic factors, such as dominance status and female availability. Given the absence of strong fecundity effects in adulthood and the male-biased nature of the benefits, we conservatively assumed a 3% reduction in male fecundity and a 1% reduction in female fecundity in non-menopausal populations. Additionally, we assumed that male offspring cease receiving maternal assistance after approximately age 9 due to dispersal.

#### 8. Chimpanzees

##### Dispersal, mating, kinship dynamics

Chimpanzees typically exhibit female-biased dispersal, which promotes outbreeding. Most females leave their natal communities before breeding, with over 90% of females dispersing at some sites. Male chimpanzees generally exhibit philopatry, meaning they remain in their natal communities^23^. Thus, we take female dispersal rate (*d*_*f*_) to be 90% and male dispersal rate (*d*_*m*_) to be 0%.

Female chimpanzees typically avoid inbreeding^110^. Since they usually disperse to a new group after reaching sexual maturity^23^, their reproduction is mostly within that new group, and we refer to that as inbreeding. Thus, we take the inbreeding rate to be 100%.

##### Help provided by older females

Studies on different chimpanzee communities highlight the significant impact of maternal presence on offspring survival and, to a lesser extent, fecundity. Nakamura et al. reported that in the Mahale chimpanzee community, males who lost their mothers before the age of five faced a 4.07-fold increase in mortality risk, while those orphaned between ages five and thirteen experienced a 2.38-fold increase^34^. However, males who lost their mothers after age thirteen did not show a significant increase in mortality risk. Similarly, Stanton et al. found that in the Gombe chimpanzee community, males orphaned at any age between 0 and 14.99 years had significantly lower survival rates than non-orphans and died earlier than expected^36^. Females also experienced reduced survival when orphaned before age ten, but maternal loss between 10 and 14.99 years did not significantly impact their mortality risk.

Beyond immediate survival, Samuni et al. observed in the Taï Forest chimpanzee communities that maternal investment extends beyond lactation into early adulthood, positively influencing offspring’s physical development^35^. Data from the same population suggest that early maternal loss negatively impacts the reproductive success of sons^33^, although this relationship appears weaker in other chimpanzee populations^111^.

Overall, the literature supports the idea that maternal presence plays a crucial role in juvenile survival and development, particularly for males. The reduced impact on females is likely due to their early dispersal upon reaching sexual maturity^23^, which limits the window during which they receive maternal care. Based on these findings, we assume that maternal care extends up to ages 10–15 for females.

Given this dispersal pattern, we assume that female fecundity does not differ between those with menopausal and non-menopausal mothers or grandmothers. For males, we expect a rather small fecundity advantage in populations with menopausal females, as the literature presents conflicting evidence. Given the similarly small differences observed in baboons, we apply a 5% improvement in male fecundity for populations with menopausal females.

Regarding survival, evidence suggests that non-orphans experience the strongest survival benefits at younger ages^34^. However, research on Sebitoli chimpanzees suggests that environmental stressors like pesticide exposure can increase early-life vulnerability, potentially amplifying the importance of maternal care for offspring survival^112^. Because chimpanzees in the broader Mahale region are also exposed to environmental stressors such as habitat degradation and fragmentation due to human activities activities^113^, we adopt conservative survival-effect sizes to avoid overestimating survival benefits. Specifically, we use values closer to those observed in Asian elephants and assume that, at each age class, male offspring without post-reproductive care have 92% of the survival probability of males with post-reproductive care. Because females disperse earlier and receive less prolonged maternal support, we assume a smaller effect for daughters: females without post-reproductive care have 95% of the survival probability of females with it.

### (v) Post-reproductive representation (PrR) calculation

The PrR is the proportion of an individual’s adult lifespan that is spent in the post-reproductive phase. We calculate it following Levitis C Lackey^114^, using life-tables,

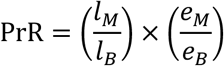

Here *t*_*B*_ and *t*_*M*_ are survival probabilities at ages *B* (the onset of reproductive lifespan) and *M* (the end of reproductive lifespan), respectively; and *e*_*B*_ and *e*_*M*_ are the remaining life expectancies at ages *B* and *M*, respectively.

The term 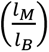 captures the fraction of individuals who survive from age *B* to age *M*, reflecting survival into the post-reproductive phase. The term 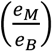 reflects the remaining life expectancy at age *M* relative to that at age *B*, indicating the longevity of the post-reproductive phase compared to the total adult lifespan. Since most of the life tables we found do not contain the life expectancy values, we calculate them using the life table approach^115^.

From the survival probabilities, *l*(*x*), we calculate *T*(*x*), the cumulative sum of all future person-years lived from age x onward:

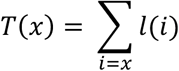

Then, life expectancy at age x, denoted as *e*_*x*_, is calculated as:

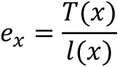

This gives the average number of years remaining for an individual who has survived to age x.

The PrR predicted by our model are based on the value of *M* determined by the model – the end of the reproductive lifespan or the beginning of the post-reproductive lifespan – defined as the age at which *λ* > 0. These predicted values are compared to PrR values reported in the literature (see text for description), as summarized in Table 2.

**Table 2:**
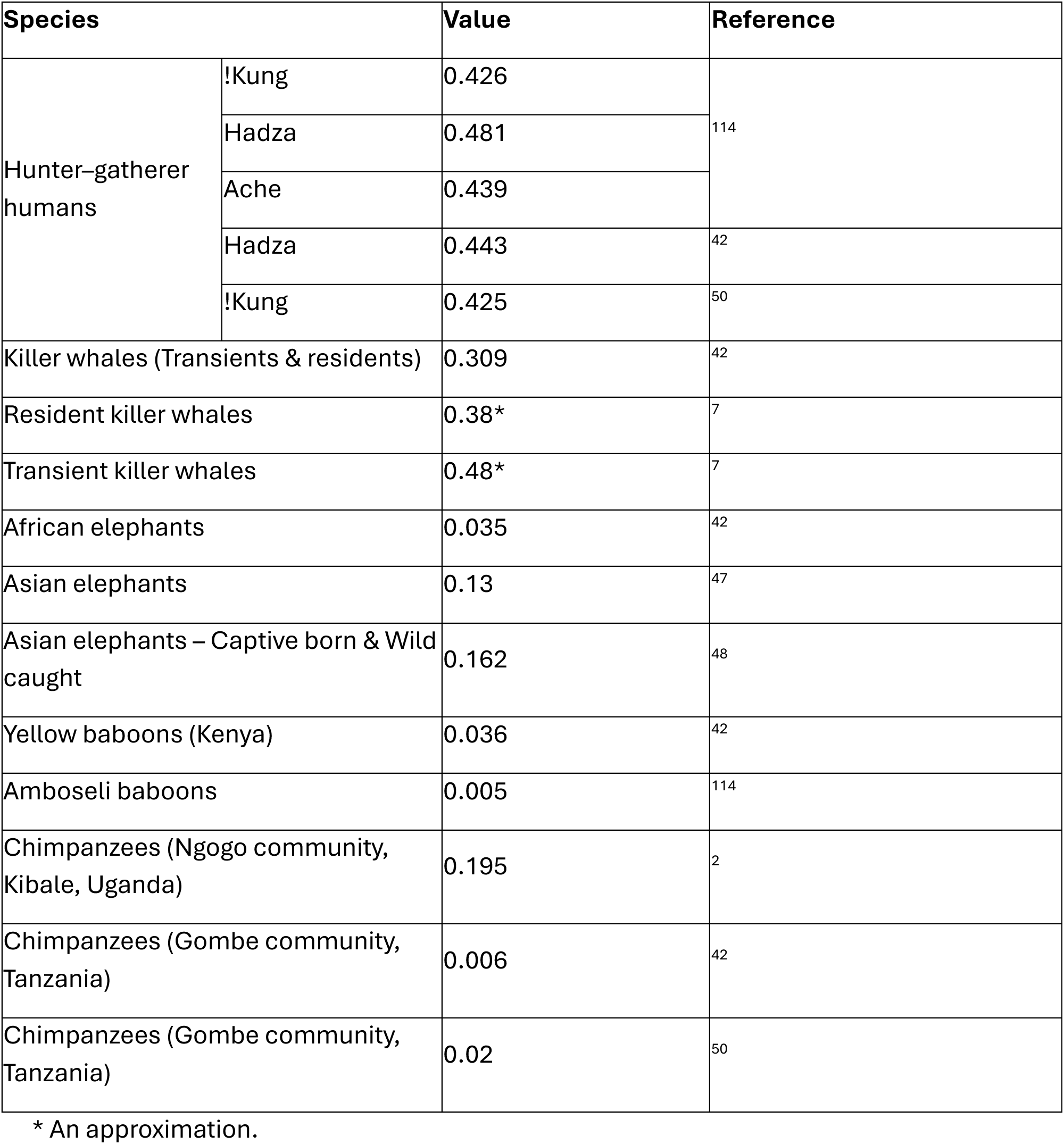
Empirical PrR values.

### (vi) Sobol sensitivity analysis

Given the complexity of the model and its reliance on help parameters, Sobol sensitivity analysis was performed to assess the relative importance of model parameters and their interactions on model output (λ)^57^. This variance-based global sensitivity analysis technique decomposes the output variance into contributions from individual parameters (first-order effects) and their interactions (higher-order effects). The first-order Sobol index was used to measure the direct effect of varying a single parameter, while the total Sobol index captured both the main effect and interactions with other parameters, providing an overall sensitivity measure.

The parameters analysed included the help in survival (*t*_*f*_O*x*_j_P, *t*_*m*_O*x*_j_P) and fecundity (*m*_*f*_O*x*_j_P, *m*_*m*_O*x*_j_P) provided to females and males respectively, as well as the effect of changes in maternal mortality due to continued reproduction at later ages. All other parameters were held constant, reflecting the specific conditions of the killer whale case. The analysis was implemented using the SALib Python library^58,59^, with a total of 10,000 samples generated using Sobol’s quasi-random sequence.

## REFERENCES

1. Ellis, S., Franks, D. W., Nielsen, M. L. K., Weiss, M. N. & Croft, D. P. The evolution of menopause in toothed whales. Nature 627, 579–585 (2024).

2. Wood, B. M. et al. Demographic and hormonal evidence for menopause in wild chimpanzees. Science 382, eadd5473 (2023).

3. Laisk, T. et al. Demographic and evolutionary trends in ovarian function and aging. Human Reproduction Update https://doi.org/10.1093/humupd/dmy031 (2018) doi:10.1093/humupd/dmy031.

4. Johnstone, R. A. & Cant, M. A. The evolution of menopause in cetaceans and humans: the role of demography. Proc. R. Soc. B. 277, 3765–3771 (2010).

5. Lawson Handley, L. J. & Perrin, N. Advances in our understanding of mammalian sex-biased dispersal. Molecular Ecology 16, 1559–1578 (2007).

6. Croft, D. P. et al. Reproductive Conflict and the Evolution of Menopause in Killer Whales. Current Biology 27, 298–304 (2017).

7. Nielsen, M. L. K. et al. A long postreproductive life span is a shared trait among genetically distinct killer whale populations. Ecology and Evolution 11, 9123–9136 (2021).

8. Brent, L. J. N. et al. Ecological Knowledge, Leadership, and the Evolution of Menopause in Killer Whales. Current Biology 25, 746–750 (2015).

9. Foster, E. A. et al. Adaptive Prolonged Postreproductive Life Span in Killer Whales. Science 337, 1313–1313 (2012).

10. Nattrass, S., et al. Postreproductive killer whale grandmothers improve the survival of their grandoffspring. Proc. Natl. Acad. Sci. U.S.A. 116, 26669–26673 (2019).

11. Wright, B. M., Stredulinsky, E. H., Ellis, G. M. & Ford, J. K. B. Kin-directed food sharing promotes lifetime natal philopatry of both sexes in a population of fish-eating killer whales, Orcinus orca. Animal Behaviour 115, 81–95 (2016).

12. Kasuya, T. Life history and reproductive biology of the short-finned pilot whale, Globicephala macrorhynchus, off the Pacific coast of Japan. Rep. int. Whal. Commn. 259–310 (1984).

13. Colbeck, G. J. et al. Groups of related belugas (*Delphinapterus leucas*) travel together during their seasonal migrations in and around Hudson Bay. Proc. R. Soc. B. 280, 20122552 (2013).

14. Marcoux, M., Auger-Méthé, M. & Humphries, M. M. Encounter frequencies and grouping patterns of narwhals in Koluktoo Bay, Baffin Island. Polar Biol 32, 1705–1716 (2009).

15. Martien, K. et al. Fidelity to natal social groups and mating within and between social groups in an endangered false killer whale population. Endang. Species. Res. 40, 219–230 (2019).

16. O’Corry-Crowe, G. et al. Data from: Migratory culture, population structure and stock identity in North Pacific beluga whales (Delphinapterus leucas). 54657 bytes Dryad 10.5061/DRYAD.6B70G11 (2019).

17. Palsbøl, P. J. Population structure and seasonal movements of narwhals, Monodon monoceros, determined from mtDNA analysis. Nature 78, 284–292 (1997).

18. Eriksson, J. et al. Y-chromosome analysis confirms highly sex-biased dispersal and suggests a low male effective population size in bonobos (*Pan paniscus*). Molecular Ecology 15, 939–949 (2006).

19. Jensen-Seaman, M. I., Deinard, A. S. & Kidd, K. K. Modern African Ape Populations as Genetic and Demographic Models of the Last Common Ancestor of Humans, Chimpanzees, and Gorillas. Journal of Heredity 92, 475–480 (2001).

20. Langergraber, K. E. et al. The Genetic Signature of Sex-Biased Migration in Patrilocal Chimpanzees and Humans. PLoS ONE 2, e973 (2007).

21. Muller, M. N. et al. Sexual dimorphism in chimpanzee (Pan troglodytes schweinfurthii) and human age-specific fertility. Journal of Human Evolution 144, 102795 (2020).

22. Nishida, T. et al. Demography, female life history, and reproductive profiles among the chimpanzees of Mahale. American J Primatol 59, 99–121 (2003).

23. Pusey, A. E. & Schroepfer-Walker, K. Female competition in chimpanzees. Phil. Trans. R. Soc. B 368, 20130077 (2013).

24. Hammer, M. F. et al. Hierarchical Patterns of Global Human Y-Chromosome Diversity. Molecular Biology and Evolution 18, 1189–1203 (2001).

25. Ségurel, L. et al. Sex-Specific Genetic Structure and Social Organization in Central Asia: Insights from a Multi-Locus Study. PLoS Genet 4, e1000200 (2008).

26. Seielstad, M. T., Minch, E. & Cavalli-Sforza, L. L. Genetic evidence for a higher female migration rate in humans. Nat Genet 20, 278–280 (1998).

27. Wilder, J. A., Kingan, S. B., Mobasher, Z., Pilkington, M. M. & Hammer, M. F. Global patterns of human mitochondrial DNA and Y-chromosome structure are not influenced by higher migration rates of females versus males. Nat Genet 36, 1122–1125 (2004).

28. Alvarez, H. P. Residence Groups Among Hunter-Gatherers: A View of the Claims and Evidence for Patrilocal Bands. in Kinship and Behavior in Primates (eds Chapais, B. & Berman, C. M.) 420–442 (Oxford University Press New York, NY, 2004). doi:10.1093/oso/9780195148893.003.0018.

29. Ember, C. R. Myths about Hunter-Gatherers. Ethnology 17, 439 (1978).

30. Marlowe, F. W. Marital Residence among Foragers. Current Anthropology 45, 277–284 (2004).

31. Hawkes, K., O’Connell, J. F., Jones, N. G. B., Alvarez, H. & Charnov, E. L. Grandmothering, menopause, and the evolution of human life histories. Proc. Natl. Acad. Sci. U.S.A. 99, 1336–1339 (1998).

32. Sear, R. & Mace, R. Who keeps children alive? A review of the effects of kin on child survival. Evolution and Human Behavior 29, 1–18 (2008).

33. Crockford, C., Samuni, L., Vigilant, L. & Wittig, R. M. Postweaning maternal care increases male chimpanzee reproductive success. Sci. Adv. 6, eaaz5746 (2020).

34. Nakamura, M., Hayaki, H., Hosaka, K., Itoh, N. & Zamma, K. Brief Communication: Orphaned male Chimpanzees die young even after weaning. American J Phys Anthropol 153, 139–143 (2014).

35. Samuni, L. et al. Maternal effects on offspring growth indicate post-weaning juvenile dependence in chimpanzees (Pan troglodytes verus). Front Zool 17, 1 (2020).

36. Stanton, M. A., Lonsdorf, E. V., Murray, C. M. & Pusey, A. E. Consequences of maternal loss before and after weaning in male and female wild chimpanzees. Behav Ecol Sociobiol 74, 22 (2020).

37. Weiss, M. N. et al. Costly lifetime maternal investment in killer whales. Current Biology 33, 744–748.e3 (2023).

38. Charlesworth, B. & Charnov, E. L. Kin selection in age-structured populations. Journal of Theoretical Biology 88, 103–119 (1981).

39. Hamilton, W. D. The genetical evolution of social behaviour. I. Journal of Theoretical Biology 7, 1–16 (1964).

40. Kirkwood, T. B. L. & Rose, M. R. Evolution of senescence: late survival sacrificed for reproduction. Philosophical Transactions of the Royal Society of London. Series B: Biological Sciences 332, 15–24 (1991).

41. Shanley, D. P. & Kirkwood, T. B. L. Evolution of the human menopause. Bioessays 23, 282–287 (2001).

42. Ellis, S. et al. Postreproductive lifespans are rare in mammals. Ecology and Evolution 8, 2482–2494 (2018).

43. Pavard, S., E. Metcalf, C. J. & Heyer, E. Senescence of reproduction may explain adaptive menopause in humans: A test of the “mother” hypothesis. American J Phys Anthropol 136, 194–203 (2008).

44. Ellis, S. et al. Analyses of ovarian activity reveal repeated evolution of post-reproductive lifespans in toothed whales. Sci Rep 8, 12833 (2018).

45. Kappeler, P. M. & Pereira, M. E. Primate Life Histories and Socioecology. (University of Chicago Press, 2003).

46. Frankham, R. CONSERVATION GENETICS. Annu. Rev. Genet. 29, 305–327 (1995).

47. Lahdenperä, M., Mar, K. U. & Lummaa, V. Reproductive cessation and post-reproductive lifespan in Asian elephants and pre-industrial humans. Front Zool 11, 54 (2014).

48. Chapman, S. N., Jackson, J., Htut, W., Lummaa, V. & Lahdenperä, M. Asian elephants exhibit post-reproductive lifespans. BMC Evol Biol 19, 193 (2019).

49. Lahdenperä, M., Mar, K. U. & Lummaa, V. Nearby grandmother enhances calf survival and reproduction in Asian elephants. Sci Rep 6, 27213 (2016).

50. Alberts, S. C., et al. Reproductive aging patterns in primates reveal that humans are distinct. Proc. Natl. Acad. Sci. U.S.A. 110, 13440–13445 (2013).

51. Bronikowski, A. M. et al. Female and male life tables for seven wild primate species. Sci Data 3, 160006 (2016).

52. Nishida, T. C Hiraiwa-Hasegawa, M. Chimpanzees and Bonobos: Cooperative Relationships among Males. Primate Societies 165–178 (1986) 10.7208/9780226220468-017.

53. Herndon, J. G. et al. Menopause occurs late in life in the captive chimpanzee (Pan troglodytes). AGE 34, 1145–1156 (2012).

54. Jones, J. H., Wilson, M. L., Murray, C. & Pusey, A. Phenotypic quality influences fertility in Gombe chimpanzees. Journal of Animal Ecology 79, 1262–1269 (2010).

55. Pruetz, J. Apes on the Edge: Chimpanzee Life on the West African Savanna. (University of Chicago Press, 2025).

56. Nielsen, M. L. K. et al. Temporal dynamics of mother–offspring relationships in Bigg’s killer whales: opportunities for kin-directed help by post-reproductive females. Proc. R. Soc. B. 290, 20230139 (2023).

57. Sobol, I. M. Global sensitivity indices for nonlinear mathematical models and their Monte Carlo estimates. Mathematics and Computers in Simulation 55, 271–280 (2001).

58. Herman, J. & Usher, W. SALib: An open-source Python library for Sensitivity Analysis. JOSS 2, 97 (2017).

59. Iwanaga, T., Usher, W. & Herman, J. Toward SALib 2.0: Advancing the accessibility and interpretability of global sensitivity analyses. SESMO 4, 18155 (2022).

60. Hawkes, K. How grandmother effects plus individual variation in frailty shape fertility and mortality: Guidance from human–chimpanzee comparisons. Proc. Natl. Acad. Sci. U.S.A. 107, 8977–8984 (2010).

61. Atsalis, S. & Margulis, S. W. Sexual and Hormonal Cycles in Geriatric Gorilla gorilla gorilla. Int J Primatol 27, 1663–1687 (2006).

62. Kohama, S. G. & Urbanski, H. F. The aged female rhesus macaque as a translational model for human menopause and hormone therapy. Hormones and Behavior 166, 105658 (2024).

63. Smit, N. & Robbins, M. M. Post-reproductive lifespan in wild mountain gorillas. Proc. Natl. Acad. Sci. U.S.A. 122, e2510998122 (2025).

64. Stokes, E. J., Parnell, R. J. & Olejniczak, C. Female dispersal and reproductive success in wild western lowland gorillas (Gorilla gorilla gorilla). Behavioral Ecology and Sociobiology 54, 329–339 (2003).

65. Robbins, A. M. et al. Mothers may shape the variations in social organization among gorillas. R. Soc. open sci. 3, 160533 (2016).

66. Pusey, A. E., Wilson, M. L. & Anthony Collins, D. Human impacts, disease risk, and population dynamics in the chimpanzees of Gombe National Park, Tanzania. American J Primatol 70, 738–744 (2008).

67. Wood, B. M., Watts, D. P., Mitani, J. C. & Langergraber, K. E. Favorable ecological circumstances promote life expectancy in chimpanzees similar to that of human hunter-gatherers. Journal of Human Evolution 105, 41–56 (2017).

68. Muller, M. N. & Wrangham, R. W. Mortality rates among Kanyawara chimpanzees. Journal of Human Evolution 66, 107–114 (2014).

69. Watts, D. The Ngogo Chimpanzee Project. https://campuspress.yale.edu/ngogochimp/project/.

70. Helle, S., Lummaa, V. & Jokela, J. Sons Reduced Maternal Longevity in Preindustrial Humans. Science 296, 1085–1085 (2002).

71. Helle, S. & Lummaa, V. A trade-off between having many sons and shorter maternal post-reproductive survival in pre-industrial Finland. Biol. Lett. 9, 20130034 (2013).

72. Alfano, M. Daughters, dowries, deliveries: The effect of marital payments on fertility choices in India. Journal of Development Economics 125, 89–104 (2017).

73. Grafen, A. A theory of Fisher’s reproductive value. J. Math. Biol. 53, 15–60 (2006).

74. Otto, S. P. & Day, T. A Biologist’s Guide to Mathematical Modeling in Ecology and Evolution. (Princeton University Press, 2011). doi:10.2307/j.ctvcm4hnd.

75. Leslie, P. H. ON THE USE OF MATRICES IN CERTAIN POPULATION MATHEMATICS. Biometrika 33, 183–212 (1945).

76. Olesiuk, P. F., Ellis, G. M. & Ford, J. K. B. Cycle biologique et dynamique de la population des épaulards (Orcinus orca) résidents du nord de la Colombie-Britannique.

77. Bigg, M. A., Olesiuk, P. F., Ellis, G. M., Ford, J. K. B. & Balcomb, K. C. Social organization and genealogy of resident killer whales (Orcinus orca) in the coastal waters of British Columbia and Washington State. Report of the International Whaling Commission https://www.researchgate.net/profile/Sally-Mizroch/publication/291157559_Report_of_the_workshop_on_individual_recognition_and_the_estimation_of_cetacean_population_parameters/links/5807cdf008ae5ed04bfe7e78/Report-of-the-workshop-on-individual-recognition-and-the-estimation-of-cetacean-population-parameters.pdf#page=391 (1990).

78. Ford, J. K. B., Ellis, G. M., Durban, J. W., Department of Fisheries and Oceans, Ottawa, ON(Canada) & Canadian Science Advisory Secretariat, Ottawa, ON(Canada). An assessment of the potential for recovery of west coast transient killer whales using coastal waters of British Columbia. (2007).

79. Ford, J. K. B. & Ellis, G. M. *Transients: Mammal-Hunting Killer Whales of B.C., Washington State, and Southeast Alaska*. (University of British Columbia Press, 1999). doi:10.59962/9780774856300.

80. Chelliah, K., Bukka, H. & Sukumar, R. Modeling harvest rates and numbers from age and sex ratios: A demonstration for elephant populations. Biological Conservation 165, 54–61 (2013).

81. Eguiguren, A., Konrad Clarke, C. M. & Cantor, M. Sperm Whale Reproductive Strategies: Current Knowledge and Future Directions. in Sex in Cetaceans (eds Würsig, B. & Orbach, D. N.) 443–467 (Springer International Publishing, Cham, 2023). doi:10.1007/978-3-031-35651-3_19.

82. Whitehead, H. Sperm Whale. in Encyclopedia of Marine Mammals 919–925 (Elsevier, 2018). doi:10.1016/B978-0-12-804327-1.00242-9.

83. Sukumar, R. Ecology of the Asian elephant in southern India. I. Movement and habitat utilization patterns. J. Trop. Ecol. 5, 1–18 (1989).

84. Sukumar, R., Ramakrishnan, U. & Santosh, J. A. Impact of poaching on an Asian elephant population in Periyar, southern India: a model of demography and tusk harvest. Animal Conservation 1, 281–291 (1998).

85. Hill, K. R. & Hurtado, A. M. *Ache Life History: The Ecology and Demography of a Foraging People*. (Aldine de Gruyter, New York, NY, 1996).

86. Gurven, M. & Kaplan, H. Longevity Among Hunter-Gatherers: A Cross-Cultural Examination. Population & Development Rev 33, 321–365 (2007).

87. Hill, K. R. et al. Co-Residence Patterns in Hunter-Gatherer Societies Show Unique Human Social Structure. Science 331, 1286–1289 (2011).

88. Gowans, S., Würsig, B. & Karczmarski, L. The Social Structure and Strategies of Delphinids: Predictions Based on an Ecological Framework. in Advances in Marine Biology vol. 53 195–294 (Elsevier, 2007).

89. Barrett-Lennard, L. G. Population structure and mating patterns of Killer Whales (Orcinus orca) as revealed by DNA analysis. https://doi.org/10.14288/1.0099652 (2009) doi:10.14288/1.0099652.

90. Kardos, M. et al. Inbreeding depression explains killer whale population dynamics. Nat Ecol Evol 7, 675–686 (2023).

91. Gero, S., Gordon, J. & Whitehead, H. Calves as social hubs: dynamics of the social network within sperm whale units. Proc. R. Soc. B. 280, 20131113 (2013).

92. Girardet, J. et al. Long Distance Runners in the Marine Realm: New Insights Into Genetic Diversity, Kin Relationships and Social Fidelity of Indian Ocean Male Sperm Whales. Front. Mar. Sci. 9, 815684 (2022).

93. Gero, S. et al. Behavior and social structure of the sperm whales of Dominica, West Indies. Marine Mammal Science 30, 905–922 (2014).

94. Van Der Linde, M. L. & Eriksson, I. K. An assessment of sperm whale occurrence and social structure off São Miguel Island, Azores using fluke and dorsal identification photographs. Marine Mammal Science 36, 47–65 (2020).

95. Coakes, A. K. & Whitehead, H. Social structure and mating system of sperm whales off northern Chile. Can. J. Zool. 82, 1360–1369 (2004).

96. Whitehead, H. The behaviour of mature male sperm whales on the Galápagos Islands breeding grounds. Can. J. Zool. 71, 689–699 (1993).

97. Jaquet, N. & Gendron, D. The social organization of sperm whales in the Gulf of California and comparisons with other populations. J. Mar. Biol. Ass. 89, 975–983 (2009).

98. O’Connell-Rodwell, C. E. et al. Male African Elephant (Loxodonta africana) Behavioral Responses to Estrous Call Playbacks May Inform Conservation Management Tools. Animals 12, 1162 (2022).

99. Parker, J. M. et al. Poaching of African elephants indirectly decreases population growth through lowered orphan survival. Current Biology 31, 4156–4162.e5 (2021).

100. Sarano, F. et al. Nursing Behavior in Sperm Whales (Physeter macrocephalus). AB&C 10, 105–131 (2023).

101. Srinivasaiah, N., Kumar, V., Vaidyanathan, S., Sukumar, R. & Sinha, A. All-Male Groups in Asian Elephants: A Novel, Adaptive Social Strategy in Increasingly Anthropogenic Landscapes of Southern India. Sci Rep 9, 8678 (2019).

102. Lynch, E. C., Lummaa, V., Htut, W. & Lahdenperä, M. Evolutionary significance of maternal kinship in a long-lived mammal. Phil. Trans. R. Soc. B 374, 20180067 (2019).

103. Seltmann, M. W., Helle, S., Htut, W. & Lahdenperä, M. Males have more aggressive and less sociable personalities than females in semi-captive Asian elephants. Sci Rep 9, 2668 (2019).

104. Fortunato, L. Lineal kinship organization in cross-specific perspective. Phil. Trans. R. Soc. B 374, 20190005 (2019).

105. Fortunato, L. & Jordan, F. Your place or mine? A phylogenetic comparative analysis of marital residence in Indo-European and Austronesian societies. Phil. Trans. R. Soc. B 365, 3913–3922 (2010).

106. Onyango, P. O., Gesquiere, L. R., Altmann, J. & Alberts, S. C. Puberty and dispersal in a wild primate population. Hormones and Behavior 64, 240–249 (2013).

107. Alberts, S. C. & Altmann, J. Preparation and activation: determinants of age at reproductive maturity in male baboons. (1995).

108. Alberts, S. C. & Altmann, J. Balancing Costs and Opportunities: Dispersal in Male Baboons. The American Naturalist 145, 279–306 (1995).

109. Zipple, M. N., Archie, E. A., Tung, J., Altmann, J. & Alberts, S. C. Intergenerational effects of early adversity on survival in wild baboons. eLife 8, e47433 (2019).

110. White, L. C., Städele, V., Ramirez Amaya, S., Langergraber, K. & Vigilant, L. Female chimpanzees avoid inbreeding even in the presence of substantial bisexual philopatry. R. Soc. Open Sci. 11, 230967 (2024).

111. Surbeck, M. et al. Males with a mother living in their group have higher paternity success in bonobos but not chimpanzees. Current Biology 29, R354–R355 (2019).

112. Krief, S. et al. Agricultural expansion as risk to endangered wildlife: Pesticide exposure in wild chimpanzees and baboons displaying facial dysplasia. Science of The Total Environment 598, 647–656 (2017).

113. Chitayat, A. B., Wich, S. A., Lewis, M., Stewart, F. A. & Piel, A. K. Ecological correlates of chimpanzee (Pan troglodytes schweinfurthii) density in Mahale Mountains National Park, Tanzania. PLoS ONE 16, e0246628 (2021).

114. Levitis, D. A. & Lackey, L. B. A measure for describing and comparing postreproductive life span as a population trait. Methods Ecol Evol 2, 446–453 (2011).

115. Keyfitz, N. & Caswell, H. *Applied Mathematical Demography*. (Springer-Verlag, New York, 2005). doi:10.1007/b139042.

