## Supplementary Figures and Tables for "Evolution of Menopause and the Mama’s Boy Hypothesis"

### SUPPLEMENTARY MATERIAL

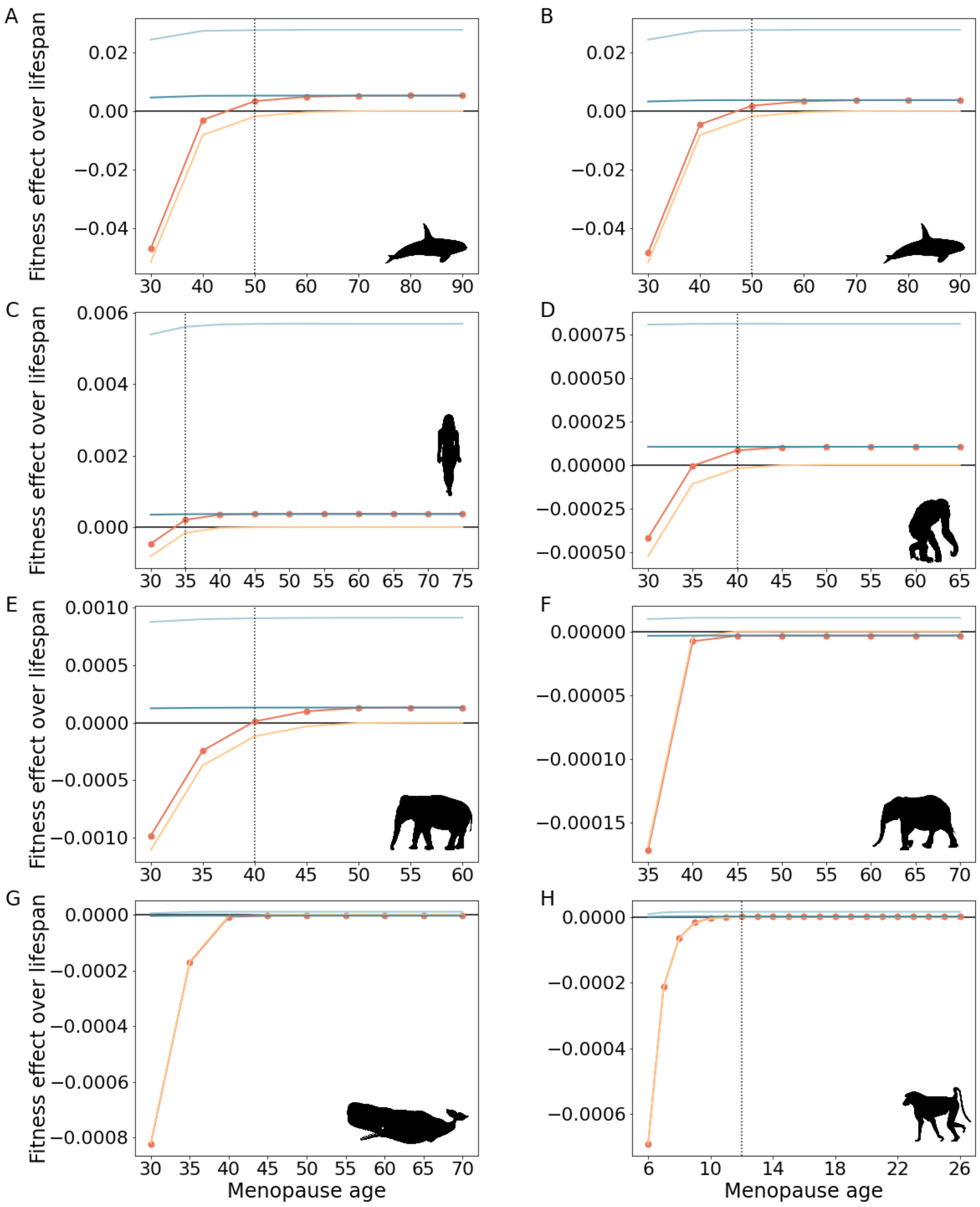


Figure S1: **$\lambda$, cost , benefit and relatedness-weighted benefit values for different menopause onset ages, under the stop early pathway for the evolution of menopause**. The x-axis represents the age where the donor stops reproducing and starts helping instead (see Methods). A) Resident killer whales, B) Transient killer whales, with 10-year increments. C) Humans, D) Chimpanzees, E) Asian elephants, F) African elephants, G) Sperm whales, with 5-year increments. H) Amboseli baboons with 1-year increments. The red line represents $\lambda$ , the yellow line represents the cost calculation for the donor, the light blue line represents the benefit calculation, the blue line represents the benefit calculation while accounting for the kinship dynamics, and the vertical dotted line represents the age where menopause first becomes favorable. The absence of the vertical dotted line implies $\lambda \leq 0$ for all ages.

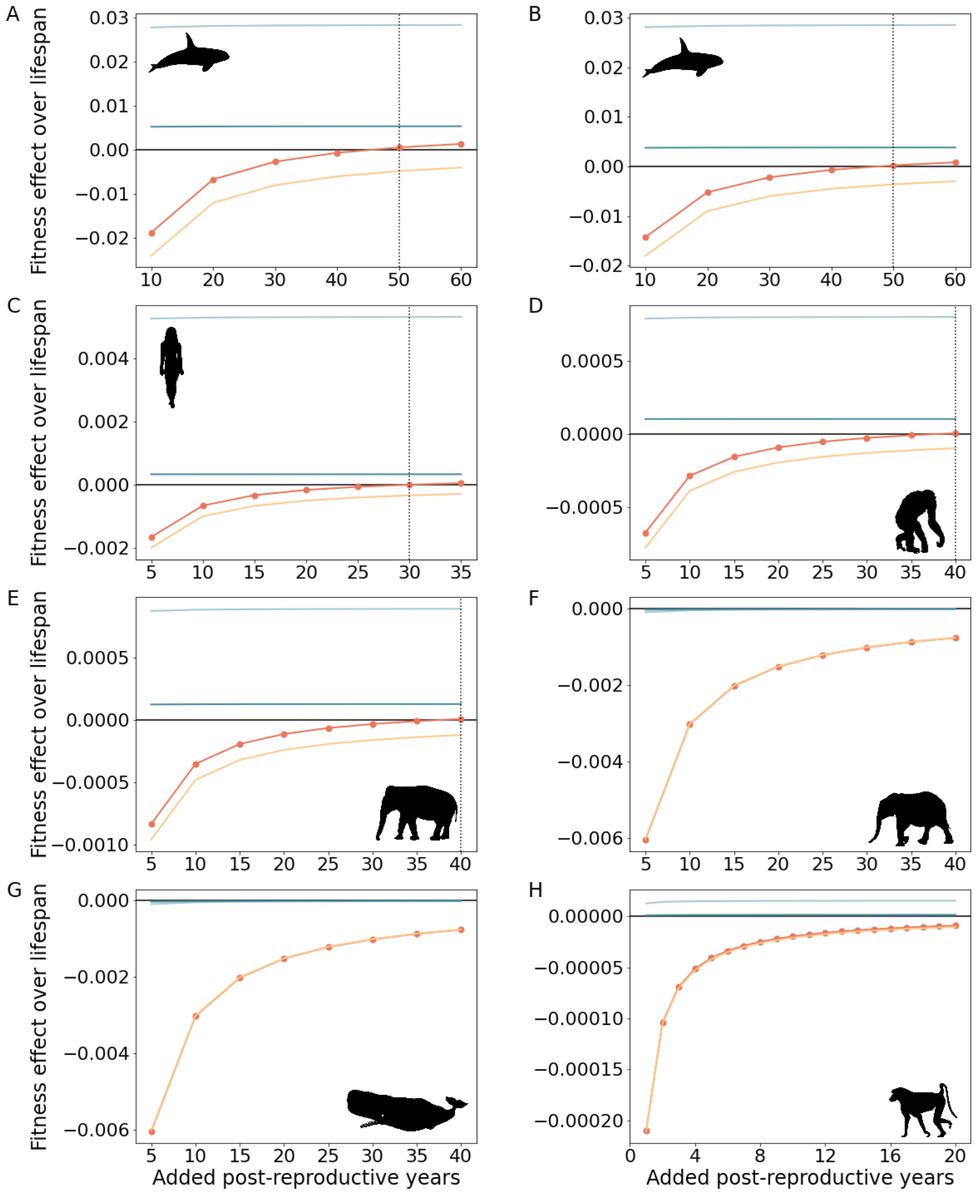


Figure S2: **$\lambda$, cost , benefit and relatedness-weighted benefit across increasing post-reproductive lifespan, under the live long pathway for the evolution of menopause.** The x-axis represents the number of added post-reproductive years, in year increments corresponding to the species' age interval in their matrices. A) Resident killer whales, B) Transient killer whales, with 10-year increments. C) Humans, D) Chimpanzees, E) Asian elephants, F) African elephants, G) Sperm whales, with 5-year increments. H) Amboseli baboons with 1-year increments. The red line represents $\lambda$, the yellow line represents the cost calculation for the donor, the light blue line represents the benefit calculation, the blue line represents the benefit calculation while accounting for the kinship dynamics, and the vertical dotted line represents the age where menopause first becomes favorable. The absence of the vertical dotted line implies $\lambda \leq 0$ for all ages.

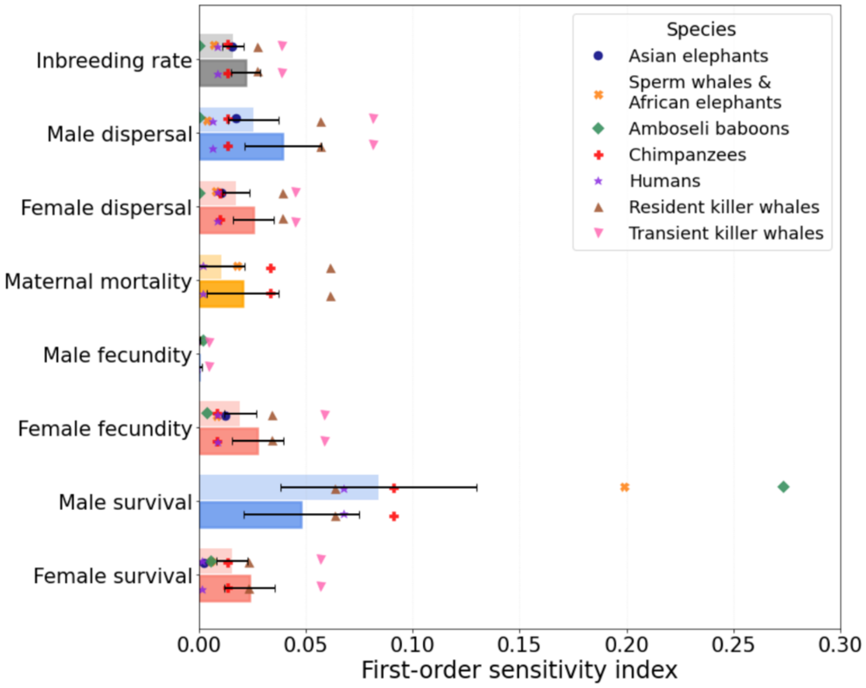


Figure S3: **First-order Sobol sensitivity indices for the evolution of menopause.** The first-order sensitivity indices measure the contribution of each parameter (y-axis) to the variance in the invasion criterion λ. Bars represent the contribution of each parameter to model variance. Parameters include helping effects on survival and fecundity of younger females and males, and changes in maternal mortality associated with continued reproduction at later ages. Markers show indices in different species. Dark and light bars show the average indices in menopausal species and all species, respectively.

**Low dispersal and high inbreeding promote the evolution of menopause.** We next focus on resident killer whales. Our model predicts reproductive cessation at ~45-55 years. To examine the invasion dynamics, we vary the female and male dispersal rates ($d_f$ and $d_m$) and the inbreeding rate ($m$) while holding all other parameters at resident killer whale values (Figure S1A). Thus, the direct and indirect fitness costs and benefits of menopause are fixed (as they do not depend on $d_f$, $d_m$ and $m$); only the focal female's relatedness to her pod changes as the parameters vary.

As expected, increasing the dispersal of either sex lowers a donor's relatedness to her pod[1] and thus reduces the invasion criterion λ, making menopause less favored by selection (Figure S4A). Conversely, increasing the inbreeding rate raises relatedness[1] and increases λ, favoring menopause (Figure S4B-C). Although high inbreeding with minimal dispersal strongly favors menopause, it would also elevate inbreeding depression, which is likely to be maladaptive[2]. These patterns accord with kin-selection theory[1], highlighting the role of sex-specific dispersal and mating patterns in the evolution of menopause.

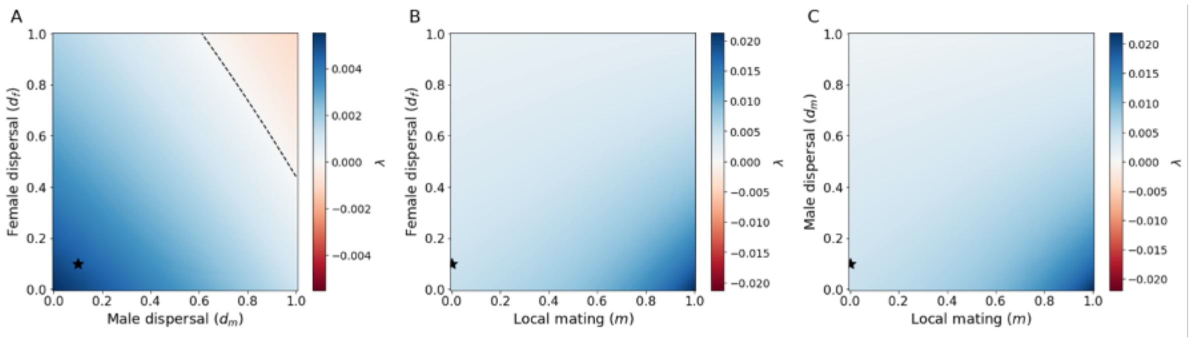


Figure S4**: Low dispersal and high inbreeding favor the evolution of menopause.** Panels show the invasion criterion λ at the menopause age predicted for resident killer whales while varying two parameters, and holding the third at the resident killer whale defaults ($d_f = 0.1, d_m = 0.1, m = 0$; black star). Menopause evolves in the blue regions (λ>0). Dashed contour lines marks λ = 0.

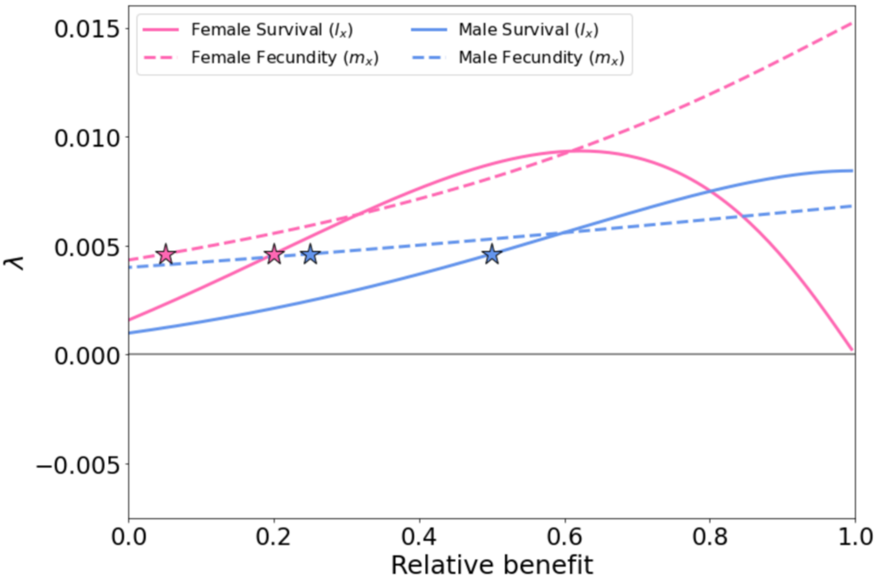


Figure S5: **Sensitivity of $\lambda$ to post-reproductive help in killer whales.** The figure shows the effect of increasing help on survival (solid line) and fecundity (dashed line) when directed to females (pink) or males (blue). The x-axis represents the relative benefit ($1 - \frac{non-menopausal\ value}{menopausal\ value}$), where only the numerator (the non-menopausal parameter) changes while the denominator (the menopausal parameter) remains fixed. 0 represents the baseline empirical values of the menopausal population ($\frac{non-menopausal\ value}{menopausal\ value} = 1$), and 1 represents the maximal benefit for menopause ($non - menopausal\ value \ll\ menopausal\ value$) and thus the highest λ.

**Longer post-reproductive lifespan can enable menopause under the ‘live long’ route in humans and killer whales.** While the stop early pathway focuses on early reproductive cessation producing a post-reproductive lifespan without increased longevity, the live long pathway asks whether shifting resources from reproduction to survival can increase longevity via a post-reproductive lifespan. Under disposable-soma assumptions of a finite energy budget, greater maintenance reduces fecundity, whereas prioritizing fecundity increases mortality and accelerates senescence[3]. We therefore tested whether this survival–fecundity trade-off allows a menopause-inducing modifier to invade when post-reproductive lifespan is extended (Figure S6), mapped the trade-off costs that permit invasion, and quantified how selection strength (λ) changes with each additional post-reproductive year.

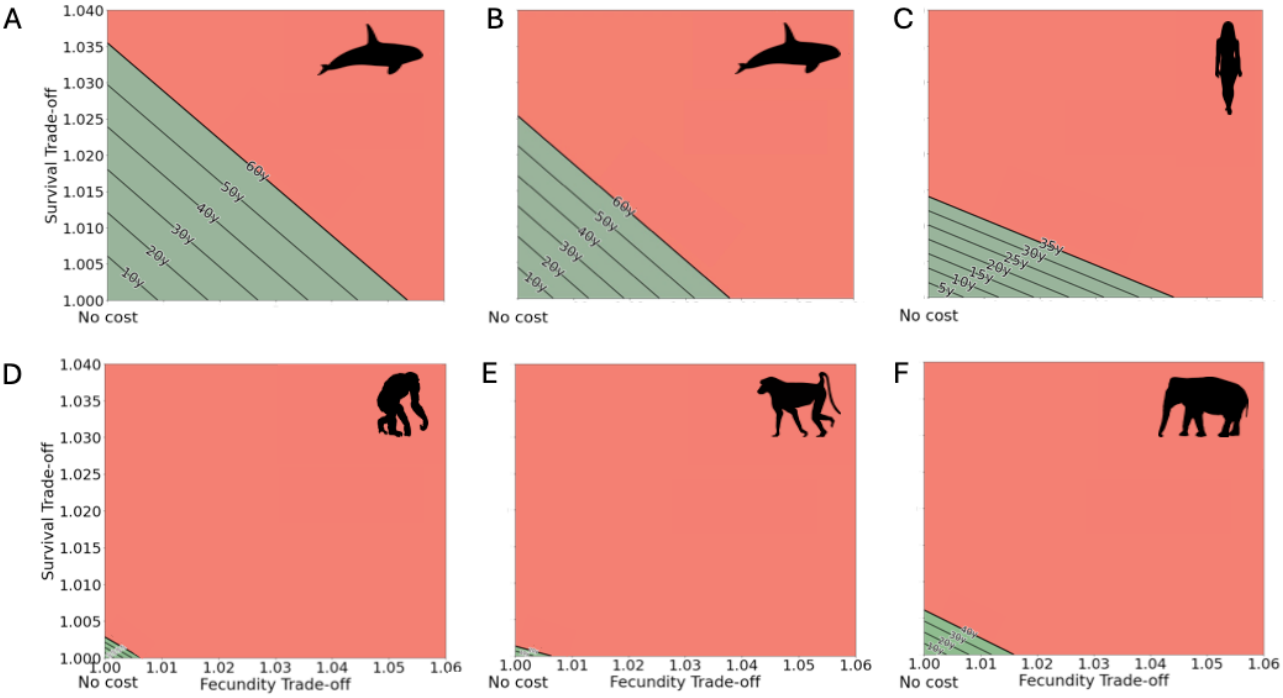


Figure S6: **Longer post-reproductive lifespan can enable menopause under the ‘live long’ route in humans and killer whales.** Axes show trade-off (cost) multipliers on fecundity (x) and early-life survival (y) associated with reallocating effort to longevity (=1, no cost; >1, stronger costs/lower rates). Diagonal contours give the λ=0 boundary for the labeled increase in post-reproductive lifespan (PRLS); for each label, parameter combinations below the corresponding contour yield λ>0 (green; menopause can invade), whereas combinations above yield λ<0 (red; menopause cannot evolve). (A) Resident killer whales (10-year increments). (B) Transient killer whales (10-year increments). (C) Humans (5-year increments). (D) Chimpanzees (5-y shown up to 10-y); (E) Amboseli baboons (1-y shown up to 10-y); (F) Asian elephants (5-y shown up to 10-y). Sperm whales and African elephants are omitted because no explored trade-off combinations obtained λ>0.

**Table S1: Sex-specific Leslie matrix for a resident/transient killer whale population.** A) Female population values ($L^f$); B) Male population values ($L^m$).

A

| | | | | | | | | | |
|---|---|---|---|---|---|---|---|---|---|
| 0 | 1 | 2 | 2 | 0 | 0 | 0 | 0 | 0 | 0 |
| 0.781 | 0 | 0 | 0 | 0 | 0 | 0 | 0 | 0 | 0 |
| 0 | 0.973 | 0 | 0 | 0 | 0 | 0 | 0 | 0 | 0 |
| 0 | 0 | 0.996 | 0 | 0 | 0 | 0 | 0 | 0 | 0 |
| 0 | 0 | 0 | 0.986 | 0 | 0 | 0 | 0 | 0 | 0 |
| 0 | 0 | 0 | 0 | 0.964 | 0 | 0 | 0 | 0 | 0 |
| 0 | 0 | 0 | 0 | 0 | 0.942 | 0 | 0 | 0 | 0 |
| 0 | 0 | 0 | 0 | 0 | 0 | 0.898 | 0 | 0 | 0 |
| 0 | 0 | 0 | 0 | 0 | 0 | 0 | 0.862 | 0 | 0 |
| 0 | 0 | 0 | 0 | 0 | 0 | 0 | 0 | 0.862 | 0 |

B

$$\begin{bmatrix}
0 & 1 & 2 & 2 & 0 & 0 & 0 & 0 & 0 & 0 \\
0.781 & 0 & 0 & 0 & 0 & 0 & 0 & 0 & 0 & 0 \\
0 & 0.931 & 0 & 0 & 0 & 0 & 0 & 0 & 0 & 0 \\
0 & 0 & 0.915 & 0 & 0 & 0 & 0 & 0 & 0 & 0 \\
0 & 0 & 0 & 0.859 & 0 & 0 & 0 & 0 & 0 & 0 \\
0 & 0 & 0 & 0 & 0.859 & 0 & 0 & 0 & 0 & 0 \\
0 & 0 & 0 & 0 & 0 & 0.859 & 0 & 0 & 0 & 0 \\
0 & 0 & 0 & 0 & 0 & 0 & 0 & 0 & 0 & 0 \\
0 & 0 & 0 & 0 & 0 & 0 & 0 & 0 & 0 & 0 \\
0 & 0 & 0 & 0 & 0 & 0 & 0 & 0 & 0 & 0
\end{bmatrix}$$

**Table S2: Sex-specific Leslie matrix for a sperm whale and African elephant population.** A) Female population values ($L^f$); B) Male population values ($L^m$).

A

| | | | | | | | | | | | | | | |
|---|---|---|---|---|---|---|---|---|---|---|---|---|---|---|
| 0 | 0 | 1 | 0.95 | 0.902 | 0.857 | 0.815 | 0.774 | 0.077 | 0.008 | 0.001 | 0 | 0 | 0 | 0 |
| 0.84 | 0 | 0 | 0 | 0 | 0 | 0 | 0 | 0 | 0 | 0 | 0 | 0 | 0 | 0 |
| 0 | 0.99 | 0 | 0 | 0 | 0 | 0 | 0 | 0 | 0 | 0 | 0 | 0 | 0 | 0 |
| 0 | 0 | 0.99 | 0 | 0 | 0 | 0 | 0 | 0 | 0 | 0 | 0 | 0 | 0 | 0 |
| 0 | 0 | 0 | 0.97 | 0 | 0 | 0 | 0 | 0 | 0 | 0 | 0 | 0 | 0 | 0 |
| 0 | 0 | 0 | 0 | 0.99 | 0 | 0 | 0 | 0 | 0 | 0 | 0 | 0 | 0 | 0 |
| 0 | 0 | 0 | 0 | 0 | 0.95 | 0 | 0 | 0 | 0 | 0 | 0 | 0 | 0 | 0 |
| 0 | 0 | 0 | 0 | 0 | 0 | 0.95 | 0 | 0 | 0 | 0 | 0 | 0 | 0 | 0 |
| 0 | 0 | 0 | 0 | 0 | 0 | 0 | 0.95 | 0 | 0 | 0 | 0 | 0 | 0 | 0 |
| 0 | 0 | 0 | 0 | 0 | 0 | 0 | 0 | 0.9 | 0 | 0 | 0 | 0 | 0 | 0 |
| 0 | 0 | 0 | 0 | 0 | 0 | 0 | 0 | 0 | 0.93 | 0 | 0 | 0 | 0 | 0 |
| 0 | 0 | 0 | 0 | 0 | 0 | 0 | 0 | 0 | 0 | 0.74 | 0 | 0 | 0 | 0 |
| 0 | 0 | 0 | 0 | 0 | 0 | 0 | 0 | 0 | 0 | 0 | 0.74 | 0 | 0 | 0 |
| 0 | 0 | 0 | 0 | 0 | 0 | 0 | 0 | 0 | 0 | 0 | 0 | 0.59 | 0 | 0 |
| 0 | 0 | 0 | 0 | 0 | 0 | 0 | 0 | 0 | 0 | 0 | 0 | 0 | 0.47 | 0 |

B

$$
\begin{bmatrix}
0 & 0 & 0 & 0 & 1 & 1 & 1 & 1 & 1 & 1 & 1 & 1 & 0 & 0 & 0 \\
0.79 & 0 & 0 & 0 & 0 & 0 & 0 & 0 & 0 & 0 & 0 & 0 & 0 & 0 & 0 \\
0 & 0.96 & 0 & 0 & 0 & 0 & 0 & 0 & 0 & 0 & 0 & 0 & 0 & 0 & 0 \\
0 & 0 & 0.96 & 0 & 0 & 0 & 0 & 0 & 0 & 0 & 0 & 0 & 0 & 0 & 0 \\
0 & 0 & 0 & 0.89 & 0 & 0 & 0 & 0 & 0 & 0 & 0 & 0 & 0 & 0 & 0 \\
0 & 0 & 0 & 0 & 0.91 & 0 & 0 & 0 & 0 & 0 & 0 & 0 & 0 & 0 & 0 \\
0 & 0 & 0 & 0 & 0 & 0.9 & 0 & 0 & 0 & 0 & 0 & 0 & 0 & 0 & 0 \\
0 & 0 & 0 & 0 & 0 & 0 & 0.88 & 0 & 0 & 0 & 0 & 0 & 0 & 0 & 0 \\
0 & 0 & 0 & 0 & 0 & 0 & 0 & 0.88 & 0 & 0 & 0 & 0 & 0 & 0 & 0 \\
0 & 0 & 0 & 0 & 0 & 0 & 0 & 0 & 0.89 & 0 & 0 & 0 & 0 & 0 & 0 \\
0 & 0 & 0 & 0 & 0 & 0 & 0 & 0 & 0 & 0.79 & 0 & 0 & 0 & 0 & 0 \\
0 & 0 & 0 & 0 & 0 & 0 & 0 & 0 & 0 & 0 & 0.87 & 0 & 0 & 0 & 0 \\
0 & 0 & 0 & 0 & 0 & 0 & 0 & 0 & 0 & 0 & 0 & 0.75 & 0 & 0 & 0 \\
0 & 0 & 0 & 0 & 0 & 0 & 0 & 0 & 0 & 0 & 0 & 0 & 0.696 & 0 & 0 \\
0 & 0 & 0 & 0 & 0 & 0 & 0 & 0 & 0 & 0 & 0 & 0 & 0 & 0.6 & 0
\end{bmatrix}
$$

**Table S3:** Sex-specific Leslie matrix for an Asian elephant population. A) Female population values ($L^f$); B) Male population values ($L^m$).

A

| | | | | | | | | | | | | |
|---|---|---|---|---|---|---|---|---|---|---|---|---|
| 0 | 0 | 0.28 | 0.28 | 0.4 | 0.4 | 0.4 | 0.40 | 0.4 | 0.4 | 0.1 | 0.1 | 0 |
| 0.76 | 0 | 0 | 0 | 0 | 0 | 0 | 0 | 0 | 0 | 0 | 0 | 0 |
| 0 | 0.93 | 0 | 0 | 0 | 0 | 0 | 0 | 0 | 0 | 0 | 0 | 0 |
| 0 | 0 | 0.93 | 0 | 0 | 0 | 0 | 0 | 0 | 0 | 0 | 0 | 0 |
| 0 | 0 | 0 | 0.86 | 0 | 0 | 0 | 0 | 0 | 0 | 0 | 0 | 0 |
| 0 | 0 | 0 | 0 | 0.95 | 0 | 0 | 0 | 0 | 0 | 0 | 0 | 0 |
| 0 | 0 | 0 | 0 | 0 | 0.95 | 0 | 0 | 0 | 0 | 0 | 0 | 0 |
| 0 | 0 | 0 | 0 | 0 | 0 | 0.95 | 0 | 0 | 0 | 0 | 0 | 0 |
| 0 | 0 | 0 | 0 | 0 | 0 | 0 | 0.95 | 0 | 0 | 0 | 0 | 0 |
| 0 | 0 | 0 | 0 | 0 | 0 | 0 | 0 | 0.95 | 0 | 0 | 0 | 0 |
| 0 | 0 | 0 | 0 | 0 | 0 | 0 | 0 | 0 | 0.95 | 0 | 0 | 0 |
| 0 | 0 | 0 | 0 | 0 | 0 | 0 | 0 | 0 | 0 | 0.59 | 0 | 0 |
| 0 | 0 | 0 | 0 | 0 | 0 | 0 | 0 | 0 | 0 | 0 | 0.59 | 0 |

B

$$
\begin{bmatrix}
0 & 0 & 0.28 & 0.28 & 0.4 & 0.4 & 0.4 & 0.4 & 0.4 & 0.4 & 0.1 & 0.1 & 0 \\
0.64 & 0 & 0 & 0 & 0 & 0 & 0 & 0 & 0 & 0 & 0 & 0 & 0 \\
0 & 0.72 & 0 & 0 & 0 & 0 & 0 & 0 & 0 & 0 & 0 & 0 & 0 \\
0 & 0 & 0.82 & 0 & 0 & 0 & 0 & 0 & 0 & 0 & 0 & 0 & 0 \\
0 & 0 & 0 & 0.82 & 0 & 0 & 0 & 0 & 0 & 0 & 0 & 0 & 0 \\
0 & 0 & 0 & 0 & 0.82 & 0 & 0 & 0 & 0 & 0 & 0 & 0 & 0 \\
0 & 0 & 0 & 0 & 0 & 0.82 & 0 & 0 & 0 & 0 & 0 & 0 & 0 \\
0 & 0 & 0 & 0 & 0 & 0 & 0.82 & 0 & 0 & 0 & 0 & 0 & 0 \\
0 & 0 & 0 & 0 & 0 & 0 & 0 & 0.82 & 0 & 0 & 0 & 0 & 0 \\
0 & 0 & 0 & 0 & 0 & 0 & 0 & 0 & 0.82 & 0 & 0 & 0 & 0 \\
0 & 0 & 0 & 0 & 0 & 0 & 0 & 0 & 0 & 0.82 & 0 & 0 & 0 \\
0 & 0 & 0 & 0 & 0 & 0 & 0 & 0 & 0 & 0 & 0.59 & 0 & 0 \\
0 & 0 & 0 & 0 & 0 & 0 & 0 & 0 & 0 & 0 & 0 & 0.59 & 0
\end{bmatrix}
$$

**Table S4: Sex-specific Leslie matrix for a human population**. A) Female population values ($L^f$); B) Male population values ($L^m$).

A

| | | | | | | | | | | | | | | | |
|---|---|---|---|---|---|---|---|---|---|---|---|---|---|---|---|
| 0 | 0 | 0.143 | 1.446 | 1.641 | 1.88 | 1.552 | 1.194 | 0.536 | 0 | 0 | 0 | 0 | 0 | 0 | 0 |
| 0.837 | 0 | 0 | 0 | 0 | 0 | 0 | 0 | 0 | 0 | 0 | 0 | 0 | 0 | 0 | 0 |
| 0 | 0.868 | 0 | 0 | 0 | 0 | 0 | 0 | 0 | 0 | 0 | 0 | 0 | 0 | 0 | 0 |
| 0 | 0 | 0.956 | 0 | 0 | 0 | 0 | 0 | 0 | 0 | 0 | 0 | 0 | 0 | 0 | 0 |
| 0 | 0 | 0 | 0.961 | 0 | 0 | 0 | 0 | 0 | 0 | 0 | 0 | 0 | 0 | 0 | 0 |
| 0 | 0 | 0 | 0 | 0.986 | 0 | 0 | 0 | 0 | 0 | 0 | 0 | 0 | 0 | 0 | 0 |
| 0 | 0 | 0 | 0 | 0 | 0.984 | 0 | 0 | 0 | 0 | 0 | 0 | 0 | 0 | 0 | 0 |
| 0 | 0 | 0 | 0 | 0 | 0 | 0.996 | 0 | 0 | 0 | 0 | 0 | 0 | 0 | 0 | 0 |
| 0 | 0 | 0 | 0 | 0 | 0 | 0 | 0.968 | 0 | 0 | 0 | 0 | 0 | 0 | 0 | 0 |
| 0 | 0 | 0 | 0 | 0 | 0 | 0 | 0 | 0.996 | 0 | 0 | 0 | 0 | 0 | 0 | 0 |
| 0 | 0 | 0 | 0 | 0 | 0 | 0 | 0 | 0 | 0.980 | 0 | 0 | 0 | 0 | 0 | 0 |
| 0 | 0 | 0 | 0 | 0 | 0 | 0 | 0 | 0 | 0 | 0.911 | 0 | 0 | 0 | 0 | 0 |
| 0 | 0 | 0 | 0 | 0 | 0 | 0 | 0 | 0 | 0 | 0 | 0.957 | 0 | 0 | 0 | 0 |
| 0 | 0 | 0 | 0 | 0 | 0 | 0 | 0 | 0 | 0 | 0 | 0 | 0.880 | 0 | 0 | 0 |
| 0 | 0 | 0 | 0 | 0 | 0 | 0 | 0 | 0 | 0 | 0 | 0 | 0 | 0.909 | 0 | 0 |
| 0 | 0 | 0 | 0 | 0 | 0 | 0 | 0 | 0 | 0 | 0 | 0 | 0 | 0 | 1 | 0 |

B

$$
\begin{bmatrix}
0 & 0 & 0 & 0.342 & 1.143 & 1.469 & 1.17 & 0.794 & 0.747 & 0.436 & 0.146 & 0.172 & 0 & 0 & 0 & 0 \\
0.848 & 0 & 0 & 0 & 0 & 0 & 0 & 0 & 0 & 0 & 0 & 0 & 0 & 0 & 0 & 0 \\
0 & 0.929 & 0 & 0 & 0 & 0 & 0 & 0 & 0 & 0 & 0 & 0 & 0 & 0 & 0 & 0 \\
0 & 0 & 1 & 0 & 0 & 0 & 0 & 0 & 0 & 0 & 0 & 0 & 0 & 0 & 0 & 0 \\
0 & 0 & 0 & 1 & 0 & 0 & 0 & 0 & 0 & 0 & 0 & 0 & 0 & 0 & 0 & 0 \\
0 & 0 & 0 & 0 & 0.975 & 0 & 0 & 0 & 0 & 0 & 0 & 0 & 0 & 0 & 0 & 0 \\
0 & 0 & 0 & 0 & 0 & 0.968 & 0 & 0 & 0 & 0 & 0 & 0 & 0 & 0 & 0 & 0 \\
0 & 0 & 0 & 0 & 0 & 0 & 1 & 0 & 0 & 0 & 0 & 0 & 0 & 0 & 0 & 0 \\
0 & 0 & 0 & 0 & 0 & 0 & 0 & 0.996 & 0 & 0 & 0 & 0 & 0 & 0 & 0 & 0 \\
0 & 0 & 0 & 0 & 0 & 0 & 0 & 0 & 0.972 & 0 & 0 & 0 & 0 & 0 & 0 & 0 \\
0 & 0 & 0 & 0 & 0 & 0 & 0 & 0 & 0 & 0.952 & 0 & 0 & 0 & 0 & 0 & 0 \\
0 & 0 & 0 & 0 & 0 & 0 & 0 & 0 & 0 & 0 & 0.952 & 0 & 0 & 0 & 0 & 0 \\
0 & 0 & 0 & 0 & 0 & 0 & 0 & 0 & 0 & 0 & 0 & 0.982 & 0 & 0 & 0 & 0 \\
0 & 0 & 0 & 0 & 0 & 0 & 0 & 0 & 0 & 0 & 0 & 0 & 0.900 & 0 & 0 & 0 \\
0 & 0 & 0 & 0 & 0 & 0 & 0 & 0 & 0 & 0 & 0 & 0 & 0 & 0.662 & 0 & 0 \\
0 & 0 & 0 & 0 & 0 & 0 & 0 & 0 & 0 & 0 & 0 & 0 & 0 & 0 & 0.705 & 0
\end{bmatrix}
$$

**Table S5: Sex-specific Leslie matrix for an Amboseli baboon population**. A) Female population values ($L^f$); B) Male population values ($L^m$).

A

| | | | | | | | | | | | | | | | | | | | | | | | | | | |
|---|---|---|---|---|---|---|---|---|---|---|---|---|---|---|---|---|---|---|---|---|---|---|---|---|---|---|
| 0 | 0 | 0 | 0 | 0.128 | 0.272 | 0.275 | 0.265 | 0.260 | 0.224 | 0.198 | 0.231 | 0.262 | 0.260 | 0.284 | 0.301 | 0.266 | 0.217 | 0.244 | 0.184 | 0.240 | 0.322 | 0.131 | 0.214 | 0 | 0 | 0 |
| 0.794 | 0 | 0 | 0 | 0 | 0 | 0 | 0 | 0 | 0 | 0 | 0 | 0 | 0 | 0 | 0 | 0 | 0 | 0 | 0 | 0 | 0 | 0 | 0 | 0 | 0 | 0 |
| 0 | 0.888 | 0 | 0 | 0 | 0 | 0 | 0 | 0 | 0 | 0 | 0 | 0 | 0 | 0 | 0 | 0 | 0 | 0 | 0 | 0 | 0 | 0 | 0 | 0 | 0 | 0 |
| 0 | 0 | 0.937 | 0 | 0 | 0 | 0 | 0 | 0 | 0 | 0 | 0 | 0 | 0 | 0 | 0 | 0 | 0 | 0 | 0 | 0 | 0 | 0 | 0 | 0 | 0 | 0 |
| 0 | 0 | 0 | 0.969 | 0 | 0 | 0 | 0 | 0 | 0 | 0 | 0 | 0 | 0 | 0 | 0 | 0 | 0 | 0 | 0 | 0 | 0 | 0 | 0 | 0 | 0 | 0 |
| 0 | 0 | 0 | 0 | 0.953 | 0 | 0 | 0 | 0 | 0 | 0 | 0 | 0 | 0 | 0 | 0 | 0 | 0 | 0 | 0 | 0 | 0 | 0 | 0 | 0 | 0 | 0 |
| 0 | 0 | 0 | 0 | 0 | 0.944 | 0 | 0 | 0 | 0 | 0 | 0 | 0 | 0 | 0 | 0 | 0 | 0 | 0 | 0 | 0 | 0 | 0 | 0 | 0 | 0 | 0 |
| 0 | 0 | 0 | 0 | 0 | 0 | 0.948 | 0 | 0 | 0 | 0 | 0 | 0 | 0 | 0 | 0 | 0 | 0 | 0 | 0 | 0 | 0 | 0 | 0 | 0 | 0 | 0 |
| 0 | 0 | 0 | 0 | 0 | 0 | 0 | 0.948 | 0 | 0 | 0 | 0 | 0 | 0 | 0 | 0 | 0 | 0 | 0 | 0 | 0 | 0 | 0 | 0 | 0 | 0 | 0 |
| 0 | 0 | 0 | 0 | 0 | 0 | 0 | 0 | 0.943 | 0 | 0 | 0 | 0 | 0 | 0 | 0 | 0 | 0 | 0 | 0 | 0 | 0 | 0 | 0 | 0 | 0 | 0 |
| 0 | 0 | 0 | 0 | 0 | 0 | 0 | 0 | 0 | 0.923 | 0 | 0 | 0 | 0 | 0 | 0 | 0 | 0 | 0 | 0 | 0 | 0 | 0 | 0 | 0 | 0 | 0 |
| 0 | 0 | 0 | 0 | 0 | 0 | 0 | 0 | 0 | 0 | 0.891 | 0 | 0 | 0 | 0 | 0 | 0 | 0 | 0 | 0 | 0 | 0 | 0 | 0 | 0 | 0 | 0 |
| 0 | 0 | 0 | 0 | 0 | 0 | 0 | 0 | 0 | 0 | 0 | 0.894 | 0 | 0 | 0 | 0 | 0 | 0 | 0 | 0 | 0 | 0 | 0 | 0 | 0 | 0 | 0 |
| 0 | 0 | 0 | 0 | 0 | 0 | 0 | 0 | 0 | 0 | 0 | 0 | 0.942 | 0 | 0 | 0 | 0 | 0 | 0 | 0 | 0 | 0 | 0 | 0 | 0 | 0 | 0 |
| 0 | 0 | 0 | 0 | 0 | 0 | 0 | 0 | 0 | 0 | 0 | 0 | 0 | 0.916 | 0 | 0 | 0 | 0 | 0 | 0 | 0 | 0 | 0 | 0 | 0 | 0 | 0 |
| 0 | 0 | 0 | 0 | 0 | 0 | 0 | 0 | 0 | 0 | 0 | 0 | 0 | 0 | 0.870 | 0 | 0 | 0 | 0 | 0 | 0 | 0 | 0 | 0 | 0 | 0 | 0 |
| 0 | 0 | 0 | 0 | 0 | 0 | 0 | 0 | 0 | 0 | 0 | 0 | 0 | 0 | 0 | 0.873 | 0 | 0 | 0 | 0 | 0 | 0 | 0 | 0 | 0 | 0 | 0 |
| 0 | 0 | 0 | 0 | 0 | 0 | 0 | 0 | 0 | 0 | 0 | 0 | 0 | 0 | 0 | 0 | 0.788 | 0 | 0 | 0 | 0 | 0 | 0 | 0 | 0 | 0 | 0 |
| 0 | 0 | 0 | 0 | 0 | 0 | 0 | 0 | 0 | 0 | 0 | 0 | 0 | 0 | 0 | 0 | 0 | 0.633 | 0 | 0 | 0 | 0 | 0 | 0 | 0 | 0 | 0 |
| 0 | 0 | 0 | 0 | 0 | 0 | 0 | 0 | 0 | 0 | 0 | 0 | 0 | 0 | 0 | 0 | 0 | 0 | 0.628 | 0 | 0 | 0 | 0 | 0 | 0 | 0 | 0 |
| 0 | 0 | 0 | 0 | 0 | 0 | 0 | 0 | 0 | 0 | 0 | 0 | 0 | 0 | 0 | 0 | 0 | 0 | 0 | 0.865 | 0 | 0 | 0 | 0 | 0 | 0 | 0 |
| 0 | 0 | 0 | 0 | 0 | 0 | 0 | 0 | 0 | 0 | 0 | 0 | 0 | 0 | 0 | 0 | 0 | 0 | 0 | 0 | 0.746 | 0 | 0 | 0 | 0 | 0 | 0 |
| 0 | 0 | 0 | 0 | 0 | 0 | 0 | 0 | 0 | 0 | 0 | 0 | 0 | 0 | 0 | 0 | 0 | 0 | 0 | 0 | 0 | 0.552 | 0 | 0 | 0 | 0 | 0 |
| 0 | 0 | 0 | 0 | 0 | 0 | 0 | 0 | 0 | 0 | 0 | 0 | 0 | 0 | 0 | 0 | 0 | 0 | 0 | 0 | 0 | 0 | 0.609 | 0 | 0 | 0 | 0 |
| 0 | 0 | 0 | 0 | 0 | 0 | 0 | 0 | 0 | 0 | 0 | 0 | 0 | 0 | 0 | 0 | 0 | 0 | 0 | 0 | 0 | 0 | 0 | 0.417 | 0 | 0 | 0 |
| 0 | 0 | 0 | 0 | 0 | 0 | 0 | 0 | 0 | 0 | 0 | 0 | 0 | 0 | 0 | 0 | 0 | 0 | 0 | 0 | 0 | 0 | 0 | 0 | 0.380 | 0 | 0 |
| 0 | 0 | 0 | 0 | 0 | 0 | 0 | 0 | 0 | 0 | 0 | 0 | 0 | 0 | 0 | 0 | 0 | 0 | 0 | 0 | 0 | 0 | 0 | 0 | 0 | 0.500 | 0 |

B

| | | | | | | | | | | | | | | | | | | | | | | | | | | |
|---|---|---|---|---|---|---|---|---|---|---|---|---|---|---|---|---|---|---|---|---|---|---|---|---|---|---|
| 0 | 0 | 0 | 0 | 0 | 0.373 | 0.776 | 0.779 | 0.766 | 0.742 | 0.722 | 0.707 | 0.665 | 0.647 | 0.595 | 0.475 | 0.433 | 0.625 | 0.586 | 0.213 | 0 | 0 | 0 | 0 | 0 | 0 | 0 |
| 0.782 | 0 | 0 | 0 | 0 | 0 | 0 | 0 | 0 | 0 | 0 | 0 | 0 | 0 | 0 | 0 | 0 | 0 | 0 | 0 | 0 | 0 | 0 | 0 | 0 | 0 | 0 |
| 0 | 0.912 | 0 | 0 | 0 | 0 | 0 | 0 | 0 | 0 | 0 | 0 | 0 | 0 | 0 | 0 | 0 | 0 | 0 | 0 | 0 | 0 | 0 | 0 | 0 | 0 | 0 |
| 0 | 0 | 0.934 | 0 | 0 | 0 | 0 | 0 | 0 | 0 | 0 | 0 | 0 | 0 | 0 | 0 | 0 | 0 | 0 | 0 | 0 | 0 | 0 | 0 | 0 | 0 | 0 |
| 0 | 0 | 0 | 0.917 | 0 | 0 | 0 | 0 | 0 | 0 | 0 | 0 | 0 | 0 | 0 | 0 | 0 | 0 | 0 | 0 | 0 | 0 | 0 | 0 | 0 | 0 | 0 |
| 0 | 0 | 0 | 0 | 0.959 | 0 | 0 | 0 | 0 | 0 | 0 | 0 | 0 | 0 | 0 | 0 | 0 | 0 | 0 | 0 | 0 | 0 | 0 | 0 | 0 | 0 | 0 |
| 0 | 0 | 0 | 0 | 0 | 0.943 | 0 | 0 | 0 | 0 | 0 | 0 | 0 | 0 | 0 | 0 | 0 | 0 | 0 | 0 | 0 | 0 | 0 | 0 | 0 | 0 | 0 |
| 0 | 0 | 0 | 0 | 0 | 0 | 0.939 | 0 | 0 | 0 | 0 | 0 | 0 | 0 | 0 | 0 | 0 | 0 | 0 | 0 | 0 | 0 | 0 | 0 | 0 | 0 | 0 |
| 0 | 0 | 0 | 0 | 0 | 0 | 0 | 0.942 | 0 | 0 | 0 | 0 | 0 | 0 | 0 | 0 | 0 | 0 | 0 | 0 | 0 | 0 | 0 | 0 | 0 | 0 | 0 |
| 0 | 0 | 0 | 0 | 0 | 0 | 0 | 0 | 0.931 | 0 | 0 | 0 | 0 | 0 | 0 | 0 | 0 | 0 | 0 | 0 | 0 | 0 | 0 | 0 | 0 | 0 | 0 |
| 0 | 0 | 0 | 0 | 0 | 0 | 0 | 0 | 0 | 0.911 | 0 | 0 | 0 | 0 | 0 | 0 | 0 | 0 | 0 | 0 | 0 | 0 | 0 | 0 | 0 | 0 | 0 |
| 0 | 0 | 0 | 0 | 0 | 0 | 0 | 0 | 0 | 0 | 0.894 | 0 | 0 | 0 | 0 | 0 | 0 | 0 | 0 | 0 | 0 | 0 | 0 | 0 | 0 | 0 | 0 |
| 0 | 0 | 0 | 0 | 0 | 0 | 0 | 0 | 0 | 0 | 0 | 0.882 | 0 | 0 | 0 | 0 | 0 | 0 | 0 | 0 | 0 | 0 | 0 | 0 | 0 | 0 | 0 |
| 0 | 0 | 0 | 0 | 0 | 0 | 0 | 0 | 0 | 0 | 0 | 0 | 0.846 | 0 | 0 | 0 | 0 | 0 | 0 | 0 | 0 | 0 | 0 | 0 | 0 | 0 | 0 |
| 0 | 0 | 0 | 0 | 0 | 0 | 0 | 0 | 0 | 0 | 0 | 0 | 0 | 0.830 | 0 | 0 | 0 | 0 | 0 | 0 | 0 | 0 | 0 | 0 | 0 | 0 | 0 |
| 0 | 0 | 0 | 0 | 0 | 0 | 0 | 0 | 0 | 0 | 0 | 0 | 0 | 0 | 0.782 | 0 | 0 | 0 | 0 | 0 | 0 | 0 | 0 | 0 | 0 | 0 | 0 |
| 0 | 0 | 0 | 0 | 0 | 0 | 0 | 0 | 0 | 0 | 0 | 0 | 0 | 0 | 0 | 0.667 | 0 | 0 | 0 | 0 | 0 | 0 | 0 | 0 | 0 | 0 | 0 |
| 0 | 0 | 0 | 0 | 0 | 0 | 0 | 0 | 0 | 0 | 0 | 0 | 0 | 0 | 0 | 0 | 0.625 | 0 | 0 | 0 | 0 | 0 | 0 | 0 | 0 | 0 | 0 |
| 0 | 0 | 0 | 0 | 0 | 0 | 0 | 0 | 0 | 0 | 0 | 0 | 0 | 0 | 0 | 0 | 0 | 0.800 | 0 | 0 | 0 | 0 | 0 | 0 | 0 | 0 | 0 |
| 0 | 0 | 0 | 0 | 0 | 0 | 0 | 0 | 0 | 0 | 0 | 0 | 0 | 0 | 0 | 0 | 0 | 0 | 0.750 | 0 | 0 | 0 | 0 | 0 | 0 | 0 | 0 |
| 0 | 0 | 0 | 0 | 0 | 0 | 0 | 0 | 0 | 0 | 0 | 0 | 0 | 0 | 0 | 0 | 0 | 0 | 0 | 0.333 | 0 | 0 | 0 | 0 | 0 | 0 | 0 |
| 0 | 0 | 0 | 0 | 0 | 0 | 0 | 0 | 0 | 0 | 0 | 0 | 0 | 0 | 0 | 0 | 0 | 0 | 0 | 0 | 0 | 0 | 0 | 0 | 0 | 0 | 0 |
| 0 | 0 | 0 | 0 | 0 | 0 | 0 | 0 | 0 | 0 | 0 | 0 | 0 | 0 | 0 | 0 | 0 | 0 | 0 | 0 | 0 | 0 | 0 | 0 | 0 | 0 | 0 |
| 0 | 0 | 0 | 0 | 0 | 0 | 0 | 0 | 0 | 0 | 0 | 0 | 0 | 0 | 0 | 0 | 0 | 0 | 0 | 0 | 0 | 0 | 0 | 0 | 0 | 0 | 0 |
| 0 | 0 | 0 | 0 | 0 | 0 | 0 | 0 | 0 | 0 | 0 | 0 | 0 | 0 | 0 | 0 | 0 | 0 | 0 | 0 | 0 | 0 | 0 | 0 | 0 | 0 | 0 |
| 0 | 0 | 0 | 0 | 0 | 0 | 0 | 0 | 0 | 0 | 0 | 0 | 0 | 0 | 0 | 0 | 0 | 0 | 0 | 0 | 0 | 0 | 0 | 0 | 0 | 0 | 0 |
| 0 | 0 | 0 | 0 | 0 | 0 | 0 | 0 | 0 | 0 | 0 | 0 | 0 | 0 | 0 | 0 | 0 | 0 | 0 | 0 | 0 | 0 | 0 | 0 | 0 | 0 | 0 |

**Table S6: Sex-specific Leslie matrix for a chimpanzee population**. A) Female population values ($L^f$); B) Male population values ($L^m$).

A

| | | | | | | | | | | | | | |
|---|---|---|---|---|---|---|---|---|---|---|---|---|---|
| 0 | 0.053 | 0.685 | 1.077 | 1.069 | 1.121 | 0.900 | 0.793 | 0.620 | 0.345 | 0.052 | 0 | 0 | 0 |
| 0.764 | 0 | 0 | 0 | 0 | 0 | 0 | 0 | 0 | 0 | 0 | 0 | 0 | 0 |
| 0 | 0.921 | 0 | 0 | 0 | 0 | 0 | 0 | 0 | 0 | 0 | 0 | 0 | 0 |
| 0 | 0 | 1.000 | 0 | 0 | 0 | 0 | 0 | 0 | 0 | 0 | 0 | 0 | 0 |
| 0 | 0 | 0 | 0.980 | 0 | 0 | 0 | 0 | 0 | 0 | 0 | 0 | 0 | 0 |
| 0 | 0 | 0 | 0 | 0.960 | 0 | 0 | 0 | 0 | 0 | 0 | 0 | 0 | 0 |
| 0 | 0 | 0 | 0 | 0 | 0.950 | 0 | 0 | 0 | 0 | 0 | 0 | 0 | 0 |
| 0 | 0 | 0 | 0 | 0 | 0 | 0.902 | 0 | 0 | 0 | 0 | 0 | 0 | 0 |
| 0 | 0 | 0 | 0 | 0 | 0 | 0 | 1.000 | 0 | 0 | 0 | 0 | 0 | 0 |
| 0 | 0 | 0 | 0 | 0 | 0 | 0 | 0 | 0.884 | 0 | 0 | 0 | 0 | 0 |
| 0 | 0 | 0 | 0 | 0 | 0 | 0 | 0 | 0 | 0.818 | 0 | 0 | 0 | 0 |
| 0 | 0 | 0 | 0 | 0 | 0 | 0 | 0 | 0 | 0 | 0.712 | 0 | 0 | 0 |
| 0 | 0 | 0 | 0 | 0 | 0 | 0 | 0 | 0 | 0 | 0 | 0.688 | 0 | 0 |
| 0 | 0 | 0 | 0 | 0 | 0 | 0 | 0 | 0 | 0 | 0 | 0 | 0.503 | 0 |

B

| | | | | | | | | | | | | | |
|---|---|---|---|---|---|---|---|---|---|---|---|---|---|
| 0 | 0 | 0.455 | 1.909 | 1.086 | 1.071 | 0.967 | 1.092 | 0.376 | 1.236 | 0.564 | 0 | 0 | 0 |
| 0.820 | 0 | 0 | 0 | 0 | 0 | 0 | 0 | 0 | 0 | 0 | 0 | 0 | 0 |
| 0 | 1.000 | 0 | 0 | 0 | 0 | 0 | 0 | 0 | 0 | 0 | 0 | 0 | 0 |
| 0 | 0 | 1.000 | 0 | 0 | 0 | 0 | 0 | 0 | 0 | 0 | 0 | 0 | 0 |
| 0 | 0 | 0 | 0.951 | 0 | 0 | 0 | 0 | 0 | 0 | 0 | 0 | 0 | 0 |
| 0 | 0 | 0 | 0 | 0.756 | 0 | 0 | 0 | 0 | 0 | 0 | 0 | 0 | 0 |
| 0 | 0 | 0 | 0 | 0 | 0.883 | 0 | 0 | 0 | 0 | 0 | 0 | 0 | 0 |
| 0 | 0 | 0 | 0 | 0 | 0 | 0.845 | 0 | 0 | 0 | 0 | 0 | 0 | 0 |
| 0 | 0 | 0 | 0 | 0 | 0 | 0 | 0.752 | 0 | 0 | 0 | 0 | 0 | 0 |
| 0 | 0 | 0 | 0 | 0 | 0 | 0 | 0 | 0.625 | 0 | 0 | 0 | 0 | 0 |
| 0 | 0 | 0 | 0 | 0 | 0 | 0 | 0 | 0 | 0.622 | 0 | 0 | 0 | 0 |
| 0 | 0 | 0 | 0 | 0 | 0 | 0 | 0 | 0 | 0 | 0.335 | 0 | 0 | 0 |
| 0 | 0 | 0 | 0 | 0 | 0 | 0 | 0 | 0 | 0 | 0 | 0 | 0 | 0 |
| 0 | 0 | 0 | 0 | 0 | 0 | 0 | 0 | 0 | 0 | 0 | 0 | 0 | 0 |

**Table S7: Selected parameter sets illustrating survival and fecundity trade-offs across added post-reproductive age classes.**

| Species | Years Added | Survival Trade-off | Fecundity Trade-off |
|---|---|---|---|
| **Resident killer whales** | 10 | 1.01 | 1.025 |
| | 20 | 1.005 | 1.0125 |
| | 30 | 1.0033 | 1.0083 |
| | 40 | 1.0025 | 1.00625 |
| | 50 | 1.002 | 1.005 |
| | 60 | 1.0017 | 1.0042 |
| **Transient killer whales** | 10 | 1.01 | 1.015 |
| | 20 | 1.005 | 1.0075 |
| | 30 | 1.0033 | 1.005 |
| | 40 | 1.0025 | 1.00375 |
| | 50 | 1.002 | 1.003 |
| | 60 | 1.0017 | 1.0025 |
| **Humans** | 5 | 1.008 | 1.012 |
| | 10 | 1.004 | 1.006 |
| | 15 | 1.0027 | 1.004 |
| | 20 | 1.002 | 1.003 |
| | 25 | 1.0016 | 1.0024 |
| | 30 | 1.0013 | 1.002 |
| | 35 | 1.0011 | 1.0017 |
| **African elephants** | 5 | 1.008 | 1.012 |
| | 10 | 1.004 | 1.006 |
| | 15 | 1.0027 | 1.004 |
| | 20 | 1.002 | 1.003 |
| | 25 | 1.0016 | 1.0024 |
| | 30 | 1.0013 | 1.002 |
| | 35 | 1.0011 | 1.0017 |
| | 40 | 1.001 | 1.0015 |
| **Sperm whales** | 5 | 1.008 | 1.012 |
| | 10 | 1.004 | 1.006 |
| | 15 | 1.0027 | 1.004 |
| | 20 | 1.002 | 1.003 |
| | 25 | 1.0016 | 1.0024 |
| | 30 | 1.0013 | 1.002 |
| | 35 | 1.0011 | 1.0017 |
| | 40 | 1.001 | 1.0015 |
| | 5 | 1.003 | 1.007 |

| | | | |
|---|---|---|---|
| **Asian elephants** | 10 | 1.0015 | 1.0035 |
| | 15 | 1.001 | 1.0023 |
| | 20 | 1.00075 | 1.00175 |
| | 25 | 1.0006 | 1.0014 |
| | 30 | 1.0005 | 1.0012 |
| | 35 | 1.0004 | 1.001 |
| | 40 | 1.0004 | 1.0009 |
| **Amboseli baboons** | 1 | 1.008 | 1.012 |
| | 2 | 1.004 | 1.006 |
| | 3 | 1.0027 | 1.004 |
| | 4 | 1.002 | 1.003 |
| | 5 | 1.0016 | 1.0024 |
| | 6 | 1.0013 | 1.002 |
| | 7 | 1.0011 | 1.0017 |
| | 8 | 1.001 | 1.0015 |
| | 9 | 1.0009 | 1.0013 |
| | 10 | 1.0008 | 1.0012 |
| | 11 | 1.0007 | 1.0011 |
| | 12 | 1.0007 | 1.001 |
| | 13 | 1.0006 | 1.0009 |
| | 14 | 1.0006 | 1.0009 |
| | 15 | 1.0005 | 1.0008 |
| | 16 | 1.0005 | 1.0008 |
| | 17 | 1.0005 | 1.0007 |
| | 18 | 1.0004 | 1.0007 |
| | 19 | 1.0004 | 1.0006 |
| | 20 | 1.0004 | 1.0006 |
| **Chimpanzees** | 5 | 1.0018 | 1.002 |
| | 10 | 1.0009 | 1.001 |
| | 15 | 1.0006 | 1.0007 |
| | 20 | 1.00045 | 1.0005 |
| | 25 | 1.00036 | 1.0004 |
| | 30 | 1.0003 | 1.0003 |
| | 35 | 1.00026 | 1.00029 |
| | 40 | 1.00023 | 1.00025 |

**SI REFERENCES**

1. Johnstone, R. A. & Cant, M. A. The evolution of menopause in cetaceans and humans: the role of demography. *Proc. R. Soc. B.* **277**, 3765–3771 (2010).
2. Kardos, M. *et al.* Inbreeding depression explains killer whale population dynamics. *Nat Ecol Evol* **7**, 675–686 (2023).
3. Kirkwood, T. B. L. & Rose, M. R. Evolution of senescence: late survival sacrificed for reproduction. *Philosophical Transactions of the Royal Society of London. Series B: Biological Sciences* **332**, 15–24 (1991).
