## Supplementary Code for "Evolution of Menopause and the Mama’s Boy Hypothesis": all_results.ipynb

```
# %% [code]
import numpy as np 
from pandas import DataFrame, read_csv
import matplotlib.pyplot as plt
import seaborn as sns
from copy import deepcopy
from scipy.linalg import leslie
import matplotlib.image as mpimg
%matplotlib inline

# %% [code]
import io, os, sys, types
import glob

# %% [code]
from IPython import get_ipython
from nbformat import read
from IPython.core.interactiveshell import InteractiveShell

# %% [code]
def find_notebook(fullname, path=None):
    """find a notebook, given its fully qualified name and an optional path

    This turns "foo.bar" into "foo/bar.ipynb"
    and tries turning "Foo_Bar" into "Foo Bar" if Foo_Bar
    does not exist.
    """
    name = fullname.rsplit('.', 1)[-1]
    if not path:
        path = ['']
    for d in path:
        nb_path = os.path.join(d, name + ".ipynb")
        if os.path.isfile(nb_path):
            return nb_path
        # let import Notebook_Name find "Notebook Name.ipynb"
        nb_path = nb_path.replace("_", " ")
        if os.path.isfile(nb_path):
            return nb_path

# %% [code]
class NotebookLoader(object):
    """Module Loader for Jupyter Notebooks"""
    def __init__(self, path=None):
        self.shell = InteractiveShell.instance()
        self.path = path

    def load_module(self, fullname):
        """import a notebook as a module"""
        path = find_notebook(fullname, self.path)

        print ("importing Jupyter notebook from %s" % path)

        # load the notebook object
        with io.open(path, 'r', encoding='utf-8') as f:
            nb = read(f, 4)


        # create the module and add it to sys.modules
        # if name in sys.modules:
        #    return sys.modules[name]
        mod = types.ModuleType(fullname)
        mod.__file__ = path
        mod.__loader__ = self
        mod.__dict__['get_ipython'] = get_ipython
        sys.modules[fullname] = mod

        # extra work to ensure that magics that would affect the user_ns
        # actually affect the notebook module's ns
        save_user_ns = self.shell.user_ns
        self.shell.user_ns = mod.__dict__

        try:
          for cell in nb.cells:
            if cell.cell_type == 'code':
                # transform the input to executable Python
                code = self.shell.input_transformer_manager.transform_cell(cell.source)
                # run the code in themodule
                exec(code, mod.__dict__)
        finally:
            self.shell.user_ns = save_user_ns
        return mod

# %% [code]
class NotebookFinder(object):
    """Module finder that locates Jupyter Notebooks"""
    def __init__(self):
        self.loaders = {}

    def find_module(self, fullname, path=None):
        nb_path = find_notebook(fullname, path)
        if not nb_path:
            return

        key = path
        if path:
            # lists aren't hashable
            key = os.path.sep.join(path)

        if key not in self.loaders:
            self.loaders[key] = NotebookLoader(path)
        return self.loaders[key]

# %% [code]
sys.meta_path.append(NotebookFinder())

# %% [code]
from Johnstone_Cant import IBD_age_0, no_age_rs, IBD_no_age, IBD_older_age_v2, rff_age_v2

# %% [code]
from project_functions import survival2x, leslie_, Px, mx, age_distribution, tot_rBC_all_donor_life, tot_rBC, survival_effect, fecundity_effect, reproductive_value, \
different_onset_ages, latex_array, ci_intervals, plot_post_reproductive_life, PRL, PrR_from_model, PrR_from_LM, ci_intervals_by_PrR, plot_post_reproductive_life_boxplot, remaining_life_expectancy, calc_rcoeff, different_longevities, extend_LM_both_sexes, add_image, fig_panel_labels

# %% [code]
import os
import glob
import numpy as np
import pandas as pd
import matplotlib.pyplot as plt

# ================= SETTINGS =================
RESULTS_DIR = "sa_results"
FIGSIZE = (26, 10)
XMAX = 1.1
SHOW_ERRORBARS = False  # set True to add ±conf on dots

# your parameter order/colors (matching your single-species plot)
param_order_raw = [
    'lx_F', 'lx_M',
    'mx_F', 'mx_M',
    'mm',
    'df', 'dm', 'm'
]

parameter_names = [
    'Inbreeding rate',
    'Male dispersal',
    'Female dispersal',
    'Maternal mortality',
    'Male fecundity',
    'Female fecundity',
    'Male survival',
    'Female survival'
][::-1]   # keep your top-to-bottom logic

bar_colors = [
    'salmon',          # Female survival
    'cornflowerblue',  # Male survival
    'salmon',          # Female fecundity
    'cornflowerblue',  # Male fecundity
    'orange',          # Maternal mortality
    'salmon',          # Female dispersal
    'cornflowerblue',  # Male dispersal
    'gray'             # Local mating
]

display_map = {
    'lx_F': 'Female survival',
    'lx_M': 'Male survival',
    'mx_F': 'Female fecundity',
    'mx_M': 'Male fecundity',
    'mm'  : 'Maternal mortality',
    'df'  : 'Female dispersal',
    'dm'  : 'Male dispersal',
    'm'   : 'Inbreeding rate'
}

display_order = parameter_names


# menopausal subset by FILE PREFIX (first part of CSV file name)
menopausal_sp = ['chimps', 'hum', 'resident_KW', 'transient_KW']

# readable dot colors for species
named_colors = [
    "navy", "tab:orange", "seagreen", "red",
    "blueviolet", "sienna", "hotpink", "tab:gray",
    "tab:olive", "tab:cyan", "steelblue", "goldenrod",
    "mediumseagreen", "crimson", "darkorchid"
]

# Distinct marker shapes (one per species, in the same order as species_list)
marker_cycle = ['o', 'X', 'D', 'P', '*', '^', 'v']  # star, diamond, tri up/down, filled plus, point, filled X
species_marker_map = {sp: marker_cycle[i % len(marker_cycle)] for i, sp in enumerate(species_list)}

# ============================================


# -------- LOAD & PREPARE --------
files = sorted(glob.glob(os.path.join(RESULTS_DIR, "*_sobol.csv")))
if not files:
    raise FileNotFoundError(f"No CSVs found in {RESULTS_DIR}")

dfs, file_to_species = [], {}
for f in files:
    df = pd.read_csv(f)
    req = {'species','param','S1','S1_conf','ST','ST_conf'}
    if not req.issubset(df.columns):
        raise ValueError(f"File {f} missing required columns {req}.")
    dfs.append(df)
    file_to_species[f] = str(df['species'].iloc[0])

all_df = pd.concat(dfs, ignore_index=True)
all_df = all_df[all_df['param'].isin(param_order_raw)].copy()
all_df['param_display'] = all_df['param'].map(display_map)
all_df['param_display'] = pd.Categorical(all_df['param_display'],
                                         categories=display_order,
                                         ordered=True)

species_list = sorted(all_df['species'].unique())
species_colors = [named_colors[i % len(named_colors)] for i in range(len(species_list))]
species_color_map = {sp: species_colors[i] for i, sp in enumerate(species_list)}

# menopausal subset -> species names (from filename prefixes)
menopausal_species = {
    file_to_species[f]
    for f in files
    if any(os.path.basename(f).startswith(prefix) for prefix in menopausal_sp)
}
menopausal_species = sorted(menopausal_species)

meno_df = all_df[all_df['species'].isin(menopausal_species)].copy()

# means + standard error (all vs menopausal)
def compute_stats(df):
    """Compute mean, std, n, and sem for S1 and ST per parameter."""
    stats = (df.groupby('param_display', observed=True)
               .agg(S1_mean=('S1', 'mean'),
                    S1_std=('S1', 'std'),
                    S1_n=('S1', 'count'),
                    ST_mean=('ST', 'mean'),
                    ST_std=('ST', 'std'),
                    ST_n=('ST', 'count'))
               .reindex(display_order))
    stats['S1_sem'] = stats['S1_std'] / np.sqrt(stats['S1_n'])
    stats['ST_sem'] = stats['ST_std'] / np.sqrt(stats['ST_n'])
    return stats

stats_all = compute_stats(all_df)
stats_meno = compute_stats(meno_df) if not meno_df.empty else None


# -------- PLOTTING --------
def plot_panel(ax, metric_col, xlabel, xmax=XMAX, in_panel_legend=False, labels=None):
    y = np.arange(len(display_order))
    delta = 0.18  # vertical offset for bar pair

    # --- choose metric columns ---
    if metric_col == 'S1':
        all_mean = stats_all['S1_mean'].values
        all_sem  = stats_all['S1_sem'].values
        if stats_meno is not None:
            meno_mean = stats_meno['S1_mean'].values
            meno_sem  = stats_meno['S1_sem'].values
    else:  # 'ST'
        all_mean = stats_all['ST_mean'].values
        all_sem  = stats_all['ST_sem'].values
        if stats_meno is not None:
            meno_mean = stats_meno['ST_mean'].values
            meno_sem  = stats_meno['ST_sem'].values

    # --- 1. bars (background) ---
    ax.barh(y + delta, all_mean, height=0.34,
            color=bar_colors, edgecolor='none', alpha=0.35)
    if stats_meno is not None:
        ax.barh(y - delta, meno_mean, height=0.34,
                color=bar_colors, edgecolor=bar_colors,
                linewidth=2.5, alpha=0.85)

    # --- 2. dots (species values) ---
    handles_done = set()
    for sp in species_list:
        col = species_color_map[sp]
        sub = all_df[all_df['species'] == sp].set_index('param_display').reindex(display_order)
        xx = sub[metric_col].values

        # ALL dots (upper bar)
        yy_all = y + delta + np.random.uniform(-0.03, 0.03, size=len(y))
        mk = species_marker_map[sp]
        ax.plot(xx, yy_all, linestyle='None', marker=mk, markersize=9,
                color=col, alpha=0.85,
                label=sp if sp not in handles_done else None, markeredgecolor=col, markeredgewidth=0.6)
        
        handles_done.add(sp)

        # MENO dots (lower bar)
        if sp in menopausal_species:
            yy_m = y - delta + np.random.uniform(-0.03, 0.03, size=len(y))
            mk = species_marker_map[sp]
            ax.plot(xx, yy_m, linestyle='None', marker=mk, markersize=9,
                        color=col, alpha=0.85, markeredgecolor=col, markeredgewidth=0.6)

        # optional confidence error bars for dots
        if SHOW_ERRORBARS:
            xerr = sub[f"{metric_col}_conf"].values
            ax.errorbar(xx, yy_all, xerr=xerr, fmt='none', ecolor=col, alpha=0.5, capsize=3, zorder=2.1)
            if sp in menopausal_species:
                ax.errorbar(xx, yy_m, xerr=xerr, fmt='none', ecolor=col, alpha=0.5, capsize=3, zorder=2.1)

    # --- 3. SEM error bars (drawn on top) ---
    ax.errorbar(all_mean, y + delta,
                xerr=all_sem,
                fmt='none',
                ecolor='black',
                elinewidth=2,
                capsize=4,
                zorder=3)
    if stats_meno is not None:
        ax.errorbar(meno_mean, y - delta,
                    xerr=meno_sem,
                    fmt='none',
                    ecolor='black',
                    elinewidth=2,
                    capsize=4,
                    zorder=3)

    # --- axes styling ---
    ax.set_yticks(y)
    ax.set_yticklabels(display_order, fontsize=23)
    ax.set_xlabel(xlabel, fontsize=24)
    ax.tick_params(axis='both', which='major', labelsize=21)
    ax.set_xlim(0, xmax)
    ax.grid(axis='x', linestyle=':', alpha=0.3)
    if in_panel_legend:
        ax.legend(labels, title="Species", loc='upper right', fontsize=18, title_fontsize=19)
        


fig, axes = plt.subplots(1, 2, figsize=FIGSIZE)
plt.subplots_adjust(wspace=0.7)
labels = ['Asian elephants', 'Sperm whales &\nAfrican elephants', 'Amboseli baboons', 'Chimpanzees', 'Humans', 'Resident killer whales', 'Transient killer whales']

plot_panel(axes[0], 'S1', 'First-order sensitivity index', xmax=0.3)#, in_panel_legend=True, labels=labels)
plot_panel(axes[1], 'ST', 'Total sensitivity index')

# Species legend (outside, right)
handles, _ = axes[0].get_legend_handles_labels()
if handles:
    axes[0].legend(handles, labels, title="Species", loc='upper right', fontsize=18, title_fontsize=19)
               # bbox_to_anchor=(0.98, 0.5), )
    axes[1].legend(handles, labels, title="Species", loc='upper right', fontsize=16, title_fontsize=17)

plt.tight_layout(rect=[0, 0, 0.95, 1])  # leave room for legend on the right
plt.show()

# Console helper (file → species → color)
print("\n=== Species / file / color ===")
for f in files:
    sp = file_to_species[f]
    col = species_color_map[sp]
    print(f"{os.path.basename(f):<35} → {sp:<25} → {col}")
print("\nBar encoding:")
print("  Filled bar (color, alpha=0.35) = All species mean")
print("  Outline bar (edge only)        = Menopausal subset mean")
print("  Dots on upper bar              = All species")
print("  Dots on lower bar              = Menopausal species only")


# %% [code]
print("\n=== Species color reference ===")
for file, sp in zip(files, species_list):
    rgb = tuple(c for c in species_color_map[sp])
    print(f"{file:<40}  →  {sp:<25}  color {rgb}")


# %% [markdown]
### Humans

# %% [code]
F_mx_5yr = [0.   , 0.   , 0.143, 1.446, 1.641, 1.88 , 1.552, 1.194, 0.536,  0.   , 0.   , 0.   , 0.   , 0.   , 0.   , 0.   ]
M_mx_5yr = [0.   , 0.   , 0.   , 0.342, 1.143, 1.469, 1.17 , 0.794, 0.747,  0.436, 0.146, 0.172, 0.   , 0.   , 0.   , 0.   ]
F_px = [0.8366    , 0.86755917, 0.95591072, 0.96137215, 0.9856072 ,  0.9841801 , 0.99629057, 0.96773193, 0.99551138, 
            0.98003221,  0.91127177, 0.95744681, 0.88022599, 0.90928541, 1.        ]
M_px = [0.8484    , 0.92880717, 1.        , 1.        , 0.9748731 ,  0.96849779, 1.        , 0.99623656, 0.9719374 , 
            0.9516935 , 0.95186698, 0.98222495, 0.90015601, 0.66169844, 0.70508119]


surv_FM = np.array([F_px, M_px]) 
fec_FM = np.array([F_mx_5yr, M_mx_5yr]) 
human_LM_mp = leslie_(fec_FM, surv_FM)

human_LM = deepcopy(human_LM_mp) # No menopause
d = human_LM.shape[-1] // 2
human_LM[0,9:d-2] = np.array([(d-2-9- i+1) / (d-2-9 + 2*i) for i in range(d-9-2)]) * 0.3

human_interval = 5
human_LM_mp.shape


# %% [markdown]
### Resident/ Transient killer whales

# %% [code]
d = 10
L_mp = np.zeros((2*d, 2*d))
fecundity = np.zeros(d)
fecundity[1:4] = [1, 2, 2]
fecundity_both_sex = np.array([fecundity, fecundity])
survival_f = np.array([3903/5000, 389/400, 2491/2500, 1971/2000, 9641/10000, 4711/5000, 4491/5000, 431/500, 431/500])
survival_m = np.array([3903/5000, 0.9315, 0.9149, 0.8588, 0.8588, 0.8588, 0, 0, 0])
survival_both_sex = np.array([survival_f, survival_m])
KW_L_mp = leslie_(fecundity_both_sex, survival_both_sex)

KW_LM = deepcopy(KW_L_mp) # no menopause
KW_LM[0,4:10] = np.array([2,2,2,1,0.5,0.2]) #based on elephants
competition_factor = 1/1.67
competition_start_age = 4
KW_LM[0, competition_start_age:10] *= competition_factor

KW_interval = 10

# %% [markdown]
### Asian elephants

# %% [code]
fec_F = np.array([0,0,0.28,0.28,0.4,0.4,0.4,0.4,0.4,0.4,0.1,0.1,0])
fec_both_sex = np.array([fec_F, fec_F])
surv_F = np.array([0.76, 0.93, 0.93, 0.86, 0.95, 0.95, 0.95, 0.95, 0.95, 0.95, 0.59, 0.59])
surv_M = np.array([0.64, 0.72, 0.82, 0.82, 0.82, 0.82, 0.82, 0.82, 0.82, 0.82, 0.59, 0.59])
surv_both_sex = np.array([surv_F, surv_M])
AsianEle_LM = leslie_(fec_both_sex, surv_both_sex)

AsianEle_interval = 5

# %% [markdown]
### African elephants/ Sperm whales

# %% [code]
fec_F = np.array([0,0,1,0.95,0.9025,0.857,0.8145,0.7737,0.0774,0.0077,0.0008,0.0001,0,0,0])
fec_M = np.ones(15)
fec_M[:4] = 0
fec_M[12:] = 0
fec_both_sex = np.array([fec_F, fec_M])
surv_F = np.array([0.84,0.99,0.99,0.97,0.99,0.95,0.95,0.95,0.90,0.93,0.74,0.74,0.59,0.47])
surv_M = np.array([0.79,0.96,0.96,0.89,0.91,0.90,0.88,0.88,0.89,0.79,0.87,0.75,0.696,0.6])
surv_both_sex = np.array([surv_F, surv_M])
SW_LM = leslie_(fec_both_sex, surv_both_sex)

SW_interval = 5

# %% [markdown]
### Amboseli baboons

# %% [code]
female_px = [0.794, 0.888, 0.937, 0.969, 0.953, 0.944, 0.948, 0.948, 0.943, 0.923, 0.891, 0.894, 0.942, 
             0.916, 0.87, 0.873, 0.788, 0.633, 0.628, 0.865, 0.746, 0.552, 0.609, 0.417, 0.38, 0.5]
male_px = [0.782, 0.912, 0.934, 0.917, 0.959, 0.943, 0.939, 0.942, 0.931, 0.911, 0.894, 
           0.882, 0.846, 0.83, 0.782, 0.667, 0.625, 0.8, 0.75, 0.333, 0, 0, 0, 0, 0, 0]

female_mx = [0, 0, 0, 0, 0.128, 0.272, 0.275, 0.265, 0.26, 0.224, 0.198, 0.231, 0.262, 0.26, 
             0.284, 0.301, 0.266, 0.217, 0.244, 0.184, 0.24, 0.322, 0.131, 0.214, 0, 0, 0]
male_mx = [0, 0, 0, 0, 0, 0.373, 0.776, 0.779, 0.766, 0.742, 0.722, 0.707, 0.665, 
           0.647, 0.595, 0.475, 0.433, 0.625, 0.586, 0.213, 0, 0, 0, 0, 0, 0, 0]

all_px = np.array([female_px, male_px])
all_mx = np.array([female_mx, male_mx])
baboons_LM = leslie_(all_mx, all_px)

baboons_interval = 1

# %% [markdown]
### Chimpanzees

# %% [code]
male_chimp_mx = [0.0, 0.0, 0.45499999999999996, 1.9089999999999998, 1.086, 1.071, 0.9670000000000001, 1.092, 0.376, 1.236, 0.564, 0.0, 0, 0]
female_chimp_mx = [0.0, 0.053205238, 0.684635425, 1.077490993, 1.069306749, 1.1214980780000001, 0.8998109299999999, 0.793325069, 0.620417416, 0.34493894499999994, 0.051598229, 0.0, 0, 0]
male_chimp_px = [0.81979744, 1.0, 1.0, 0.9506, 0.7560955800000001, 0.8832, 0.8454999999999999, 0.7518119999999999, 0.6246240000000001, 0.6224999999999999, 0.335, 0.0, 0.0]
female_chimp_px = [0.7636, 0.9215, 1.0, 0.98, 0.96, 0.95, 0.9025, 1.0, 0.8835999999999999, 0.8184, 0.71208, 0.6880000000000001, 0.5025000000000001]


print(len(male_chimp_mx), len(female_chimp_mx), len(male_chimp_px), len(female_chimp_px))
chimp_LM = leslie_(np.array([female_chimp_mx, male_chimp_mx]), np.array([female_chimp_px, male_chimp_px]))
chimp_interval = 5

# %% [code]
all_LMs = [human_LM, KW_LM, KW_LM, chimp_LM, AsianEle_LM, SW_LM, SW_LM, baboons_LM]
all_intervals = [human_interval, KW_interval, KW_interval, chimp_interval, AsianEle_interval, SW_interval, SW_interval, baboons_interval]
CI_intervals = ci_intervals(all_LMs, all_intervals)
CI_intervals

# %% [code]
mp_onsets = [35, 50, 50, 40, 45, 74, 74, 26]
# percentages = PRL(mp_onsets, all_LMs, all_intervals)

# %% [code]
RKW_eM, RKW_eB = 22, 39 
TKW_eM, TKW_eB = 32, 46

lX, _ = survival2x(KW_LM)
RKW_lM, RKW_lB = lX[(100 - RKW_eM) // 10], lX[(100 - RKW_eB) // 10]
TKW_lM, TKW_lB = lX[(100 - TKW_eM) // 10], lX[(100 - TKW_eB) // 10]

(lX[(90 - RKW_eM) // 10] / lX[(90 - RKW_eB) // 10]) * (22 / 39)

# %% [code]
# Actual PRRs - https://onlinelibrary.wiley.com/doi/full/10.1002/ece3.3856
african_elephant_PRR = 0.035
# kw_PRR = 0.309
baboon_PRR = 0.036 # (yellow baboons)

# Actual PRR for Asian elephants and humans - https://frontiersinzoology.biomedcentral.com/articles/10.1186/s12983-014-0054-0
asian_elephants_PRR = 0.13

# Actual human PrR - Ellis, S. et al. Postreproductive lifespans are rare in mammals. Ecol. Evol. 8, 2482–2494 (2018).
human_PRR = 0.443 # or 0.51 -> https://onlinelibrary.wiley.com/doi/full/10.1002/ece3.3856 ; also from https://pmc.ncbi.nlm.nih.gov/articles/PMC3212949/ supplementary

# Actual PRR for KW - 
RKW_PrR = (RKW_lM / RKW_lB) * (RKW_eM / RKW_eB)
TKW_PrR = (TKW_lM / TKW_lB) * (TKW_eM / TKW_eB)

# actual_PRRs = [human_PRR, RKW_PrR, TKW_PrR, asian_elephants_PRR, african_elephant_PRR, african_elephant_PRR, baboon_PRR]

# Here, for KW is from A_long_postreproductive_life_span_is_a_shared_trait by Nielsen et al., 2021
#actual_PRRs = [human_PRR, RKW_PrR, TKW_PrR, chimp_PrR, asian_elephants_PRR, african_elephant_PRR, african_elephant_PRR, baboon_PRR]
actual_PRRs = [[0.426, 0.481, 0.439, 0.443, 0.425], [0.309, 0.38], [0.309, 0.48], [0.02, 0.006, 0.195], [0.13, 0.162], [0.035], [0.035], [0.036, 0.005]]
# colors = [['mediumpurple', 'mediumpurple', 'mediumpurple', 'pink', 'royalblue'], ['pink', 'skyblue'], ['pink', 'skyblue'], ['teal', 'thistle'], ['pink'], ['pink'], ['pink', 'mediumpurple']]
colors2 = [['k', 'k', 'k', 'k', 'k'], ['k', 'k'], ['k', 'k'], ['k', 'k', 'k'], ['k', 'k'], ['k'], ['k'], ['k', 'k']]


# %% [code]
# plot_post_reproductive_life(['Humans', 'Resident/ Transient\nkiller whales', 'Asian Elephants', 'African Elephants', 'Sperm whales', 'Amboseli baboons'], percentages, CI_intervals, actual_PRRs)

# %% [code]
PrR_from_model(mp_onsets, all_LMs, all_intervals)

# %% [code]
human_phylopic = mpimg.imread('/Users/Omer/Downloads/295x1024.png')
RKW_phylopic = mpimg.imread('/Users/Omer/Downloads/1024x503.png')
TKW_phylopic = mpimg.imread('/Users/Omer/Downloads/1024x503.png')
AsianE_phylopic = mpimg.imread('/Users/Omer/Downloads/1024x691.png')
AfricanE_phylopic = mpimg.imread('/Users/Omer/Downloads/1024x660.png')
SW_phylopic = mpimg.imread('/Users/Omer/Downloads/1536x518.png')
baboon_phylopic = mpimg.imread('/Users/Omer/Downloads/1024x810.png')
chimp_phylopic = mpimg.imread("/Users/Omer/Documents/DirectPhD/Lab/Yoav/Data/Chimps/chimp_phylopic.png")

phylopics = [human_phylopic, RKW_phylopic, TKW_phylopic, chimp_phylopic, AsianE_phylopic, AfricanE_phylopic, SW_phylopic, baboon_phylopic]
yposition_pics = PrR_from_model(mp_onsets, all_LMs, all_intervals) + 0.1

# %% [code]
PrR_stop_early_allspecies = PrR_from_model(mp_onsets, all_LMs, all_intervals)
PrR_stop_early_allspecies[-3:] = 0

# %% [code]

plot_post_reproductive_life(['Humans', 'Resident\nkiller whales', 'Transient\nkiller whales', 'Chimpanzees', 'Asian Elephants', 'African Elephants', 'Sperm whales', 'Amboseli baboons'], \
                            PrR_stop_early_allspecies, \
                            CI_intervals/2, \
                            actual_PRRs, \
                            colors2, \
                            barcolor='royalblue', \
                            alphabar=0.2, \
                            phylopics=phylopics,
                            yposition_pics=yposition_pics, \
                            zoom=0.055, \
                            ylims=[-0.05, 0.8]) 
# plt.savefig('PrR.png') #, PrR_from_LM([human_LM_mp, KW_L_mp, KW_L_mp, *all_LMs[3:]], all_intervals))


# %% [code]
chimp_PrR_live_long = 0.0102027
hum_PrR_live_long = 0.36319344
RKW_PrR_live_long = TKW_PrR_live_long = 0.49490401
AfE_SW_PrR_live_long = 0
baboon_PrR_live_long = 0
AsianE_PrR_live_long = 0.02118374

PrR_live_long = [hum_PrR_live_long, RKW_PrR_live_long, RKW_PrR_live_long, chimp_PrR_live_long, AsianE_PrR_live_long, AfE_SW_PrR_live_long, AfE_SW_PrR_live_long, baboon_PrR_live_long]

# %% [code]
y_pos_long_live = np.array(PrR_live_long) + np.ones(len(PrR_live_long)) * 0.1
y_pos_long_live[-4] += 0.15
y_pos_long_live[0] += 0.12
y_pos_long_live[3] += 0.15
plot_post_reproductive_life(['Humans', 'Resident\nkiller whales', 'Transient\nkiller whales', 'Chimpanzees', 'Asian Elephants', 'African Elephants', 'Sperm whales', 'Amboseli baboons'], \
                            PrR_live_long, \
                            CI_intervals/2, \
                            actual_PRRs, \
                            colors2, \
                            barcolor='royalblue', \
                            alphabar=0.2, \
                            phylopics=phylopics,
                            yposition_pics=y_pos_long_live, \
                            zoom=0.055, \
                            ylims=[-0.05, 0.8]) 


# %% [markdown]
# Live longer results sum

# %% [markdown]
### Killer whales

# %% [code]
df_RKW = dm_RKW = 0.1 # male, female dispersal occurs at a probability 0.1
m_RKW = 0 # indicative for non-local reproduction as in orcas
nf_RKW = nm_RKW = 3
a_RKW = 10 # number of age classes to view

# %% [code]
r_to_pod_RKW = calc_rcoeff(L_mp=KW_L_mp, m=m_RKW, df=df_RKW, dm=dm_RKW, nf=nf_RKW, nm=nm_RKW, a=a_RKW)
r_to_pod_10yr_RKW = np.concatenate([r_to_pod_RKW[:,::5], (r_to_pod_RKW[:, -1]).reshape(2,1)], axis=1)
r_to_pod_10yr_RKW

# %% [code]
additional_years_after_rep_try_RKW = 6
female_surv_tradeoff_arr_try_RKW = [1 + (1.01-1) / (i + 1) for i in range(additional_years_after_rep_try_RKW)]
female_fec_tradeoff_arr_try_RKW = [1 + (1.025-1) / (i + 1) for i in range(additional_years_after_rep_try_RKW)]
ε = 10**(-60) 
age_classes_RKW = np.arange(10, 61, 10)
RKW_res = different_longevities(KW_L_mp, r_to_pod_10yr_RKW, female_fec_tradeoff_arr_try_RKW, female_surv_tradeoff_arr_try_RKW, additional_years_after_rep_try_RKW, Δmaternal_mortality=1, mx_contribution=np.array([0.95, 0.75]), lx_contribution=np.array([0.8, 0.5]), sex_ratio=[0.6,0.4], return_all=True)

# %% [code]
df_TKW = dm_TKW = 0.35 # male, female dispersal occurs at a probability 0.1
m_TKW = 0.2 # indicative for non-local reproduction as in orcas
nf_TKW = nm_TKW = 3
a_TKW = 10 # number of age classes to view

# %% [code]
r_to_pod_TKW = calc_rcoeff(L_mp=KW_L_mp, m=m_TKW, df=df_TKW, dm=dm_TKW, nf=nf_TKW, nm=nm_TKW, a=a_TKW)
r_to_pod_10yr_TKW = np.concatenate([r_to_pod_TKW[:,::5], (r_to_pod_TKW[:, -1]).reshape(2,1)], axis=1)
r_to_pod_10yr_TKW

# %% [code]
additional_years_after_rep_try_TKW = 6
female_surv_tradeoff_arr_try_TKW = [1 + (1.01-1) / (i + 1) for i in range(additional_years_after_rep_try_TKW)]
female_fec_tradeoff_arr_try_TKW = [1 + (1.015-1) / (i + 1) for i in range(additional_years_after_rep_try_TKW)]
ε = 10**(-60) 
age_classes_RKW = np.arange(10, 61, 10)
TKW_res = different_longevities(KW_L_mp, r_to_pod_10yr_TKW, female_fec_tradeoff_arr_try_TKW, female_surv_tradeoff_arr_try_TKW, additional_years_after_rep_try_TKW, Δmaternal_mortality=1, mx_contribution=np.array([0.95, 0.75]), lx_contribution=np.array([0.8, 0.5]), sex_ratio=[0.6,0.4], return_all=True)

# %% [markdown]
### Humans

# %% [code]
# Dispersal parameters
df_hum = 0.95
dm_hum = 0.05
m_hum = 1
nf_hum = nm_hum = 3
a_hum = 16 # number of age classes to view
r_to_pod_hum = calc_rcoeff(L_mp=human_LM_mp, m=m_hum, df=df_hum, dm=dm_hum, nf=nf_hum, nm=nm_hum, a=a_hum)

r_to_pod_5yr_hum = r_to_pod_hum[:,::6][:, :-1] 
r_to_pod_5yr_hum.shape
r_to_pod_hum.shape

# %% [code]
additional_years_after_rep_try_hum = 7
female_surv_tradeoff_arr_try_hum = [1 + (1.008-1) / (i + 1) for i in range(additional_years_after_rep_try_hum)]
female_fec_tradeoff_arr_try_hum = [1 + (1.012-1) / (i + 1) for i in range(additional_years_after_rep_try_hum)]
hum_res = different_longevities(human_LM_mp, r_to_pod_5yr_hum, female_fec_tradeoff_arr_try_hum, female_surv_tradeoff_arr_try_hum, additional_years_after_rep_try_hum, Δmaternal_mortality=0.97, mx_contribution=np.array([0.75, 0.75]), lx_contribution=np.array([0.67, 0.67]), sex_ratio=[0.5,0.5], return_all=True)


# %% [markdown]
### Sperm whales & African elephants

# %% [code]
additional_years_after_rep_try_SW_Af = 8
female_surv_tradeoff_arr_try_SW_Af = [1 + (1.008-1) / (i + 1) for i in range(additional_years_after_rep_try_SW_Af)]
female_fec_tradeoff_arr_try_SW_Af = [1 + (1.012-1) / (i + 1) for i in range(additional_years_after_rep_try_SW_Af)]


# %% [code]
new_Lmp_SW = extend_LM_both_sexes(SW_LM, num_extra_classes=additional_years_after_rep_try_SW_Af, postpeak_phase_start=9)
new_d_SW = new_Lmp_SW.shape[0] // 2
df_SW = 0.15
dm_SW = 0.995 # male, female dispersal occurs at a probability 0.9, 0
m_SW = 0.01 # indicative for non-local reproduction as in orcas
nf_SW = nm_SW = 3
a_SW = new_d_SW # number of age classes to view
r_to_pod_SW = calc_rcoeff(L_mp=new_Lmp_SW, m=m_SW, df=df_SW, dm=dm_SW, nf=nf_SW, nm=nm_SW, a=a_SW)

r_to_pod_5yr_SW = np.concatenate([r_to_pod_SW[:,::7], (r_to_pod_SW[:, -2:]).reshape(2,2)], axis=1) 
r_to_pod_5yr_SW.shape

# %% [code]
SW_Af_res = different_longevities(new_Lmp_SW, r_to_pod_5yr_SW, female_fec_tradeoff_arr_try_SW_Af, female_surv_tradeoff_arr_try_SW_Af, additional_years_after_rep_try_SW_Af, Δmaternal_mortality=1, mx_contribution=[1.134, 1], lx_contribution=[0.967, 0.967], sex_ratio=[0.5,0.5], return_all=True)


# %% [markdown]
### Asian elephants

# %% [code]
num_extra_classes_AE = 8
new_Lmp_AE = extend_LM_both_sexes(AsianEle_LM, num_extra_classes=num_extra_classes_AE, postpeak_phase_start=9)
df_AE = 0.15 
dm_AE = 0.95 # male, female dispersal occurs at a probability 0.9
m_AE = 0.05 # indicative for non-local reproduction as in orcas
nf_AE = nm_AE = 3
a_AE = new_Lmp_AE.shape[0] // 2 # number of age classes to view

r_to_pod_AE = calc_rcoeff(L_mp=new_Lmp_AE, m=m_AE, df=df_AE, dm=dm_AE, nf=nf_AE, nm=nm_AE, a=a_AE)

r_to_pod_5yr_AE = np.concatenate([r_to_pod_AE[:,::5], (r_to_pod_AE[:, (r_to_pod_AE.shape[-1]-3):]).reshape(2,3)], axis=1)
r_to_pod_5yr_AE.shape

# %% [code]
female_surv_tradeoff_arr_try_AE = [1 + (1.003-1) / (i + 1) for i in range(num_extra_classes_AE)]
female_fec_tradeoff_arr_try_AE = [1 + (1.007-1) / (i + 1) for i in range(num_extra_classes_AE)]

AE_res = different_longevities(new_Lmp_AE, r_to_pod_5yr_AE, female_fec_tradeoff_arr_try_AE, female_surv_tradeoff_arr_try_AE, num_extra_classes_AE, Δmaternal_mortality=1, mx_contribution=[0.812, 1], lx_contribution=[0.91, 0.91], sex_ratio=[0.5,0.5], return_all=True)


# %% [markdown]
### Chimps

# %% [code]
num_extra_classes_chmp = 8
new_Lmp_chmp = extend_LM_both_sexes(chimp_LM, num_extra_classes=num_extra_classes_chmp, postpeak_phase_start=8)
df_chmp = 0.9
dm_chmp = 0
m_chmp = 1
nf_chmp = nm_chmp = 3
a_chmp = 14 + num_extra_classes_chmp # number of age classes to view
r_to_pod_5yr_chimp = calc_rcoeff(L_mp=new_Lmp_chmp, m=m_chmp, df=df_chmp, dm=dm_chmp, nf=nf_chmp, nm=nm_chmp, a=a_chmp)[:,::4][:, :-1]

# %% [code]
female_surv_tradeoff_arr_try_chmp = [1 + (1.0018-1) / (i + 1) for i in range(num_extra_classes_chmp)]
female_fec_tradeoff_arr_try_chmp = [1 + (1.002-1) / (i + 1) for i in range(num_extra_classes_chmp)]
chmp_res = different_longevities(new_Lmp_chmp, r_to_pod_5yr_chimp, female_fec_tradeoff_arr_try_chmp, female_surv_tradeoff_arr_try_chmp, num_extra_classes_chmp, Δmaternal_mortality=1, mx_contribution=[1, 0.95], lx_contribution=[0.95, 0.93], sex_ratio=[0.5,0.5], return_all=True)


# %% [markdown]
### Amboseli baboons

# %% [code]
num_extra_classes_baboon = 20
new_Lmp_bab = extend_LM_both_sexes(baboons_LM, num_extra_classes=num_extra_classes_baboon, postpeak_phase_start=14)
new_d_bab = new_Lmp_bab.shape[0] // 2

# %% [code]
df_bab = 0.15
dm_bab = 0.95
m_bab = 0.33420596902319577
nf_bab = nm_bab = 3
a_bab = new_d_bab # number of age classes to view
r_to_group_bab = calc_rcoeff(L_mp=new_Lmp_bab, m=m_bab, df=df_bab, dm=dm_bab, nf=nf_bab, nm=nm_bab, a=a_bab)
r_to_group_5yr_bab = np.concatenate([r_to_group_bab[:,::5], (r_to_group_bab[:, -1]).reshape(2,1)], axis=1)

# %% [code]
female_surv_tradeoff_arr_try_bab = [1 + (1.008-1) / (i + 1) for i in range(num_extra_classes_baboon)]
female_fec_tradeoff_arr_try_bab = [1 + (1.012-1) / (i + 1) for i in range(num_extra_classes_baboon)]
bab_res = different_longevities(new_Lmp_bab, r_to_group_5yr_bab, female_fec_tradeoff_arr_try_bab, female_surv_tradeoff_arr_try_bab, num_extra_classes_baboon, Δmaternal_mortality=1, mx_contribution=[0.99, 0.97], lx_contribution=[0.98, 0.97], sex_ratio=[0.5,0.5], return_all=True)


# %% [markdown]
### Plotting all together - live long

# %% [code]
def fig_panel_labels(axes, letters=None, uppercase=True,
                     xcoord=-0.17, ycoord=0.92, panel_label_size=None):
    import string
    from matplotlib.font_manager import FontProperties
    import numpy as np
    import matplotlib.pyplot as plt

    # Convert rcParam to a numeric size
    if panel_label_size is None:
        fp = FontProperties(size=plt.rcParams['axes.titlesize'])
        panel_label_size = fp.get_size() * 1.3

    # Generate default letters if needed
    if letters is None:
        letters = string.ascii_uppercase if uppercase else string.ascii_lowercase

    axes_arr = np.array(axes, dtype=object)
    flat_axes = list(axes_arr.flat)
    n_axes = len(flat_axes)

    # Truncate / slice letters to the number of axes
    letters = list(letters[:n_axes])

    nrows, ncols = axes_arr.shape if axes_arr.ndim == 2 else (1, n_axes)

    # --- handle xcoord ---
    if hasattr(xcoord, "__len__") and not isinstance(xcoord, str):
        xcoord_list = list(xcoord)
        if axes_arr.ndim == 2 and len(xcoord_list) == ncols:
            # per-column xcoords
            xs = []
            for i in range(n_axes):
                col = i % ncols
                xs.append(xcoord_list[col])
        elif len(xcoord_list) == n_axes:
            # per-panel xcoords
            xs = xcoord_list
        else:
            raise ValueError("xcoord must be a scalar, length ncols, or length n_axes")
    else:
        xs = [xcoord] * n_axes

    # --- handle ycoord (mirror logic, but per-row) ---
    if hasattr(ycoord, "__len__") and not isinstance(ycoord, str):
        ycoord_list = list(ycoord)
        if axes_arr.ndim == 2 and len(ycoord_list) == nrows:
            # per-row ycoords
            ys = []
            for i in range(n_axes):
                row = i // ncols
                ys.append(ycoord_list[row])
        elif len(ycoord_list) == n_axes:
            # per-panel ycoords
            ys = ycoord_list
        else:
            raise ValueError("ycoord must be a scalar, length nrows, or length n_axes")
    else:
        ys = [ycoord] * n_axes

    # --- annotate ---
    annots = []
    for ax, letter, xc, yc in zip(flat_axes, letters, xs, ys):
        annots.append(
            ax.annotate(
                letter, (xc, yc),
                xycoords='axes fraction',
                fontsize=panel_label_size
            )
        )
    return annots


# %% [code]
# rBC_arr = [rBC_RKW, rBC_TKW, rBC_hum, rBC_SW_Af, rBC_SW_Af, rBC_AE, rBC_chmp, rBC_bab]
# C_arr = [C_RKW, C_TKW, C_hum, C_SW_Af, C_SW_Af, C_AE, C_chmp, C_bab]
# B_arr = [B_RKW, B_TKW, B_hum, B_SW_Af, B_SW_Af, B_AE, B_chmp, B_bab]
# rxB_arr = [rxB_RKW, rxB_TKW, rxB_hum, rxB_SW_Af, rxB_SW_Af, rxB_AE, rxB_chmp, rxB_bab]
species_res = [RKW_res, TKW_res, hum_res, chmp_res, AE_res, SW_Af_res, SW_Af_res, bab_res]
new_all_intervals = [KW_interval, KW_interval, human_interval, chimp_interval, AsianEle_interval, SW_interval, SW_interval, baboons_interval]
new_phylopics = [RKW_phylopic, TKW_phylopic, human_phylopic, chimp_phylopic, AsianE_phylopic, AfricanE_phylopic, SW_phylopic, baboon_phylopic]

phylopositions = [(16, 0.022), (16, 0.022), (7, 0.0037), (36, -0.0006), (35, -0.00075), (35, -0.0055), (34, -0.0055), (18, -0.00019)]
phylozoom = [0.095, 0.095, 0.095, 0.15, 0.095, 0.1, 0.095, 0.08]

def plot_all_species_lambdas(species_res, phylopics, phylopositions, phylozoom, all_intervals, x_label, fs=22, lw=2, saveby=None, letters=None, uppercase=True, xcoord=-0.25, ycoord=0.92, panel_label_size=24):
    fig, axes = plt.subplots(4,2, figsize=(18,25))#, sharey=True, sharex=True)
    plt.subplots_adjust(wspace=0.45, hspace=0.15)

    for i, curr_spcs in enumerate(species_res):
        row = i//2 
        col = i%2 if i!=1 else 1
        ax = axes[row, col]
        age_classes = np.arange(1 * all_intervals[i], curr_spcs[0].shape[-1] * all_intervals[i] + 1, all_intervals[i])
        rBC, C, B, rxB = curr_spcs
        ax.axhline(0,c='k')
        ax.plot(age_classes, np.sum(rBC, axis=0), label='λ', marker='o', c='#E9765B', lw=lw, markersize=7)
        ax.plot(age_classes, np.sum(C, axis=0), label='C', ls='-', c='#FFC98B', lw=lw)
        ax.plot(age_classes, np.sum(B, axis=0), label='B', ls='-', c='#A5CAD2', lw=lw)
        ax.plot(age_classes, np.sum(rxB, axis=0), label='r*B', ls='-', c='#478BA2', lw=lw)
        add_image(ax, phylopics[i], xy=phylopositions[i], zoom=phylozoom[i])
        if np.any(np.sum(rBC, axis=0) > 0):
            ax.axvline(all_intervals[i]*(np.where(np.sum(rBC, axis=0)>ε)[0][0] +1), c='k', ls=':', label='first positive')
 
        if row == len(species_res)//2 -1:
            ax.set_xlabel(x_label, fontsize=fs+2)
        if col == 0:
            ax.set_ylabel('Fitness effect over lifespan', fontsize=fs+2)
            ax.yaxis.set_label_coords(-0.25, 0.5)
        ax.tick_params(axis='both', which='major', labelsize=fs)
        ax.set_xticks(age_classes if all_intervals[i] != 1 else np.arange(0, curr_spcs[0].shape[-1] * all_intervals[i] + 1, all_intervals[i] * 4))
        # ax.set_xlim(age_classes[0], age_classes[-1] + 3)
        # sns.despine()


    fig_panel_labels(axes, letters=letters, uppercase=uppercase, xcoord=xcoord, ycoord=ycoord, panel_label_size=panel_label_size)
    if saveby is not None:
        plt.savefig('{}.png'.format(saveby))
    plt.show()

# %% [code]
xcoords = [-0.38, -0.3]      # same for both rows, per column
ycoords = [0.97, 0.97, 0.97, 0.97]   # same for both columns, per row

plot_all_species_lambdas(species_res, new_phylopics, phylopositions, phylozoom, new_all_intervals, 'Added post-reproductive years', fs=22, saveby='livelong_res', xcoord=xcoords, ycoord=ycoords)

# %% [markdown]
# Stop early results sum

# %% [markdown]
### Killer whales

# %% [code]
RKW_res_Stop = different_onset_ages(KW_LM, r_to_pod_10yr_RKW, Δmaternal_mortality=1, mx_contribution=np.array([0.95, 0.75]), lx_contribution=np.array([0.8, 0.5]), sex_ratio=[0.6,0.4], return_all=True, ε=10^(-50))

# %% [code]
TKW_res_Stop = different_onset_ages(KW_LM, r_to_pod_10yr_TKW, Δmaternal_mortality=1, mx_contribution=np.array([0.95, 0.75]), lx_contribution=np.array([0.8, 0.5]), sex_ratio=[0.6,0.4], return_all=True, ε=10^(-50))

# %% [markdown]
### Humans

# %% [code]
hum_res_Stop = different_onset_ages(human_LM, r_to_pod_5yr_hum, Δmaternal_mortality=0.97, mx_contribution=np.array([0.75, 0.75]), lx_contribution=np.array([0.67, 0.67]), sex_ratio=[0.5,0.5], return_all=True, ε=10^(-50))


# %% [markdown]
### Sperm whales & African elephants

# %% [code]
SW_Af_res_Stop = different_onset_ages(SW_LM, r_to_pod_5yr_SW, Δmaternal_mortality=1, mx_contribution=[1.134, 1], lx_contribution=[0.967, 0.967], sex_ratio=[0.5,0.5], return_all=True, ε=10^(-50))


# %% [markdown]
### Asian elephants

# %% [code]
AE_res_Stop = different_onset_ages(AsianEle_LM, r_to_pod_5yr_AE, Δmaternal_mortality=1, mx_contribution=[0.812, 1], lx_contribution=[0.91, 0.91], sex_ratio=[0.5,0.5], return_all=True, ε=10^(-50))


# %% [markdown]
### Chimps

# %% [code]
chmp_res_Stop = different_onset_ages(chimp_LM, r_to_pod_5yr_chimp, Δmaternal_mortality=1, mx_contribution=[1, 0.95], lx_contribution=[0.95, 0.93], sex_ratio=[0.5,0.5], return_all=True, ε=10^(-50))


# %% [markdown]
### Amboseli baboons

# %% [code]
bab_res_Stop = different_onset_ages(baboons_LM, r_to_group_5yr_bab, Δmaternal_mortality=1, mx_contribution=[0.99, 0.97], lx_contribution=[0.98, 0.97], sex_ratio=[0.5,0.5], return_all=True, ε=10^(-50))


# %% [code]
def plot_all_species_lambdas_stop(species_res, phylopics, phylopositions, phylozoom, all_intervals, x_label, fs=22, lw=2, saveby=None, letters=None, uppercase=True, xcoord=-0.25, ycoord=0.92, panel_label_size=24, begin_plotting_idx=None):
    fig, axes = plt.subplots(4,2, figsize=(18,25))#, sharey=True, sharex=True)
    plt.subplots_adjust(wspace=0.45, hspace=0.15)

    if begin_plotting_idx is None:
        begin_plotting_idx = [0] * len(species_res)
    
    for i, curr_spcs in enumerate(species_res):
        row = i//2 
        col = i%2 if i!=1 else 1
        ax = axes[row, col]
        rBC, C, B, rxB = curr_spcs
        start_age = begin_plotting_idx[i]
        age_classes = np.arange(start_age * all_intervals[i], rBC.shape[-1] * all_intervals[i], all_intervals[i])
        print(age_classes, start_age)
        ax.axhline(0,c='k')
        ax.plot(age_classes, np.sum(rBC, axis=0)[start_age:], label='λ', marker='o', c='#E9765B', lw=lw, markersize=7)
        ax.plot(age_classes, np.sum(C, axis=0)[start_age:], label='C', ls='-', c='#FFC98B', lw=lw)
        ax.plot(age_classes, np.sum(B, axis=0)[start_age:], label='B', ls='-', c='#A5CAD2', lw=lw)
        ax.plot(age_classes, np.sum(rxB, axis=0)[start_age:], label='r*B', ls='-', c='#478BA2', lw=lw)
        add_image(ax, phylopics[i], xy=phylopositions[i], zoom=phylozoom[i])
        if np.any(np.sum(rBC, axis=0) > 0):
            ax.axvline(all_intervals[i]*(np.where(np.sum(rBC, axis=0)>ε)[0][0]), c='k', ls=':', label='first positive')
 
        if row == len(species_res)//2 -1:
            ax.set_xlabel(x_label, fontsize=fs+2)
        if col == 0:
            ax.set_ylabel('Fitness effect over lifespan', fontsize=fs+2)
            ax.yaxis.set_label_coords(-0.25, 0.5)
        ax.tick_params(axis='both', which='major', labelsize=fs)
        ax.set_xticks(age_classes if all_intervals[i] != 1 else np.arange(start_age, curr_spcs[0].shape[-1] * all_intervals[i] + 1, all_intervals[i] * 4))
        # ax.set_xlim(age_classes[0], age_classes[-1] + 3)


    fig_panel_labels(axes, letters=letters, uppercase=uppercase, xcoord=xcoord, ycoord=ycoord, panel_label_size=panel_label_size)
    if saveby is not None:
        plt.savefig('{}.png'.format(saveby))
    plt.show()

# %% [code]
species_res_Stop = [RKW_res_Stop, TKW_res_Stop, hum_res_Stop, chmp_res_Stop, AE_res_Stop, SW_Af_res_Stop, SW_Af_res_Stop, bab_res_Stop]
begin_plotting_idx = [3, 3, 6, 6, 6, 7, 6, 6, 9]
phylopositions_stop = [(83, -0.045), (83, -0.045), (73, 0.002), (62, -0.00038), (57, -0.00087), (65, -0.00015), (63, -0.00075), (24.5, -0.0006)]
xcoords = [-0.38, -0.27]      # same for both rows, per column
ycoords = [0.98, 0.98, 0.98, 0.98]   # same for both columns, per row
plot_all_species_lambdas_stop(species_res_Stop, new_phylopics, phylopositions_stop, phylozoom, new_all_intervals, 'Menopause age', fs=22, saveby='stopearly_res', xcoord=xcoords, ycoord=ycoords, begin_plotting_idx=begin_plotting_idx)


# %% [code]
male_symbol = mpimg.imread('/Users/Omer/Downloads/pTodqKkpc-3982000212.png')
female_symbol = mpimg.imread('/Users/Omer/Downloads/8czr7L9Ki-3456542989.png')

# %% [code]


# %% [markdown]
## Relatedness Values 

# %% [markdown]
### Sperm whales & African elephants

# %% [code]
df_SW = 0.15
dm_SW = 0.995 # male, female dispersal occurs at a probability 0.9, 0
m_SW = 0.01 # indicative for non-local reproduction as in orcas
nf_SW = nm_SW = 3
a_SW = SW_LM.shape[0]//2 # number of age classes to view
r_to_pod_SW = calc_rcoeff(SW_LM, m=m_SW, df=df_SW, dm=dm_SW, nf=nf_SW, nm=nm_SW, a=a_SW)

r_to_pod_5yr_SW_origin = np.concatenate([r_to_pod_SW[:,::7], (r_to_pod_SW[:, -2:]).reshape(2,2)], axis=1) 
r_to_pod_5yr_SW_origin.shape

# %% [markdown]
### Asian elephants

# %% [code]
df_AE = 0.15 
dm_AE = 0.95 # male, female dispersal occurs at a probability 0.9
m_AE = 0.05 # indicative for non-local reproduction as in orcas
nf_AE = nm_AE = 3
a_AE = AsianEle_LM.shape[0] // 2 # number of age classes to view

r_to_pod_AE = calc_rcoeff(L_mp=AsianEle_LM, m=m_AE, df=df_AE, dm=dm_AE, nf=nf_AE, nm=nm_AE, a=a_AE)

r_to_pod_5yr_AE_origin = np.concatenate([r_to_pod_AE[:,::5], (r_to_pod_AE[:, (r_to_pod_AE.shape[-1]-3):]).reshape(2,3)], axis=1)
r_to_pod_5yr_AE_origin.shape

# %% [markdown]
### Chimps

# %% [code]
df_chmp = 0.9
dm_chmp = 0
m_chmp = 1
nf_chmp = nm_chmp = 3
a_chmp = chimp_LM.shape[0] // 2 # number of age classes to view
r_to_pod_5yr_chimp = calc_rcoeff(L_mp=chimp_LM, m=m_chmp, df=df_chmp, dm=dm_chmp, nf=nf_chmp, nm=nm_chmp, a=a_chmp)[:,::4][:, :-1]

# %% [markdown]
### Amboseli baboons

# %% [code]
df_bab = 0.15
dm_bab = 0.95
m_bab = 0.33420596902319577
nf_bab = nm_bab = 3
a_bab = baboons_LM.shape[0]//2 # number of age classes to view
r_to_group_bab = calc_rcoeff(L_mp=baboons_LM, m=m_bab, df=df_bab, dm=dm_bab, nf=nf_bab, nm=nm_bab, a=a_bab)
r_to_group_5yr_bab = np.concatenate([r_to_group_bab[:,::5], (r_to_group_bab[:, -1]).reshape(2,1)], axis=1)

# %% [code]
import seaborn as sns

# %% [code]
def relatedness_plots(species_relatedness, phylopics, phylopositions, phylozoom, sex_symb_pstns, sex_symb_zoom, all_intervals, sex_ratio_per_spcs=None, lw=2, fs=22, saveby=None, letters=None, uppercase=True, xcoord=-0.25, ycoord=0.92, panel_label_size=27, begin_plotting_idx=None):
    fig, axes = plt.subplots(2, 4, figsize=(30,12), sharey=True)
    plt.subplots_adjust(wspace=0.15, hspace=0.15)

    if sex_ratio_per_spcs is None:
        sex_ratio_per_spcs = [(0.5, 0.5)] * len(species_relatedness)
        
    for i, curr_relatedness in enumerate(species_relatedness):
        row = i//4
        col = i%4 
        ax = axes[row, col]
        rf, rm = curr_relatedness
        age_classes = np.arange(0 * all_intervals[i], rf.shape[-1] * all_intervals[i], all_intervals[i])
        # print(age_classes, start_age)
        sex_ratio = sex_ratio_per_spcs[i]
        ravg = sex_ratio[0]*rf+sex_ratio[1]*rm
        # ax.axhline(0,c='k')
        
        ax.plot(age_classes, rf, label='$r_f$', c='salmon', lw=lw, marker='o')
        ax.plot(age_classes, rm, label='$r_m$', c='cornflowerblue', lw=lw, marker='o')
        ax.plot(age_classes, ravg, c='#FFC98B', label='$r_{avg}$', lw=lw, ls='--')# marker='o')
        
        add_image(ax, male_symbol, xy=sex_symb_pstns[i][1], zoom=sex_symb_zoom[i][1])
        add_image(ax, female_symbol, xy=sex_symb_pstns[i][0], zoom=sex_symb_zoom[i][0])
        add_image(ax, phylopics[i], xy=phylopositions[i], zoom=phylozoom[i])
        
        if row == len(species_res)//4 -1:
            ax.set_xlabel('Female age', fontsize=fs+2)
        if col == 0:
            ax.set_ylabel('Relatedness', fontsize=fs+2)
            ax.yaxis.set_label_coords(-0.15, 0.5)
            
        ax.tick_params(axis='both', which='major', labelsize=fs)
        ax.set_xticks(np.arange(0, rf.shape[-1] * all_intervals[i], 10))# if all_intervals[i] != 1 else np.arange(start_age, species_relatedness[0].shape[-1] * all_intervals[i] + 1, all_intervals[i] * 4))
        sns.despine()
    fig_panel_labels(axes, letters=letters, uppercase=uppercase, xcoord=xcoord, ycoord=ycoord, panel_label_size=panel_label_size)
    if saveby is not None:
        plt.savefig('{}.png'.format(saveby))
    plt.show()

# %% [code]
species_relatedness = [r_to_pod_10yr_RKW, r_to_pod_10yr_TKW, r_to_pod_5yr_hum,  r_to_pod_5yr_chimp, r_to_pod_5yr_AE, r_to_pod_5yr_SW_origin, r_to_pod_5yr_SW_origin, r_to_group_5yr_bab]
sex_ratio_per_species = [(0.6, 0.4)] * 2 + [(0.5, 0.5)] * 6
phylopositions_relatedness = [(15, 0.29), (15, 0.29), (3.5, 0.26), (4.5, 0.28), (12, 0.285), (10, 0.285), (15, 0.29), (4, 0.28)]
new_phylopics = [RKW_phylopic, TKW_phylopic, human_phylopic,  chimp_phylopic, AsianE_phylopic, AfricanE_phylopic, SW_phylopic, baboon_phylopic]

sex_symb_pstns = [[(0.5, 0.215), (6, 0.075)], 
                  [(0.5, 0.17), (5, 0.06)],
                  [(13, 0.035), (13, 0.18)],
                  [(11, 0.05), (11, 0.165)],
                  [(6, 0.165), (6, 0.03)],
                  [(5, 0.16), (5, 0.035)],
                  [(6, 0.165), (6, 0.03)],
                  [(2, 0.045), (2, 0.175)]
                 ]
sex_symb_zoom = [(0.011, 0.014)] * 8

# %% [code]
xcoords = [-0.28, -0.1, -0.1, -0.1]      # same for both rows, per column
ycoords = [0.98, 0.98]  
phylozoom = [0.13, 0.13, 0.12, 0.15, 0.095, 0.11, 0.095, 0.095]

relatedness_plots(species_relatedness, new_phylopics, phylopositions_relatedness, phylozoom, sex_symb_pstns, sex_symb_zoom, new_all_intervals, sex_ratio_per_spcs=sex_ratio_per_species, ycoord=ycoords, xcoord=xcoords, saveby='relatedness_allspecies', lw=3)


# %% [code]


# %% [code]
```
