## Supplementary Code for "Evolution of Menopause and the Mama’s Boy Hypothesis": AmB_stop_early.ipynb

```
# %% [markdown]
# Baboons (stop early)

# %% [code]
import numpy as np
from copy import deepcopy
import matplotlib.pyplot as plt
import pandas as pd
from io import StringIO
import matplotlib.image as mpimg
from SALib.sample import saltelli
from SALib.analyze import sobol

# %% [code]
import io, os, sys, types

# %% [code]
from IPython import get_ipython
from nbformat import read
from IPython.core.interactiveshell import InteractiveShell

# %% [code]
def find_notebook(fullname, path=None):
    """find a notebook, given its fully qualified name and an optional path

    This turns "foo.bar" into "foo/bar.ipynb"
    and tries turning "Foo_Bar" into "Foo Bar" if Foo_Bar
    does not exist.
    """
    name = fullname.rsplit('.', 1)[-1]
    if not path:
        path = ['']
    for d in path:
        nb_path = os.path.join(d, name + ".ipynb")
        if os.path.isfile(nb_path):
            return nb_path
        # let import Notebook_Name find "Notebook Name.ipynb"
        nb_path = nb_path.replace("_", " ")
        if os.path.isfile(nb_path):
            return nb_path

# %% [code]
class NotebookLoader(object):
    """Module Loader for Jupyter Notebooks"""
    def __init__(self, path=None):
        self.shell = InteractiveShell.instance()
        self.path = path

    def load_module(self, fullname):
        """import a notebook as a module"""
        path = find_notebook(fullname, self.path)

        print ("importing Jupyter notebook from %s" % path)

        # load the notebook object
        with io.open(path, 'r', encoding='utf-8') as f:
            nb = read(f, 4)


        # create the module and add it to sys.modules
        # if name in sys.modules:
        #    return sys.modules[name]
        mod = types.ModuleType(fullname)
        mod.__file__ = path
        mod.__loader__ = self
        mod.__dict__['get_ipython'] = get_ipython
        sys.modules[fullname] = mod

        # extra work to ensure that magics that would affect the user_ns
        # actually affect the notebook module's ns
        save_user_ns = self.shell.user_ns
        self.shell.user_ns = mod.__dict__

        try:
          for cell in nb.cells:
            if cell.cell_type == 'code':
                # transform the input to executable Python
                code = self.shell.input_transformer_manager.transform_cell(cell.source)
                # run the code in themodule
                exec(code, mod.__dict__)
        finally:
            self.shell.user_ns = save_user_ns
        return mod

# %% [code]
class NotebookFinder(object):
    """Module finder that locates Jupyter Notebooks"""
    def __init__(self):
        self.loaders = {}

    def find_module(self, fullname, path=None):
        nb_path = find_notebook(fullname, path)
        if not nb_path:
            return

        key = path
        if path:
            # lists aren't hashable
            key = os.path.sep.join(path)

        if key not in self.loaders:
            self.loaders[key] = NotebookLoader(path)
        return self.loaders[key]

# %% [code]
sys.meta_path.append(NotebookFinder())

# %% [code]
from Johnstone_Cant import IBD_age_0, no_age_rs, IBD_no_age, IBD_older_age_v2, rff_age_v2

# %% [code]
from project_functions import survival2x, leslie_, Px, mx, age_distribution, tot_rBC_all_donor_life, tot_rBC, survival_effect, fecundity_effect, reproductive_value, different_onset_ages, latex_array, add_image, model, LM_to_block_tables_v2, save_sobol_results, calc_rcoeff

# %% [code]
def convert_to_surv(lx):
    div_by = 1/lx[:-1]
    surv_values = deepcopy(lx)
    surv_values[1:] *= div_by
    return surv_values

# %% [markdown]
![image.png](attachment:78b2e25f-8fd2-43c9-a5c0-2aee11b46c41.png)

# %% [code]
# Values from "Primate life histories and socioecology"
female_px = [0.794, 0.888, 0.937, 0.969, 0.953, 0.944, 0.948, 0.948, 0.943, 0.923, 0.891, 0.894, 0.942, 
             0.916, 0.87, 0.873, 0.788, 0.633, 0.628, 0.865, 0.746, 0.552, 0.609, 0.417, 0.38, 0.5]
male_px = [0.782, 0.912, 0.934, 0.917, 0.959, 0.943, 0.939, 0.942, 0.931, 0.911, 0.894, 
           0.882, 0.846, 0.83, 0.782, 0.667, 0.625, 0.8, 0.75, 0.333, 0, 0, 0, 0, 0, 0]

female_mx = [0, 0, 0, 0, 0.128, 0.272, 0.275, 0.265, 0.26, 0.224, 0.198, 0.231, 0.262, 0.26, 
             0.284, 0.301, 0.266, 0.217, 0.244, 0.184, 0.24, 0.322, 0.131, 0.214, 0, 0, 0]
male_mx = [0, 0, 0, 0, 0, 0.373, 0.776, 0.779, 0.766, 0.742, 0.722, 0.707, 0.665, 
           0.647, 0.595, 0.475, 0.433, 0.625, 0.586, 0.213, 0, 0, 0, 0, 0, 0, 0]

len(female_px), len(female_mx), len(male_px), len(male_mx)

# %% [code]
all_px = np.array([female_px, male_px])
all_mx = np.array([female_mx, male_mx])

# %% [code]
baboons_LM = leslie_(all_mx, all_px)
baboons_LM.shape

# %% [markdown]
### Males disperse while females remain in their natal groups, in natural pipulations of East africa.  (Bronikowski et. al, 2002). 
They migrate at age 7 so they can only receive help up until that age. They get no reproductive assist up to that age because they start reproducing at age 7. (JAY F. STORZ ET AL., 2002).
m = 0.5 because "50% of males leave the natal group without mating" (JAY F. STORZ ET AL., 2002)
df = 0.1, dm = 0.9 because males disperse. 

# %% [code]


# %% [code]
def calc_m(fecundity, age_start, age_end, survival=None):
    if survival is None:
        survival = np.ones(len(fecundity))
    if type(fecundity) != np.ndarray:
        fecundity = np.array(fecundity)
    return sum(fecundity[age_start : age_end] * survival[age_start : age_end]) / sum(fecundity * survival)

# %% [code]
local_mating_fraction = calc_m(male_mx, 5, 11, survival=lx_baboons[1,:])
local_mating_fraction * 0.5

# %% [code]
# Dispersal parameters
df = 0.15
dm = 0.95
m = local_mating_fraction * 0.5
# m=0
nf = nm = 3
a = 27 # number of age classes to view

mu = np.zeros((2,a)) # array for survival probabilities per age-group, where each age group is in 10-year intervals
for i in range(a-1):
    mu[:,i] = baboons_LM[i+1,i], baboons_LM[a+i+1,a+i]
rf_to_f, rm_to_f, rf_to_m, rm_to_m, avg_rf = rff_age_v2(df, dm, nf, nm, m, a, 1-mu)
r_to_group = np.array([rf_to_f, rm_to_f])

r_to_group_5yr = r_to_group[:,::5] # np.concatenate([r_to_pod[:,::4], (r_to_pod[:, -4:-1]).reshape(2,3)], axis=1)
# r_to_pod_5yr
# r_to_pod[:,::3].shape
r_to_group_5yr.shape

# %% [code]
age_classes = np.arange(0,a)
age_classes

# %% [code]
male_symbol = mpimg.imread('/Users/Omer/Downloads/pTodqKkpc-3982000212.png')
female_symbol = mpimg.imread('/Users/Omer/Downloads/8czr7L9Ki-3456542989.png')
baboon_phylopic = mpimg.imread('/Users/Omer/Downloads/1024x810.png')

# %% [code]
lw, fs = 2.5, 17

fig, ax = plt.subplots(figsize=(8,5))
ax.plot(age_classes[:], r_to_group_5yr[0,:], label='$r_f$', c='salmon', lw=lw, marker='o')
ax.plot(age_classes[:], r_to_group_5yr[1,:], label='$r_m$', c='cornflowerblue', lw=lw, marker='o')
ax.plot(age_classes[:], 0.5*r_to_group_5yr[0,:]+0.5*r_to_group_5yr[1,:], c='#FFC98B', label='$r_{avg}$', lw=lw, ls='--')
ax.set_xlabel('Female age', fontsize=fs+2)
ax.set_ylabel('Relatedness coefficient', fontsize=fs+2)
ax.tick_params(axis='both', which='major', labelsize=fs)
# ax.set_title('Relatedness Coefficient of Female to Pod Throughout Life (1-year intervals)', fontsize=fs)
# ax.legend(fontsize=fs-3, loc='lower right')
# ax.axhline(0, c='k')

add_image(ax, female_symbol, xy=(3, 0.18), zoom=0.012)
add_image(ax, baboon_phylopic, xy=(23.5, 0.045), zoom=0.08)
plt.savefig('fig_R_baboons.png', dpi=400, bbox_inches = 'tight')

plt.show()

# %% [code]


# %% [code]
plt.subplots_adjust(wspace=0.3)
ε = 10**(-50) 
rbc, c, b, rb = different_onset_ages(baboons_LM, r_to_group_5yr, Δmaternal_mortality=1, mx_contribution=[0.99, 0.97], lx_contribution=[0.98, 0.97], return_all=True, ε=ε, sex_ratio=[0.5,0.5], max_male_help=9) 
rbc, c, b, rb = rbc[:,8:], c[:,8:], b[:,8:], rb[:,8:]

fig, ax = plt.subplots(figsize=(8,5))
ax.axhline(0,c='k')
ax.plot(age_classes[8:], np.sum(rbc, axis=0), label='λ', marker='o', c='#E9765B', lw=lw, markersize=7)
ax.plot(age_classes[8:], np.sum(c, axis=0), label='C', ls='-', c='#FFC98B', lw=lw)
ax.plot(age_classes[8:], np.sum(b, axis=0), label='B', ls='-', c='#A5CAD2', lw=lw)
ax.plot(age_classes[8:], np.sum(rb, axis=0), label='r*B', ls='-', c='#478BA2', lw=lw)

# first_pos
ax.axvline((np.where(np.sum(rbc, axis=0)>ε)[0][0]+8), c='k', ls=':', label='first positive')
ax.set_xlabel('Menopause age', fontsize=fs+2)
ax.set_ylabel('Benefit and cost over lifespan', fontsize=fs+2)
# ax.set_ylabel('λ', fontsize=fs+2)
ax.tick_params(axis='both', which='major', labelsize=fs)
ax.set_xlim(5,28)
ax.set_xticks(np.arange(6,28,2))
plt.savefig('fig2Aa.png', dpi=400, bbox_inches = 'tight')
add_image(ax, baboon_phylopic, xy=(25.2, -5.7e-05), zoom=0.08)
plt.show()

np.sum(rbc, axis=0)


# %% [code]


# %% [code]
from SALib.sample import saltelli 
from SALib.analyze import sobol
problem = {
    'num_vars': 8,
    'names': ['lx_F', 'lx_M', 'mx_F', 'mx_M', 'mm',
              'df', 'dm', 'm'],
    'bounds': [
        [0, 1],  # lx_F
        [0, 1],  # lx_M
        [0, 1],  # mx_F
        [0, 1],  # mx_M
        [0, 1],  # mm
        [0, 1],  # df  (example biologically reasonable range)
        [0, 1],  # dm
        [0, 1],    # m
    ]
}


# %% [code]
def model2(x, L_no_effects, onset=5, sex_ratio=[0.5, 0.5], max_male_help=None, max_female_help=None, adjusted_age_dist_rho=True, ε=10**(-20), age_dist=None, ρ=None):

    # life-history modifiers
    lx_cont = np.array([x[0], x[1]])
    mx_cont = np.array([x[2], x[3]])
    mm = x[4]

    # dispersal & mating parameters
    df, dm, m = x[5], x[6], x[7]

    # constants (kept fixed for Sobol)
    nf = nm = 3

    # ---- relatedness calculation (generalized) ----
    r_to_pod = calc_rcoeff(L_mp=L_no_effects, m=m, df=df, dm=dm, nf=nf, nm=nm, a=L_no_effects.shape[0]//2)

    r_to_pod_5yr = r_to_group[:,::5]
    # ----------------------------------------------

    if adjusted_age_dist_rho:
        lamb = different_onset_ages(
            L_no_effects,
            r_to_pod_5yr,
            Δmaternal_mortality=mm,
            lx_contribution=lx_cont,
            mx_contribution=mx_cont,
            sex_ratio=[0.6, 0.4],
            ε=ε,
            return_all=True
        )[0]
    else:
        lamb = different_onset_ages(
            L_no_effects,
            r_to_pod_5yr,
            Δmaternal_mortality=mm,
            lx_contribution=lx_cont,
            mx_contribution=mx_cont,
            sex_ratio=[0.6, 0.4],
            ε=ε,
            return_all=True,
            age_dist=age_distribution(L_mp, ρ),
            ρ=ρ
        )[0]

    return np.sum(lamb, axis=0)[onset]


# %% [code]
N = 1024

param_values = saltelli.sample(problem, N, calc_second_order=True)
Y = np.array([model2(x, baboons_LM, onset=26) for x in param_values])
Si = sobol.analyze(problem, Y)


# %% [code]
parameter_names = [
    'Female survival',
    'Male survival',
    'Female fecundity',
    'Male fecundity',
    'Maternal mortality',
    'Female dispersal',
    'Male dispersal',
    'Inbreeding rate'
]
param_names = [
    'lx_F', 'lx_M',
    'mx_F', 'mx_M',
    'mm',
    'df', 'dm', 'm'
]

# %% [code]
save_sobol_results(
    species_name="baboon",
    Si=Si,
    param_names=param_names,
    outdir="sa_results"
)


# %% [code]


# %% [markdown]
## Bibliography
> Kappeler, Peter M; Pereira, Michael Eric. 2003. Primate life histories and socioecology. Chicago : University of Chicago Press
> https://archive.org/details/primatelifehisto0000unse/page/74/mode/2up
>
> Anne M. Bronikowski, Susan C. Alberts, Jeanne Altmann, Craig Packer, K. Dee Carey, and Marc Tatar. 2002. The aging baboon: Comparative demography in a non-human primate, PNAS, 99-14. 
> https://www.pnas.org/doi/full/10.1073/pnas.142675599
>
> JAY F. STORZ, UMA RAMAKRISHNAN, AND SUSAN C. ALBERTS. 2002. GENETIC EFFECTIVE SIZE OF A WILD PRIMATE POPULATION: INFLUENCE OF CURRENT AND HISTORICAL DEMOGRAPHY. https://swh.princeton.edu/~baboon/publications/Alberts%20&%20Altmann%20J.%201995.pdf
```
