## Supplementary Code for "Evolution of Menopause and the Mama’s Boy Hypothesis": AsE_live_long.ipynb

```
# %% [markdown]
### Imports

# %% [code]
import numpy as np
from pandas import DataFrame, read_csv
import matplotlib.pyplot as plt
import seaborn as sns
from copy import deepcopy
from scipy.linalg import leslie
%matplotlib inline
import matplotlib.image as mpimg

# %% [code]
import io, os, sys, types

# %% [code]
from IPython import get_ipython
from nbformat import read
from IPython.core.interactiveshell import InteractiveShell

# %% [code]
def find_notebook(fullname, path=None):
    """find a notebook, given its fully qualified name and an optional path

    This turns "foo.bar" into "foo/bar.ipynb"
    and tries turning "Foo_Bar" into "Foo Bar" if Foo_Bar
    does not exist.
    """
    name = fullname.rsplit('.', 1)[-1]
    if not path:
        path = ['']
    for d in path:
        nb_path = os.path.join(d, name + ".ipynb")
        if os.path.isfile(nb_path):
            return nb_path
        # let import Notebook_Name find "Notebook Name.ipynb"
        nb_path = nb_path.replace("_", " ")
        if os.path.isfile(nb_path):
            return nb_path

# %% [code]
class NotebookLoader(object):
    """Module Loader for Jupyter Notebooks"""
    def __init__(self, path=None):
        self.shell = InteractiveShell.instance()
        self.path = path

    def load_module(self, fullname):
        """import a notebook as a module"""
        path = find_notebook(fullname, self.path)

        print ("importing Jupyter notebook from %s" % path)

        # load the notebook object
        with io.open(path, 'r', encoding='utf-8') as f:
            nb = read(f, 4)


        # create the module and add it to sys.modules
        # if name in sys.modules:
        #    return sys.modules[name]
        mod = types.ModuleType(fullname)
        mod.__file__ = path
        mod.__loader__ = self
        mod.__dict__['get_ipython'] = get_ipython
        sys.modules[fullname] = mod

        # extra work to ensure that magics that would affect the user_ns
        # actually affect the notebook module's ns
        save_user_ns = self.shell.user_ns
        self.shell.user_ns = mod.__dict__

        try:
          for cell in nb.cells:
            if cell.cell_type == 'code':
                # transform the input to executable Python
                code = self.shell.input_transformer_manager.transform_cell(cell.source)
                # run the code in themodule
                exec(code, mod.__dict__)
        finally:
            self.shell.user_ns = save_user_ns
        return mod

# %% [code]
class NotebookFinder(object):
    """Module finder that locates Jupyter Notebooks"""
    def __init__(self):
        self.loaders = {}

    def find_module(self, fullname, path=None):
        nb_path = find_notebook(fullname, path)
        if not nb_path:
            return

        key = path
        if path:
            # lists aren't hashable
            key = os.path.sep.join(path)

        if key not in self.loaders:
            self.loaders[key] = NotebookLoader(path)
        return self.loaders[key]

# %% [code]
sys.meta_path.append(NotebookFinder())

# %% [code]
from Johnstone_Cant import IBD_age_0, no_age_rs, IBD_no_age, IBD_older_age_v2, rff_age_v2

# %% [code]
from project_functions import sex_weight, survival2x, leslie_, Px, mx, age_distribution, tot_rBC_all_donor_life, tot_rBC, survival_effect, fecundity_effect, reproductive_value, different_longevities, latex_array, add_image, evaluate_survival_fecundity_tradeoffs, extend_LM_both_sexes,plot_lambda_zero_contours, PrR_from_model 


# %% [markdown]
### Original LM

# %% [code]
fec_F = np.array([0,0,0.28,0.28,0.4,0.4,0.4,0.4,0.4,0.4,0.1,0.1,0])
fec_both_sex = np.array([fec_F, fec_F])
surv_F = np.array([0.76, 0.93, 0.93, 0.86, 0.95, 0.95, 0.95, 0.95, 0.95, 0.95, 0.59, 0.59])
surv_M = np.array([0.64, 0.72, 0.82, 0.82, 0.82, 0.82, 0.82, 0.82, 0.82, 0.82, 0.59, 0.59])
surv_both_sex = np.array([surv_F, surv_M])
LM = leslie_(fec_both_sex, surv_both_sex)
d =LM.shape[0] // 2
d

# %% [markdown]
### Extended survival LM

# %% [code]
num_extra_classes = 8
new_Lmp = extend_LM_both_sexes(LM, num_extra_classes=num_extra_classes, postpeak_phase_start=9)
new_d = new_Lmp.shape[0] // 2
new_d

# %% [code]
(new_Lmp.shape[0] // 2) * 5 - 40
np.where(new_Lmp[0, :new_d] != 0)

# %% [code]
PrR_from_model(menopause_onset=[(new_Lmp.shape[0] // 2) * 5 - 40], leslie_matrices=[new_Lmp], age_intervals=[5])

# %% [code]
df = 0.15 
dm = 0.95 # male, female dispersal occurs at a probability 0.9
m = 0.05 # indicative for non-local reproduction as in orcas
nf = nm = 3
a = new_d # number of age classes to view
mu = np.zeros((2,a)) # array for survival probabilities per age-group, where each age group is in 10-year intervals
for i in range(a-1):
    mu[:,i] = new_Lmp[i+1,i], new_Lmp[a+i+1,a+i]
rf_to_f, rm_to_f, rf_to_m, rm_to_m, avg_rf = rff_age_v2(df, dm, nf, nm, m, a, 1-mu)
r_to_pod = np.array([rf_to_f, rm_to_f])

r_to_pod_5yr = np.concatenate([r_to_pod[:,::5], (r_to_pod[:, (r_to_pod.shape[-1]-3):]).reshape(2,3)], axis=1)
r_to_pod_5yr.shape

# %% [code]
male_symbol = mpimg.imread('/Users/Omer/Downloads/pTodqKkpc-3982000212.png')
female_symbol = mpimg.imread('/Users/Omer/Downloads/8czr7L9Ki-3456542989.png')
AsianE_phylopic = mpimg.imread('/Users/Omer/Downloads/1024x691.png')

# %% [code]
age_classes = np.arange(0,61 + num_extra_classes*5,5)

lw, fs = 2.5, 17

fig, ax = plt.subplots(figsize=(9,5))
ax.plot(age_classes[:], r_to_pod_5yr[0,:], label='$r_f$', c='salmon', lw=lw, marker='o')
ax.plot(age_classes[:], r_to_pod_5yr[1,:], label='$r_m$', c='cornflowerblue', lw=lw, marker='o')
ax.plot(age_classes[:], 0.5*r_to_pod_5yr[0,:]+0.5*r_to_pod_5yr[1,:], c='#FFC98B', label='$r_{avg}$', lw=lw, ls='--')# marker='o')
ax.set_xlabel('Female age', fontsize=fs+2)
ax.set_ylabel('Relatedness coefficient', fontsize=fs+2)
ax.tick_params(axis='both', which='major', labelsize=fs)
add_image(ax, male_symbol, xy=(5, 0.0265), zoom=0.0155)
add_image(ax, female_symbol, xy=(5, 0.165), zoom=0.012)
add_image(ax, AsianE_phylopic, xy=(95, 0.04), zoom=0.08)
# ax.set_title('Relatedness Coefficient of Female to Pod Throughout Life (5-year intervals)', fontsize=fs)
# ax.legend(fontsize=fs-3, loc='best', bbox_to_anchor=(0.5, 0., 0.5, 0.5))
plt.savefig('AE_r.png')

plt.show()

# %% [code]
additional_years_after_rep = num_extra_classes
surv_range = np.linspace(1.1, 1, 50)  # Survival trade-off values
fec_range = np.linspace(1.1, 1, 50)   # Fecundity trade-off values
female_fec_tradeoff_arr = np.linspace(1.5,1,additional_years_after_rep)
female_surv_tradeoff_arr = np.linspace(1.15,1,additional_years_after_rep)
mx_contribution=[0.812, 1]
lx_contribution=[0.91, 0.91]

# %% [code]
results = evaluate_survival_fecundity_tradeoffs(
    surv_range,
    fec_range,
    additional_years_after_rep,
    new_Lmp, 
    r_to_pod_5yr,
    female_fec_tradeoff_arr,
    female_surv_tradeoff_arr,
    Δmaternal_mortality=1,
    mx_contribution=mx_contribution, 
    lx_contribution=lx_contribution,
    sex_ratio=[0.5, 0.5],
    return_all=True)

# %% [code]
fig, ax = plot_lambda_zero_contours(
    results, surv_range, fec_range, additional_years_after_rep,
    years_per_step=5, 
    fallback_when_no_zero='max',
    highlight_year=additional_years_after_rep*5,                    # <— bold + shade below the 30y line
    shade_below_highlight=True, figsize=(12,10), non_meno_col='salmon', meno_col='darkseagreen', fs=20, alpha=1, skip=[5,15,25,35])
add_image(ax, AsianE_phylopic, xy=(1.0485, 1.033), zoom=0.25)

ax.set_xlim(1.0, 1.06)
ax.set_ylim(1.0, 1.04)
ax.set_xlabel('Fecundity Trade-off', fontsize=34)
ax.set_ylabel('Survival Trade-off', fontsize=0)
ax.tick_params(axis='x', which='major', labelsize=32)
ax.tick_params(axis='y', which='major', labelsize=0)
plt.savefig('LL_AsianE.png', bbox_inches="tight")
plt.show()

# %% [code]
X, Y = np.meshgrid(fec_range, surv_range)  # X = fecundity (horizontal), Y = survival (vertical)

# %% [code]
fig, axes = plt.subplots(2, 4, figsize=(30, 13))
plt.subplots_adjust(wspace=0.3, hspace=0.35)

letters = ['A', 'B', 'C', 'D', 'E', 'F', 'G', 'H']

for additional_year in range(additional_years_after_rep):
    row = additional_year // 4
    col = additional_year % 4
    print(row,col)
    ax = axes[row, col]
    
    Z = np.array([[r['rBC'][additional_year] for r in results if r['surv'] == s and r['fec'] == f]
                  for s in surv_range for f in fec_range])
    Z = Z.reshape(len(surv_range), len(fec_range))
    
    vmax = np.max([abs(np.max(Z)), abs(np.min(Z))])
    vmin = -vmax

    c = ax.pcolormesh(X, Y, Z, shading='auto', cmap='coolwarm', vmin=vmin, vmax=vmax)

    if np.min(Z) < 0 and np.max(Z) > 0:
        contour = ax.contour(X, Y, Z, levels=[0], colors='black', linewidths=1.5)
        ax.clabel(contour, fmt={0: "0"}, inline=True, fontsize=12)

    ax.set_xlabel("Fecundity Trade-off", fontsize=16)
    ax.set_ylabel("Survival Trade-off", fontsize=16)
    ax.set_title(f"{letters[additional_year]}) λ at {(additional_year+1)*5} Post-Reproductive Years", fontsize=17)
    ax.tick_params(axis='both', labelsize=14)

# # Hide unused subplots (if any)
# for i in range(7, 9):  # Remaining empty subplots in 3x3 grid
#     fig.delaxes(axes.flatten()[i])

# # Add colorbar
cbar = fig.colorbar(c, ax=axes.ravel().tolist(), shrink=0.9)
cbar.set_label('λ', fontsize=15)
cbar.ax.tick_params(labelsize=13)

plt.show()


# %% [code]
additional_years_after_rep_try = 8
female_surv_tradeoff_arr_try = [1 + (1.003-1) / (i + 1) for i in range(additional_years_after_rep_try)]
female_fec_tradeoff_arr_try = [1 + (1.007-1) / (i + 1) for i in range(additional_years_after_rep_try)]

print("female_surv_tradeoff_arr =", female_surv_tradeoff_arr_try)
print("female_fec_tradeoff_arr =", female_fec_tradeoff_arr_try)

# %% [code]
rBC, C, B, rxB = different_longevities(new_Lmp, r_to_pod_5yr, female_fec_tradeoff_arr_try, female_surv_tradeoff_arr_try, additional_years_after_rep_try, Δmaternal_mortality=1, mx_contribution=mx_contribution, lx_contribution=lx_contribution, sex_ratio=[0.5,0.5], return_all=True)
np.sum(rBC, axis=0)

# %% [code]
ε = 10**(-60) 
age_classes = np.arange(5, num_extra_classes*5+1, 5)
# rBC, C, B, rxB = different_longevities(L_mp, r_to_pod_10yr, female_fec_tradeoff_arr_try, female_surv_tradeoff_arr_try, additional_years_after_rep_try, Δmaternal_mortality=1, mx_contribution=np.array([0.95, 0.75]), lx_contribution=np.array([0.8, 0.5]), sex_ratio=[0.6,0.4], return_all=True)

fig, ax = plt.subplots(figsize=(10,7))
ax.axhline(0,c='k')
ax.plot(age_classes, np.sum(rBC, axis=0), label='λ', marker='o', c='#E9765B', lw=lw, markersize=7)
ax.plot(age_classes, np.sum(C, axis=0), label='C', ls='-', c='#FFC98B', lw=lw)
ax.plot(age_classes, np.sum(B, axis=0), label='B', ls='-', c='#A5CAD2', lw=lw)
ax.plot(age_classes, np.sum(rxB, axis=0), label='r*B', ls='-', c='#478BA2', lw=lw)
add_image(ax, AsianE_phylopic, xy=(35, -0.00075), zoom=0.09)
ax.axvline(5*(np.where(np.sum(rBC, axis=0)>0)[0][0] +1), c='k', ls=':', label='1st positive')
ax.set_xlabel('Added post-reproductive years', fontsize=fs+2)
ax.set_ylabel('Benefit and cost over lifespan', fontsize=fs+2)
# ax.set_ylabel('λ', fontsize=fs+2)
ax.tick_params(axis='both', which='major', labelsize=fs)
# ax.set_xlim(0,91)
ax.set_xticks(np.arange(5,num_extra_classes*5+1,5))

plt.show()


# %% [code]
```
