## Supplementary Code for "Evolution of Menopause and the Mama’s Boy Hypothesis": hum_stop_early.ipynb

```
# %% [markdown]
# Humans (stop early)

# %% [code]
import numpy as np
from copy import deepcopy
import matplotlib.pyplot as plt
import pandas as pd
from io import StringIO
import matplotlib.image as mpimg

# %% [code]
import io, os, sys, types

# %% [code]
from IPython import get_ipython
from nbformat import read
from IPython.core.interactiveshell import InteractiveShell

# %% [code]
def find_notebook(fullname, path=None):
    """find a notebook, given its fully qualified name and an optional path

    This turns "foo.bar" into "foo/bar.ipynb"
    and tries turning "Foo_Bar" into "Foo Bar" if Foo_Bar
    does not exist.
    """
    name = fullname.rsplit('.', 1)[-1]
    if not path:
        path = ['']
    for d in path:
        nb_path = os.path.join(d, name + ".ipynb")
        if os.path.isfile(nb_path):
            return nb_path
        # let import Notebook_Name find "Notebook Name.ipynb"
        nb_path = nb_path.replace("_", " ")
        if os.path.isfile(nb_path):
            return nb_path

# %% [code]
class NotebookLoader(object):
    """Module Loader for Jupyter Notebooks"""
    def __init__(self, path=None):
        self.shell = InteractiveShell.instance()
        self.path = path

    def load_module(self, fullname):
        """import a notebook as a module"""
        path = find_notebook(fullname, self.path)

        print ("importing Jupyter notebook from %s" % path)

        # load the notebook object
        with io.open(path, 'r', encoding='utf-8') as f:
            nb = read(f, 4)


        # create the module and add it to sys.modules
        # if name in sys.modules:
        #    return sys.modules[name]
        mod = types.ModuleType(fullname)
        mod.__file__ = path
        mod.__loader__ = self
        mod.__dict__['get_ipython'] = get_ipython
        sys.modules[fullname] = mod

        # extra work to ensure that magics that would affect the user_ns
        # actually affect the notebook module's ns
        save_user_ns = self.shell.user_ns
        self.shell.user_ns = mod.__dict__

        try:
          for cell in nb.cells:
            if cell.cell_type == 'code':
                # transform the input to executable Python
                code = self.shell.input_transformer_manager.transform_cell(cell.source)
                # run the code in themodule
                exec(code, mod.__dict__)
        finally:
            self.shell.user_ns = save_user_ns
        return mod

# %% [code]
class NotebookFinder(object):
    """Module finder that locates Jupyter Notebooks"""
    def __init__(self):
        self.loaders = {}

    def find_module(self, fullname, path=None):
        nb_path = find_notebook(fullname, path)
        if not nb_path:
            return

        key = path
        if path:
            # lists aren't hashable
            key = os.path.sep.join(path)

        if key not in self.loaders:
            self.loaders[key] = NotebookLoader(path)
        return self.loaders[key]

# %% [code]
sys.meta_path.append(NotebookFinder())

# %% [code]
from Johnstone_Cant import IBD_age_0, no_age_rs, IBD_no_age, IBD_older_age_v2, rff_age_v2

# %% [code]
from project_functions import leslie_, Px, mx, age_distribution, tot_rBC_all_donor_life, tot_rBC, survival_effect, fecundity_effect, reproductive_value, different_onset_ages, latex_array, add_image, LM_to_block_tables_v1, save_sobol_results, calc_rcoeff

# %% [code]
def convert_to_surv(lx):
    div_by = 1/lx[:-1]
    surv_values = deepcopy(lx)
    surv_values[1:] *= div_by
    return surv_values

# %% [code]
F_mx = np.array([0,0,0,0,0,0,0,0,0,0,0,0,0.005988024,0.012269939,0.025641026,0.088435374,0.090909091,0.136690647,0.172932331,0.279069767,0.25984252,0.31092437,0.276785714,0.333333333,
                 0.183673469,0.32967033,0.333333333,0.303797468,0.24,0.271428571,0.353846154,0.360655738,0.224137931,0.254901961,0.395833333,0.234042553,0.391304348,0.260869565,0.25, 0.255813953,
                 0.214285714,0.214285714,0.225,0.305555556,0.125,0.193548387,0.037037037,0.038461538,0.041666667,0,0,0,0,0,0,0,0,0,0,0,0,0,0,0,0,0,0,0,0,0,0,0,0,0,0,0,0,0,0,0,0,0,0,0,0,0,0,0,0,0])
F_mx.shape
F_mx_5yr = np.array([sum(F_mx[i:i+5]) for i in range(0,91,5)])

# %% [code]
AcheF_mx = pd.read_csv('/Users/Omer/Documents/DirectPhD/Lab/Yoav/Human data/Ache Life History The Ecology and Demography of a Foraging People (Hill & Hurtado)/Age-Specific_Fertility_Data_of_Ache_Women_on_Reservations.csv')
AcheM_mx = pd.read_csv('/Users/Omer/Documents/DirectPhD/Lab/Yoav/Human data/Ache Life History The Ecology and Demography of a Foraging People (Hill & Hurtado)/Age-specific_Fertility_Data_Males_Reservations.csv')

# %% [code]
AcheF_mx.iloc[32:36, 3] = [0.217, 0, 0.176, 0]

# %% [code]
F_mx = np.zeros(75)
F_mx[9:47] = np.array(AcheF_mx.iloc[:, 3])
F_mx_5yr = np.array([sum(F_mx[i:i+5]) for i in range(0,76,5)])
F_mx_5yr

# %% [code]
AcheM_mx.iloc[-2:, 3] = [0.143, 0.029]

# %% [code]
M_mx = np.zeros(75)
M_mx[14:60] = np.array(AcheM_mx.iloc[:, 3])
M_mx_5yr = np.array([sum(M_mx[i:i+5]) for i in range(0,76,5)])
M_mx_5yr

# %% [code]
plt.plot(F_mx_5yr)
plt.plot(M_mx_5yr)

# %% [code]
AcheF_lx = pd.read_csv('/Users/Omer/Documents/DirectPhD/Lab/Yoav/Human data/Ache Life History The Ecology and Demography of a Foraging People (Hill & Hurtado)/Females_Survivorship_and_Expected_Life.csv')
AcheM_lx = pd.read_csv('/Users/Omer/Documents/DirectPhD/Lab/Yoav/Human data/Ache Life History The Ecology and Demography of a Foraging People (Hill & Hurtado)/Males_Survivorship_and_Expected_Life.csv')

# %% [code]
AcheF_lx

# %% [code]
AcheF_lx_5yr = np.array([np.mean(np.array(AcheF_lx['Survivorship (s_x)'])[i:i+5]) for i in range(0,76,5)])
AcheF_px_5yr = convert_to_surv(AcheF_lx_5yr)

AcheM_lx_5yr = np.array([np.mean(np.array(AcheM_lx['Survivorship (s_x)'])[i:i+5]) for i in range(0,76,5)])
AcheM_px_5yr = convert_to_surv(AcheM_lx_5yr)

# print(AcheF_px_5yr)
plt.plot(AcheF_lx_5yr)
plt.plot(AcheM_lx_5yr)

# %% [code]
AcheF_px_5yr.shape, AcheM_px_5yr.shape, F_mx_5yr.shape

# %% [code]
F_px = AcheF_px_5yr[:-1]
M_px = AcheM_px_5yr[:-1]

# %% [code]
np.array([F_mx_5yr, M_mx_5yr]), np.array([F_px, M_px])

# %% [code]
hum_LM = leslie_(np.array([F_mx_5yr, M_mx_5yr]), np.array([F_px, M_px]))

# %% [code]
# Dispersal parameters
df = 0.95
dm = 0.05
m = 1
nf = nm = 3
a = 16 # number of age classes to view

mu = np.zeros((2,a)) # array for survival probabilities per age-group, where each age group is in 10-year intervals
for i in range(a-1):
    mu[:,i] = hum_LM[i+1,i], hum_LM[a+i+1,a+i]
rf_to_f, rm_to_f, rf_to_m, rm_to_m, avg_rf = rff_age_v2(df, dm, nf, nm, m, a, 1-mu)
r_to_pod = np.array([rf_to_f, rm_to_f])

r_to_pod_5yr = r_to_pod[:,::6][:, :-1] # np.concatenate([r_to_pod[:,::4], (r_to_pod[:, -4:-1]).reshape(2,3)], axis=1)
# r_to_pod_5yr
# r_to_pod[:,::3].shape
r_to_pod_5yr.shape

# %% [code]
age_classes = np.arange(0,a*5,5)
age_classes

# %% [code]
L_no_effects = deepcopy(hum_LM)
d = hum_LM.shape[-1] // 2
L_no_effects[0,9:d-2] = np.array([(d-2-9- i+1) / (d-2-9 + 2*i) for i in range(d-9-2)]) * 0.3
L_no_effects[0,:d]
L_no_effects[0,d-6:d]*=1/1.67

# %% [code]
strng = latex_array(L_no_effects[d:,d:])
new_strng = strng.replace('.0000', '').replace('00 ', '')
new_strng

# %% [markdown]
$$
\begin{bmatrix}0 & 0 & 0 & 0.3420 & 1.1430 & 1.4690 & 1.17& 0.7940 & 0.7470 & 0.4360 & 0.1460 & 0.1720 & 0 & 0 & 0 & 0 \\\\0.8366 & 0 & 0 & 0 & 0 & 0 & 0 & 0 & 0 & 0 & 0 & 0 & 0 & 0 & 0 & 0 \\\\0 & 0.8676 & 0 & 0 & 0 & 0 & 0 & 0 & 0 & 0 & 0 & 0 & 0 & 0 & 0 & 0 \\\\0 & 0 & 0.9559 & 0 & 0 & 0 & 0 & 0 & 0 & 0 & 0 & 0 & 0 & 0 & 0 & 0 \\\\0 & 0 & 0 & 0.9614 & 0 & 0 & 0 & 0 & 0 & 0 & 0 & 0 & 0 & 0 & 0 & 0 \\\\0 & 0 & 0 & 0 & 0.9856 & 0 & 0 & 0 & 0 & 0 & 0 & 0 & 0 & 0 & 0 & 0 \\\\0 & 0 & 0 & 0 & 0 & 0.9842 & 0 & 0 & 0 & 0 & 0 & 0 & 0 & 0 & 0 & 0 \\\\0 & 0 & 0 & 0 & 0 & 0 & 0.9963 & 0 & 0 & 0 & 0 & 0 & 0 & 0 & 0 & 0 \\\\0 & 0 & 0 & 0 & 0 & 0 & 0 & 0.9677 & 0 & 0 & 0 & 0 & 0 & 0 & 0 & 0 \\\\0 & 0 & 0 & 0 & 0 & 0 & 0 & 0 & 0.9955 & 0 & 0 & 0 & 0 & 0 & 0 & 0 \\\\0 & 0 & 0 & 0 & 0 & 0 & 0 & 0 & 0 & 0.98& 0 & 0 & 0 & 0 & 0 & 0 \\\\0 & 0 & 0 & 0 & 0 & 0 & 0 & 0 & 0 & 0 & 0.9113 & 0 & 0 & 0 & 0 & 0 \\\\0 & 0 & 0 & 0 & 0 & 0 & 0 & 0 & 0 & 0 & 0 & 0.9574 & 0 & 0 & 0 & 0 \\\\0 & 0 & 0 & 0 & 0 & 0 & 0 & 0 & 0 & 0 & 0 & 0 & 0.8802 & 0 & 0 & 0 \\\\0 & 0 & 0 & 0 & 0 & 0 & 0 & 0 & 0 & 0 & 0 & 0 & 0 & 0.9093 & 0 & 0 \\\\0 & 0 & 0 & 0 & 0 & 0 & 0 & 0 & 0 & 0 & 0 & 0 & 0 & 0 & 1 & 0 \\\\\end{bmatrix}
$$

# %% [code]
male_symbol = mpimg.imread('/Users/Omer/Downloads/pTodqKkpc-3982000212.png')
female_symbol = mpimg.imread('/Users/Omer/Downloads/8czr7L9Ki-3456542989.png')
human_phylopic = mpimg.imread('/Users/Omer/Downloads/295x1024.png')

# %% [code]
lw, fs = 2.5, 17

fig, ax = plt.subplots(figsize=(8,5))
ax.plot(age_classes[:], r_to_pod_5yr[0,:], label='$r_f$', c='salmon', lw=lw, marker='o')
ax.plot(age_classes[:], r_to_pod_5yr[1,:], label='$r_m$', c='cornflowerblue', lw=lw, marker='o')
# ax.plot(age_classes[:], 0.5*r_to_pod_5yr[0,:]+0.5*r_to_pod_5yr[1,:], c='#FFC98B', label='$r_{avg}$', lw=lw, marker='o')
ax.plot(age_classes[:], 0.5*r_to_pod_5yr[0,:]+0.5*r_to_pod_5yr[1,:], c='#FFC98B', label='$r_{avg}$', lw=lw, ls='--')
ax.set_xlabel('Female age', fontsize=fs+2)
ax.set_ylabel('Relatedness Coefficient', fontsize=fs+2)
ax.tick_params(axis='both', which='major', labelsize=fs)
ax.set_xticks(np.arange(0,76,10))
# ax.set_title('Relatedness Coefficient of Female to Group Throughout Life (5-year intervals)', fontsize=fs)
# ax.legend(fontsize=fs-3)
ax.axhline(0, c='k')
add_image(ax, male_symbol, xy=(13, 0.18), zoom=0.015)
add_image(ax, female_symbol, xy=(13, 0.035), zoom=0.012)
add_image(ax, human_phylopic, xy=(71, 0.06), zoom=0.08)
plt.savefig('Hum_r.png', dpi=400, bbox_inches = 'tight')

plt.show()

# %% [code]
age_classes_to_plot = age_classes[6:]
r_to_pod_5yr_original = r_to_pod_5yr

# %% [code]


# %% [code]
plt.subplots_adjust(wspace=0.3)
ε = 10**(-60) 
rbc, c, b, rb = different_onset_ages(L_no_effects, r_to_pod_5yr_original, Δmaternal_mortality=0.97, mx_contribution=[0.75, 0.75], lx_contribution=[0.67, 0.67], return_all=True, ε=ε, sex_ratio=[0.5,0.5], max_female_help=6) 
rbc, c, b, rb = rbc[:,6:], c[:,6:], b[:,6:], rb[:,6:]

fig, ax = plt.subplots(figsize=(8,5))
ax.axhline(0,c='k')
ax.plot(age_classes_to_plot, np.sum(rbc, axis=0), label='λ', marker='o', c='#E9765B', lw=lw, markersize=7)
ax.plot(age_classes_to_plot, np.sum(c, axis=0), label='C', ls='-', c='#FFC98B', lw=lw)
ax.plot(age_classes_to_plot, np.sum(b, axis=0), label='B', ls='-', c='#A5CAD2', lw=lw)
ax.plot(age_classes_to_plot, np.sum(rb, axis=0), label='r*B', ls='-', c='#478BA2', lw=lw)
ax.plot(age_classes_to_plot, np.sum(rbc, axis=0), label='λ', marker='o', c='#E9765B', lw=lw, markersize=7)
# first_pos
ax.axvline(5*(np.where(np.sum(rbc, axis=0)>ε)[0][0]+6), c='k', ls=':', label='first positive')
ax.set_xlabel('Menopause age', fontsize=fs+2)
ax.set_ylabel('Benefit and cost over lifespan', fontsize=fs+2)
ax.tick_params(axis='both', which='major', labelsize=fs)
# ax.set_xlim(30,91)
ax.set_xticks(np.arange(30,76,5))
# ax.set_yscale('log')
add_image(ax, human_phylopic, xy=(73, 0.002), zoom=0.1)
plt.savefig('fig2A.png', dpi=400, bbox_inches = 'tight')
# ax.set_yscale('symlog')

plt.show()

print(np.sum(rbc, axis=0).max()/10**(-7))
np.sum(rbc, axis=0)

# %% [markdown]
## SA

# %% [code]
from SALib.sample import saltelli 
from SALib.analyze import sobol
problem = {
    'num_vars': 8,
    'names': ['lx_F', 'lx_M', 'mx_F', 'mx_M', 'mm',
              'df', 'dm', 'm'],
    'bounds': [
        [0, 1],  # lx_F
        [0, 1],  # lx_M
        [0, 1],  # mx_F
        [0, 1],  # mx_M
        [0, 1],  # mm
        [0, 1],  # df  (example biologically reasonable range)
        [0, 1],  # dm
        [0, 1],    # m
    ]
}


# %% [code]
def model2(x, L_no_effects, onset=5, sex_ratio=[0.5, 0.5], max_male_help=None, max_female_help=None, adjusted_age_dist_rho=True, ε=10**(-20), age_dist=None, ρ=None):

    # life-history modifiers
    lx_cont = np.array([x[0], x[1]])
    mx_cont = np.array([x[2], x[3]])
    mm = x[4]

    # dispersal & mating parameters
    df, dm, m = x[5], x[6], x[7]

    # constants (kept fixed for Sobol)
    nf = nm = 3

    # ---- relatedness calculation (generalized) ----
    r_to_pod = calc_rcoeff(L_mp=L_no_effects, m=m, df=df, dm=dm, nf=nf, nm=nm, a=L_no_effects.shape[0]//2)

    r_to_pod_5yr = r_to_pod_5yr = r_to_pod[:,::6][:, :-1]
    # ----------------------------------------------

    if adjusted_age_dist_rho:
        lamb = different_onset_ages(
            L_no_effects,
            r_to_pod_5yr,
            Δmaternal_mortality=mm,
            lx_contribution=lx_cont,
            mx_contribution=mx_cont,
            sex_ratio=[0.6, 0.4],
            ε=ε,
            return_all=True
        )[0]
    else:
        lamb = different_onset_ages(
            L_no_effects,
            r_to_pod_5yr,
            Δmaternal_mortality=mm,
            lx_contribution=lx_cont,
            mx_contribution=mx_cont,
            sex_ratio=[0.6, 0.4],
            ε=ε,
            return_all=True,
            age_dist=age_distribution(L_mp, ρ),
            ρ=ρ
        )[0]

    return np.sum(lamb, axis=0)[onset]


# %% [code]
N = 1024

param_values = saltelli.sample(problem, N, calc_second_order=True)
Y = np.array([model2(x, L_no_effects, onset=8) for x in param_values])
Si = sobol.analyze(problem, Y)


# %% [code]
parameter_names = [
    'Female survival',
    'Male survival',
    'Female fecundity',
    'Male fecundity',
    'Maternal mortality',
    'Female dispersal',
    'Male dispersal',
    'Inbreeding rate'
]
param_names = [
    'lx_F', 'lx_M',
    'mx_F', 'mx_M',
    'mm',
    'df', 'dm', 'm'
]

# %% [code]
save_sobol_results(
    species_name="hum",
    Si=Si,
    param_names=param_names,
    outdir="sa_results"
)


# %% [code]
```
