## Supplementary Code for "Evolution of Menopause and the Mama’s Boy Hypothesis": Johnstone_Cant.ipynb

```
# %% [code]
import numpy as np
import matplotlib.pyplot as plt
import seaborn as sns
import copy
%matplotlib inline

# %% [markdown]
## Model parameters
- $1-\mu_{f}$ = probability of survival in a group if the breeder is a female
- $1-\mu_{m}$ = probability of survival in a group if the breeder is a male
- $d_{f}$ = female offspring dispersal probability
- $d_{m}$ = male offspring dispersal probability
- $m$ = a fraction of the young produced in a group and are fathered by males within that group (chosen at random) = the fraction of matings that occur locally
- $n_{f}$ = population size of female breeders
- $n_{m}$ = population size of male breeders
- $t$ (time) = proceeds in a series of discrete steps or cycles, during each of which the following events occur one after the other:
    - Reproduction - breeders mate; females reproduce a large number of offspring (1:1 sexes assumed). fraction $m$ are fathered by males inside that group; $1-m$ are fathered outside of that group.
    - Dispersal - offspring disperese with a probability of $\frac{d_{m}}{d_{f}}$; each arriving in a random group if they do so.
    - Adult mortality - each breeder in a group survives with probability $1-\mu_{f}$ or $1-\mu_{m}$
    - Competition - offspring in a group (IMMIGRANT or NATIVE) compete on an equal basis with others of their own sex for any vacated breeding positions; those who fail to obtain a position - die! the cycle then repeats.
- $a$ = age. Mean relatedness of female breeders to males or females in her group, changes with age.

# %% [code]
def IBD_no_age(df,dm,nf,nm,muf,mum,m):
    ''' returns np.ndarray with the probabilities that two gene copies, sampled randomly with replacement (immediately prior to reproduction) 
    from the same locus in:
    - the same individual, either male or female respectively (gf,gm), or ->
    - two females in the same patch (gff), or -> 
    - a male and a female from the same patch (gfm, gmf), or ->
    - two males in the same patch (gmm),
    are identical by descent (IBD). 
    each row in the matrix will include the coefficients of gf, gff, gmf, gmm, respectively. 
    ''' 
    a1 = 0.25*(muf**2)*(1-df)**2
    b1 = muf*(1-muf)
    c1 = (1-muf)**2
    # derived from gf:
    row1 = [1,0,-0.5*m,0] # sol = -0.5
    # derived from gff:
    
    row2 = [a1*(1/nf +(m**2)/nm)+b1/nf, 
                      a1*((nf-1)/nf)+b1*((nf-1)/nf)+c1-1,
                      a1*2*m+b1*m,
                      a1*(m**2)*(nm-1)/nm] # sol = 0
    
    a2 = muf*mum*(1-df)*(1-dm)*0.25
    b2 = muf*(1-mum)*(1-df)*0.5
    c2 = (1-muf)*mum*(1-dm)*0.5
    d2 = (1-muf)*(1-mum)
    
    row3 = [a2*(1/nf + (m**2)/nm)+b2*(m/nm)+c2*(1/nf),
                      a2*((nf-1)/nf)+c2*((nf-1)/nf),
                      a2*2*m+b2+m*c2+d2-1,
                      a2*((m**2)*(nm-1)/nm)+b2*(m*(nm-1)/nm)] # sol = 0
    
    a3 = (mum**2*(1-dm)**2)*0.25
    b3 = mum*(1-mum)*(1-dm)
    c3 = (1-mum)**2
    
    row4 = [a3*((m**2)/nm + 1/nf)+b3*m/nm,
                      a3*(nf-1)/nf,
                      a3*2*m + b3,
                      a3*(m**2)*(nm-1)/nm + b3*m*(nm-1)/nm + c3-1] # sol = 0
    
    A = np.array([row1,row2,row3,row4])
    y = np.array([-0.5,0,0,0])
    x = np.linalg.solve(A,y) #x[0] = gf, x[1] = gff, x[2] = gmf, x[3] = gmm
    
    return x

# %% [code]
def no_age_rs(df,dm,nf,nm,muf,mum,m):
    gf, gff, gfm, gmm = IBD_no_age(df,dm,nf,nm,muf,mum,m)
    rff = gff/gf
    rfm = gfm/gf # = rmf
    rmm = gmm/gf
    return rff,rfm,rmm

# %% [code]
def IBD_age_0(df,dm,nf,nm,muf,mum,m):
    ''' returns tuple with the probabilities that two alleles sampled randomly from the same locus in:
    - two different females
    - a male and a female
    - a female and a male
    - a male and a male
    in the same patch, are IBD, respectively, given that one of them is of age = 0 and the other is of whatever age.
    NOTE: gf = gm'''
    
    gf, gff, gfm, gmm = IBD_no_age(df,dm,nf,nm,muf,mum,m)
    gm = gf
    
    # gff where one female is of age = 0, the other is of any age
    gff_0 = muf*((1-df)**2)*0.25*((gf/nf + (nf-1)*gff/nf) + 2*m*gfm + (m**2)*(gf/nm + gmm*(nm-1)/nm)) + \
            (1-muf)*(1-df)*0.5*((gf/nf + gff*(nf-1)/nf) + m*gfm) 
    
    # gmf for newly born female individual of age a = 0, male is of any age
    gmf_0 = mum*(1-df)*(1-dm)*0.25*((gf/nf + gff*(nf-1)/nf) + 2*m*gfm + (m**2)*(gm/nm + gmm*(nm-1)/nm)) + \
            (1-mum)*(1-df)*0.5*(m*(gm/nm + gmm*(nm-1)/nm) + gfm)
    
    # gfm for newly born male individual of age a = 0, female is of any age
    gfm_0 = muf*(1-df)*(1-dm)*0.25*((gf/nf + gff*(nf-1)/nf) + 2*m*gfm + (m**2)*(gm/nm + gmm*(nm-1)/nm)) + \
            (1-muf)*(1-dm)*0.5*((gf/nf + gff*(nf-1)/nf) + m*gfm)
    
    # gmm where one male is of age=0, the other is of any age
    gmm_0 = mum*((1-dm)**2)*0.25*((gf/nf + (nf-1)*gff/nf) + 2*m*gfm + (m**2)*(gf/nm + gmm*(nm-1)/nm)) + \
            (1-mum)*(1-dm)*0.5*(m*(gm/nm + gmm*(nm-1)/nm) + gfm)
    
    return gff_0, gmf_0, gfm_0, gmm_0

# %% [code]
def IBD_older_age(df,dm,nf,nm,muf,mum,m,a): # a is age
    ''' NOTE: If two genes are sampled randomly, one from local female of age >0, and one from another
    randomly chosen female in the same patch, then with probability muf the other female is of age 0 and 
        was born in the previous time-step, while with probability (1-muf) the other female is of age > 0 
        and was present in the patch as an adult during the previous time-step.
        Suppose the other female is newly born:
        - with probability = 1-df, she was born in the local patch (if not, p(gene copies are IBD)=0)
    '''
    gff_0, gmf_0, gfm_0, gmm_0 = IBD_age_0(df,dm,nf,nm,muf,mum,m) # initial age
    gf = gm = IBD_no_age(df,dm,nf,nm,muf,mum,m)[0]
    gff_prev, gmf_prev, gfm_prev, gmm_prev = gff_0, gmf_0, gfm_0, gmm_0
    ages = np.linspace(0,a,int(a/muf))
    gff_arr = np.zeros(ages.size)
    gmf_arr = np.zeros(ages.size)
    gfm_arr = np.zeros(ages.size)
    gmm_arr = np.zeros(ages.size)
    for t in range(ages.size):
        gff_arr[t] = gff_prev
        gmf_arr[t] = gmf_prev
        gfm_arr[t] = gfm_prev
        gmm_arr[t] = gmm_prev
        gff_prev, gmf_prev, gfm_prev, gmm_prev = 0.5*muf*(1-df)*((gf/nf + gff_arr[t]*(nf-1)/nf)+m*gmf_arr[t]) + (1-muf)*gff_arr[t], \
        0.5*mum*(1-dm)*((gf/nf + gff_arr[t]*(nf-1)/nf)+m*gmf_arr[t]) + (1-mum)*gmf_arr[t], \
        0.5*muf*(1-df)*(m*(gm/nm + gmm_arr[t]*(nm-1)/nm)+gfm_arr[t]) + (1-muf)*gfm_arr[t], \
        0.5*mum*(1-dm)*(m*(gm/nm +gmm_arr[t]*(nm-1)/nm)+gfm_arr[t]) + (1-mum)*gmm_arr[t]
   
    return gff_arr, gmf_arr, gfm_arr, gmm_arr

# %% [code]
def IBD_older_age_v2(df,dm,nf,nm,m,a,mu): # a is age; the paper states that sex differences don't affect the result so we won't consider them in our calculation
    ''' NOTE: If two genes are sampled randomly, one from local female of age >0, and one from another
        randomly chosen female in the same patch, then with probability muf the other female is of age 0 and 
        was born in the previous time-step, while with probability (1-muf) the other female is of age > 0 
        and was present in the patch as an adult during the previous time-step.
        Suppose the other female is newly born:
        - with probability = 1-df, she was born in the local patch (if not, p(gene copies are IBD)=0)'''
    
    muf, mum = mu[:,0]
    # initial age 
    gff_0, gmf_0, gfm_0, gmm_0 = IBD_age_0(df, dm, nf, nm, muf, mum, m) 
    gf = gm = IBD_no_age(df, dm, nf, nm, muf, mum, m)[0]
    gff_prev, gmf_prev, gfm_prev, gmm_prev = gff_0, gmf_0, gfm_0, gmm_0
    ages = np.linspace(0,a,int(a/muf))
    g_arr = np.zeros((4, ages.size)) # g[0,:] = gff values ; g[1,:] = gmf values ; g[2,:] = gfm values ; g[3,:] = gmm values

    i = 1
    for t in range(ages.size):
        g_arr[:, t] = gff_prev, gmf_prev, gfm_prev, gmm_prev
        if t!=0 and t%10 == 0:
            muf, mum = mu[:,i]
            i+=1
        
        gff_prev, gmf_prev, gfm_prev, gmm_prev = 0.5*muf*(1-df)*((gf/nf + g_arr[0, t]*(nf-1)/nf)+m*g_arr[1, t]) + (1-muf)*g_arr[0, t], 0.5*mum*(1-dm)*((gf/nf + g_arr[0, t]*(nf-1)/nf)+m*g_arr[1, t]) + (1-mum)*g_arr[1, t], 0.5*muf*(1-df)*(m*(gm/nm + g_arr[3, t]*(nm-1)/nm)+g_arr[2, t]) + (1-muf)*g_arr[2, t], 0.5*mum*(1-dm)*(m*(gm/nm +g_arr[3, t]*(nm-1)/nm)+g_arr[2, t]) + (1-mum)*g_arr[3, t]
   
    return g_arr

# %% [code]
def rff_age(df,dm,nf,nm,muf,mum,m,a):
    ''' returns np.ndarrays with relatedness coefficients conditional on age a, derived from the identity probabilities '''
    gf = gm = IBD_no_age(df,dm,nf,nm,muf,mum,m)[0]
    gff_a, gmf_a, gfm_a, gmm_a = IBD_older_age(df,dm,nf,nm,muf,mum,m,a)
    rff_a, rmf_a, rfm_a, rmm_a = gff_a/gf, gmf_a/gm, gfm_a/gf, gmm_a/gm
    avrg_r = (rff_a + rmf_a)/2
    return rff_a, rmf_a, rfm_a, rmm_a, avrg_r

# %% [code]
def rff_age_v2(df, dm, nf, nm, m, a, mu):
    '''returns np.ndarrays with relatedness coefficients conditional on age a, derived from the identity probabilities.'''

    muf, mum = mu[:,0]
    gf = gm = IBD_no_age(df, dm, nf, nm, muf, mum, m)[0]
    gff_a, gmf_a, gfm_a, gmm_a = IBD_older_age_v2(df, dm, nf, nm, m, a, mu)
    rff_a, rmf_a, rfm_a, rmm_a = gff_a/gf, gmf_a/gm, gfm_a/gf, gmm_a/gm
    avrg_r = (rff_a + rmf_a)/2

    return rff_a, rmf_a, rfm_a, rmm_a, avrg_r

# %% [markdown]
values for [local, non-local] reproduction. combinations of dispersal within local and non-local: [low_female, low_male], [high_female, low_male], [low_female, high male], [high_female, high_male]

# %% [code]
rep_patterns = {'local':1, 'non_local':0} # m values - the fraction of matings that occur locally
disp_patters = {'low_female':0.15, 'high_female':0.85, 'low_male':0.15, 'high_male':0.85} # low d = 0.15 in both species, high d = 0.85 in both
disp_combs = [['low_female', 'low_male'], ['high_female', 'low_male'], ['low_female', 'high_male'], ['high_female', 'high_male']]
all_combs = copy.deepcopy(disp_combs)
all_combs1 = copy.deepcopy(all_combs)
all_combs.extend(all_combs1)
reps = [rep for rep in rep_patterns.keys() for i in range(len(disp_combs))]

for idx,rep in enumerate(reps):
    comb = all_combs[idx]
    comb.append(rep)

# %% [code]
muf = 0.1
mum = 0.1
nf = nm = 3
a = 3.3
ages = np.linspace(0,a,int(a/muf))
r_vals = {}

for i,rep_comb in enumerate(all_combs):
    m = rep_patterns[rep_comb[2]]
    df, dm = disp_patters[rep_comb[0]], disp_patters[rep_comb[1]]
    rff_a, rmf_a, rfm_a, rmm_a, avg_r = rff_age(df,dm,nf,nm,muf,mum,m,a) # we care only for changing age of female, so we only take rff_a and rmf_a values and their avg
    r_vals[(rep_comb[2],rep_comb[0],rep_comb[1])] = list([rff_a,rmf_a,avg_r])
    # print([rff_a,rmf_a,avg_r])
    r_vals[i] = (rep_comb[2],rep_comb[0],rep_comb[1])

# %% [code]
def plot_r():
    rows = 4
    cols = 2
    fig, axes = plt.subplots(nrows=rows, ncols=cols, dpi=200, sharex = 'col', sharey = 'row',figsize=(5,5))
    
    idx = 0
    for row in range(rows):
        for col in range(cols):
            rprd = r_vals[idx]
            vals = r_vals[rprd]
            axes[row][col].set_title(f"{(rprd[1][:rprd[1].rfind('_')]).title()} {(rprd[1][rprd[1].rfind('_')+1:]).title()}, {(rprd[2][:rprd[2].rfind('_')]).title()} {(rprd[2][rprd[2].rfind('_')+1:]).title()}", fontsize=8)
            axes[row][col].plot(ages,vals[0],color = "#BA6E9E", label = "$r_{f}$", linewidth = 1) # rff
            axes[row][col].plot(ages,vals[1],linestyle = ':', color = '#5C888B', label = "$r_{m}$", linewidth = 1) # rmf
            axes[row][col].plot(ages,vals[2], linestyle = '--', color = '#000000', label = "$r_{avg}$", linewidth = 1) # avg
            axes[row][col].set_xlim([0,a])
            if row == len(axes)-1:
                axes[row][col].set_xlabel('Age (relative to mean\ngeneration time)', fontsize=9)
            if col == 0:
                if row<rows//2:
                    loc = "Local"
                else:
                    loc = "Non-local"
                axes[row][col].set_ylabel(f'Relatedness\n({loc})', fontsize=9)
            idx+=1
            sns.despine()
    
    axes[-1][0].set_xticks(np.arange(a), fontsize=7)
    axes[-1][1].set_xticks(np.arange(a), fontsize=7)
    
    handles,labels = axes[-1][-1].get_legend_handles_labels()
    
    fig.legend(handles,labels,fontsize=6)
    plt.tight_layout()   
    plt.show()

# %% [code]
plot_r()

# %% [markdown]
## Symbols

# %% [code]
# import sympy as sy

# %% [code]
# def IBD_no_age_sy():
#     ''' returns np.ndarray with the probabilities that two gene copies, sampled randomly with replacement (immediately prior to reproduction) 
#     from the same locus in:
#     - the same individual, either male or female respectively (gf,gm), or ->
#     - two females in the same patch (gff), or -> 
#     - a male and a female from the same patch (gfm, gmf), or ->
#     - two males in the same patch (gmm),
#     are identical by descent (IBD). 
#     each row in the matrix will include the coefficients of gf, gff, gmf, gmm, respectively. 
#     '''
#     df,dm,nf,nm,muf,mum,m = sy.symbols('d_f d_m n_f n_m μ_f μ_m m')
#     a1 = 0.25*(muf**2)*(1-df)**2
#     b1 = muf*(1-muf)
#     c1 = (1-muf)**2
#     # derived from gf:
#     row1 = [1,0,-0.5*m,0] # sol = -0.5
#     print(row1)
#     # derived from gff:
    
#     row2 = [a1*(1/nf +(m**2)/nm)+b1/nf, 
#                       a1*((nf-1)/nf)+b1*((nf-1)/nf)+c1-1,
#                       a1*2*m+b1*m,
#                       a1*(m**2)*(nm-1)/nm] # sol = 0

#     print(row2)
        
#     a2 = muf*mum*(1-df)*(1-dm)*0.25
#     b2 = muf*(1-mum)*(1-df)*0.5
#     c2 = (1-muf)*mum*(1-dm)*0.5
#     d2 = (1-muf)*(1-mum)
    
#     row3 = [a2*(1/nf + (m**2)/nm)+b2*(m/nm)+c2*(1/nf),
#                       a2*((nf-1)/nf)+c2*((nf-1)/nf),
#                       a2*2*m+b2+m*c2+d2-1,
#                       a2*((m**2)*(nm-1)/nm)+b2*(m*(nm-1)/nm)] # sol = 0

#     print(row3)
    
#     a3 = (mum**2*(1-dm)**2)*0.25
#     b3 = mum*(1-mum)*(1-dm)
#     c3 = (1-mum)**2
    
#     row4 = [a3*((m**2)/nm + 1/nf)+b3*m/nm,
#                       a3*(nf-1)/nf,
#                       a3*2*m + b3,
#                       a3*(m**2)*(nm-1)/nm + b3*m*(nm-1)/nm + c3-1] # sol = 0
#     print(row4)
    
#     A = sy.Matrix([row1,row2,row3,row4])
#     print(A)
#     y = sy.Matrix([-0.5,0,0,0])
#     print(y)
#     x = A.solve(y) #x[0] = gf, x[1] = gff, x[2] = gmf, x[3] = gmm
    
#     return x

# %% [code]
# IBD_no_age_sy()

# %% [code]
```
