## Supplementary Code for "Evolution of Menopause and the Mama’s Boy Hypothesis": KW_live_long.ipynb

```
# %% [code]
import numpy as np
from pandas import DataFrame, read_csv
import matplotlib.pyplot as plt
import seaborn as sns
from copy import deepcopy
from scipy.linalg import leslie
%matplotlib inline
import matplotlib.image as mpimg

# %% [code]
import io, os, sys, types

# %% [code]
from IPython import get_ipython
from nbformat import read
from IPython.core.interactiveshell import InteractiveShell

# %% [code]
def find_notebook(fullname, path=None):
    """find a notebook, given its fully qualified name and an optional path

    This turns "foo.bar" into "foo/bar.ipynb"
    and tries turning "Foo_Bar" into "Foo Bar" if Foo_Bar
    does not exist.
    """
    name = fullname.rsplit('.', 1)[-1]
    if not path:
        path = ['']
    for d in path:
        nb_path = os.path.join(d, name + ".ipynb")
        if os.path.isfile(nb_path):
            return nb_path
        # let import Notebook_Name find "Notebook Name.ipynb"
        nb_path = nb_path.replace("_", " ")
        if os.path.isfile(nb_path):
            return nb_path

# %% [code]
class NotebookLoader(object):
    """Module Loader for Jupyter Notebooks"""
    def __init__(self, path=None):
        self.shell = InteractiveShell.instance()
        self.path = path

    def load_module(self, fullname):
        """import a notebook as a module"""
        path = find_notebook(fullname, self.path)

        print ("importing Jupyter notebook from %s" % path)

        # load the notebook object
        with io.open(path, 'r', encoding='utf-8') as f:
            nb = read(f, 4)


        # create the module and add it to sys.modules
        # if name in sys.modules:
        #    return sys.modules[name]
        mod = types.ModuleType(fullname)
        mod.__file__ = path
        mod.__loader__ = self
        mod.__dict__['get_ipython'] = get_ipython
        sys.modules[fullname] = mod

        # extra work to ensure that magics that would affect the user_ns
        # actually affect the notebook module's ns
        save_user_ns = self.shell.user_ns
        self.shell.user_ns = mod.__dict__

        try:
          for cell in nb.cells:
            if cell.cell_type == 'code':
                # transform the input to executable Python
                code = self.shell.input_transformer_manager.transform_cell(cell.source)
                # run the code in themodule
                exec(code, mod.__dict__)
        finally:
            self.shell.user_ns = save_user_ns
        return mod

# %% [code]
class NotebookFinder(object):
    """Module finder that locates Jupyter Notebooks"""
    def __init__(self):
        self.loaders = {}

    def find_module(self, fullname, path=None):
        nb_path = find_notebook(fullname, path)
        if not nb_path:
            return

        key = path
        if path:
            # lists aren't hashable
            key = os.path.sep.join(path)

        if key not in self.loaders:
            self.loaders[key] = NotebookLoader(path)
        return self.loaders[key]

# %% [code]
sys.meta_path.append(NotebookFinder())

# %% [code]
from Johnstone_Cant import IBD_age_0, no_age_rs, IBD_no_age, IBD_older_age_v2, rff_age_v2

# %% [code]
from project_functions import sex_weight, survival2x, leslie_, Px, mx, age_distribution, tot_rBC_all_donor_life, tot_rBC, survival_effect, fecundity_effect, reproductive_value, different_longevities, latex_array, add_image, evaluate_survival_fecundity_tradeoffs, plot_lambda_zero_contours, PrR_from_model


# %% [code]
d = 10 # looking at 10 age classes: 0-10,10-20,20-30,30-40,40-50,50-60,60-70,70-80,80-90,90-100
L_mp = np.zeros((2*d, 2*d))
fecundity = np.zeros(d)
fecundity[1:4] = [1, 2, 2]
fecundity_both_sex = np.array([fecundity, fecundity])
survival_f = np.array([3903/5000, 389/400, 2491/2500, 1971/2000, 9641/10000, 4711/5000, 4491/5000, 431/500, 431/500])
survival_m = np.array([3903/5000, 0.9315, 0.9149, 0.8588, 0.8588, 0.8588, 0, 0, 0])
survival_both_sex = np.array([survival_f, survival_m])
L_mp = leslie_(fecundity_both_sex, survival_both_sex)

# %% [code]
mx_contribution=[0.95, 0.75]
lx_contribution=[0.8, 0.5]
female_fec_tradeoff_arr = np.linspace(1.5,1,6)
female_surv_tradeoff_arr = np.linspace(1.15,1,6)
additional_years_after_rep = 6
# L_nomps = create_longer_life_nomp_LMs(L_mp, lx_contribution, mx_contribution, female_fec_tradeoff_arr, female_surv_tradeoff_arr, additional_years_after_rep, Δmaternal_mortality=0.97)

# %% [code]
male_symbol = mpimg.imread('/Users/Omer/Downloads/pTodqKkpc-3982000212.png')
female_symbol = mpimg.imread('/Users/Omer/Downloads/8czr7L9Ki-3456542989.png')
RKW_phylopic = mpimg.imread('/Users/Omer/Downloads/1024x503.png')

# %% [markdown]
# Resident

# %% [code]
df = dm = 0.1 # male, female dispersal occurs at a probability 0.1
m = 0 # indicative for non-local reproduction as in orcas
nf = nm = 3
a = 10 # number of age classes to view
mu = np.zeros((2,10)) # array for survival probabilities per age-group, where each age group is in 10-year intervals
for i in range(9):
    mu[:,i] = L_mp[i+1,i], L_mp[10+i+1,10+i]
rf_to_f, rm_to_f, rf_to_m, rm_to_m, avg_rf = rff_age_v2(df, dm, nf, nm, m, a, 1-mu)
r_to_pod = np.array([rf_to_f, rm_to_f])

r_to_pod_10yr = np.concatenate([r_to_pod[:,::5], (r_to_pod[:, -1]).reshape(2,1)], axis=1)
age_classes = np.arange(0,100,10)

lw, fs = 2.5, 17

fig, ax = plt.subplots(figsize=(8,5))
ax.plot(age_classes, r_to_pod_10yr[0,:], label='$r_f$', c='salmon', lw=lw, marker='o')
ax.plot(age_classes, r_to_pod_10yr[1,:], label='$r_m$', c='cornflowerblue', lw=lw, marker='o')
# ax.axvline(40,c='gray', ls=':', label='$r$ at ageD=40')
ax.plot(age_classes, 0.6*r_to_pod_10yr[0,:]+0.4*r_to_pod_10yr[1,:], c='#FFC98B', label='$r_{avg}$', lw=lw, ls='--')# marker='o')
ax.set_xlabel('Female age', fontsize=fs+2)
ax.set_ylabel('Relatedness coefficient', fontsize=fs+2)
ax.tick_params(axis='both', which='major', labelsize=fs)
ax.set_xticks(np.arange(0,100,10))
add_image(ax, male_symbol, xy=(6, 0.075), zoom=0.014)
add_image(ax, female_symbol, xy=(0.5, 0.205), zoom=0.011)
# ax.set_title('Relatedness Coefficient of Female to Pod Throughout Life (10-year intervals)', fontsize=fs)
# ax.legend(fontsize=fs-3)
add_image(ax, RKW_phylopic, xy=(82, 0.08), zoom=0.1)
plt.savefig('RKW_r.png')

plt.show()

# %% [markdown]
# Transient

# %% [code]
dfT = dmT = 0.35 # male, female dispersal occurs at a probability 0.1
mT = 0.2 # indicative for non-local reproduction as in orcas
nfT = nmT = 3
aT = 10 # number of age classes to view
muT = np.zeros((2,10)) # array for survival probabilities per age-group, where each age group is in 10-year intervals
for i in range(9):
    muT[:,i] = L_mp[i+1,i], L_mp[10+i+1,10+i]
rf_to_fT, rm_to_fT, rf_to_mT, rm_to_mT, avg_rfT = rff_age_v2(dfT, dmT, nfT, nmT, mT, aT, 1-muT)
r_to_podT = np.array([rf_to_fT, rm_to_fT])

r_to_pod_10yr_T = np.concatenate([r_to_podT[:,::5], (r_to_podT[:, -1]).reshape(2,1)], axis=1)
age_classes = np.arange(0,100,10)

lw, fs = 2.5, 17

fig, ax = plt.subplots(figsize=(8,5))
ax.plot(age_classes[:], r_to_pod_10yr_T[0,:], label='$r_f$', c='salmon', lw=lw, marker='o')
ax.plot(age_classes[:], r_to_pod_10yr_T[1,:], label='$r_m$', c='cornflowerblue', lw=lw, marker='o')
# ax.axvline(40,c='gray', ls=':', label='$r$ at ageD=40')
ax.plot(age_classes[:], 0.6*r_to_pod_10yr_T[0,:]+0.4*r_to_pod_10yr_T[1,:], c='#FFC98B', label='$r_{avg}$', lw=lw, ls='--')# marker='o')
ax.set_xlabel('Female age', fontsize=fs+2)
ax.set_ylabel('Relatedness coefficient', fontsize=fs+2)
ax.set_xticks(np.arange(0,100,10))
ax.tick_params(axis='both', which='major', labelsize=fs)
add_image(ax, male_symbol, xy=(5, 0.065), zoom=0.014)
add_image(ax, female_symbol, xy=(0.5, 0.1325), zoom=0.011)
# ax.set_title('Relatedness Coefficient of Female to Pod Throughout Life (10-year intervals)', fontsize=fs)
# ax.legend(fontsize=fs-3)
add_image(ax, RKW_phylopic, xy=(82, 0.0655), zoom=0.1)
plt.savefig('TKW_r.png')

plt.show()

# %% [code]
# Define parameter ranges to test
additional_years_after_rep = 6
surv_range = np.linspace(1.1, 1, 50)  # Survival trade-off values
fec_range = np.linspace(1.1, 1, 50)   # Fecundity trade-off values

# %% [markdown]
## Resident results

# %% [code]
res = evaluate_survival_fecundity_tradeoffs(
    surv_range,
    fec_range,
    additional_years_after_rep,
    L_mp, 
    r_to_pod_10yr,
    female_fec_tradeoff_arr,
    female_surv_tradeoff_arr,
    Δmaternal_mortality=1,
    mx_contribution=np.array([0.95, 0.75]),
    lx_contribution=np.array([0.8, 0.5]),
    sex_ratio=[0.6, 0.4],
    return_all=True)

# %% [code]


# %% [code]
fig, ax = plot_lambda_zero_contours(
    res, surv_range, fec_range, additional_years_after_rep,
    years_per_step=10,
    fallback_when_no_zero='max',
    highlight_year=60,                    # <— bold + shade below the 30y line
    shade_below_highlight=True, figsize=(12, 10), non_meno_col='salmon', meno_col='#99B49B', fs=30, alpha=1)
add_image(ax, RKW_phylopic, xy=(1.047, 1.035), zoom=0.29)

ax.set_xlim(1.0, 1.06)
ax.set_ylim(1.0, 1.04)
ax.set_xlabel('Fecundity Trade-off', fontsize=0)
ax.set_ylabel('Survival Trade-off', fontsize=34)
ax.tick_params(axis='x', which='major', labelsize=0)
ax.tick_params(axis='y', which='major', labelsize=32)
ax.text(
    0.0, -0.04,                 # x=0 (left edge), slightly below axis
    'No cost',
    transform=ax.transAxes,
    ha='center',
    va='top',
    fontsize=28,
    color='black'
)
plt.savefig('LL_RKW.png', bbox_inches="tight")

plt.show()


# %% [code]
results = res

# %% [code]
fec_range[37], surv_range[41]

# %% [code]
additional_years_after_rep_try = 6
female_surv_tradeoff_arr_try = [1 + (1.01-1) / (i + 1) for i in range(additional_years_after_rep_try)]
female_fec_tradeoff_arr_try = [1 + (1.025-1) / (i + 1) for i in range(additional_years_after_rep_try)]

female_surv_tradeoff_arr_try, female_fec_tradeoff_arr_try

# %% [code]
rBC, C, B, rxB = different_longevities(L_mp, r_to_pod_10yr, female_fec_tradeoff_arr_try, female_surv_tradeoff_arr_try, additional_years_after_rep_try, Δmaternal_mortality=1, mx_contribution=np.array([0.95, 0.75]), lx_contribution=np.array([0.8, 0.5]), sex_ratio=[0.6,0.4], return_all=True)
np.sum(rBC, axis=0)

# %% [code]
RKW_phylopic = mpimg.imread('/Users/Omer/Downloads/1024x503.png')

# %% [code]
ε = 10**(-60) 
age_classes = np.arange(10, 61, 10)
rBC, C, B, rxB = different_longevities(L_mp, r_to_pod_10yr, female_fec_tradeoff_arr_try, female_surv_tradeoff_arr_try, additional_years_after_rep_try, Δmaternal_mortality=1, mx_contribution=np.array([0.95, 0.75]), lx_contribution=np.array([0.8, 0.5]), sex_ratio=[0.6,0.4], return_all=True)

fig, ax = plt.subplots(figsize=(10,7))
ax.axhline(0,c='k')
ax.plot(age_classes, np.sum(rBC, axis=0), label='λ', marker='o', c='#E9765B', lw=lw, markersize=7)
ax.plot(age_classes, np.sum(C, axis=0), label='C', ls='-', c='#FFC98B', lw=lw)
ax.plot(age_classes, np.sum(B, axis=0), label='B', ls='-', c='#A5CAD2', lw=lw)
ax.plot(age_classes, np.sum(rxB, axis=0), label='r*B', ls='-', c='#478BA2', lw=lw)
add_image(ax, RKW_phylopic, xy=(14, 0.05), zoom=0.1)
ax.axvline(10*(np.where(np.sum(rBC, axis=0)>0)[0][0] +1), c='k', ls=':', label='1st positive')
ax.set_xlabel('Added post-reproductive years', fontsize=fs+2)
ax.set_ylabel('Benefit and cost over lifespan', fontsize=fs+2)
# ax.set_ylabel('λ', fontsize=fs+2)
ax.tick_params(axis='both', which='major', labelsize=fs)
# ax.set_xlim(0,91)
ax.set_xticks(np.arange(10,70,10))
# ax.set_yscale('log')
# plt.savefig('lambda_rkw.png')
# ax.set_title('\n$d_f = d_m = 0$;  $m=1$;  $n_f = n_m = 30$', fontsize=fs+4)
# ax.set_title('\n$d_f = d_m = 0.1$;  $m=0$;  $n_f = n_m = 3$', fontsize=fs+4)
# sns.despine()
# ax.legend(loc='upper right', fontsize=fs - 3)
# ax.set_title('\n$λ$, C, B and r*B values as a function of different menopause\nonset ages for female donor', fontsize=fs)
ax.set_ylim(-0.025, 0.06)
plt.savefig('RKW_live_longer.png', bbox_inches='tight')
plt.show()


# %% [markdown]
## Transient results

# %% [code]
res_T = evaluate_survival_fecundity_tradeoffs(
    surv_range,
    fec_range,
    additional_years_after_rep,
    L_mp, 
    r_to_pod_10yr_T,
    female_fec_tradeoff_arr,
    female_surv_tradeoff_arr,
    Δmaternal_mortality=1,
    mx_contribution=np.array([0.95, 0.75]),
    lx_contribution=np.array([0.8, 0.5]),
    sex_ratio=[0.6, 0.4],
    return_all=True)

# %% [code]


# %% [code]
fig, ax = plot_lambda_zero_contours(
    res_T, surv_range, fec_range, additional_years_after_rep,
    years_per_step=10,
    fallback_when_no_zero='max',
    highlight_year=60,                    # <— bold + shade below the 30y line
    shade_below_highlight=True, figsize=(12, 10), non_meno_col='salmon', meno_col='#99B49B', fs=30,alpha=1)
add_image(ax, RKW_phylopic, xy=(1.047, 1.035), zoom=0.29)
ax.set_xlim(1.0, 1.06)
ax.set_ylim(1.0, 1.04)
ax.set_xlabel('Fecundity Trade-off', fontsize=0)
ax.set_ylabel('Survival Trade-off', fontsize=0)
ax.tick_params(axis='both', which='major', labelsize=0)
plt.savefig('LL_tkw.png', bbox_inches="tight")

plt.show()


# %% [code]
results = res_T

# %% [code]
additional_years_after_rep_try = 6
female_surv_tradeoff_arr_try = [1 + (1.01-1) / (i + 1) for i in range(additional_years_after_rep_try)]
female_fec_tradeoff_arr_try = [1 + (1.015-1) / (i + 1) for i in range(additional_years_after_rep_try)]

female_surv_tradeoff_arr_try, female_fec_tradeoff_arr_try

# %% [code]
rBC, C, B, rxB = different_longevities(L_mp, r_to_pod_10yr_T, female_fec_tradeoff_arr_try, female_surv_tradeoff_arr_try, additional_years_after_rep_try, Δmaternal_mortality=1, mx_contribution=np.array([0.95, 0.75]), lx_contribution=np.array([0.8, 0.5]), sex_ratio=[0.6,0.4], return_all=True)
np.sum(rBC, axis=0)

# %% [code]
TKW_phylopic = mpimg.imread('/Users/Omer/Downloads/1024x503.png')

# %% [code]
ε = 10**(-60) 
age_classes = np.arange(10, 61, 10)
rBC, C, B, rxB = different_longevities(L_mp, r_to_pod_10yr_T, female_fec_tradeoff_arr_try, female_surv_tradeoff_arr_try, additional_years_after_rep_try, Δmaternal_mortality=1, mx_contribution=np.array([0.95, 0.75]), lx_contribution=np.array([0.8, 0.5]), sex_ratio=[0.6,0.4], return_all=True)

fig, ax = plt.subplots(figsize=(10,7))
ax.axhline(0,c='k')
ax.plot(age_classes, np.sum(rBC, axis=0), label='λ', marker='o', c='#E9765B', lw=lw, markersize=7)
ax.plot(age_classes, np.sum(C, axis=0), label='C', ls='-', c='#FFC98B', lw=lw)
ax.plot(age_classes, np.sum(B, axis=0), label='B', ls='-', c='#A5CAD2', lw=lw)
ax.plot(age_classes, np.sum(rxB, axis=0), label='r*B', ls='-', c='#478BA2', lw=lw)
add_image(ax, RKW_phylopic, xy=(14, 0.05), zoom=0.1)
ax.axvline(10*(np.where(np.sum(rBC, axis=0)>0)[0][0] +1), c='k', ls=':', label='1st positive')
ax.set_xlabel('Added post-reproductive years', fontsize=fs+2)
ax.set_ylabel('Benefit and cost over lifespan', fontsize=fs+2)
# ax.set_ylabel('λ', fontsize=fs+2)
ax.tick_params(axis='both', which='major', labelsize=fs)
# ax.set_xlim(0,91)
ax.set_xticks(np.arange(10,70,10))
# ax.set_yscale('log')
# plt.savefig('lambda_rkw.png')
# ax.set_title('\n$d_f = d_m = 0$;  $m=1$;  $n_f = n_m = 30$', fontsize=fs+4)
# ax.set_title('\n$d_f = d_m = 0.1$;  $m=0$;  $n_f = n_m = 3$', fontsize=fs+4)
# sns.despine()
# ax.legend(loc='upper right', fontsize=fs - 3)
# ax.set_title('\n$λ$, C, B and r*B values as a function of different menopause\nonset ages for female donor', fontsize=fs)
ax.set_ylim(-0.025, 0.06)
plt.savefig('TKW_live_longer.png', bbox_inches='tight')
plt.show()
```
