## Supplementary Code for "Evolution of Menopause and the Mama’s Boy Hypothesis": project_funcs.ipynb

```
# %% [markdown]
#  Kin selection in age-structured populations - FUNCTIONS
This model shows the spread, if occurs, of a rare mutant allele $A_{2}$ ("menopause allele") that appears in a $A_1$ ("no menopause allele") locus, in a homogenous $A_{1}A_{1}$ ("no-menopause genotype") population. The allele makes the carrier altruistic. 

This model is based on the framework in:
> Charlesworth, B. & Charnov, E.L. 1981. **Kin selection in age-structured populations**, Journal of Theoretical Biology. 88, 103-119. https://doi.org/10.1016/0022-5193(81)90330-1.

# %% [code]
import numpy as np
from pandas import DataFrame, read_csv
import matplotlib.pyplot as plt
import seaborn as sns
from copy import deepcopy
from scipy.linalg import leslie
%matplotlib inline
import matplotlib.image as mpimg
from matplotlib.offsetbox import OffsetImage, AnnotationBbox
from SALib.sample import saltelli
from SALib.analyze import sobol
import traceback
import matplotlib.patheffects as pe
from matplotlib.path import Path
from matplotlib.patches import PathPatch

# %% [markdown]
## Model assumptions
- Infintiely large, random mating popualtion
- Exponential population growth
- Initially allele $A_{1}$ (no menopause) is fixed
- Primary sex ratio is $c$, and it's independent of the parents' age
> According to Ellis et al. (2021), the common sex-ratio within orca pods is f:m = ~0.6:0.4
- Female demographic dominance - the fecundity of a mating depends only on maternal age.
- $A_{2}A_{2}$ genotype is in very low frequency for a long period of time _(negligible)_
- The result assumes the effect of $A_{2}$ on the frequency of behavioral interactions is so low that we can neglect the possibility of an individual's experiencing _both_ the effect of $A_{2}$ its own behaviour _and_ the effect of $A_{2}$ on the behaviour of its relatives ; or that the same individual cannot act as both donor and recipient simultaneously
- The effect of one interaction between donor and recipient is assumed to not endure over more than one time interval. Therefore, `I sum interactions over different ages`
- The number of donors and recipients engaged in interactions at a given time are the same

# %% [markdown]
## Model parameters
- $\rho\ :$ intrinsic rate of increase of the population _(leading eigenvalue, λ, of Leslie matrix)_
- $\alpha\ :$ $\text{Age}_{recipient} - \text{Age}_{donor}$ = $x_j - x_i$
- $d\ :$ maximal number of age classes.
    - male orcas live ~40-50 years (Olesiuk et al., 2005)
    - females orcas live ~90-100 years, so $d\ =\ 10$ (age classes) if looked at with 10 year intervals (Olesiuk et al., 2005)     
- $a_{g,j}(x_i, r_g)\ :$ expected number of interactions of an $A_{1}A_{2}$ female aged $x_i$ as donor towards $A_{1}A_{1}$ recipient of sex g ($g \in \set{f\ (\text{female}), m\ (\text{male})}$), age $x_j$ and relatedness $r_g$. This is crucial as the calculated $r_g$ is the average relatedness between the donor and each sex but the age distribution isn't uniform so weights should be given for each age+sex-specific interactions. So the age distribution is used for that.
    for example, a cost expression for donor's fecundity will look like: 
    $$
    COST = e^{-\rho x_i}\sum_{g}\sum_{j}a_{g,j}(x_i, r_g)l(x_i)\delta_i m(x_i)
    $$
    
> **I calculate the r values from the relatedenss model in the paper**: Johnstone R.A. & Cant M.A. 2010. The evolution of menopause in cetaceans and humans: the role of demography. Proc. R. Soc. B.2773765–3771
http://doi.org/10.1098/rspb.2010.0988

# %% [markdown]
### Relatedness coefficients
The functions are in a different notebook (Johnstone_Cant.ipynb) so I import them using this code $\rightarrow$ https://jupyter-notebook.readthedocs.io/en/stable/examples/Notebook/Importing%20Notebooks.html 

# %% [code]
import io, os, sys, types

# %% [code]
from IPython import get_ipython
from nbformat import read
from IPython.core.interactiveshell import InteractiveShell

# %% [code]
def find_notebook(fullname, path=None):
    """find a notebook, given its fully qualified name and an optional path

    This turns "foo.bar" into "foo/bar.ipynb"
    and tries turning "Foo_Bar" into "Foo Bar" if Foo_Bar
    does not exist.
    """
    name = fullname.rsplit('.', 1)[-1]
    if not path:
        path = ['']
    for d in path:
        nb_path = os.path.join(d, name + ".ipynb")
        if os.path.isfile(nb_path):
            return nb_path
        # let import Notebook_Name find "Notebook Name.ipynb"
        nb_path = nb_path.replace("_", " ")
        if os.path.isfile(nb_path):
            return nb_path

# %% [code]
class NotebookLoader(object):
    """Module Loader for Jupyter Notebooks"""
    def __init__(self, path=None):
        self.shell = InteractiveShell.instance()
        self.path = path

    def load_module(self, fullname):
        """import a notebook as a module"""
        path = find_notebook(fullname, self.path)

        print ("importing Jupyter notebook from %s" % path)

        # load the notebook object
        with io.open(path, 'r', encoding='utf-8') as f:
            nb = read(f, 4)


        # create the module and add it to sys.modules
        # if name in sys.modules:
        #    return sys.modules[name]
        mod = types.ModuleType(fullname)
        mod.__file__ = path
        mod.__loader__ = self
        mod.__dict__['get_ipython'] = get_ipython
        sys.modules[fullname] = mod

        # extra work to ensure that magics that would affect the user_ns
        # actually affect the notebook module's ns
        save_user_ns = self.shell.user_ns
        self.shell.user_ns = mod.__dict__

        try:
          for cell in nb.cells:
            if cell.cell_type == 'code':
                # transform the input to executable Python
                code = self.shell.input_transformer_manager.transform_cell(cell.source)
                # run the code in themodule
                exec(code, mod.__dict__)
        finally:
            self.shell.user_ns = save_user_ns
        return mod

# %% [code]
class NotebookFinder(object):
    """Module finder that locates Jupyter Notebooks"""
    def __init__(self):
        self.loaders = {}

    def find_module(self, fullname, path=None):
        nb_path = find_notebook(fullname, path)
        if not nb_path:
            return

        key = path
        if path:
            # lists aren't hashable
            key = os.path.sep.join(path)

        if key not in self.loaders:
            self.loaders[key] = NotebookLoader(path)
        return self.loaders[key]

# %% [code]
sys.meta_path.append(NotebookFinder())

# %% [code]
from Johnstone_Cant import IBD_age_0, no_age_rs, IBD_no_age, IBD_older_age_v2, rff_age_v2

# %% [markdown]
## Model variables and functions

# %% [markdown]
#### Survival probabilities
- $P_g(0)$ where $g \in {f, m}$ is the probability of survival of sex g from conception to age class 1
- $P_g(x)\ =$ probabilities of survival from age $x$ to $x+1$ $(x\geq1)$
- $l_g(x)=P_g(0)P_g(1)...P_g(x-1)$ , $(x\geq1)$ $=$ probabilities of survival from conception to age x:
$$
(1)\ \ \ l_{g}(x) = \Pi_{u=0}^{x-1}P_{g}(u)
$$

# %% [code]
def Px(L):
    '''    
    Parameters
    -----------
    L : np.ndarray 
            sex-specific Leslie matrix

    Description
    -----------
    p(x) in the model = probability to survive to age x+1.
    This function returns a (2, d) shaped numpy array where d is the number of age classes, containing the survival probabilities from the leslie matrix, both for female (f; row 0 in the returned array) and male (m; row 1). 
    The first element in each row is 1 because it is the probability to survive to age 0 (if an individual is born, it stays alive in the first age class). 
    This value does not appear in the matrix because the matrix only shows the probabilities to survive to the *next* age class and since the first class has no previous class, it is not shown. 
    I will note that this function is specified for a leslie matrix of shape (2d, 2d), because it is like two (d,d) matrices concatenated where each matrix is sex-specific.
    '''
    
    # because the matrix is of shape (2d, 2d) where d is the number of age classes, I divide by 2 to get d. The name "d" for this variable comes from the presented model, where d is the maximal age (class) of the population.
    d = L.shape[0]//2
    # diagonal of the upper left matrix - females. The leslie matrix shows the probability to survive to the next age class, so the diagonal in this matrix has 9 age classes even though there are actually 10. That's because it's a given that the last age class will not survive to the next one. I take the diagonal of L[1:d, :d-1] and not L[:d, :d] because the first row in the leslie matrix is the fecundity values, not the survival values (hence rows 1:d), and the last column does not actually hold any survival values because it is the last age class (so p(next age)=0).
    Pxf = np.diagonal(L[1:d, :d-1]) 
    # diagonal of the lower right matrix - males
    Pxm = np.diagonal(L[d+1:, d:-1]) 
    # Since the "survival2x" function (=l(x)) looks at the probability to survive to a certain age (multiplication of all probabilities before that age), I shift everything by 1 so instead of getting [p(10-20), p(20-30),...,p(100-110)] I get [p(0-10), p(10-20),...,p(90-100)] where p(0-10) is 1 because if you are born you survive your birth.
    return np.concatenate([np.array([[1],[1]]), np.array([Pxf, Pxm])], axis=1) 


# %% [code]
def survival2x(L): 
    ''' 
    Parameters
    ----------
    L : np.ndarray 
            sex-specific Leslie matrix
    
    Description
    -----------
    This function is l(x) in the model equation.
    It returns a numpy array with shape (2, d), where 2 is for sex-specificy (row 0=f, row 1=m) and d is the number of age classes. 
    Each element in the returned array represents the probability of that sex and age to survive *to that age* (cumulative product of the probabilities before). Notice this is different from the probabilities in the leslie matrix because there each survival probability is about
    surviving to *the next age*. So, in each row (=sex), index 0 is the probability to survive to age 0-10 (if you are born you survive your birth) which is p=1, index 1 is the prob. to survive to age 10-20, which is p(age 0-10) * p(age 10-20) and so on (eq. 1).
    ''' 
    
    # For l(x), we need p(x) of females and p(x) of males
    Pxf, Pxm = Px(L) 
    # Cumulative product - each element is the product of the elements up to their point
    lx = np.array([np.cumprod(Pxf), np.cumprod(Pxm)])
    
    return lx

# %% [markdown]
#### Age distribution
I calculate the age distribution using equation 10.4.4 in Otto and Day's "A Biologist's Guide to Mathematical Modeling in Ecology and Evolution" book (p. 408):
$$(2)\ \ \ a_{g,j}(x_i, r_g) = \frac{1}{\sum_{k=1}^{d}l_g(x_k)\rho^{-(x_k-1)}}\ l_g(x_j)\rho^{-(x_j-1)}
$$
where $a_{g,j}(x_i, r_g)$ is the fraction of age class x and sex g in the population.

# %% [code]
def age_distribution(L, ρ, sex_ratio=np.array([[0.6],[0.4]])): 
    '''
    Parameters
    ----------
    ρ : float 
            population growth rate 
    L : np.ndarray 
            sex-specific Leslie matrix
    sex_ratio : np.ndarray
            sex_ratio[0] is for F and sex_ratio[1] is for M. 
    
    Description
    -----------
    sex_ratio[1] = c in the model equation (therefore sex_ratio[0]=1-c). The literature indicates ~60% of the group are females and the rest are males, so I use this as a default value.
    This function returns the sex-specific age distribution in the Orca population, according to equation 10.19 in the book "A Biologist's Guide to Mathematical Modeling in Ecology and Evolution" by Sarah P. Otto and Troy Day, page 407.  
    '''
    
    # d is the number of age classes. Also make sure growth rate is a float.
    ρ, d = float(ρ), L.shape[0]//2
    # Cumulative survival probabilities
    lx = survival2x(L)
    assert sex_ratio.shape[0] == lx.shape[0] 
    # normalization by sex ratio
    sum_over = np.arange(0,d,1) # for normalization
    normalize = (np.sum(sex_ratio*lx*np.power(ρ, -sum_over))) 
    # Calculation of the age distribution
    age_dist = sex_ratio*lx*np.power(ρ, -sum_over)/normalize

    return age_dist

# %% [markdown]
#### Fecundity
- $m_f(x)\ =$ _#female_ offspring a female of age x produces (counted here at the zygote stage) $\rightarrow$ the total number of offspring of a female aged x is thus $\frac{m_f(x)}{1-c}$.
- $m_m(x)\ =$ _#male_ offspring a male of age x produces $\rightarrow$ # total # offspring of a male aged x is $\frac{m_{m}(x)}{c}$.

# %% [code]
def mx(L): 
    '''
    Parameters
    ----------
    L : np.ndarray 
            sex-specific Leslie matrix

    Description
    -----------
    mx = m(x) in the model = fecundity at age x. 
    This function returns a (2, d) numpy array of the sex-speific fecundity values at each age class.
    '''
    
    # d is the number of age classes
    d = L.shape[0]//2
    # fecundity of females in the first row, first d columns; fecundity of males in the d-th row, last d columns
    mx_F_and_M = np.array([L[0,:d], L[d,d:]]) 

    return mx_F_and_M

# %% [markdown]
#### $\Delta$ survival probabilities
- $δ_DP(x_{i})$ = change in survival probability from age class $x_{i}$ to $x_{i}+1$ for a female donor as a result of a single altruistic interaction event.
- $δ_RP(x_{j})$ = change in survival probability of recipient. 
#### $\Delta$ fecundity
- $δ_Dm(x_{i})$ = change in donor fecundity
- $δ_Rm(x_{j})$ = change in recipient fecundity.

> NOTE: because the actual Leslie Matrix for Orca populations describes a menopausal one, the changes we consider to build the non-menopausal Leslie matrix represent the transition from a menopausal to a non-menopausal pop. (e.g, positive no. of offspring for females at ages > 40, decrease in young-survival, etc.).

# %% [code]
def leslie_(m, p):
    '''
    Parameters
    ----------
    m : np.ndarray 
            sex specific fecundity values in shape (2, d) where row 0 is females' fecundity values and row 1 is males', and d is the number of age classes.
    p : np.ndarray
            sex specific survival probabilities in shape (2,d-1) because there are d-1 survival probability values in the Leslie matrix.

    Description
    -----------
    This function is based on scipy's function constructing a leslie matrix ("leslie" function), but only of shape (d,d). 
    Since I use a specific format of a leslie matrix looking at males and females separately and not the population as a whole, I construct a leslie matrix of shape (2d,2d) by using the scipy function twice, the values of which I insert into an empty (2d, 2d) array. 
    That is what this function then returns and it will have the format of the leslie matrix I use in my model. This function will be necessary to construct a new leslie matrix for every possible onset age of menopause because different ages will cause fecundity=0 start appearing in different places in the matrix for females. 
    As will be seen, menopause will also have an effect on survival of the recipients, so I also construct a new leslie matrix for every possible onset age of menopause for the recipients. More on that in the "different_onset_ages" function.
    '''
    
    # d is the number of age classes
    d = m.shape[-1]
    # Setting an empty array of shape (2d, 2d) where the leslie matrices of both sexes will be inserted
    Ltot = np.zeros((2*d,2*d)) 
    # Leslie matrix construction for females and males separately
    p[p>1] = 1
    p[p<0] = 0
    m[m<0] = 0
    Lf, Lm = leslie(m[0,:], p[0,:]), leslie(m[1,:], p[1,:]) 
    # Inserting the female leslie matrix into the upper left corner of the empty array
    Ltot[:d, :d] = Lf 
    # Inserting the male leslie matrix into the lower right corner of the empty array
    Ltot[d:, d:] = Lm 

    return Ltot

# %% [markdown]
## Modeling altruism's survival effect
This expression includes the terms that represent the effect the altruistic behavior has on the survival of the donor (`cost`) and on the recipient (`benefit`).
**If the sum of terms is positive, $A_{2}$ $-$ the menopause allele $-$ will spread**.
- To include fecundity, add the relevant fecundity effect (next section).
- What I refer here as cost has to be negative by altruism definition, but we can certainly check other cases. 

The gene $A_2$ increases if the sum of terms in the simplified expression is positive (see Methods).

# %% [code]
def sex_weight(m_or_f, sex_ratio=[0.6,0.4]):
    '''
    Parameters
    ----------
    m_or_f : int (0 | 1)
            m_or_f = 0 indicates the sex is female so h(female) = 1; 
            m_or_f = 1 indicates the sex is male so h(male) = (1-c)/c = sex_ratio[0]/sex_ratio[1]
    sex_ratio : list
            sex_ratio[0] is for F and sex_ratio[1] is for M. sum(sex_ratio) should be 1.
        
    Description
    -----------
    This function is h(g) as shown in the equations above: in the model, represented by: (1-c) = sex_ratio[0] = female fraction, c = sex_ratio[1] = male fraction.
    It gives the relevant weight to each sex.
    '''
    
    weight = 1 if m_or_f == 0 else sex_ratio[0]/sex_ratio[1] 
    return weight

# %% [code]
def δPx(L_nomp, L_mp): 
    '''
    Parameters
    ----------
    L_nomp : np.ndarray
            Leslie matrix - without menopause
    L_mp : np.ndarray
            Leslie matrix - with menopause

    Description
    -----------
    This is δp(x) in the model equation, where x is the age class; δp(x) = p(x)_mp - p(x)_nomp where mp=menopause, nomp=no menopause and p(x) = probability to survive to age x+1. 
    This function calculates the change (δ) in survival probabilities between two leslie matrices - without menopause (L_nomp), the theoretical scenario, and with menopause (L_mp), the actual scenario.
    It returns an array of (2, d-1) values indicating the differences in probabilities to survive to the *next* age class in each index.
    '''

    # δp(x) = p(x)_mp - p(x)_nomp so we need to calculate p(x)_mp and p(x)_nomp
    px_nomp, px_mp = Px(L_nomp), Px(L_mp)
    # Calculation: only return d-1 values, ignoring the first age class because it is 1 in both matrices. so here indexes=[0,9) represent survival probabilities to age classes [1,10)
    δp = px_mp[:, 1:] - px_nomp[:, 1:] 

    return δp

# %% [code]
def reproductive_value(lx, mx, ρ):
    '''
    Parameters
    ----------
    lx : np.ndarray
            a vector of female cumulative survival probabilities (see "survival2x"). Male values can be added in a second row.
            (Since some computations require only female values because she's the donor, I allowed a seperate handling of male and female values where the male values are optional in the vector.)
    mx : np.ndarray
            a vector of female fecundity values. Male values can be added in a second row.
    ρ : float 
            population growth rate.
    
    Description
    -----------
    This is v(x) in the model equation where x is the age class. 
    This function calculates the sex specific reproductive value of each age class (eq. 4), according to Charlesworth's model (1981). It's a measure for the contribution of an individual of a given age and sex to the future growth of the population.  
    Therefore, older parents will have lower values of the reproductive value weightings, whereas younger parents will have higher reproductive value weightings. This helps understand why the middle ages are more interesting when looking at menopause onset age. 
    At older ages, the reproductive value is low, so perhaps reproduction cessation in these ages is not as costly as in the younger ages where the reproductive value is high.
    '''
    
    # the number of rows in lx and mx must be the same
    assert lx.shape[0] == mx.shape[0] 
    
    # number of age classes = number of columns in lx
    d = lx.shape[-1]
    # k(x) of females (and males but it's optional); k(x) is the net reproduction of each age class, looking at both the fecundity of each age class and the probability to survive to that age class. 
    kx = lx * mx 
    
    if lx.shape[0] == 2 and mx.shape[0] == 2: # if male values are given to the function
        # 2-row age classes
        ages = np.array([np.arange(0,d,1), np.arange(0,d,1)]) 
        # A matrix of coefficients to use for matrix multiplication
        multiby = np.zeros((2,d)) 
        # print(lx[lx>0])
        # the only relevant values are where the recipients survive, otherwise the reproductive value is 0. 
        multiby[lx>0] = np.exp(ρ*ages[lx>0])/lx[lx>0] 
        # matrix multiplication of the coefficients and the kx values.
        vx = multiby * np.flip(np.cumsum(np.flip(np.exp(-ρ*ages)*kx, axis=1), axis=1), axis=1) # The innermost flip is essential because np.cumsum which is essential for efficient summing over future values works from the first value up to some point, but I need it to work from that point to the end. Meaning, I need to sum the values starting from a certain age to the end, and not up to that age, as cumsum would give. The outermost flip is because I need the matrix to return to the correct age indices, now that the correct values are present.
    
    else: # if only female values are given; all above explanations on the calculations apply here in the same manner.
        # 1-row age classes
        ages = np.arange(0,d,1) 
        multiby = np.zeros(d)
        multiby[lx>0] = np.exp(ρ*ages[lx>0])/lx[lx>0]
        vx = multiby * np.flip(np.cumsum(np.flip(np.exp(-ρ*ages)*kx)))

    return vx

# %% [code]
def survival_effect(L_nomp, L_mpR, L_mpD, ρ, age_dist, r_to_pod, sex_ratio=[0.6,0.4], return_all=False, ε=10**(-5)): 
    '''
    Parameters
    ----------
    L_nomp : np.ndarray
            (nomp stands for no-menopause) A leslie matrix representing the population without meopause. Meaning, one where the female reproduces until the end of her life (with reasonable fecundity values due to reproductive scenescence). 
            This is the theoretical scenario.
    L_mpR : np.ndarray
            (mp=menopause, R=recipient) A leslie matrix representing the population with menopause, from the recipients' angle. 
            Female recipients will have fecundity values as the donor (because we consider the donor's relatives by multiplying by r, the relatedness coefficient, which is also the probability these individuals will have the menopause allele), which can change by some factor if we consider the donor helps reproduction. 
            The survival values will also change by some factor compared to the donor, because we assume the donor helps the recipient survive. 
            The males' values can be changed in a similar manner, where fecundity and survival in a menopausal population are increased compared to a non-menopausal one. 
            This is the actual scenario because when old females are absent (analogous to non-donor old females in the throretical scenario), young males and females exhibit decreased survival and fecundity (more specifications on that in the "different_onset_ages" function).
    L_mpD : np.ndarray 
            (D=donor) A leslie matrix representing the population with menopause, from the donors' angle. According to the specific onset age, it will have 0's in the fecundity values of ages starting from the onset age. 
            Also, the fecundity values that are not 0 and the survival values will be lower by some factor than that of the recipients' since we assume the donor cannot be a recipient so she does not get any help with survival and reproduction. 
            That's because we assume the initial frequency of menopause in the population is low enough so that two donors will not interact (i.e, the donor cannot be also a recipient).
    ρ : float
            The population growth rate. It is the leading eigenvalue of the whole population's Leslie matrix.
    age_dist : np.ndarray
            sex-specific age distribution. 
    r_to_pod : np.ndarray
            the donors' local relatedness to males and females in the pod (Orca group) in all her life (10 age classes). 
    sex_ratio : list
            proportion of females and males in the group.
    return_all : boolean 
            whether to return all the values (C, B and rB also) or only the rB-C values. 
    ε : float
            a very small number representing a threshold for a number to be considered as 0. This is because some values are very small, sometimes because of python's inaccuracy, so I ignore them. 
            However, since there is a multiplication by a negative exponent, relevant values can be very small. So I take a very small ε (default is 10^(-5)) to make sure I don't ignore values that are actually relevant.

    Description
    -----------
    This function calculates the survival effect menopause has on the population - both the donor (menopausal female) and the recipient, throughout the entire life of the donor.
    First it calculates the survival cost (C) of menopause for the donor at each age and its benefit (B) for the recipient at each age. The survival benefit is then multiplied by r, the relatedness coefficient, to provide rB values. 
    Then the effective rB-C value is calculated from the survival effects on the donor and recipient. 
    Each donor age class is a column in the returned (2,d) array and is aligned with the corresponding rB-C value, which are the sum of all rB-C values over all recipient ages. 
    The separation to 2 recipient sexes is maintained (that's why 2 rows). C, B and rB values in the same format can also be returned (view "return_all").
    '''
    
    # d age classes
    d = L_nomp.shape[0]//2
    # Calculate cumulative survival probabilities for the recipients (R) and donors (D)
    lx_mpR, lx_mpD = survival2x(L_mpR), survival2x(L_mpD) 
    # Calculate fecundity values for the recipients and donors
    mx_mpR, mx_mpD = mx(L_mpR), mx(L_mpD) 
    # Discount factor is e^(-ρ*age)
    ages = np.arange(0,d+1,1)
    discount = np.exp(-(ρ)*ages) 
    
    # Looking only at female donors' survival probabilities because they are the ones exhibiting menopause
    lxD = lx_mpD[0] 
    # Calculating v(x)=reproductive value for female donors 
    rep_valD =  reproductive_value(lxD, mx_mpD[0,:], ρ)
    # Calculating v(x) for all recipients
    rep_valR =  reproductive_value(lx_mpR, mx_mpR, ρ) 
    
    # Calculate the survival differences of the donor due to menopause. Then, consider the reproductive value for correct weighting of that survival difference.
    δPxD = np.concatenate([δPx(L_nomp, L_mpD)[0] * rep_valD[1:], [0]], axis=0) #  The [1:] is because when we look at the first age class, we will only look at the values after it because of how the model is constructed. So the first age class gets the value of the second, the second gets that of the third and so on. The last one gets 0 and that's why it's concatenated.
    # Calculate the survival differences of the recipient due to menopause.
    δPxR = np.concatenate([δPx(L_nomp, L_mpR) * rep_valR[:, 1:], np.zeros((2,1))], axis=1)
    
    # The rB - C (r=relatedness coefficient, B=benefit, C=cost - like in Hamilton's rule) values will be saved in an array with shape (2, d, d) - 2=female+male, first d=number of donor age groups, second d=number of recipient age groups. So I calculate the survival effect for each recipient age group. In the end, I will sum over the recipient age groups to get the total effect of menopause on the population. Following that, I will sum over the donor age groups to get the total effect of menopause on the donor's life (but in a different function - "tot_rBC_all_donor_life").
    # rBC = np.zeros((2, d, d))
    rBC = np.zeros((2, d))
    # The cost of altruism - only the donor's values
    # COST = np.zeros((2, d, d)) 
    COST = np.zeros((2, d)) 
    # The benefit of altruism - only the recipient's values
    BENEFIT = np.zeros((2, d, d)) 
    # The benefit of altruism, when considering the relatedness of the donor to the recipient. This is the value that will be used in the rB-C expression.
    rxB = np.zeros((2, d, d)) 

    for g in range(2): # iterating over 2 rows, that of the female (0) and that of the male (1)

        for ageD in range(d): # iterating over the donor age groups
            # The donor's probability to survive to ageD. This is l(x_i) in the model equation
            p_D_survival = lxD[ageD] 
            # The change in the donor's survival probability to get to age x_i+1=ageD+1. This is δ_{D}P(x_i) in the model equation
            Δsurvival_D = δPxD[ageD] 
            # The cost (donor) = probability to survive to ageD * change in survival probability to next age (weighted by v(ageD+1)) * discount factor
            cost = p_D_survival * Δsurvival_D * discount[ageD+1] 
            
            for ageR in range(ageD):#range(d):# # iterating over the recipient age groups
                if lx_mpR[g, ageR]<=0: # if the recipient cannot survive to that age
                    # stop iterating over this recipient, all of the next values will =0. More relevant to males which have p(x)=0 earlier than females.
                    break; 
                
                # Current weight of the interaction, according to the sex-specific age distribution of the recipients
                interaction_weight = age_dist[g, ageR]
                # The change in the recipient's survival probability at age x_j=ageR, to get to the next age. This is δ_{R}P(x_j) in the model equation
                Δsurvival_R = δPxR[g, ageR] 
                # Relatedness coefficient of the donor to the recipient
                r = r_to_pod[g, ageD]

                # Normalizing the donor's cost by interaction's weight
                # cost *= interaction_weight 
                # Benefit (recipient) calculation
                benefit = p_D_survival * Δsurvival_R * interaction_weight * sex_weight(g, sex_ratio=sex_ratio) * discount[ageD+1]
                # Multiplying the benefit by the relatedness coefficient
                rXb = r*benefit 
                
                # If the cost is too close to 0, we consider it as 0. Otherwise, we save the value in the array. This explanation applies to all the following lines in this loop.
                # COST[g, ageD, ageR] = cost if abs(cost) > ε else 0 
                BENEFIT[g, ageD, ageR] = benefit if abs(benefit) > ε else 0
                rxB[g, ageD, ageR] = rXb if abs(rXb) > ε else 0
                # rBC[g, ageD, ageR] = COST[g, ageD, ageR] + rxB[g, ageD, ageR]

            COST[g, ageD] = cost if abs(cost) > ε else 0 
            rBC[g, ageD] = COST[g, ageD] + np.sum(rxB, axis=2)[g, ageD]
            
    if return_all:
        return np.array([rBC, COST, np.sum(BENEFIT, axis=2), np.sum(rxB, axis=2)])
        # return np.array([np.sum(rBC, axis=2), np.sum(COST, axis=2), np.sum(BENEFIT, axis=2), np.sum(rxB, axis=2)]) # return all the values: rB-C (as described below), C, B, rB
    else:
        return np.sum(rBC, axis=2) # array containing: 2 rows, for sex-specificity (0 = female, 1 = male); 10 columns per row, each for different donor ages. Each column contains the total survival effect of menopause on the population, for a specific donor age.


# %% [markdown]
## Modeling altrusim's fecundity effects
Perturbations to the characteristic equation introduced by interactions involving the fecundities of either donor or recipient. 
See Methods for detailed maths.

# %% [code]
def fecundity_effect(L_nomp, L_mpR, L_mpD, ρ, age_dist, r_to_pod, sex_ratio=[0.6,0.4], return_all=False, ε=10**(-5)):
    '''
    Parameters
    ----------
    see "survival_effect".

    Description
    -----------
    This function calculates the fecundity effect menopause has on the population - both the donor (menopausal female) and the recipient, throughout the entire life of the donor.
    First it calculates the fecundity cost (C) of menopause for the donor at each age and its benefit (B) for the recipient at each age. The fecundity benefit is then multiplied by r, the relatedness coefficient, to provide rB values. 
    Then the effective rB-C value is calculated from the fecundity effects on the donor and recipient. 
    '''

    # d age classes
    d = L_nomp.shape[0]//2
    # The only survival values we need are those of the donor because she is the one exhibiting menopause and the entire altruistic act is based on her survival 
    lx_mpD = survival2x(L_mpD) 
    # The fecundity values in the absence of menopause, in the presence of menopause from the recipients' angle, and in the presence of menopause from the donors' angle
    mx_nomp, mx_mpR, mx_mpD = mx(L_nomp), mx(L_mpR), mx(L_mpD) 
    # The effect of menopause on the donor's fecundity, only female
    δmD = mx_mpD[0] - mx_nomp[0]
    # The effect of menopause on the recipient's fecundity, both male and female
    δmR = mx_mpR - mx_nomp 
    # Discount factor is e^(-ρ*age)
    ages = np.arange(0,d,1)
    discount = np.exp(-(ρ)*ages) 
    
    # The rB - C values will be saved in an array with shape (2, d, d) - 2=female+male, first d=number of donor age groups, second d=number of recipient age groups. So I calculate the survival effect for each recipient age group. In the end, I will sum over the recipient age groups to get the total effect of menopause on the population. Following that, I will sum over the donor age groups to get the total effect of menopause on the donor's life.
    rBC = np.zeros((2,d,d)) 
    # rBC = np.zeros((2,d)) 
    # The cost of altruism - only the donor's values
    COST = np.zeros((2, d, d)) 
    # COST = np.zeros((2, d))
    # The benefit of altruism - only the recipient's values
    BENEFIT = np.zeros((2, d, d)) 
    # The benefit of altruism, when considering the relatedness of the donor to the recipient. This is the value that will be used in the rB-C expression.
    rxB = np.zeros((2, d, d)) 

    for g in range(2): # iterating over 2 rows, that of the female (0) and that of the male (1)

        for ageD in range(d): # iterating over the donor age groups
            # The donor's probability to survive to age x_i=ageD. l(x_i) in the model equation
            p_D_survival = lx_mpD[0, ageD]
            # The change in the donor's fecundity at age x_i=ageD. δm(x_i) in the model equation
            Δfecundity_D = δmD[ageD] 

            # The cost of altruism (donor)
            # cost = Δfecundity_D * p_D_survival * discount[ageD]
            # print(ageD, ':', Δfecundity_D, p_D_survival, discount[ageD])
            
            for ageR in range(ageD):#range(d): # iterating over the recipient age groups
                # Current weight of the interaction, according to the sex-specific age distribution of the recipients
                interaction_weight = age_dist[g, ageR] 
                # The change in recipient's fecundity at age x_j=ageR. δm(x_j) in the model equation
                Δfecundity_R = δmR[g, ageR] 
                # Relatedness coefficient of the donor to the recipient
                r = r_to_pod[g, ageD] 
                
                # # The cost of altruism (donor)
                cost = interaction_weight * Δfecundity_D * p_D_survival * discount[ageD] 
                # The benefit of altruism (recipient)
                benefit = sex_weight(g, sex_ratio=sex_ratio) * interaction_weight * Δfecundity_R * p_D_survival * discount[ageD] 
                # The benefit of altruism, when considering the relatedness of the donor to the recipient
                rxb = r*benefit 
                
                # If the cost is very small, we consider it as 0. Otherwise, we save the value in the array. This explanation applies to all the following lines in this loop.
                COST[g, ageD, ageR] = cost if abs(cost) > ε else 0
                BENEFIT[g, ageD, ageR] = benefit if abs(benefit) > ε else 0
                rxB[g, ageD, ageR] = rxb if abs(rxb) > ε else 0
                rBC[g, ageD, ageR] = COST[g, ageD, ageR] + rxB[g, ageD, ageR]
        
            # COST[g, ageD] = cost if abs(cost) > ε else 0        
            # # print(COST[g, ageD])
            # rBC[g, ageD] = COST[g, ageD] + np.sum(rxB, axis=2)[g, ageD]

    # print(np.array([rBC, COST, np.sum(BENEFIT, axis=2), np.sum(rxB, axis=2)[g, ageD]]))
    if return_all:
        # return np.array([rBC, COST, np.sum(BENEFIT, axis=2), np.sum(rxB, axis=2)])
        return np.array([np.sum(rBC, axis=2), np.sum(COST, axis=2), np.sum(BENEFIT, axis=2), np.sum(rxB, axis=2)]) # return all the values: rB-C (as described below), C, B, rB
    else:
        return np.sum(rBC, axis=2) # array containing: 2 rows, for sex-specificity (0 = female, 1 = male); 10 columns per row, each for different donor ages. Each column contains the total survival effect of menopause on the population, for a specific donor age.
    

# %% [code]
def fecundity_effect_cost_noAgeDist(L_nomp, L_mpR, L_mpD, ρ, age_dist, r_to_pod, sex_ratio=[0.6,0.4], return_all=False, ε=10**(-5)):
    '''
    Parameters
    ----------
    see "survival_effect".

    Description
    -----------
    This function calculates the fecundity effect menopause has on the population - both the donor (menopausal female) and the recipient, throughout the entire life of the donor.
    First it calculates the fecundity cost (C) of menopause for the donor at each age and its benefit (B) for the recipient at each age. The fecundity benefit is then multiplied by r, the relatedness coefficient, to provide rB values. 
    Then the effective rB-C value is calculated from the fecundity effects on the donor and recipient. 
    '''

    # d age classes
    d = L_nomp.shape[0]//2
    # The only survival values we need are those of the donor because she is the one exhibiting menopause and the entire altruistic act is based on her survival 
    lx_mpD = survival2x(L_mpD) 
    # The fecundity values in the absence of menopause, in the presence of menopause from the recipients' angle, and in the presence of menopause from the donors' angle
    mx_nomp, mx_mpR, mx_mpD = mx(L_nomp), mx(L_mpR), mx(L_mpD) 
    # The effect of menopause on the donor's fecundity, only female
    δmD = mx_mpD[0] - mx_nomp[0]
    # The effect of menopause on the recipient's fecundity, both male and female
    δmR = mx_mpR - mx_nomp 
    # Discount factor is e^(-ρ*age)
    ages = np.arange(0,d,1)
    discount = np.exp(-(ρ)*ages) 
    
    # The rB - C values will be saved in an array with shape (2, d, d) - 2=female+male, first d=number of donor age groups, second d=number of recipient age groups. So I calculate the survival effect for each recipient age group. In the end, I will sum over the recipient age groups to get the total effect of menopause on the population. Following that, I will sum over the donor age groups to get the total effect of menopause on the donor's life.
    # rBC = np.zeros((2,d,d)) 
    rBC = np.zeros((2,d)) 
    # The cost of altruism - only the donor's values
    # COST = np.zeros((2, d, d)) 
    COST = np.zeros((2, d))
    # The benefit of altruism - only the recipient's values
    BENEFIT = np.zeros((2, d, d)) 
    # The benefit of altruism, when considering the relatedness of the donor to the recipient. This is the value that will be used in the rB-C expression.
    rxB = np.zeros((2, d, d)) 

    for g in range(2): # iterating over 2 rows, that of the female (0) and that of the male (1)

        for ageD in range(d): # iterating over the donor age groups
            # The donor's probability to survive to age x_i=ageD. l(x_i) in the model equation
            p_D_survival = lx_mpD[0, ageD]
            # The change in the donor's fecundity at age x_i=ageD. δm(x_i) in the model equation
            Δfecundity_D = δmD[ageD] 

            # The cost of altruism (donor)
            cost = Δfecundity_D * p_D_survival * discount[ageD]
            # print(ageD, ':', Δfecundity_D, p_D_survival, discount[ageD])
            
            for ageR in range(ageD):#range(d): # iterating over the recipient age groups
                # Current weight of the interaction, according to the sex-specific age distribution of the recipients
                interaction_weight = age_dist[g, ageR] 
                # The change in recipient's fecundity at age x_j=ageR. δm(x_j) in the model equation
                Δfecundity_R = δmR[g, ageR] 
                # Relatedness coefficient of the donor to the recipient
                r = r_to_pod[g, ageD] 
                
                # # The cost of altruism (donor)
                # cost = interaction_weight * Δfecundity_D * p_D_survival * discount[ageD] 
                # The benefit of altruism (recipient)
                benefit = sex_weight(g, sex_ratio=sex_ratio) * interaction_weight * Δfecundity_R * p_D_survival * discount[ageD] 
                # The benefit of altruism, when considering the relatedness of the donor to the recipient
                rxb = r*benefit 
                
                # If the cost is very small, we consider it as 0. Otherwise, we save the value in the array. This explanation applies to all the following lines in this loop.
                # COST[g, ageD, ageR] = cost if abs(cost) > ε else 0
                BENEFIT[g, ageD, ageR] = benefit if abs(benefit) > ε else 0
                rxB[g, ageD, ageR] = rxb if abs(rxb) > ε else 0
                # rBC[g, ageD, ageR] = COST[g, ageD, ageR] + rxB[g, ageD, ageR]
        
            COST[g, ageD] = cost if abs(cost) > ε else 0        
            # print(COST[g, ageD])
            rBC[g, ageD] = COST[g, ageD] + np.sum(rxB, axis=2)[g, ageD]

    # print(np.array([rBC, COST, np.sum(BENEFIT, axis=2), np.sum(rxB, axis=2)[g, ageD]]))
    if return_all:
        return np.array([rBC, COST, np.sum(BENEFIT, axis=2), np.sum(rxB, axis=2)])
        # return np.array([np.sum(rBC, axis=2), np.sum(COST, axis=2), np.sum(BENEFIT, axis=2), np.sum(rxB, axis=2)]) # return all the values: rB-C (as described below), C, B, rB
    else:
        return np.sum(rBC, axis=2) # array containing: 2 rows, for sex-specificity (0 = female, 1 = male); 10 columns per row, each for different donor ages. Each column contains the total survival effect of menopause on the population, for a specific donor age.
    

# %% [markdown]
## Calculating total effects over the donor ages
$\lambda_{tot} = \lambda_{surv} + \lambda_{fec}$

# %% [code]
def tot_rBC(L_nomp, L_mpR, L_mpD, ρ, age_dist, r_to_pod, sex_ratio=[0.6,0.4], return_all=False, ε=10**(-5)):
    '''
    Parameters 
    ----------
    see "survival_effect".

    Description
    -----------
    This function calculates the total effect menopause has on the population, considering both survival and fecundity effects on donors (menopausal female) and recipients by summing them, over each donor age class. 
    It does so by calling the "survival_effect" and "fecundity_effect" functions and summing the values they return. As known, they return arrays of shape (2,d) so this is also the shape of the returned array here.
    This array will hold the total rB-C values achieved from both survival and fecundity effects, in each donor age. C, B and rB values in the same format can also be returned (with "return_all").
    '''

    # rB-C values of survival effect
    rBC_survival = survival_effect(L_nomp, L_mpR, L_mpD, ρ, age_dist, r_to_pod, sex_ratio=sex_ratio, return_all=return_all, ε=ε) 
    # rB-C values of fecundity effect
    rBC_fecundity = fecundity_effect(L_nomp, L_mpR, L_mpD, ρ, age_dist, r_to_pod, sex_ratio=sex_ratio, return_all=return_all, ε=ε) 
    # Summing the two effects
    rBC_all_effects = rBC_survival + rBC_fecundity 

    return rBC_all_effects

# %% [code]
def tot_rBC_costNoAgeDist(L_nomp, L_mpR, L_mpD, ρ, age_dist, r_to_pod, sex_ratio=[0.6,0.4], return_all=False, ε=10**(-5)):
    '''
    Parameters 
    ----------
    see "survival_effect".

    Description
    -----------
    This function calculates the total effect menopause has on the population, considering both survival and fecundity effects on donors (menopausal female) and recipients by summing them, over each donor age class. 
    It does so by calling the "survival_effect" and "fecundity_effect" functions and summing the values they return. As known, they return arrays of shape (2,d) so this is also the shape of the returned array here.
    This array will hold the total rB-C values achieved from both survival and fecundity effects, in each donor age. C, B and rB values in the same format can also be returned (with "return_all").
    '''

    # rB-C values of survival effect
    rBC_survival = survival_effect(L_nomp, L_mpR, L_mpD, ρ, age_dist, r_to_pod, sex_ratio=sex_ratio, return_all=return_all, ε=ε) 
    # rB-C values of fecundity effect
    rBC_fecundity = fecundity_effect_cost_noAgeDist(L_nomp, L_mpR, L_mpD, ρ, age_dist, r_to_pod, sex_ratio=sex_ratio, return_all=return_all, ε=ε) 
    # Summing the two effects
    rBC_all_effects = rBC_survival + rBC_fecundity 

    return rBC_all_effects

# %% [markdown]
## Summing all contributions over donor ages

# %% [code]
def tot_rBC_all_donor_life(L_nomp, L_mpR, L_mpD, ρ, age_dist, r_to_pod, sex_ratio=[0.6,0.4], return_all=False, ε=10**(-5)):
    '''
    Parameters
    ----------
    see "survival_effect".

    Description
    -----------
    This function calculates the total effect menopause has on the population by taking the rB-C values considering both survival and fecundity effects (from the tot_rBC function) and then summing all the calculated values appearing in each donor age to be left with just one rB-C value for each recipient sex. 
    Therefore, the returned array will be of shape (2,1). These two values and their sum will tell us whether it is beneficial for the donor to stop reproducing with the conditions this function receives (leslie matrices) or not, which imply the onset age, survival effects and fecundity effects. 
    I do this summing over donor ages because according to the model, all the help the donors give throughout their lives can be summed unedr the assumption that A2 allele has a small effect on the frequency of behavioral intercations. 
    '''

    # Sum the survival and fecundity effects
    # tot_effects = tot_rBC(L_nomp, L_mpR, L_mpD, ρ, age_dist, r_to_pod, sex_ratio=sex_ratio, return_all=return_all, ε=ε) 
    tot_effects = tot_rBC_costNoAgeDist(L_nomp, L_mpR, L_mpD, ρ, age_dist, r_to_pod, sex_ratio=sex_ratio, return_all=return_all, ε=ε) 
    # Sum all effects of donor's altruism throughout her life and return the sum 
    return [np.sum(tot_effects[element], axis=1) for element in range(tot_effects.shape[0])] 

# %% [markdown]
## Calculating the values over different menopause onset ages
= when the zeros in the fecundity of the females start appearing

# %% [markdown]
### Stop early pathway

# %% [code]
def different_onset_ages(L_no_effects, r_to_pod, Δmaternal_mortality=1, mx_contribution=np.array([0.95, 0.95]), lx_contribution=np.array([0.95, 0.95]), sex_ratio=[0.6,0.4], return_all=False, ε=10**(-5), max_male_help=None, max_female_help=None, age_dist=None, ρ=None, plot_age_dist=False):

    # d age classes
    d = L_no_effects.shape[0]//2
    if type(lx_contribution) != np.ndarray: # if the user did not input an array, convert it to one, this format is necessary for the calculations
        lx_contribution = np.array(lx_contribution)
    if type(mx_contribution) != np.ndarray: # if the user did not input an array, convert it to one, this format is necessary for the calculations
        mx_contribution = np.array(mx_contribution)
    
    # Calculate the survival probabilities of the population without any effects (no menopause and no reduction in survival due to lack of help)
    px_no_effects= Px(L_no_effects)
    # Calculate the fecundity values of the population without any effects (no menopause and no reduction in survival due to lack of help)
    mx_no_effects = mx(L_no_effects) 
    
    # Create an array to be filled with the contribution factor to divide the survival values by, for males and females separately 
    lx_contribution_arr = np.ones((2,d)) 
    lx_contribution_arr[1, :] = lx_contribution[1]

    # Matrices to save the results: C, B, rB
    COST = np.zeros((2, d)) 
    BENEFIT = np.zeros((2, d))
    rxB = np.zeros((2, d))
    if plot_age_dist:
        age_dist_each_onset = np.zeros((d, 2, d))
        rho_each_onset = np.zeros(d)

    if max_male_help is None:
        max_male_help = d
    if max_female_help is None:
        max_female_help = d
    
    for onset in range(d): # iterating over all possible onset ages
        # Indexes of the ages that will get a benefit from the donor, the ages are < onset age because only younger individuals can be recipients
        indexes_benefit_female = np.arange(0, min(onset, max_female_help), 1) 
        indexes_benefit_male = np.arange(0, min(onset, max_male_help), 1) 
        # Indexes of the ages of menopause, in which a fecundity cost is "paid" by the donor (but survival in menopause ages *can* be improved)
        indexes_cost_female = np.arange(onset, d, 1) 
        indexes_cost_male = np.arange(min(onset, max_male_help), d, 1) 
        # The donor's survival is improved by ΔPRL_survival in menopause ages, so lower the survival values of non-menopausal females in these ages by that factor
        lx_contribution_arr[0,:] = lx_contribution[0] 
        lx_contribution_arr[0, indexes_cost_female] *= Δmaternal_mortality 

        ##### Non-menopausal population leslie matrix: positive fecundity values in the altruists' menopausal ages, but with general lowered survival and fecundity due to lack of help #####
        # Generate a non-menopausal population leslie matrix, with the lowered survival and fecundity values but with positive fecundity values in the older ages (mx_no_effects), as opposed to 0. This is the theoretical scenario.
        L_nomp_weffects = leslie_(mx_no_effects*(mx_contribution[:, np.newaxis]), px_no_effects[:, 1:]*(lx_contribution_arr[:,1:])) 
        
        # Get the survival probabilities of the non-menopausal population with the lowered recipient survival 
        px_nomp_weffects = Px(L_nomp_weffects) 
        # Get the fecundity values of the non-menopausal population with the lowered fecundity but positive fecundity in the menopausal ages
        mx_nomp_weffects = mx(L_nomp_weffects) 
    
        ##### Recipient (R) Leslie matrix #####
        # Take the "worst" survival values, i.e, the ones that are not improved by the donor's help, and then change them to the "best" survival values, i.e, the ones that are improved by the donor's help, in the indexes of menopause help (indexes_benefit).
        px_mpR = deepcopy(px_nomp_weffects) 
        px_mpR[0, indexes_benefit_female] = px_no_effects[0, indexes_benefit_female] # change the worst survival values in the indexes of being helped to the best ones, represented by those in the L_no_effects matrix 
        px_mpR[1, indexes_benefit_male] = px_no_effects[1, indexes_benefit_male]
        px_mpR[0, indexes_cost_female] = px_no_effects[0, indexes_cost_female]*lx_contribution[0]
        # Take the "worst" fecundity values, i.e, the ones that are not improved by the donor's help, and then change them to the "best" fecundity values, i.e, the ones that are improved by the donor's help, in the indexes of menopause help (indexes_benefit).
        mx_mpR = deepcopy(mx_nomp_weffects) 
        mx_mpR[0, indexes_benefit_female] = mx_no_effects[0, indexes_benefit_female] # change the worst fecundity values in the indexes of being helped to the best ones, represented by those in the L_no_effects matrix
        mx_mpR[1, indexes_benefit_male] = mx_no_effects[1, indexes_benefit_male]
        mx_mpR[0, indexes_cost_female] = np.zeros(indexes_cost_female.shape[0]) # since the recipient is assumed to be related to the donor by r, the relatedness coefficient, the female recipient's fecundity in the donor's menopause ages is 0 
        # Generate the leslie matrix of the recipient population, with the improved survival and fecundity values. 
        L_mpR = leslie_(mx_mpR, px_mpR[:, 1:]) 
        
        ##### Donor (D) Leslie matrix #####
        # I don't increase the survival and fecundity values of the donor in the ages < onset because she is not a recipient because of the assumption that initially this trait is rare, so she does not get any help from any donors.
        # Take the "worst" survival values, i.e, the ones that are not improved by the donor's help, because the donor is not a recipient. Only change the post reproductive life (indexes_cost) survival to the better values, because the donor's survival is improved by ΔPRL_survival in these ages.
        ''' This part may be the reason for the weird lambda as a funtion of help graph:
        # px_mpD = np.array([deepcopy(px_no_effects[0]), deepcopy(px_nomp_weffects[1])])
        # mx_mpD = np.array([deepcopy(mx_no_effects[0]), deepcopy(mx_nomp_weffects[1])])
        # mx_mpD[0, indexes_cost_female] = np.zeros(indexes_cost_female.shape[0])
        '''
        px_mpD = deepcopy(px_nomp_weffects) 
        px_mpD[0, indexes_cost_female] = px_no_effects[0, indexes_cost_female]*lx_contribution[0] # the donor's survival in her menopause ages is improved by ΔPRL_survival but not by lx_contribution[0]. 
         # Take the "worst" fecundity values, i.e, the ones that are not improved by the donor's help, because the donor is not a recipient. Only change the post reproductive life (indexes_cost) fecundity to 0 because of menopause
        mx_mpD = deepcopy(mx_nomp_weffects)
        mx_mpD[0, indexes_cost_female] = np.zeros(indexes_cost_female.shape[0]) # the donor's fecundity in her menopause ages is 0
        # generate the leslie matrix of the donor population 
        L_mpD = leslie_(mx_mpD, px_mpD[:, 1:]) 

        # Calculate the population growth rate of the non-menopausal population since that is the dominant population in the model, as menopause is rare in the beginning. The matrix needs to be in a (d,d) shape for the eigenvalue calculation so I take the average according to the sex ratio. 
        if ρ is None and age_dist is None:    
            ρ = np.linalg.eigvals(sex_ratio[0]*L_nomp_weffects[:d,:d]+sex_ratio[1]*L_nomp_weffects[d:,d:]).max().real 
            # Calculate the age distribution of the non-menopausal population (L_nomp_weffects)
            age_dist = age_distribution(L_nomp_weffects, ρ, sex_ratio=np.array([[sex_ratio[0]], [sex_ratio[1]]])) 
        
        # Calculate the total effect of menopause on the population, using L_mpR as the leslie matrix of the recipient population and L_mpD as the leslie matrix of the donor population. L_nomp_weffects is the leslie matrix of the non-menopausal population, which is the theoretical scenario.
        results = tot_rBC_all_donor_life(L_nomp_weffects, L_mpR, L_mpD, ρ, age_dist, r_to_pod, sex_ratio=sex_ratio, return_all=return_all, ε=ε) 
        # results = tot_rBC_all_donor_life_costNoAgeDist(L_nomp_weffects, L_mpR, L_mpD, ρ, age_dist, r_to_pod, sex_ratio=sex_ratio, return_all=return_all, ε=ε) 
        
        if plot_age_dist:
            age_dist_each_onset[onset, :, :] = age_dist
            rho_each_onset[onset] = ρ   
        
        # Save the results
        COST[:,onset] = results[1]
        BENEFIT[:,onset] = results[2]
        rxB[:,onset] = results[3]
    
    # Calculate rB-C
    rBC = rxB + COST # Cost is non-positive so it is a subtraction, rB-abs(C) = rB+C
    if return_all and not plot_age_dist:
        return rBC, COST, BENEFIT, rxB
    elif return_all and plot_age_dist:
        return rBC, COST, BENEFIT, rxB, (age_dist_each_onset, rho_each_onset)
    else:
        return rBC 


# %% [code]
def latex_array(numpy_array, epsilon=10^(-6)):
    # Start the string with the LaTeX matrix environment
    latex_str = "\\begin{bmatrix}\n"
    
    # Loop through each row in the array
    for row in numpy_array:
        # Join the elements of the row with " & " for LaTeX matrix formatting
        # print(str(x), str(int(x)))
        to_join = [str(int(x)) if int(x) - float(x) < epsilon else f"{x:.4f}" for x in row] 
        latex_str += " & ".join(to_join)
        latex_str += " \\\\"  # End each row with a new line and LaTeX row separator
        
    # Close the LaTeX matrix environment
    latex_str += "\\end{bmatrix}"
    
    return latex_str

# %% [code]
def plot_post_reproductive_life(species, percentages, ci_intervals, actual_values, colors, LM_values=None, barcolor='gold', alphabar=0.2, phylopics=None, yposition_pics=None, zoom=None, ylims=None):
    """
    Plots a bar plot of the percentage of post-reproductive life for various species.

    Parameters:
    - species (list): A list of species names (strings).
    - percentages (list): A list of percentages representing post-reproductive life.
    - ci_intervals (list): A list of confidence intervals (as floats or lists of lower/upper bounds).
    - actual_values (list of lists): Each species has multiple actual values, represented as a nested list.
    - colors (list of lists): Colors corresponding to actual values.
    - LM_values (list, optional): Values to be plotted as "LM" dots.
    - barcolor (str, optional): Color for the bars (default: 'gold').
    - alphabar (float, optional): Alpha transparency for bars (default: 0.2).
    - phylopics (list, optional): List of images to overlay on the plot (ensure add_image function is defined).
    - yposition_pics (list, optional): Y-axis positions for the images.
    - ylims (list with 2 elements, optional): Y-axis bottom and upper boundaries.
    """
    x_positions = np.arange(len(species))  # X-axis positions for the bars

    # Plotting
    fig, ax = plt.subplots(figsize=(10, 8))
    ax.bar(x_positions, percentages, yerr=ci_intervals, capsize=5, color=barcolor, alpha=alphabar)
    for i in range(len(x_positions)):
        if phylopics is not None:
            add_image(ax, phylopics[i], xy=(i, yposition_pics[i]), zoom=zoom)
        for j in range(len(actual_values[i])):
            ax.scatter(x_positions[i], actual_values[i][j], color=colors[i][j], zorder=5, label='According to literature', alpha=0.95, s=50)  # Add dots for actual values
    if LM_values is not None:
        ax.scatter(x_positions, LM_values, color='k', zorder=5, label='According to LM', size=3)  # Add dots for the actual LM values
    ax.set_xticks(x_positions)
    ax.set_xticklabels(species, rotation=45, ha='right', fontsize=17)
    ax.tick_params(axis='y', labelsize=18)
    ax.set_ylabel('PrR', fontsize=21)
    if ylims is not None:
        ax.set_ylim(ylims)
    # plt.xlabel('Species', fontsize=15)
    # plt.title('Percentage of Post-Reproductive Life Across Species')
    # plt.legend()
    plt.tight_layout()
    plt.savefig('PrR_results.png')
    plt.show()

# %% [code]
def ci_intervals(leslie_matrices, age_intervals):

    ci_interval_arr = np.zeros(len(leslie_matrices))
    
    for i, LM in enumerate(leslie_matrices): 
        total_lifespan = age_intervals[i] * LM.shape[0]//2
        print(age_intervals[i], total_lifespan)
        ci_interval_arr[i] = age_intervals[i] / total_lifespan

    return ci_interval_arr

# %% [code]
def remaining_life_expectancy(lx, x, age_interval):
    """
    Calculate the expected number of years remaining at age x.
    
    Parameters:
    lx (array): Survival probabilities (proportion of individuals alive at each age).
    x (int): Age at which to calculate remaining life expectancy.
    age_interval (int): Corresponding age interval for lx values.
    
    Returns:
    float: Expected number of years remaining at age x.
    """
    # Calculate L(x) - the number of person-years lived in each interval
    # Lx = [(lx[i] + lx[i + 1]) / 2 for i in range(len(lx) - 1)]
    
    # Calculate T(x) - the cumulative sum of future person-years from age x onward
    # Tx = np.cumsum(Lx[::-1])[::-1]  # Reverse cumulative sum
    Tx = np.cumsum(lx[::-1])[::-1]  # Reverse cumulative sum

    # Step 5: Calculate e_x = T(x) / l(x)
    ex = Tx[x] / lx[x]
    
    return ex

# %% [code]
def PRL(menopause_onset, leslie_matrices, age_intervals):
    total_lifespans = [(leslie_matrices[i].shape[0] // 2) * age_intervals[i] for i in range(len(leslie_matrices))]
    print(total_lifespans)
    return [(total_lifespans[i] - menopause_onset[i]) / total_lifespans[i] for i in range(len(total_lifespans))]

# %% [code]
def PrR_from_LM(leslie_matrices, age_intervals):
    
    fecund_lifespan_stats = np.empty((len(leslie_matrices), 4)) # columns are lM, lB, eM, eB and rows are species
    PrRs = np.empty(len(leslie_matrices))
    total_lifespans = [(leslie_matrices[i].shape[0] // 2) * age_intervals[i] for i in range(len(leslie_matrices))]
    
    for i, LM in enumerate(leslie_matrices):
        d = LM.shape[0] // 2
        fecund_lifespan_indices = np.where(LM[0, :d] != 0)[0]
        start_fecund, end_fecund = fecund_lifespan_indices[0], fecund_lifespan_indices[-1] + 1
        lX, _ = survival2x(LM)
        lB = lX[start_fecund] if start_fecund != 0 else 1
        lM = lX[end_fecund] if end_fecund != d else 0
        eM = total_lifespans[i] - end_fecund*age_intervals[i]
        eB = total_lifespans[i] - start_fecund*age_intervals[i]
        print(i, ':', start_fecund, end_fecund)
        print('lM = {}, lB = {}, eM = {}, eB = {}'.format(lM, lB, eM, eB))

        fecund_lifespan_stats[i] = lM, lB, eM, eB
        PrRs[i] = (lM / lB) * (eM / eB)

    return PrRs

# %% [code]
def PrR_from_model(menopause_onset, leslie_matrices, age_intervals): 
    
    total_lifespans = [(leslie_matrices[i].shape[0] // 2) * age_intervals[i] for i in range(len(leslie_matrices))]
    PrRs = np.empty(len(leslie_matrices))
    fecund_lifespan_stats = np.empty((len(leslie_matrices), 4)) # columns are lM, lB, eM, eB and rows are species

    for i, LM in enumerate(leslie_matrices):
        d = LM.shape[0] // 2
        fecund_lifespan_indices = np.where(LM[0, :d] != 0)[0]
        start_fecund, end_fecund = fecund_lifespan_indices[0], menopause_onset[i] // age_intervals[i]
        lX, _ = survival2x(LM)
        lB = lX[start_fecund] if start_fecund != 0 else 1
        lM = lX[end_fecund] if end_fecund != d else 0
        eM = remaining_life_expectancy(lX, end_fecund, age_intervals[i]) # total_lifespans[i] - end_fecund*age_intervals[i]
        eB = remaining_life_expectancy(lX, start_fecund, age_intervals[i]) #total_lifespans[i] - start_fecund*age_intervals[i]
        # print(i, ':', start_fecund, end_fecund, '\n')
        print('lM = {}, lB = {}, eM = {}, eB = {}'.format(lM, lB, eM, eB))
        
        fecund_lifespan_stats[i] = lM, lB, eM, eB
        PrRs[i] = (lM / lB) * (eM / eB)

    return PrRs

# %% [code]
def ci_intervals_by_PrR(menopause_onset, leslie_matrices, age_intervals):
    if type(menopause_onset) != np.ndarray:
        menopause_onset = np.array(menopause_onset)
    if type(age_intervals) != np.ndarray:
        age_intervals = np.array(age_intervals)

    before_onset = menopause_onset - age_intervals
    after_onset = menopause_onset + age_intervals
    print(before_onset, ';', after_onset)
    
    low_bound = PrR_from_model(menopause_onset, leslie_matrices, age_intervals) - PrR_from_model(before_onset, leslie_matrices, age_intervals)
    high_bound = PrR_from_model(after_onset, leslie_matrices, age_intervals) - PrR_from_model(menopause_onset, leslie_matrices, age_intervals)
    # print(low_bound, ';', high_bound)

    return np.array([low_bound, high_bound])

# %% [code]
def add_image(ax, img, xy, zoom=0.1):
    imagebox = OffsetImage(img, zoom=zoom)
    ab = AnnotationBbox(imagebox, xy, frameon=False)
    ax.add_artist(ab)

# %% [code]
def model(x, L_no_effects, r_to_pod, onset=5, sex_ratio=[0.5, 0.5], max_male_help=None, max_female_help=None, adjusted_age_dist_rho=True, ε=10**(-20), age_dist=None, ρ=None):
    if (not adjusted_age_dist_rho) and age_dist is None and ρ is None:
        raise Exception("Age distibution and ρ must be inputs!!")

    if max_male_help is None:
        max_male_help = L_no_effects.shape[0] // 2
    if max_female_help is None:
        max_female_help = L_no_effects.shape[0] // 2
        
    # Replace this with your actual function
    lx_cont = np.array([x[0], x[1]])
    mx_cont = np.array([x[2], x[3]])
    if adjusted_age_dist_rho:
        lamb = different_onset_ages(L_no_effects, r_to_pod,  Δmaternal_mortality=x[4], lx_contribution=lx_cont, mx_contribution=mx_cont, sex_ratio=sex_ratio, ε=ε, return_all=True, max_male_help=max_male_help, max_female_help=max_female_help)[0]
    else:
        lamb = different_onset_ages(L_no_effects, r_to_pod, Δmaternal_mortality=x[4], lx_contribution=lx_cont, mx_contribution=mx_cont, sex_ratio=sex_ratio, ε=ε, return_all=True, age_dist=age_dist, ρ=ρ, max_male_help=max_male_help, max_female_help=max_female_help)[0]
        
    return np.sum(lamb, axis=0)[onset]

# %% [code]
def plot_post_reproductive_life_boxplot(species, percentages, ranges, actual_values, colors, barcolor='gold', alphabar=0.2, phylopics=None, yposition_pics=None):
    """
    Plots a boxplot of the percentage of post-reproductive life for various species.

    Parameters:
    - species (list): A list of species names (strings).
    - percentages (list): A list of PrR values.
    - ranges (list of tuples): A list of tuples (min, max) defining the range of the box for each species.
    - actual_values (list of lists): A list containing lists of actual PrR values for each species.
    - colors (list of lists): Colors for the dots representing the actual values.
    """
    x_positions = np.arange(len(species))  # X-axis positions for the boxplots
    
    # Extract lower and upper bounds for the ranges
    lower_bounds = [r[0] for r in ranges]
    upper_bounds = [r[1] for r in ranges]

    fig, ax = plt.subplots(figsize=(10, 8))

    # Creating the boxplot manually using `vlines` and `patches`
    for i, x in enumerate(x_positions):
        # Draw the box for each species
        plt.fill_betweenx(
            [lower_bounds[i], upper_bounds[i]],
            x - 0.3, x + 0.3,
            color=barcolor,
            alpha=alphabar,
            edgecolor='black',
            linewidth=1.5
        )
        if phylopics is not None:
            add_image(ax, phylopics[x], xy=(x, yposition_pics), zoom=0.1)
        # Add actual values as scatter points
        for j, value in enumerate(actual_values[i]):
            plt.scatter(x, value, color=colors[i][j], zorder=5, alpha=0.95, s=50)

    # Customizing the plot
    plt.xticks(x_positions, species, rotation=45, ha='right', fontsize=14)
    plt.yticks(fontsize=14)
    plt.ylabel('PrR', fontsize=17)
    plt.tight_layout()
    plt.savefig('PrR.png')
    plt.show()


# %% [markdown]
### Live longer pathway

# %% [code]
def create_non_menopausal_pop(L_mp, lx_contribution, mx_contribution, female_fec_tradeoff=1, female_surv_tradeoff=1, additional_years_after_rep=0, Δmaternal_mortality=1):
    """
    Generate a Leslie matrix representing a non-menopausal population derived from a menopausal population matrix.
    
    This function models a life history shift where females:
    - Lose survival and fecundity benefits gained from post-reproductive helping (scaled by lx_contribution and mx_contribution).
    - Reallocate resources from longevity to early-life reproduction and survival (adjusted by female_fec_tradeoff and female_surv_tradeoff).
    
    Parameters:
    - L_mp: Original menopausal Leslie matrix (combined female/male).
    - lx_contribution: Survival scaling factors (e.g., removing helping effects). List or array with 2 elements — first index is for females, second for males.
    - mx_contribution: Fecundity scaling factors (e.g., removing helping effects). List or array with 2 elements — first index is for females, second for males.
    - female_fec_tradeoff: Multiplier to increase early-life female fecundity. Any non-negative number is fine.
    - female_surv_tradeoff: Multiplier to increase early-life female survival. Any non-negative number is fine.
    - additional_years_after_rep: Number of extra age classes added after last reproduction (will be zero for absolute absence of post-reproductive life!).
    
    Returns:
    - L_nomp: Adjusted Leslie matrix representing a population different from our menopausal one.
    """
        
    d = L_mp.shape[0] // 2
    L_mp_females, L_mp_males = L_mp[:d, :d], L_mp[d:, d:]
    mx_F, mx_M = L_mp_females[0, :], L_mp_males[0, :]
    last_reproduction_age = np.where(mx_F > 0)[0][-1]+1 if np.any(mx_F > 0) else -1 # this will also be the last survival age
    female_death_age = last_reproduction_age + additional_years_after_rep
    
    assert female_death_age <= d
    
    px_F, px_M = Px(L_mp)

    # Change for females - survival
    px_F_nohelp = np.zeros(px_F.shape[0])
    px_F_nohelp[:female_death_age] = px_F[:female_death_age] * female_surv_tradeoff
    px_F_nohelp[:last_reproduction_age] *= lx_contribution[0]
    px_F_nohelp[last_reproduction_age:] = px_F[last_reproduction_age:] * lx_contribution[0]
    
    # Change for females - fecundity
    mx_F_nohelp = np.zeros(px_F.shape[0])
    mx_F_nohelp[:female_death_age] = mx_F[:female_death_age] * mx_contribution[0] * female_fec_tradeoff

    # Change for males - survival
    px_M_nohelp = deepcopy(px_M)
    px_M_nohelp[:female_death_age] = px_M[:female_death_age] * lx_contribution[1]
    
    # Change for males - fecundity
    mx_M_nohelp = deepcopy(mx_M)
    mx_M_nohelp[:female_death_age] = mx_M[:female_death_age] * mx_contribution[1]

    # Building the matrix
    mx_nohelp = np.array([mx_F_nohelp, mx_M_nohelp])
    px_nohelp = np.array([px_F_nohelp[1:], px_M_nohelp[1:]])
    px_nohelp[px_nohelp > 1] = 1
    # print(mx_nohelp, px_nohelp)
    L_nomp = leslie_(mx_nohelp, px_nohelp)

    # Menopausal versions - recipients population
    # Females
    px_F_help = np.zeros(px_F.shape[0])
    px_F_help[:female_death_age] = px_F[:female_death_age] #* female_surv_tradeoff
    px_F_help[last_reproduction_age:] = px_F[last_reproduction_age:] * Δmaternal_mortality
    mx_F_help = np.zeros(px_F.shape[0])
    mx_F_help[:female_death_age] = mx_F[:female_death_age] #* female_fec_tradeoff
    # Males
    px_M_help = np.zeros(px_M.shape[0])
    px_M_help[:female_death_age] = px_M[:female_death_age] 
    mx_M_help = np.zeros(px_M.shape[0])
    mx_M_help[:female_death_age] = mx_M[:female_death_age] 
    # Building the matrix
    mx_help = np.array([mx_F_help, mx_M_help])
    px_help = np.array([px_F_help[1:], px_M_help[1:]])
    px_help[px_nohelp > 1] = 1
    # print(mx_nohelp, px_nohelp)
    L_mpR = leslie_(mx_help, px_help)
    
    # # Menopausal versions - donors population
    px_D = np.zeros(px_F.shape[0])
    px_D[:last_reproduction_age] = px_F[:last_reproduction_age] * lx_contribution[0]
    px_D[last_reproduction_age:] = px_F[last_reproduction_age:] * Δmaternal_mortality * lx_contribution[0]
    mx_D = np.zeros(px_F.shape[0])
    mx_D[:last_reproduction_age] = mx_F[:last_reproduction_age] * mx_contribution[0]
    # Building the matrix
    mx_D_tot = np.array([mx_D, mx_M_help])
    px_D_tot = np.array([px_D[1:], px_M_help[1:]])
    px_D_tot[px_D_tot > 1] = 1
    # print(mx_nohelp, px_nohelp)
    L_mpD = leslie_(mx_D_tot, px_D_tot)

    return L_nomp, L_mpR, L_mpD

# %% [code]
def different_longevities(L_mp, r_to_pod, female_fec_tradeoff_arr, female_surv_tradeoff_arr, additional_years_after_rep, Δmaternal_mortality=1, mx_contribution=np.array([0.95, 0.95]), lx_contribution=np.array([0.95, 0.95]), sex_ratio=[0.6,0.4], return_all=False, ε=10**(-50), max_male_help=None, max_female_help=None, age_dist=None, ρ=None, plot_age_dist=False):

    assert additional_years_after_rep == len(female_surv_tradeoff_arr) == len(female_fec_tradeoff_arr)
    L_nomp_per_additional_year = []
    L_mpR_per_additional_year = []
    L_mpD_per_additional_year = []
    
    # d age classes
    d = L_mp.shape[0]//2
    if type(lx_contribution) != np.ndarray: # if the user did not input an array, convert it to one, this format is necessary for the calculations
        lx_contribution = np.array(lx_contribution)
    if type(mx_contribution) != np.ndarray: # if the user did not input an array, convert it to one, this format is necessary for the calculations
        mx_contribution = np.array(mx_contribution)
    
    # Matrices to save the results: C, B, rB
    COST = np.zeros((2, additional_years_after_rep)) 
    BENEFIT = np.zeros((2, additional_years_after_rep))
    rxB = np.zeros((2, additional_years_after_rep))
    if plot_age_dist:
        age_dist_each_onset = np.zeros((additional_years_after_rep, 2, additional_years_after_rep))
        rho_each_onset = np.zeros(additional_years_after_rep)

    if max_male_help is None:
        max_male_help = d
    if max_female_help is None:
        max_female_help = d
    
    for i in range(additional_years_after_rep):
        female_fec_tradeoff, female_surv_tradeoff = female_fec_tradeoff_arr[i], female_surv_tradeoff_arr[i]
        L_nomp, L_mpR, L_mpD = create_non_menopausal_pop(L_mp, lx_contribution, mx_contribution, female_fec_tradeoff, female_surv_tradeoff, additional_years_after_rep, Δmaternal_mortality)
        L_nomp_per_additional_year.append(L_nomp)
        L_mpR_per_additional_year.append(L_mpR)
        L_mpD_per_additional_year.append(L_mpD)
        
        # Calculate the population growth rate of the non-menopausal population since that is the dominant population in the model, as menopause is rare in the beginning. The matrix needs to be in a (d,d) shape for the eigenvalue calculation so I take the average according to the sex ratio. 
        if ρ is None and age_dist is None:    
            ρ = np.linalg.eigvals(sex_ratio[0]*L_nomp[:d,:d]+sex_ratio[1]*L_nomp[d:,d:]).max().real 
            # Calculate the age distribution of the non-menopausal population (L_nomp_weffects)
            age_dist = age_distribution(L_nomp, ρ, sex_ratio=np.array([[sex_ratio[0]], [sex_ratio[1]]])) 
        
        # Calculate the total effect of menopause on the population, using L_mpR as the leslie matrix of the recipient population and L_mpD as the leslie matrix of the donor population. L_nomp_weffects is the leslie matrix of the non-menopausal population, which is the theoretical scenario.
        results = tot_rBC_all_donor_life(L_nomp, L_mpR, L_mpD, ρ, age_dist, r_to_pod, sex_ratio=sex_ratio, return_all=return_all, ε=ε) 
        # results = tot_rBC_all_donor_life_costNoAgeDist(L_nomp_weffects, L_mpR, L_mpD, ρ, age_dist, r_to_pod, sex_ratio=sex_ratio, return_all=return_all, ε=ε) 
        
        if plot_age_dist:
            age_dist_each_onset[i, :, :] = age_dist
            rho_each_onset[onset] = ρ   
        
        # Save the results
        COST[:,i] = results[1]
        BENEFIT[:,i] = results[2]
        rxB[:,i] = results[3]
    
    # Calculate rB-C
    rBC = rxB + COST # Cost is non-positive so it is a subtraction, rB-abs(C) = rB+C
    if return_all and not plot_age_dist:
        return rBC, COST, BENEFIT, rxB
    elif return_all and plot_age_dist:
        return rBC, COST, BENEFIT, rxB, (age_dist_each_onset, rho_each_onset)
    else:
        return rBC 


# %% [code]
def evaluate_survival_fecundity_tradeoffs(
    surv_range,
    fec_range,
    additional_years_after_rep,
    L_mp, 
    r_to_pod_10yr,
    female_fec_tradeoff_arr,
    female_surv_tradeoff_arr,
    Δmaternal_mortality=1,
    mx_contribution=np.array([0.95, 0.75]),
    lx_contribution=np.array([0.8, 0.5]),
    sex_ratio=[0.6, 0.4],
    return_all=True):

    results = []
    for surv in surv_range:
        for fec in fec_range:
            # Tradeoff arrays based on current survival and fecundity values
            female_surv_tradeoff_arr = [1 + (surv - 1) / (i + 1) for i in range(additional_years_after_rep)]
            female_fec_tradeoff_arr = [1 + (fec - 1) / (i + 1) for i in range(additional_years_after_rep)]

            try:
                rBC, C, B, rxB = different_longevities(
                    L_mp,
                    r_to_pod_10yr,
                    female_fec_tradeoff_arr,
                    female_surv_tradeoff_arr,
                    additional_years_after_rep,
                    Δmaternal_mortality=Δmaternal_mortality,
                    mx_contribution=mx_contribution,
                    lx_contribution=lx_contribution,
                    sex_ratio=sex_ratio,
                    return_all=return_all
                )

                results.append({
                    'surv': surv,
                    'fec': fec,
                    'rBC': np.sum(rBC, axis=0),
                    'COST': np.sum(C, axis=0),
                    'BENEFIT': np.sum(B, axis=0),
                    'rxB': np.sum(rxB, axis=0)
                })

            except Exception as e:
                print(f"Error for surv={surv}, fec={fec}: {e}")
                traceback.print_exc()

    return results


# %% [code]
def LM_to_table(L, age_interval):
    d = L.shape[0] // 2
    ages = [i * age_interval for i in range(d)]
    
    female_p = [1] + L[1:d, 0:d].diagonal(-1).tolist()
    male_p = [1] + L[d+1:2*d, d:2*d].diagonal(-1).tolist()
    
    data = {
        "Age": ages,
        "Female p(x)": female_p,
        "Male p(x)": male_p,
        "Female m(x)": L[0, 0:d].tolist(),
        "Male m(x)": L[d, d:2*d].tolist()
    }
    print(data)
    
    return DataFrame(data)

# %% [code]
from scipy.stats import linregress
def decay_rate_of_survival_calc(postpeak_phase_start, survival_array):
    
    ages = np.arange(len(survival_array))
    # Only use the declining portion to estimate decay
    # Let's assume ages 6 and up are in the post-peak phase
    ages_decay = ages[postpeak_phase_start:]
    surv_decay = survival_array[postpeak_phase_start:]
    
    # Filter out any survival = 1 (log(1) = 0 but masks decay)
    valid = surv_decay < 1.0
    ages_decay = ages_decay[valid]
    surv_decay = surv_decay[valid]
    
    # Fit log-linear regression
    log_surv = np.log(surv_decay)
    slope, intercept, r_value, p_value, std_err = linregress(ages_decay, log_surv)
    b_estimated = -slope  # decay rate is the negative slope

    return b_estimated

# %% [code]
def extend_survival_exp_decay(survival_list, num_extra_classes, decay_rate=None, postpeak_phase_start=None):
    if decay_rate is not None and postpeak_phase_start is None:
        last_px = survival_list[-1]
        postrep_px = [last_px * np.exp(-decay_rate * i) for i in range(1, num_extra_classes + 1)]
        return survival_list + postrep_px

    assert type(postpeak_phase_start) == int 
    decay_rate = decay_rate_of_survival_calc(postpeak_phase_start, np.array(survival_list))
    last_px = survival_list[-1]
    postrep_px = [last_px * np.exp(-decay_rate * i) for i in range(1, num_extra_classes + 1)]
    
    return survival_list + postrep_px

# %% [code]
def extend_LM_both_sexes(L_mp, num_extra_classes, decay_rate=None, postpeak_phase_start=None):
    d = L_mp.shape[0] // 2
    survivals_FM = Px(L_mp)
    
    new_survival_F = extend_survival_exp_decay(list(survivals_FM[0]), num_extra_classes, decay_rate=decay_rate, postpeak_phase_start=postpeak_phase_start)
    new_survival_M = np.concatenate([survivals_FM[1], np.zeros(num_extra_classes)])
    new_survival_FM = np.array([new_survival_F[1:], new_survival_M[1:]])

    rep_F, rep_M = mx(L_mp)
    new_rep_F, new_rep_M = np.concatenate([rep_F, np.zeros(num_extra_classes)]), np.concatenate([rep_M, np.zeros(num_extra_classes)])
    new_rep_FM = np.array([new_rep_F, new_rep_M])

    new_LM = leslie_(new_rep_FM, new_survival_FM)

    return new_LM

# %% [code]
def LM_to_block_tables_v1(L, age_interval, to_csv_path_F='F.csv', to_csv_path_M='M.csv'):
    d = L.shape[0] // 2
    ages = [i * age_interval for i in range(d)]

    # Female block
    female_block = np.zeros((d, d))
    female_block[0, :] = L[0, 0:d]  # Fecundity
    for i in range(1, d):
        female_block[i, i-1] = L[i, i-1]  # Survival
    
    # Male block
    male_block = np.zeros((d, d))
    male_block[0, :] = L[d, d:2*d]  # Fecundity
    for i in range(1, d):
        male_block[i, i-1] = L[d+i, d+i-1]  # Survival

    # Convert to DataFrames with age labels
    female_df = DataFrame(female_block, index=ages, columns=ages)
    male_df = DataFrame(male_block, index=ages, columns=ages)

    # female_df.index.name = "Female Age"
    # male_df.index.name = "Male Age"

    female_df.to_csv(to_csv_path_F)
    male_df.to_csv(to_csv_path_M)
    
    return female_df, male_df

# %% [code]
import numpy as np
import pandas as pd
from pandas import DataFrame

def LM_to_block_tables_v2(L, age_interval, to_csv_path_F='F.csv', to_csv_path_M='M.csv'):
    d = L.shape[0] // 2
    ages = [i * age_interval for i in range(d)]

    # Female block
    female_block = np.zeros((d, d))
    female_block[0, :] = L[0, 0:d]  # Fecundity
    for i in range(1, d):
        female_block[i, i-1] = L[i, i-1]  # Survival
    
    # Male block
    male_block = np.zeros((d, d))
    male_block[0, :] = L[d, d:2*d]  # Fecundity
    for i in range(1, d):
        male_block[i, i-1] = L[d+i, d+i-1]  # Survival

    # Convert to DataFrames with age labels
    female_df = DataFrame(female_block, index=ages, columns=ages)
    male_df = DataFrame(male_block, index=ages, columns=ages)

    # Formatting function: 0 → "0", else round to 3 decimals
    fmt = lambda x: "0" if np.isclose(x, 0) else f"{x:.3f}"

    female_df = female_df.applymap(fmt)
    male_df = male_df.applymap(fmt)

    female_df.to_csv(to_csv_path_F, index=True)
    male_df.to_csv(to_csv_path_M, index=True)
    
    return female_df, male_df


# %% [code]
import numpy as np
import matplotlib.pyplot as plt
import matplotlib.patheffects as pe
from matplotlib.path import Path
from matplotlib.patches import PathPatch

# ---------------- helpers ----------------

def _should_plot_year(k, years, skip, highlight_year):
    """
    skip:
      - None      -> draw all
      - int >=2   -> draw every 'skip'-th (k % skip == 0)
      - iterable  -> list/set of YEARS to SKIP (e.g., [10, 20])
    highlight_year is always drawn if provided.
    """
    if highlight_year is not None and years == highlight_year:
        return True
    if skip is None:
        return True
    if isinstance(skip, int):
        return (skip <= 1) or (k % skip == 0)
    try:
        return years not in set(skip)
    except TypeError:
        return True


def _longest_contour_verts(CS):
    """Return vertices (Nx2) of the longest path in this contour result."""
    best = None
    best_L = -1.0
    for coll in CS.collections:
        for p in coll.get_paths():
            v = p.vertices
            if v.shape[0] < 2:
                continue
            seg = np.diff(v, axis=0)
            L = np.sum(np.hypot(seg[:, 0], seg[:, 1]))
            if L > best_L:
                best_L = L
                best = v
    return best


def _arc_midpoint_xy_and_angle(verts):
    """Arc-length midpoint (x,y) and local tangent angle (degrees, upright)."""
    seg = np.diff(verts, axis=0)
    lens = np.hypot(seg[:, 0], seg[:, 1])
    cum = np.concatenate([[0.0], np.cumsum(lens)])
    if cum[-1] == 0:
        x, y = float(verts[0, 0]), float(verts[0, 1])
        return x, y, 0.0
    t = 0.5 * cum[-1]
    i = np.searchsorted(cum, t) - 1
    i = np.clip(i, 0, len(cum) - 2)
    if np.isclose(cum[i + 1], cum[i]):
        w = 0.0
    else:
        w = (t - cum[i]) / (cum[i + 1] - cum[i])
    x = verts[i, 0] + w * (verts[i + 1, 0] - verts[i, 0])
    y = verts[i, 1] + w * (verts[i + 1, 1] - verts[i, 1])
    tan = verts[i + 1] - verts[i]
    ang = np.degrees(np.arctan2(tan[1], tan[0]))
    if ang > 90:
        ang -= 180
    if ang < -90:
        ang += 180
    return float(x), float(y), float(ang)


def _shade_above_below(ax, verts, X, Y, meno_col, non_meno_col, alpha=0.25):
    """
    Use the highlight contour verts to shade:
      - below the line in meno_col
      - above the line in non_meno_col
    over the full x-range.
    """
    x_min, x_max = X.min(), X.max()
    y_min, y_max = Y.min(), Y.max()

    verts = np.asarray(verts)
    xs_line = verts[:, 0]
    ys_line = verts[:, 1]

    # sort by x to make interpolation well-defined
    order = np.argsort(xs_line)
    xs_line = xs_line[order]
    ys_line = ys_line[order]

    # dense x-grid for smooth fill
    xs = np.linspace(x_min, x_max, 500)
    ys = np.interp(xs, xs_line, ys_line,
                   left=ys_line[0], right=ys_line[-1])

    # below = menopause-favoured
    ax.fill_between(xs, y_min, ys, color=meno_col, alpha=alpha, zorder=0)
    # above = non-menopause
    ax.fill_between(xs, ys, y_max, color=non_meno_col, alpha=alpha, zorder=0)


# --------------- main function ---------------

def plot_lambda_zero_contours(results, surv_range, fec_range, additional_years_after_rep,
                              years_per_step=5, fallback_when_no_zero='max',
                              highlight_year=30, shade_below_highlight=True,
                              figsize=(11, 8), legend_title_size=14, legend_label_size=12, fs=20,
                              skip=None, meno_col='lightseagreen', non_meno_col='lightcoral', alpha=0.9):
    """
    Draw contour lines over (fecundity x survival):
      - Black line for λ = 0 when it exists.
      - Optional fallback: gray contour at max(λ) when no zero crossing.
      - Bold the line at `highlight_year` and (optionally) shade:
          below it (λ>0) in `meno_col`
          above it in `non_meno_col`
      - `skip` lets you skip some contours (see _should_plot_year docstring).
    """
    ns, nf, ny = len(surv_range), len(fec_range), additional_years_after_rep

    # Rebuild Z_stack[year, i_surv, j_fec] from results
    Z_stack = np.empty((ny, ns, nf), dtype=float)
    idx = 0
    for i in range(ns):
        for j in range(nf):
            Z_stack[:, i, j] = results[idx]['rBC'][:ny]
            idx += 1

    X, Y = np.meshgrid(fec_range, surv_range)  # X=fec, Y=surv
    fig, ax = plt.subplots(figsize=figsize)

    # Capture vertices for the highlight contour (to shade later)
    highlight_verts = None
    highlight_idx = int(round(highlight_year / years_per_step)) - 1  # 0-based

    # --- pre-pass: detect highlight contour & clear for shading order ---
    for k in range(ny):
        Zk = Z_stack[k]
        years = (k + 1) * years_per_step
        ls, lw, color, zorder = '-', 1.6, 'black', 3
        has_zero = (np.nanmin(Zk) <= 0.0) and (np.nanmax(Zk) >= 0.0)
        plot_this = _should_plot_year(k, years, skip, highlight_year)

        if has_zero:
            if not (plot_this or k == highlight_idx):
                continue
            CS = ax.contour(X, Y, Zk, levels=[0], colors=color,
                            linewidths=lw, linestyles=ls, zorder=zorder)
            if k == highlight_idx and CS.collections and CS.collections[0].get_paths():
                ax.cla()  # clear so shading can be drawn beneath contours
                break
        elif fallback_when_no_zero == 'max':
            if not plot_this:
                continue
            zmax = float(np.nanmax(Zk))
            level = zmax if np.nanmin(Zk) < zmax else zmax + 1e-12
            ax.contour(X, Y, Zk, levels=[level], colors='gray',
                       linewidths=1.4, linestyles=ls, zorder=2)

    # --- redraw properly, with skipping & labels ---
    proxies, labels = [], []
    for k in range(ny):
        Zk = Z_stack[k]
        years = (k + 1) * years_per_step
        ls = '-'
        has_zero = (np.nanmin(Zk) <= 0.0) and (np.nanmax(Zk) >= 0.0)
        plot_this = _should_plot_year(k, years, skip, highlight_year)

        if has_zero:
            if not (plot_this or k == highlight_idx):
                continue
            lw = 1.6 if k != highlight_idx else 2.6
            color = 'black'
            CS = ax.contour(X, Y, Zk, levels=[0], colors=color,
                            linewidths=lw, linestyles=ls, zorder=3)

            # label at arc-length midpoint of the longest segment
            try:
                verts = _longest_contour_verts(CS)
                if verts is not None:
                    vx, vy, vang = _arc_midpoint_xy_and_angle(verts)
                    ax.text(vx, vy, f"{years}y", fontsize=fs, color=color,
                            ha='center', va='center',
                            rotation=vang, rotation_mode='anchor',
                            path_effects=[pe.withStroke(linewidth=3, foreground='white')],
                            zorder=4)
                    if k == highlight_idx:
                        highlight_verts = verts
            except Exception:
                pass

            proxies.append(plt.Line2D([0], [0], color=color, lw=lw, linestyle=ls))
            labels.append(f"{years}y (λ=0)")

        elif fallback_when_no_zero == 'max':
            if not plot_this:
                continue
            zmax = float(np.nanmax(Zk))
            level = zmax if np.nanmin(Zk) < zmax else zmax + 1e-12
            CS = ax.contour(X, Y, Zk, levels=[level], colors='gray',
                            linewidths=1.4, linestyles=ls, zorder=2)

            try:
                verts = _longest_contour_verts(CS)
                if verts is not None:
                    vx, vy, vang = _arc_midpoint_xy_and_angle(verts)
                    ax.text(vx, vy, f"{years}y", fontsize=fs, color='gray',
                            ha='center', va='center',
                            rotation=vang, rotation_mode='anchor',
                            path_effects=[pe.withStroke(linewidth=3, foreground='white')],
                            zorder=4)
            except Exception:
                pass

            proxies.append(plt.Line2D([0], [0], color='gray', lw=1.4, linestyle=ls))
            labels.append(f"{years}y (max λ={zmax:.3g})")

    # Shade regions relative to the highlight contour
    if shade_below_highlight and highlight_verts is not None:
        _shade_above_below(ax, highlight_verts, X, Y,
                           meno_col=meno_col, non_meno_col=non_meno_col, alpha=alpha)

    # Axes labels/ticks
    ax.set_xlabel('Fecundity Trade-off', fontsize=24)
    ax.set_ylabel('Survival Trade-off', fontsize=24)
    ax.tick_params(axis='both', which='major', labelsize=21)
    ax.grid(False)

    plt.tight_layout()
    return fig, ax


# %% [code]
# ---- ADD THIS BLOCK TO EACH SPECIES SCRIPT ----
import os

def save_sobol_results(species_name, Si, param_names, outdir="sa_results"):
    """
    Save per-parameter Sobol indices for one species to CSV.
    Columns: species, param, S1, S1_conf, ST, ST_conf
    """
    os.makedirs(outdir, exist_ok=True)
    df = pd.DataFrame({
        "species": species_name,
        "param": param_names,
        "S1": Si["S1"],
        "S1_conf": Si["S1_conf"],
        "ST": Si["ST"],
        "ST_conf": Si["ST_conf"],
    })
    df.to_csv(os.path.join(outdir, f"{species_name}_sobol.csv"), index=False)
    print(f"Saved: {os.path.join(outdir, f'{species_name}_sobol.csv')}")


# %% [code]
def calc_rcoeff(L_mp, m, df, dm, nf, nm, a):

    d = L_mp.shape[0] // 2
    mu = np.zeros((2,d)) # ar ray for survival probabilities per age-group, where each age group is in 10-year intervals
    
    for i in range(d-1):
        mu[:,i] = L_mp[i+1,i], L_mp[a+i+1,a+i]
        
    rf_to_f, rm_to_f, rf_to_m, rm_to_m, avg_rf = rff_age_v2(df, dm, nf, nm, m, a, 1-mu)
    r_to_pod = np.array([rf_to_f, rm_to_f])

    # print("mu shape:", mu.shape, ", a:", a, ", d:", d)
    return r_to_pod

# %% [code]
def fig_panel_labels(axes, letters=None, uppercase=True,
                     xcoord=-0.17, ycoord=0.92, panel_label_size=None):
    import string
    from matplotlib.font_manager import FontProperties
    import numpy as np

    # Convert rcParam to a numeric size
    if panel_label_size is None:
        fp = FontProperties(size=plt.rcParams['axes.titlesize'])
        panel_label_size = fp.get_size() * 1.3

    # Generate default letters if needed
    if letters is None:
        letters = string.ascii_uppercase if uppercase else string.ascii_lowercase

    axes_arr = np.array(axes, dtype=object)
    flat_axes = list(axes_arr.flat)          # make a list so we can iterate safely
    n_axes = len(flat_axes)

    # Truncate / slice letters to the number of axes
    letters = list(letters[:n_axes])

    # --- handle xcoord ---
    # scalar: same for all panels
    # sequence length == ncols: per-column xcoord
    # sequence length == n_axes: per-panel xcoord
    if hasattr(xcoord, "__len__") and not isinstance(xcoord, str):
        xcoord_list = list(xcoord)
        if axes_arr.ndim == 2 and len(xcoord_list) == axes_arr.shape[1]:
            # per-column values
            nrows, ncols = axes_arr.shape
            xs = []
            for i in range(n_axes):
                col = i % ncols
                xs.append(xcoord_list[col])
        elif len(xcoord_list) == n_axes:
            xs = xcoord_list
        else:
            raise ValueError("xcoord must be a scalar, length ncols, or length n_axes")
    else:
        xs = [xcoord] * n_axes

    ys = [ycoord] * n_axes  # keep ycoord simple (one value for all)

    # --- annotate ---
    annots = []
    for ax, letter, xc, yc in zip(flat_axes, letters, xs, ys):
        annots.append(
            ax.annotate(
                letter, (xc, yc),
                xycoords='axes fraction',
                fontsize=panel_label_size
            )
        )
    return annots


# %% [code]
def save_results_csv_LL(res, additional_years_after_rep, path='lambda_results.csv'):
    rows = []
    K = additional_years_after_rep
    for r in res:
        row = {'surv': r['surv'], 'fec': r['fec']}
        for key in ('rBC', 'COST', 'BENEFIT', 'rxB'):
            arr = np.asarray(r[key]).ravel()
            for k in range(K):
                row[f'{key}_{k+1}'] = float(arr[k])
        rows.append(row)
    pd.DataFrame(rows).to_csv(path, index=False)

# --- LOAD (rebuild list-of-dicts like your original `res`) ---
def load_results_csv_LL(path='lambda_results.csv', additional_years_after_rep=None):
    df = pd.read_csv(path)
    if additional_years_after_rep is None:
        # infer K from rBC_* columns
        K = sum(c.startswith('rBC_') for c in df.columns)
    else:
        K = additional_years_after_rep

    out = []
    for _, row in df.iterrows():
        rec = {'surv': float(row['surv']), 'fec': float(row['fec'])}
        for key in ('rBC', 'COST', 'BENEFIT', 'rxB'):
            rec[key] = np.array([row[f'{key}_{i+1}'] for i in range(K)], dtype=float)
        out.append(rec)
    return out


# %% [code]
```
