## Supplementary Code for "Evolution of Menopause and the Mama’s Boy Hypothesis": RKW_stop_early.ipynb

```
# %% [markdown]
#  Kin selection in age-structured populations - RKW

# %% [markdown]
## Imports

# %% [code]
import numpy as np
from pandas import DataFrame, read_csv
import matplotlib.pyplot as plt
import seaborn as sns
from copy import deepcopy
from scipy.linalg import leslie
%matplotlib inline
import matplotlib.image as mpimg
import pandas as pd

# %% [code]
import io, os, sys, types

# %% [code]
from IPython import get_ipython
from nbformat import read
from IPython.core.interactiveshell import InteractiveShell

# %% [code]
def find_notebook(fullname, path=None):
    """find a notebook, given its fully qualified name and an optional path

    This turns "foo.bar" into "foo/bar.ipynb"
    and tries turning "Foo_Bar" into "Foo Bar" if Foo_Bar
    does not exist.
    """
    name = fullname.rsplit('.', 1)[-1]
    if not path:
        path = ['']
    for d in path:
        nb_path = os.path.join(d, name + ".ipynb")
        if os.path.isfile(nb_path):
            return nb_path
        # let import Notebook_Name find "Notebook Name.ipynb"
        nb_path = nb_path.replace("_", " ")
        if os.path.isfile(nb_path):
            return nb_path

# %% [code]
class NotebookLoader(object):
    """Module Loader for Jupyter Notebooks"""
    def __init__(self, path=None):
        self.shell = InteractiveShell.instance()
        self.path = path

    def load_module(self, fullname):
        """import a notebook as a module"""
        path = find_notebook(fullname, self.path)

        print ("importing Jupyter notebook from %s" % path)

        # load the notebook object
        with io.open(path, 'r', encoding='utf-8') as f:
            nb = read(f, 4)


        # create the module and add it to sys.modules
        # if name in sys.modules:
        #    return sys.modules[name]
        mod = types.ModuleType(fullname)
        mod.__file__ = path
        mod.__loader__ = self
        mod.__dict__['get_ipython'] = get_ipython
        sys.modules[fullname] = mod

        # extra work to ensure that magics that would affect the user_ns
        # actually affect the notebook module's ns
        save_user_ns = self.shell.user_ns
        self.shell.user_ns = mod.__dict__

        try:
          for cell in nb.cells:
            if cell.cell_type == 'code':
                # transform the input to executable Python
                code = self.shell.input_transformer_manager.transform_cell(cell.source)
                # run the code in themodule
                exec(code, mod.__dict__)
        finally:
            self.shell.user_ns = save_user_ns
        return mod

# %% [code]
class NotebookFinder(object):
    """Module finder that locates Jupyter Notebooks"""
    def __init__(self):
        self.loaders = {}

    def find_module(self, fullname, path=None):
        nb_path = find_notebook(fullname, path)
        if not nb_path:
            return

        key = path
        if path:
            # lists aren't hashable
            key = os.path.sep.join(path)

        if key not in self.loaders:
            self.loaders[key] = NotebookLoader(path)
        return self.loaders[key]

# %% [code]
sys.meta_path.append(NotebookFinder())

# %% [code]
from Johnstone_Cant import IBD_age_0, no_age_rs, IBD_no_age, IBD_older_age_v2, rff_age_v2

# %% [code]
from project_functions import leslie_, Px, mx, age_distribution, tot_rBC_all_donor_life, tot_rBC, survival_effect, fecundity_effect, reproductive_value, different_onset_ages, latex_array, add_image, LM_to_table,LM_to_block_tables_v2, save_sobol_results 

# %% [markdown]
## Menopausal (current state) population parameters

# %% [markdown]
#### Leslie Matrix (LM) - total pop
only used for ρ (leading eigenvalue) calculation.

# %% [code]
d = 10 # looking at 10 age classes: 0-10,10-20,20-30,30-40,40-50,50-60,60-70,70-80,80-90,90-100
fecundity = np.zeros(d)
fecundity[1:4] = [1, 2, 2]
survival = np.array([3903/5000, 389/400, 2491/2500, 1971/2000, 9641/10000, 4711/5000, 4491/5000, 431/500, 431/500])
Ltot = leslie(fecundity, survival)

# %% [code]
# maximal eigenvalue = growth rate of population
λ0 = np.linalg.eigvals(Ltot).max()
assert λ0.imag == 0
ρ = λ0.real
ρ

# %% [markdown]
#### LM - by males and females

# %% [code]
d = 10
L_mp = np.zeros((2*d, 2*d))
fecundity = np.zeros(d)
fecundity[1:4] = [1, 2, 2]
fecundity_both_sex = np.array([fecundity, fecundity])
survival_f = np.array([3903/5000, 389/400, 2491/2500, 1971/2000, 9641/10000, 4711/5000, 4491/5000, 431/500, 431/500])
survival_m = np.array([3903/5000, 0.9315, 0.9149, 0.8588, 0.8588, 0.8588, 0, 0, 0])
survival_both_sex = np.array([survival_f, survival_m])
L_mp = leslie_(fecundity_both_sex, survival_both_sex)

# %% [code]
dis = DataFrame({'sex':np.concatenate([np.array(['F']*10), np.array(['M']*10)]), 'age':np.concatenate([np.arange(0,100,10)]*2), 'frequency':age_distribution(L_mp, ρ).flatten()})
pal = sns.color_palette('mako')[-2::-4]
barp = sns.barplot(data=dis, x='age', y='frequency', hue='sex', palette=pal, width=0.9)
distr = barp.get_figure()
distr.suptitle('Age distribution in killer whales')
distr.savefig('figA1.png')
np.sum(age_distribution(L_mp, ρ)) # sanity check

# %% [markdown]
## Non-menopausal population

# %% [markdown]
#### LM - no menopause - by males and females

# %% [code]
L_no_effects = deepcopy(L_mp)
L_no_effects[0,4:10] = np.array([2,1,1,0.2,0.1,0]) #based on elephants
competition_factor = 1/1.67
competition_start_age = 4
L_no_effects[0, competition_start_age:10] *= competition_factor
L_no_effects[0, competition_start_age:10]

# %% [markdown]
## Demographic parameters

# %% [code]
def calc_rcoeff(L_mp, m, df, dm, nf, nm, a):

    d = L_mp.shape[0] // 2
    mu = np.zeros((2,d)) # array for survival probabilities per age-group, where each age group is in 10-year intervals
    
    for i in range(d-1):
        mu[:,i] = L_mp[i+1,i], L_mp[10+i+1,10+i]
        
    rf_to_f, rm_to_f, rf_to_m, rm_to_m, avg_rf = rff_age_v2(df, dm, nf, nm, m, a, 1-mu)
    r_to_pod = np.array([rf_to_f, rm_to_f])

    return r_to_pod

# %% [code]
male_symbol = mpimg.imread('/Users/Omer/Downloads/pTodqKkpc-3982000212.png')
female_symbol = mpimg.imread('/Users/Omer/Downloads/8czr7L9Ki-3456542989.png')
RKW_phylopic = mpimg.imread('/Users/Omer/Downloads/1024x503.png')

# %% [code]
df = dm = 0.1 # male, female dispersal occurs at a probability 0.1
m = 0 # indicative for non-local reproduction as in orcas
nf = nm = 3
a = 10 # number of age classes to view
mu = np.zeros((2,10)) # array for survival probabilities per age-group, where each age group is in 10-year intervals
for i in range(9):
    mu[:,i] = L_mp[i+1,i], L_mp[10+i+1,10+i]
rf_to_f, rm_to_f, rf_to_m, rm_to_m, avg_rf = rff_age_v2(df, dm, nf, nm, m, a, 1-mu)
r_to_pod = np.array([rf_to_f, rm_to_f])

r_to_pod_10yr = np.concatenate([r_to_pod[:,::5], (r_to_pod[:, -1]).reshape(2,1)], axis=1)
age_classes = np.arange(0,100,10)

lw, fs = 2.5, 17

fig, ax = plt.subplots(figsize=(8,5))
ax.plot(age_classes, r_to_pod_10yr[0,:], label='$r_f$', c='salmon', lw=lw, marker='o')
ax.plot(age_classes, r_to_pod_10yr[1,:], label='$r_m$', c='cornflowerblue', lw=lw, marker='o')
# ax.axvline(40,c='gray', ls=':', label='$r$ at ageD=40')
ax.plot(age_classes, 0.6*r_to_pod_10yr[0,:]+0.4*r_to_pod_10yr[1,:], c='#FFC98B', label='$r_{avg}$', lw=lw, ls='--')# marker='o')
ax.set_xlabel('Female age', fontsize=fs+2)
ax.set_ylabel('Relatedness coefficient', fontsize=fs+2)
ax.tick_params(axis='both', which='major', labelsize=fs)
ax.set_xticks(np.arange(0,100,10))
add_image(ax, male_symbol, xy=(6, 0.075), zoom=0.014)
add_image(ax, female_symbol, xy=(0.5, 0.205), zoom=0.011)
# ax.set_title('Relatedness Coefficient of Female to Pod Throughout Life (10-year intervals)', fontsize=fs)
# ax.legend(fontsize=fs-3)
add_image(ax, RKW_phylopic, xy=(82, 0.08), zoom=0.1)
plt.savefig('RKW_r.png')

plt.show()

# %% [markdown]
## Running the Model

# %% [code]
age_classes = np.arange(0,100,10)
age_classes = age_classes[3:]
r_to_pod_10yr_original = r_to_pod_10yr
r_to_pod_10yr = r_to_pod_10yr[:,3:]

# %% [code]
ε = 10**(-60) 
rbc, c, b, rb = different_onset_ages(L_no_effects, r_to_pod_10yr_original, Δmaternal_mortality=0.97, mx_contribution=[0.95, 0.75], lx_contribution=[0.8, 0.5], return_all=True, ε=ε)#, age_dist=age_distribution(L_mp, ρ), ρ=ρ) 
rbc, c, b, rb = rbc[:,3:], c[:,3:], b[:,3:], rb[:,3:]

fig, ax = plt.subplots(figsize=(8,5))
ax.axhline(0,c='k')
ax.plot(age_classes, np.sum(rbc, axis=0), label='λ', marker='o', c='#E9765B', lw=lw, markersize=7)
ax.plot(age_classes, np.sum(c, axis=0), label='C', ls='-', c='#FFC98B', lw=lw)
ax.plot(age_classes, np.sum(b, axis=0), label='B', ls='-', c='#A5CAD2', lw=lw)
ax.plot(age_classes, np.sum(rb, axis=0), label='r*B', ls='-', c='#478BA2', lw=lw)
add_image(ax, RKW_phylopic, xy=(85, -0.047), zoom=0.1)
ax.axvline(10*(np.where(np.sum(rbc, axis=0)>ε)[0][0]+3), c='k', ls=':', label='first positive')
ax.set_xlabel('Menopause age', fontsize=fs+2)
ax.set_ylabel('Benefit and cost over lifespan', fontsize=fs+2)
# ax.set_ylabel('λ', fontsize=fs+2)
ax.tick_params(axis='both', which='major', labelsize=fs)
# ax.set_xlim(0,91)
ax.set_xticks(np.arange(30,100,10))
# ax.set_yscale('log')
plt.savefig('lambda_rkw.png')
# ax.set_title('\n$d_f = d_m = 0$;  $m=1$;  $n_f = n_m = 30$', fontsize=fs+4)
# ax.set_title('\n$d_f = d_m = 0.1$;  $m=0$;  $n_f = n_m = 3$', fontsize=fs+4)
# sns.despine()
# ax.legend(fontsize=fs-1)
# ax.set_title('\n$λ$, C, B and r*B values as a function of different menopause\nonset ages for female donor', fontsize=fs)
plt.show()

np.sum(rbc, axis=0), np.sum(c, axis=0), np.sum(b, axis=0), np.sum(rb, axis=0)


# %% [code]
ε = 10**(-60) 
rbc1, c1, b1, rb1 = different_onset_ages(L_no_effects, r_to_pod_10yr_original, Δmaternal_mortality=1, mx_contribution=[1, 1], lx_contribution=[1, 1], return_all=True, ε=ε)#, age_dist=age_distribution(L_mp, ρ), ρ=ρ) 
np.sum(rbc1, axis=0), np.sum(c1, axis=0), np.sum(b1, axis=0), np.sum(rb1, axis=0)

# %% [markdown]
## 3D graph: $\lambda$($d_f$, $d_m$)

# %% [code]
def compute_rBC(df, dm, m, nf, nm, a, mu, L_no_effects, L_mp, Δmaternal_mortality, mx_contribution, lx_contribution, sex_ratio, ε, onset):
    r_to_pod = calc_rcoeff(L_mp, m, df, dm, nf, nm, a)
    r_to_pod_10yr = np.concatenate([r_to_pod[:, ::5], r_to_pod[:, -1:].reshape(2, 1)], axis=1)
    rBC_vals, *_ = different_onset_ages(
        L_no_effects, r_to_pod_10yr, Δmaternal_mortality=Δmaternal_mortality,
        mx_contribution=mx_contribution, lx_contribution=lx_contribution,
        sex_ratio=sex_ratio, return_all=True, ε=ε
    )
    return np.sum(rBC_vals[:, onset], axis=0)

# %% [code]
def rBC_by_demography_3D_vals(L_no_effects, L_mp, default_r_to_pod, nf, nm, a, mu, 
                          default_df=0.1, default_dm=0.1, default_m=0, 
                          Δmaternal_mortality=1, mx_contribution=np.array([0.95, 0.95]), 
                          lx_contribution=np.array([0.95, 0.95]), sex_ratio=[0.6, 0.4], 
                          ε=1e-5, onset=5):

    df_range = dm_range = m_range = np.arange(0, 1.01, 0.01)
    size = df_range.shape[0]

    rBC_by_df_dm = np.empty((size, size))
    rBC_by_df_m = np.empty((size, size))
    rBC_by_dm_m = np.empty((size, size))

    for i, df in enumerate(df_range):
        for j, dm in enumerate(dm_range):
            rBC_by_df_dm[i, j] = compute_rBC(df, dm, default_m, nf, nm, a, mu,
                                             L_no_effects, L_mp, Δmaternal_mortality, 
                                             mx_contribution, lx_contribution, 
                                             sex_ratio, ε, onset)

    for i, df in enumerate(df_range):
        for j, m in enumerate(m_range):
            rBC_by_df_m[i, j] = compute_rBC(df, default_dm, m, nf, nm, a, mu,
                                            L_no_effects, L_mp, Δmaternal_mortality, 
                                            mx_contribution, lx_contribution, 
                                            sex_ratio, ε, onset)

    for i, dm in enumerate(dm_range):
        for j, m in enumerate(m_range):
            rBC_by_dm_m[i, j] = compute_rBC(default_df, dm, m, nf, nm, a, mu,
                                            L_no_effects, L_mp, Δmaternal_mortality, 
                                            mx_contribution, lx_contribution, 
                                            sex_ratio, ε, onset)

    return rBC_by_df_dm, rBC_by_df_m, rBC_by_dm_m

# %% [code]
rBC_by_demos = rBC_by_demography_3D_vals(L_no_effects, L_mp, r_to_pod_10yr_original, nf, nm, a, mu, default_df=0.1, default_dm=0.1, default_m=0, Δmaternal_mortality=0.97, mx_contribution=[0.95, 0.75], lx_contribution=[0.8, 0.5], sex_ratio=[0.6,0.4], ε=ε)

# %% [code]
def rBC_to_df_dm_3D_vals(L_no_effects, L_mp, default_r_to_pod, nf, nm, a, mu, default_df=0.1, default_dm=0.1, default_m=0, Δmaternal_mortality=1, mx_contribution=np.array([0.95, 0.95]), lx_contribution=np.array([0.95, 0.95]), sex_ratio=[0.6,0.4], ε=10**(-50), onset=5):

    rBC, C, B, rB = different_onset_ages(L_no_effects, default_r_to_pod, Δmaternal_mortality=Δmaternal_mortality, mx_contribution=mx_contribution, lx_contribution=lx_contribution, sex_ratio=sex_ratio, return_all=True, ε=ε) 
    
    df_range = dm_range = m_range = np.arange(0, 1.01, 0.01)
    
    r_by_df_dm = np.empty((df_range.shape[0], df_range.shape[0]))
    r_by_df_m = np.empty((df_range.shape[0], m_range.shape[0]))
    r_by_dm_m = np.empty((dm_range.shape[0], m_range.shape[0]))
    
    rBC_by_df_dm = np.empty((df_range.shape[0], df_range.shape[0]))
    rBC_by_df_m = np.empty((df_range.shape[0], m_range.shape[0]))
    rBC_by_dm_m = np.empty((dm_range.shape[0], m_range.shape[0]))
    
    # print(rBC_by_df_dm.shape, r_by_df_dm.shape)
    for idx_df in range(df_range.shape[0]):
        for idx_dm in range(dm_range.shape[0]):
            df = df_range[idx_df]
            dm = dm_range[idx_dm]
        
            rf_to_f, rm_to_f, rf_to_m, rm_to_m, avg_rf = rff_age_v2(df, dm, nf, nm, default_m, a, 1-mu) 
            r_to_pod = np.array([rf_to_f, rm_to_f])
            r_to_pod_10yr = np.concatenate([r_to_pod[:,::5], (r_to_pod[:, -1]).reshape(2,1)], axis=1)
            rBC_vals, c, b, rb = different_onset_ages(L_no_effects, r_to_pod_10yr, Δmaternal_mortality=Δmaternal_mortality, mx_contribution=mx_contribution, lx_contribution=lx_contribution, sex_ratio=sex_ratio, return_all=True, ε=ε)         
            rBC_on_onset = np.sum(rBC_vals[:, onset], axis=0) 
            # print(rBC_on_onset)
            rBC_by_df_dm[idx_df, idx_dm] = rBC_on_onset

    for idx_df in range(df_range.shape[0]):
        for idx_m in range(m_range.shape[0]):
            df = df_range[idx_df]
            m = m_range[idx_m]
        
            rf_to_f, rm_to_f, rf_to_m, rm_to_m, avg_rf = rff_age_v2(df, default_dm, nf, nm, m, a, 1-mu) 
            r_to_pod = np.array([rf_to_f, rm_to_f])
            r_to_pod_10yr = np.concatenate([r_to_pod[:,::5], (r_to_pod[:, -1]).reshape(2,1)], axis=1)
            rBC_vals, c, b, rb = different_onset_ages(L_no_effects, r_to_pod_10yr, Δmaternal_mortality=Δmaternal_mortality, mx_contribution=mx_contribution, lx_contribution=lx_contribution, sex_ratio=sex_ratio, return_all=True, ε=ε)         
            rBC_on_onset = np.sum(rBC_vals[:, onset], axis=0) 
            # print(rBC_on_onset)
            rBC_by_df_m[idx_df, idx_m] = rBC_on_onset

    for idx_dm in range(dm_range.shape[0]):
        for idx_m in range(m_range.shape[0]):
            dm = dm_range[idx_dm]
            m = m_range[idx_m]
        
            rf_to_f, rm_to_f, rf_to_m, rm_to_m, avg_rf = rff_age_v2(default_df, dm, nf, nm, m, a, 1-mu) 
            r_to_pod = np.array([rf_to_f, rm_to_f])
            r_to_pod_10yr = np.concatenate([r_to_pod[:,::5], (r_to_pod[:, -1]).reshape(2,1)], axis=1)
            rBC_vals, c, b, rb = different_onset_ages(L_no_effects, r_to_pod_10yr, Δmaternal_mortality=Δmaternal_mortality, mx_contribution=mx_contribution, lx_contribution=lx_contribution, sex_ratio=sex_ratio, return_all=True, ε=ε)         
            rBC_on_onset = np.sum(rBC_vals[:, onset], axis=0) 
            # print(rBC_on_onset)
            rBC_by_dm_m[idx_dm, idx_m] = rBC_on_onset

    return rBC_by_df_dm, rBC_by_df_m, rBC_by_dm_m

# %% [code]
rBC_by_demos = rBC_to_df_dm_3D_vals(L_no_effects, L_mp, r_to_pod_10yr_original, nf, nm, a, mu, default_df=0.1, default_dm=0.1, default_m=0, Δmaternal_mortality=0.97, mx_contribution=[0.95, 0.75], lx_contribution=[0.8, 0.5], sex_ratio=[0.6,0.4], ε=ε)

# %% [markdown]
## 3D plot $\lambda$($d_f$, $d_m$) (NOT color gradient)

# %% [code]
def rBC_to_df_dm_3D(rBC_by_df_dm, rBC, default_df, default_dm, ε=10**(-5)):
    
    onset = np.where(np.sum(rBC, axis=0)>ε)[0][0] 
    df_range = dm_range = np.arange(0, 1.01, 0.01)
    
    params = np.array([df_range, dm_range])
    defaults = np.array([default_df, default_dm])
    y_label = '\n\n\n'+'$λ(d_f, d_m)$'

    fig = plt.figure(figsize=(20, 14))#figsize=(8, 6))
    ax = fig.add_subplot(1, 1, 1, projection='3d')
    dot = np.sum(rBC[:, onset], axis=0) 
    
    # Create meshgrid from df_range and dm_range
    df_mesh, dm_mesh = np.meshgrid(df_range, dm_range)
    ax.plot(default_df, default_dm, dot, marker='o', markersize=12, color='royalblue', zorder=10)
    
    # Plot 3D surface
    surf = ax.plot_surface(df_mesh, dm_mesh, rBC_by_df_dm, cmap='magma', edgecolor='none')
    # Add a color bar which maps values to colors
    fig.colorbar(surf, ax=ax, shrink=0.35, aspect=7)
    surf = ax.plot_surface(df_mesh, dm_mesh, np.zeros((101, 101)), cmap='copper', edgecolor='none', alpha=0.27)

    ax.set_xlim([0, 1])  # Replace xmin and xmax with your desired limits for the x-axis
    ax.set_ylim([0, 1])  # Replace ymin and ymax with your desired limits for the y-axis

    ax.set_zlabel(y_label, fontsize=17)
    ax.set_xlabel('$d_f$', fontsize=17)
    ax.set_ylabel('$d_m$', fontsize=17)
    ax.tick_params(axis='x', labelsize=14, pad=5)  # Adjust x-axis tick label size and padding
    ax.tick_params(axis='y', labelsize=14, pad=5)  # Adjust y-axis tick label size and padding
    ax.tick_params(axis='z', labelsize=14, pad=10)  # Adjust z-axis tick label size and padding

    plt.show()
  

# %% [code]
rBC_to_df_dm_3D(rBC_by_df_dm, rbc, 0.1, 0.1, ε=ε)

# %% [markdown]
## 3D plot $\lambda$($d_f$, $d_m$) (color gradient)

# %% [code]
def fig_panel_labels(axes, letters=None, uppercase=True, xcoord=-0.17, ycoord=0.92, panel_label_size=None):
    import string
    from matplotlib.font_manager import FontProperties

    # Convert rcParam to a numeric size
    if panel_label_size is None:
        fp = FontProperties(size=plt.rcParams['axes.titlesize'])
        panel_label_size = fp.get_size() * 1.3

    if letters is None:
        letters = string.ascii_uppercase if uppercase else string.ascii_lowercase

    return [
        ax.annotate(
            letter, (xcoord, ycoord),
            xycoords='axes fraction',
            fontsize=panel_label_size
        )
        for ax, letter in zip(axes.flat, letters)
    ]


# %% [code]
def rBC_by_demographs_pclrmesh(rBC_by_demos, rBC, default_df, default_dm, default_m, onset, ε=10**(-50), letters=None, uppercase=True, xcoord=-0.17, ycoord=0.92, panel_label_size=None):

    df_range = dm_range = m_range = np.arange(0, 1.01, 0.01)
    dot = np.sum(rBC[:, onset], axis=0) 
    rBC_by_df_dm, rBC_by_df_m, rBC_by_dm_m = rBC_by_demos
    vmaxes = np.abs(rBC_by_df_dm).max(), np.abs(rBC_by_df_m).max(), np.abs(rBC_by_dm_m).max()

    fig, axes = plt.subplots(1, 3, figsize=(25, 6))
    fig.subplots_adjust(wspace=0.3)
    y_ranges = [df_range, df_range, dm_range]
    x_ranges = [dm_range, m_range, m_range]
    y_labels = [r'Female dispersal ($d_f$)', r'Female dispersal ($d_f$)', r'Male dispersal ($d_m$)']
    x_labels = [r'Male dispersal ($d_m$)', r'Local mating ($m$)', r'Local mating ($m$)']
    stars = [(default_df, default_dm), (default_df, default_m), (default_dm, default_m)]

    for ax, data, x, y, xlabel, ylabel, vmax, (y_star, x_star) in zip(
            axes, rBC_by_demos, x_ranges, y_ranges, x_labels, y_labels, vmaxes, stars):
        
        pcm = ax.pcolormesh(x, y, data, cmap='RdBu', shading='auto',
                            vmin=-vmax, vmax=vmax)
        ax.contour(x, y, data, levels=[0], colors='black')#, linestyles='dashed')
        right_edge_data = data[:, -1]
        idx = np.argmin(np.abs(right_edge_data))
        anchor_df = y[idx] # This is the df value where lambda=0 at dm=1.0
        
        # 2. Create a symmetric line (slope -1) passing through (1.0, anchor_df)
        # Equation: y - y1 = m(x - x1)  =>  df - anchor_df = -1 * (dm - 1.0)
        # Simplified: df = anchor_df + 1.0 - dm
        ref_dm = np.linspace(0, 1.0, 100)
        ref_df = anchor_df + 1.0 - ref_dm
        
        # 3. Mask the line so it stays within the [0, 1] plot boundaries
        mask = (ref_df >= 0) & (ref_df <= 1)
        ax.plot(ref_dm[mask], ref_df[mask], color='gray', linestyle='dotted', 
                linewidth=2.5, label='Symmetric expectation')
        
        # ax.legend(loc='lower left', fontsize=12)
        ax.plot(x_star, y_star, marker='*', color='k', markersize=14)
        ax.set_xlabel(xlabel, fontsize=18)
        ax.set_ylabel(ylabel, fontsize=18)
        ax.tick_params(labelsize=16)

        cbar = fig.colorbar(pcm, ax=ax, location='right')
        cbar.set_label(r'$\lambda$', fontsize=16)
        cbar.ax.tick_params(labelsize=14)

    fig_panel_labels(axes, letters=letters, uppercase=uppercase, xcoord=xcoord, ycoord=ycoord, panel_label_size=panel_label_size)
    plt.show()

# %% [code]
rBC_by_demographs_pclrmesh(rBC_by_demos, rbc, 0.1, 0.1, 0, onset=5, ε=ε, ycoord=1.04, panel_label_size=22)

# %% [code]
def rBC_by_df_dm_subplot(rBC_by_df_dm, rBC, default_df, default_dm, onset, ax, ε=10**(-5)):

    df_range = dm_range = np.arange(0, 1.01, 0.01)
    dot = np.sum(rBC[:, onset], axis=0) 

    pcm = ax.pcolormesh(df_range, dm_range, rBC_by_df_dm, cmap='RdBu', vmin=-np.max(np.abs(rBC_by_df_dm)), vmax=np.max(np.abs(rBC_by_df_dm)))
    # Add a colorbar using the mappable variable
    cbar = fig.colorbar(pcm, ax=ax)#, shrink=0.8)
    # Set the label for the colorbar and adjust its font size
    cbar.set_label('λ', fontsize=17)
    cbar.ax.tick_params(labelsize=15)
    ax.plot(default_df, default_dm, marker='*', markersize=14, color='k')
    
    ax.tick_params(axis='x', labelsize=17, pad=5)  # Adjust x-axis tick label size and padding
    ax.tick_params(axis='y', labelsize=17, pad=5)  # Adjust y-axis tick label size and padding
    ax.contour(df_range, dm_range, rBC_by_df_dm, levels=[0], colors='black', linestyles='dashed')
    

# %% [code]
def rBC_by_df_dm_plot(rBC, default_df, default_dm, L_no_effects, L_mp, r_to_pod_10yr_original, nf, nm, a, mu, Δmaternal_mortality=0.97, mx_contribution=[0.95, 0.75], lx_contribution=[0.8, 0.5], sex_ratio=[0.6,0.4], ε=10**(-5)):

    onset = np.where(np.sum(rBC, axis=0)>ε)[0][0] 
    y_label = '$λ(d_f, d_m)$'
    axis_labels = ['Female dispersal ($d_f)$', 'Male dispersal ($d_m$)']
    m_vals = np.arange(0,1.01,0.2)
    df_range = dm_range = np.arange(0, 1.01, 0.01)

    fig, axes = plt.subplots(1, m_vals.shape[-1], figsize=(45,6))
    fig.subplots_adjust(hspace=0.3, wspace=0.3)
    rBC_by_df_dm_m = np.zeros((m_vals.shape[-1], df_range.shape[-1], dm_range.shape[-1]))

    for i, m in enumerate(m_vals):
        rBC_by_df_dm_m[i] = rBC_to_df_dm_3D_vals(L_no_effects, L_mp, r_to_pod_10yr_original, nf, nm, a, mu, default_df=default_df, default_dm=default_dm, default_m=m, Δmaternal_mortality=Δmaternal_mortality, mx_contribution=mx_contribution, lx_contribution=lx_contribution, sex_ratio=sex_ratio, ε=ε)
        rBC_by_df_dm_subplot(rBC_by_df_dm_m[i], rBC, default_df, default_dm, onset, axes[i], ε=ε)
        axes[i].set_xlabel(axis_labels[0], fontsize=19)
        axes[i].set_title('m = {:.1f}'.format(m), fontsize=19)
    axes[0].set_ylabel(axis_labels[1], fontsize=19)

    plt.show()  
    

# %% [code]
rBC_by_df_dm_plot(rbc, 0.1, 0.1, L_no_effects, L_mp, r_to_pod_10yr_original, nf, nm, a, mu)

# %% [markdown]
## $\lambda$($m$, $d_f$, $d_m$)

# %% [code]
def rBC_default_dfm_changing_m(L_no_effects, L_mp, default_r_to_pod, nf, nm, a, mu, default_df=0.1, default_dm=0.1, default_m=0, Δmaternal_mortality=1, mx_contribution=np.array([0.95, 0.95]), lx_contribution=np.array([0.95, 0.95]), sex_ratio=[0.6,0.4], ε=10**(-5)):

    rBC, C, B, rB = different_onset_ages(L_no_effects, default_r_to_pod, Δmaternal_mortality=Δmaternal_mortality, mx_contribution=mx_contribution, lx_contribution=lx_contribution, sex_ratio=sex_ratio, return_all=True, ε=ε) 
    onset = np.where(np.sum(rBC, axis=0)>ε)[0][0] 

    df_range = dm_range = np.arange(0, 1.01, 0.2)
    m_range = np.arange(0, 1.01, 0.01)

    rBC_by_m = np.empty((df_range.shape[0], df_range.shape[0], m_range.shape[0]))

    for i, df in enumerate(df_range):
        for j, dm in enumerate(dm_range):
            for k, m in enumerate(m_range):
                rf_to_f, rm_to_f, rf_to_m, rm_to_m, avg_rf = rff_age_v2(df, dm, nf, nm, m, a, 1-mu)
                r_to_pod = np.array([rf_to_f, rm_to_f])
                r_to_pod_10yr = np.concatenate([r_to_pod[:,::5], (r_to_pod[:, -1]).reshape(2,1)], axis=1)
                rBC_vals, c, b, rb = different_onset_ages(L_no_effects, r_to_pod_10yr, Δmaternal_mortality=Δmaternal_mortality, mx_contribution=mx_contribution, lx_contribution=lx_contribution, sex_ratio=sex_ratio, return_all=True, ε=ε)         
                rBC_on_onset = np.sum(rBC_vals[:, onset], axis=0)
                rBC_by_m[i, j, k] = rBC_on_onset

    fig, axes = plt.subplots(df_range.shape[0], df_range.shape[0], figsize=(26,20))
    fig.subplots_adjust(hspace=0.3, wspace=0.3)
    for row in range(df_range.shape[0]): 
        axes[row, 0].set_ylabel('$d_f$ = {:.1f}\n'.format(df_range[row]) + '$\lambda$')
        for col in range(df_range.shape[0]):
            axes[0, col].set_title('$d_m$ = {:.1f}'.format(dm_range[col]))
            ax = axes[row, col]
            lambdas = rBC_by_m[row, col, :]
            ax.plot(m_range, lambdas, color='royalblue')
            ax.set_xlabel('$m$')
            ax.axhline(0, color='k')
            ax.plot(default_m, np.sum(rBC, axis=0)[onset], marker='*', color='salmon')

    plt.show()
    

# %% [code]
rBC_default_dfm_changing_m(L_no_effects, L_mp, r_to_pod_10yr_original, nf, nm, a, mu, default_df=0.1, default_dm=0.1, default_m=0, Δmaternal_mortality=0.97, mx_contribution=[0.95, 0.75], lx_contribution=[0.8, 0.5], sex_ratio=[0.6,0.4], ε=ε)

# %% [markdown]
## Sensitivity analysis for $\lambda$ according to survival, fecundity and maternal mortality changes

# %% [markdown]
### Explanation of Sensitivity Indices and Their Graphs
1. **First-order Sensitivity Indices (S1)**
- **Definition**: The first-order sensitivity index, \(S1\), measures the effect of varying a single parameter while keeping all other parameters fixed. It quantifies the direct contribution of each parameter to the output variance.
- **Interpretation**: A high \(S1\) value for a parameter indicates that this parameter alone has a significant impact on the output. If \(S1\) is close to 1, it means the output variance is almost entirely due to that parameter.
- **Graph**: The bar plot of \(S1\) indices shows the relative importance of each parameter. Error bars indicate the confidence intervals for these estimates.

2. **Total Sensitivity Indices (ST)**
- **Definition**: The total sensitivity index, \(ST\), measures the effect of varying a parameter including both its main effect and all its interaction effects with other parameters. It quantifies the total contribution of each parameter to the output variance, accounting for interactions.
- **Interpretation**: A high \(ST\) value indicates that the parameter, including its interactions with other parameters, significantly affects the output. If \(ST\) is close to 1, the parameter and its interactions are responsible for almost all the output variance.
- **Graph**: The bar plot of \(ST\) indices shows the overall importance of each parameter, including interactions. Error bars indicate the confidence intervals for these estimates.

Differences Between First-order and Total Sensitivity Indices
- **First-order Index (S1)**: Focuses only on the direct impact of each parameter on the output. It does not account for interactions with other parameters.
- **Total Sensitivity Index (ST)**: Includes both the direct impact and the interactions with other parameters, providing a more comprehensive measure of a parameter's importance.

Understanding the Graphs
- **First-order Sensitivity Indices Graph**: This graph displays the direct contribution of each parameter to the output variance. Higher bars indicate parameters that have a more significant direct effect on the output. The error bars provide a measure of the uncertainty in these estimates.
- **Total Sensitivity Indices Graph**: This graph shows the overall contribution of each parameter, including all interactions. Higher bars indicate parameters that are more important when considering both direct and interaction effects. The error bars show the uncertainty in these estimates.

# %% [code]
from SALib.sample import saltelli 
from SALib.analyze import sobol
problem = {
    'num_vars': 8,
    'names': ['lx_F', 'lx_M', 'mx_F', 'mx_M', 'mm',
              'df', 'dm', 'm'],
    'bounds': [
        [0, 1],  # lx_F
        [0, 1],  # lx_M
        [0, 1],  # mx_F
        [0, 1],  # mx_M
        [0, 1],  # mm
        [0, 1],  # df  (example biologically reasonable range)
        [0, 1],  # dm
        [0, 1],    # m
    ]
}


# %% [code]
def model2(x, L, onset=5, adjusted_age_dist_rho=True, ε=1e-50, ):

    # life-history modifiers
    lx_cont = np.array([x[0], x[1]])
    mx_cont = np.array([x[2], x[3]])
    mm = x[4]

    # dispersal & mating parameters
    df, dm, m = x[5], x[6], x[7]

    # constants (kept fixed for Sobol)
    nf = nm = 3
    a = 10

    # ---- relatedness calculation (generalized) ----
    r_to_pod = calc_rcoeff(L_mp=L_mp, m=m, df=df, dm=dm, nf=nf, nm=nm, a=a)

    r_to_pod_10yr = np.concatenate([r_to_pod[:, ::5], r_to_pod[:, -1][:, None]], axis=1)
    # ----------------------------------------------

    if adjusted_age_dist_rho:
        lamb = different_onset_ages(
            L,
            r_to_pod_10yr,
            Δmaternal_mortality=mm,
            lx_contribution=lx_cont,
            mx_contribution=mx_cont,
            sex_ratio=[0.6, 0.4],
            ε=ε,
            return_all=True
        )[0]
    else:
        lamb = different_onset_ages(
            L,
            r_to_pod_10yr,
            Δmaternal_mortality=mm,
            lx_contribution=lx_cont,
            mx_contribution=mx_cont,
            sex_ratio=[0.6, 0.4],
            ε=ε,
            return_all=True,
            age_dist=age_distribution(L_mp, ρ),
            ρ=ρ
        )[0]

    return np.sum(lamb, axis=0)[onset]


# %% [code]
N = 1024

param_values = saltelli.sample(problem, N, calc_second_order=True)
Y = np.array([model2(x, L_no_effects) for x in param_values])
Si = sobol.analyze(problem, Y)


# %% [code]
parameter_names = [
    'Female survival',
    'Male survival',
    'Female fecundity',
    'Male fecundity',
    'Maternal mortality',
    'Female dispersal',
    'Male dispersal',
    'Inbreeding rate'
]
param_names = [
    'lx_F', 'lx_M',
    'mx_F', 'mx_M',
    'mm',
    'df', 'dm', 'm'
]

# %% [code]
save_sobol_results(
    species_name="resident_KW",
    Si=Si,
    param_names=param_names,
    outdir="sa_results"
)


# %% [code]
import numpy as np
import matplotlib.pyplot as plt

S1 = Si['S1']
ST = Si['ST']
S1_conf = Si['S1_conf']
ST_conf = Si['ST_conf']

y_pos = np.arange(len(parameter_names))

fig, axes = plt.subplots(1, 2, figsize=(18, 9))
plt.subplots_adjust(wspace=0.6)

# ---------- First-order ----------
ax = axes[0]
ax.barh(
    y_pos,
    S1,
    xerr=S1_conf,
    capsize=5
)
ax.set_yticks(y_pos)
ax.set_yticklabels(parameter_names)
ax.set_xlabel('First-order sensitivity index', fontsize=21)
ax.tick_params(axis='both', which='major', labelsize=20)
ax.set_xlim(0, np.nanmax(S1 + S1_conf) * 1.15)

# ---------- Total-order ----------
ax = axes[1]
ax.barh(
    y_pos,
    ST,
     capsize=5)
    # ,
#     xerr=ST_conf
# )
ax.set_yticks(y_pos)
ax.set_yticklabels(parameter_names)
ax.set_xlabel('Total sensitivity index', fontsize=21)
ax.tick_params(axis='both', which='major', labelsize=20)
ax.set_xlim(0, np.nanmax(ST + ST_conf) * 1.15)

plt.show()


# %% [markdown]
## $\lambda$($lx$, $mx$)

# %% [code]
def λ_by_help(L_no_effects, r_to_pod, Δmaternal_mortality=1, mx_contribution=np.array([0.95, 0.95]), lx_contribution=np.array([0.95, 0.95]), sex_ratio=[0.6,0.4], ε=10**(-5), age_dist=None, ρ=None, onset='auto', plot_age_dist=False, view_plot=False):

    rBC, C, B, rB = different_onset_ages(L_no_effects, r_to_pod, Δmaternal_mortality=Δmaternal_mortality, mx_contribution=mx_contribution, lx_contribution=lx_contribution, sex_ratio=sex_ratio, return_all=True, ε=ε, age_dist=age_dist, ρ=ρ, plot_age_dist=False) 
    d = L_no_effects.shape[-1] // 2
    if onset == 'auto':    
        try:
            onset = np.where(np.sum(rBC, axis=0)>ε)[0][0] 
        except IndexError:
            onset = 4
    
    mx_contribution_range = lx_contribution_range = np.linspace(0.005,1,150)
    
    rBC_by_mx = np.empty((2, mx_contribution_range.shape[0]))
    rBC_by_lx = np.empty((2, lx_contribution_range.shape[0]))
    rB_by_mx = np.empty((2, mx_contribution_range.shape[0]))
    rB_by_lx = np.empty((2, lx_contribution_range.shape[0]))
    C_by_mx = np.empty((2, mx_contribution_range.shape[0]))
    C_by_lx = np.empty((2, lx_contribution_range.shape[0]))

    rBC_lx_fchange = np.empty((2, lx_contribution_range.shape[0]))
    rBC_lx_mchange = np.empty((2, lx_contribution_range.shape[0]))
    rB_lx_fchange = np.empty((2, lx_contribution_range.shape[0]))
    rB_lx_mchange = np.empty((2, lx_contribution_range.shape[0]))
    C_lx_fchange = np.empty((2, lx_contribution_range.shape[0]))
    C_lx_mchange = np.empty((2, lx_contribution_range.shape[0]))

    rBC_mx_fchange = np.empty((2, mx_contribution_range.shape[0]))
    rBC_mx_mchange = np.empty((2, mx_contribution_range.shape[0]))
    rB_mx_fchange = np.empty((2, mx_contribution_range.shape[0]))
    rB_mx_mchange = np.empty((2, mx_contribution_range.shape[0]))
    C_mx_fchange = np.empty((2, mx_contribution_range.shape[0]))
    C_mx_mchange = np.empty((2, mx_contribution_range.shape[0]))

    age_dists_lx_f = np.empty((15, 2, d))
    age_dists_lx_m = np.empty((15, 2, d))
    age_dists_mx_f = np.empty((15, 2, d))
    age_dists_mx_m = np.empty((15, 2, d))
    cnt_age = 0
    
    rho_arr_lx = np.empty((2, lx_contribution_range.shape[0]))
    rho_arr_mx = np.empty((2, mx_contribution_range.shape[0]))
    
    for i in range(lx_contribution_range.shape[0]):
        # rBC_lx_m, C_lx_m, B_lx_m, rB_lx_m = different_onset_ages(L_no_effects, r_to_pod, Δmaternal_mortality=Δmaternal_mortality, mx_contribution=mx_contribution, lx_contribution=[lx_contribution_range[i], lx_contribution_range[i]], sex_ratio=sex_ratio, return_all=True, ε=ε) 
        # rBC_mx_f, C_mx_f, B_mx_f, rB_mx_f = different_onset_ages(L_no_effects, r_to_pod, Δmaternal_mortality=Δmaternal_mortality, mx_contribution=[mx_contribution_range[i], mx_contribution_range[i]], lx_contribution=lx_contribution, sex_ratio=sex_ratio, return_all=True, ε=ε) 
        rBC_lx_f, C_lx_f, B_lx_f, rB_lx_f, ages_lx_f = different_onset_ages(L_no_effects, r_to_pod, Δmaternal_mortality=Δmaternal_mortality, mx_contribution=mx_contribution, lx_contribution=[lx_contribution_range[i], lx_contribution[1]], sex_ratio=sex_ratio, return_all=True, ε=ε, age_dist=age_dist, ρ=ρ, plot_age_dist=plot_age_dist) 
        rBC_lx_m, C_lx_m, B_lx_m, rB_lx_m, ages_lx_m = different_onset_ages(L_no_effects, r_to_pod, Δmaternal_mortality=Δmaternal_mortality, mx_contribution=mx_contribution, lx_contribution=[lx_contribution[0], lx_contribution_range[i]], sex_ratio=sex_ratio, return_all=True, ε=ε, age_dist=age_dist, ρ=ρ, plot_age_dist=plot_age_dist) 
        rBC_mx_f, C_mx_f, B_mx_f, rB_mx_f, ages_mx_f = different_onset_ages(L_no_effects, r_to_pod, Δmaternal_mortality=Δmaternal_mortality, mx_contribution=[mx_contribution_range[i], mx_contribution[1]], lx_contribution=lx_contribution, sex_ratio=sex_ratio, return_all=True, ε=ε, age_dist=age_dist, ρ=ρ, plot_age_dist=plot_age_dist) 
        rBC_mx_m, C_mx_m, B_mx_m, rB_mx_m, ages_mx_m = different_onset_ages(L_no_effects, r_to_pod, Δmaternal_mortality=Δmaternal_mortality, mx_contribution=[mx_contribution[0], mx_contribution_range[i]], lx_contribution=lx_contribution, sex_ratio=sex_ratio, return_all=True, ε=ε, age_dist=age_dist, ρ=ρ, plot_age_dist=plot_age_dist) 
        # print(ages_lx_f[1])
        rho_arr_lx[:, i] = ages_lx_f[1][onset], ages_lx_m[1][onset]
        rho_arr_mx[:, i] = ages_mx_f[1][onset], ages_mx_m[1][onset]
        
        rBC_by_mx[:, i] = np.sum(rBC_mx_f[:, onset], axis=0), np.sum(rBC_mx_m[:, onset], axis=0)
        rBC_by_lx[:, i] = np.sum(rBC_lx_f[:, onset], axis=0), np.sum(rBC_lx_m[:, onset], axis=0)
        rB_by_mx[:, i] = np.sum(rB_mx_f[:, onset], axis=0), np.sum(rB_mx_m[:, onset], axis=0)
        rB_by_lx[:, i] = np.sum(rB_lx_f[:, onset], axis=0), np.sum(rB_lx_m[:, onset], axis=0)
        C_by_mx[:, i] = np.sum(C_mx_f[:, onset], axis=0), np.sum(C_mx_m[:, onset], axis=0)
        C_by_lx[:, i] = np.sum(C_lx_f[:, onset], axis=0), np.sum(C_lx_m[:, onset], axis=0)

        rBC_lx_fchange[:, i] = rBC_lx_f[:, onset]
        rBC_lx_mchange[:, i] = rBC_lx_m[:, onset]
        rB_lx_fchange[:, i] = rB_lx_f[:, onset]
        rB_lx_mchange[:, i] = rB_lx_m[:, onset]
        C_lx_fchange[:, i] = C_lx_f[:, onset]
        C_lx_mchange[:, i] = C_lx_m[:, onset]

        rBC_mx_fchange[:, i] = rBC_mx_f[:, onset]
        rBC_mx_mchange[:, i] = rBC_mx_m[:, onset]
        rB_mx_fchange[:, i] = rB_mx_f[:, onset]
        rB_mx_mchange[:, i] = rB_mx_m[:, onset]
        C_mx_fchange[:, i] = C_mx_f[:, onset]
        C_mx_mchange[:, i] = C_mx_m[:, onset]

        cnt_age +=1
        
        if view_plot:
            if cnt_age % 3 == 0:
                # age_dists_lx_f[cnt_age, :, :] = ages_lx_f
                # age_dists_lx_m[cnt_age, :, :] = ages_lx_m
                # age_dists_mx_f[cnt_age, :, :] = ages_mx_f
                # age_dists_mx_m[cnt_age, :, :] = ages_mx_m
    
                plot_age_distribution(ages_lx_f[0][onset, :, :])
                plt.show()
                plot_age_distribution(ages_lx_m[0][onset, :, :])
                plt.show()
                plot_age_distribution(ages_mx_f[0][onset, :, :])
                plt.show()
                plot_age_distribution(ages_mx_f[0][onset, :, :])
                plt.show()
                
                print(cnt_age)
            
        # rBC_by_mx[i] = np.sum(rBC_mx_f[:, onset], axis=0)
        # rBC_by_lx[i] = np.sum(rBC_lx_m[:, onset], axis=0)

    fig, axes = plt.subplots(3, 2, figsize=(16,18), sharex=True)
    fig.subplots_adjust(hspace=0.15, wspace=0.3)

    ax = axes[0, 0] # this row is for females
    ax.plot(lx_contribution_range, rho_arr_lx[0], color='salmon')
    ax.plot(lx_contribution_range, rho_arr_lx[1], color='cornflowerblue')
    ax.set_ylabel('ρ', fontsize=22)
    ax.tick_params(axis='both', which='major', labelsize=18)
    # add_image(ax, female_symbol, xy, zoom=0.1)
    
    ax = axes[0, 1]
    ax.plot(lx_contribution_range, rho_arr_mx[0], color='salmon')
    ax.plot(lx_contribution_range, rho_arr_mx[1], color='cornflowerblue')
    ax.tick_params(axis='both', which='major', labelsize=18)
    # add_image(ax, female_symbol, xy, zoom=0.1)
    
    ax = axes[1, 0]
    # ax.set_ylabel('$\lambda$', fontsize=22)
    # ax.set_xlabel(r'$\frac{\mathrm{survival \ in \ a \ non-menopausal \ population}}{\mathrm{survival \ in \ a \ menopausal \ population}}$', fontsize=25)
    ax.plot(lx_contribution_range, rBC_by_lx[0, :], c='#efbbcc', lw=2)
    ax.plot(lx_contribution_range, C_by_lx[0, :], c='#db7093', lw=2)
    ax.plot(lx_contribution_range, rB_by_lx[0, :], c='#722f37', lw=2)
    ax.plot(lx_contribution[0], np.sum(rBC, axis=0)[onset], c='#efbbcc', marker='o')
    # ax.plot(lx_contribution[0], np.sum(C, axis=0)[onset], c='#FFC98B', marker='o')
    # ax.plot(lx_contribution[0], np.sum(rB, axis=0)[onset], c='#478BA2', marker='o')
    ax.axhline(0, c='k', lw=1)
    ax.tick_params(axis='both', which='major', labelsize=18)
    ax.set_ylim(-0.0075, 0.016)
    add_image(ax, female_symbol, (0.95, 0.0135), zoom=0.012)
    

    ax = axes[1, 1] # this row is for females
    # ax.set_xlabel(r'$\frac{\mathrm{fecundity \ in \ a \ non-menopausal \ population}}{\mathrm{fecundity \ in \ a \ menopausal \ population}}$', fontsize=25)
    # ax.set_ylabel('$\lambda$', fontsize=22)
    ax.plot(mx_contribution_range, rBC_by_mx[0, :], c='#efbbcc', lw=2)
    ax.plot(mx_contribution_range, C_by_mx[0, :], c='#db7093', lw=2)
    ax.plot(mx_contribution_range, rB_by_mx[0, :], c='#722f37', lw=2)
    ax.plot(mx_contribution[0], np.sum(rBC, axis=0)[onset], c='#efbbcc', marker='o')
    ax.axhline(0, c='k', lw=1)
    ax.tick_params(axis='both', which='major', labelsize=18)
    ax.set_ylim(-0.0075, 0.016)
    add_image(ax, female_symbol, (0.95, 0.0135), zoom=0.012)

    # ax.plot(mx_contribution_range, rBC_mx_mchange[0, :], c='cornflowerblue', ls='--')
    # ax.plot(mx_contribution_range, C_mx_mchange[0, :], c='#FFC98B', ls='--')
    # ax.plot(mx_contribution_range, rB_mx_mchange[0, :], c='#478BA2', ls='--')
    
    # ax.plot(mx_contribution_range, rBC_mx_mchange[1, :], c='cornflowerblue', ls=':')
    # ax.plot(mx_contribution_range, C_mx_mchange[1, :], c='#FFC98B', ls=':')
    # ax.plot(mx_contribution_range, rB_mx_mchange[1, :], c='#478BA2', ls=':')
    

    ax = axes[2, 0] # row for males
    ax.set_xlabel(r'$\frac{\mathrm{survival \ in \ a \ non-menopausal \ population}}{\mathrm{survival \ in \ a \ menopausal \ population}}$', fontsize=25)
    # ax.set_ylabel('$\lambda$', fontsize=22)
    ax.plot(lx_contribution_range, rBC_by_lx[1, :], c='#add8e6', lw=2)
    ax.plot(lx_contribution_range, C_by_lx[1, :], c='cornflowerblue', lw=2)
    ax.plot(lx_contribution_range, rB_by_lx[1, :], c='#003366', lw=2)
    ax.plot(lx_contribution[1], np.sum(rBC, axis=0)[onset], c='#add8e6', marker='o')
    ax.axhline(0, c='k', lw=1)
    ax.tick_params(axis='both', which='major', labelsize=18)
    ax.set_ylim(-0.0075, 0.016)
    add_image(ax, male_symbol, (0.95, 0.0135), zoom=0.0152)

    ax = axes[2, 1]
    ax.set_xlabel(r'$\frac{\mathrm{fecundity \ in \ a \ non-menopausal \ population}}{\mathrm{fecundity \ in \ a \ menopausal \ population}}$', fontsize=25)
    # ax.set_ylabel('$\lambda$', fontsize=22)
    ax.plot(mx_contribution_range, rBC_by_mx[1, :], c='#add8e6', lw=2)
    ax.plot(mx_contribution_range, C_by_mx[1, :], c='cornflowerblue', lw=2)
    ax.plot(mx_contribution_range, rB_by_mx[1, :], c='#003366', lw=2)
    ax.plot(mx_contribution[1], np.sum(rBC, axis=0)[onset], c='#add8e6', marker='o')
    ax.axhline(0, c='k', lw=1)
    ax.tick_params(axis='both', which='major', labelsize=18)
    ax.set_ylim(-0.0075, 0.016)
    add_image(ax, male_symbol, (0.95, 0.0135), zoom=0.0152)
    
    plt.show()

# %% [code]
def λ_by_help_A(L_no_effects, r_to_pod, Δmaternal_mortality=1,
                mx_contribution=np.array([0.95, 0.95]),
                lx_contribution=np.array([0.95, 0.95]),
                sex_ratio=[0.6, 0.4], ε=10**(-5),
                age_dist=None, ρ=None,
                onset='auto', plot_age_dist=False,
                view_plot=False, letters=None, uppercase=True, xcoord=-0.25, ycoord=0.92, panel_label_size=22):

    rBC, C, B, rB = different_onset_ages(
        L_no_effects, r_to_pod,
        Δmaternal_mortality=Δmaternal_mortality,
        mx_contribution=mx_contribution,
        lx_contribution=lx_contribution,
        sex_ratio=sex_ratio, return_all=True, ε=ε,
        age_dist=age_dist, ρ=ρ, plot_age_dist=False
    )

    d = L_no_effects.shape[-1] // 2

    if onset == 'auto':
        try:
            onset = np.where(np.sum(rBC, axis=0) > ε)[0][0]
        except IndexError:
            onset = 4

    mx_contribution_range = lx_contribution_range = np.linspace(0.005, 1, 150)

    rBC_by_mx = np.empty((2, mx_contribution_range.shape[0]))
    rBC_by_lx = np.empty((2, lx_contribution_range.shape[0]))
    rB_by_mx = np.empty((2, mx_contribution_range.shape[0]))
    rB_by_lx = np.empty((2, lx_contribution_range.shape[0]))
    C_by_mx = np.empty((2, mx_contribution_range.shape[0]))
    C_by_lx = np.empty((2, lx_contribution_range.shape[0]))

    cnt_age = 0

    for i in range(lx_contribution_range.shape[0]):

        rBC_lx_f, C_lx_f, B_lx_f, rB_lx_f, ages_lx_f = different_onset_ages(
            L_no_effects, r_to_pod,
            Δmaternal_mortality=Δmaternal_mortality,
            mx_contribution=mx_contribution,
            lx_contribution=[lx_contribution_range[i], lx_contribution[1]],
            sex_ratio=sex_ratio, return_all=True, ε=ε,
            age_dist=age_dist, ρ=ρ, plot_age_dist=plot_age_dist
        )
        rBC_lx_m, C_lx_m, B_lx_m, rB_lx_m, ages_lx_m = different_onset_ages(
            L_no_effects, r_to_pod,
            Δmaternal_mortality=Δmaternal_mortality,
            mx_contribution=mx_contribution,
            lx_contribution=[lx_contribution[0], lx_contribution_range[i]],
            sex_ratio=sex_ratio, return_all=True, ε=ε,
            age_dist=age_dist, ρ=ρ, plot_age_dist=plot_age_dist
        )
        rBC_mx_f, C_mx_f, B_mx_f, rB_mx_f, ages_mx_f = different_onset_ages(
            L_no_effects, r_to_pod,
            Δmaternal_mortality=Δmaternal_mortality,
            mx_contribution=[mx_contribution_range[i], mx_contribution[1]],
            lx_contribution=lx_contribution,
            sex_ratio=sex_ratio, return_all=True, ε=ε,
            age_dist=age_dist, ρ=ρ, plot_age_dist=plot_age_dist
        )
        rBC_mx_m, C_mx_m, B_mx_m, rB_mx_m, ages_mx_m = different_onset_ages(
            L_no_effects, r_to_pod,
            Δmaternal_mortality=Δmaternal_mortality,
            mx_contribution=[mx_contribution[0], mx_contribution_range[i]],
            lx_contribution=lx_contribution,
            sex_ratio=sex_ratio, return_all=True, ε=ε,
            age_dist=age_dist, ρ=ρ, plot_age_dist=plot_age_dist
        )

        # sum over kin categories at onset age
        rBC_by_mx[:, i] = np.sum(rBC_mx_f[:, onset], axis=0), np.sum(rBC_mx_m[:, onset], axis=0)
        rBC_by_lx[:, i] = np.sum(rBC_lx_f[:, onset], axis=0), np.sum(rBC_lx_m[:, onset], axis=0)
        rB_by_mx[:, i] = np.sum(rB_mx_f[:, onset], axis=0), np.sum(rB_mx_m[:, onset], axis=0)
        rB_by_lx[:, i] = np.sum(rB_lx_f[:, onset], axis=0), np.sum(rB_lx_m[:, onset], axis=0)
        C_by_mx[:, i] = np.sum(C_mx_f[:, onset], axis=0), np.sum(C_mx_m[:, onset], axis=0)
        C_by_lx[:, i] = np.sum(C_lx_f[:, onset], axis=0), np.sum(C_lx_m[:, onset], axis=0)

        cnt_age += 1

        if view_plot and cnt_age % 3 == 0:
            plot_age_distribution(ages_lx_f[0][onset, :, :])
            plt.show()
            plot_age_distribution(ages_lx_m[0][onset, :, :])
            plt.show()
            plot_age_distribution(ages_mx_f[0][onset, :, :])
            plt.show()
            plot_age_distribution(ages_mx_m[0][onset, :, :])
            plt.show()
            print(cnt_age)

    # ---------- X-axes (Version A): 1 - ratio ----------
    # here ratio == contribution_range
    surv_rel_A = 1.0 - lx_contribution_range      # survival panels
    fec_rel_A  = 1.0 - mx_contribution_range      # fecundity panels

    lx_contribution = np.asarray(lx_contribution)
    mx_contribution = np.asarray(mx_contribution)

    surv_rel_A_f0 = 1.0 - lx_contribution[0]
    surv_rel_A_m0 = 1.0 - lx_contribution[1]
    fec_rel_A_f0  = 1.0 - mx_contribution[0]
    fec_rel_A_m0  = 1.0 - mx_contribution[1]

    # ---------- FIGURE: 2 x 2 panels ----------

    fig, axes = plt.subplots(2, 2, figsize=(14, 10))
    fig.subplots_adjust(hspace=0.25, wspace=0.35)

    # A: females – survival
    ax = axes[0, 0]
    ax.plot(surv_rel_A, rBC_by_lx[0, :], c='#722f37', lw=2)  # λ (benefit - cost)
    ax.plot(surv_rel_A, C_by_lx[0, :],   c='#db7093', lw=2)  # cost
    ax.plot(surv_rel_A, rB_by_lx[0, :],  c='#efbbcc', lw=2)  # benefit
    ax.plot(surv_rel_A_f0, np.sum(rBC, axis=0)[onset],
            c='#722f37', marker='o')
    ax.axhline(0, c='k', lw=1)
    ax.tick_params(axis='both', which='major', labelsize=18)
    ax.set_ylim(-0.0075, 0.016)
    ax.set_ylabel(r'$\lambda$', fontsize=22)
    # ax.set_xlabel('Relative survival benefit', fontsize=18)
    add_image(ax, female_symbol, (0.025, 0.0135), zoom=0.012)

    # B: females – fecundity
    ax = axes[0, 1]
    ax.plot(fec_rel_A, rBC_by_mx[0, :], c='#722f37', lw=2)
    ax.plot(fec_rel_A, C_by_mx[0, :],   c='#db7093', lw=2)
    ax.plot(fec_rel_A, rB_by_mx[0, :],  c='#efbbcc', lw=2)
    ax.plot(fec_rel_A_f0, np.sum(rBC, axis=0)[onset],
            c='#722f37', marker='o')
    ax.axhline(0, c='k', lw=1)
    ax.tick_params(axis='both', which='major', labelsize=18)
    ax.set_ylim(-0.0075, 0.016)
    # ax.set_xlabel('Relative fecundity benefit', fontsize=18)
    add_image(ax, female_symbol, (0.025, 0.0135), zoom=0.012)

    # C: males – survival
    ax = axes[1, 0]
    ax.plot(surv_rel_A, rBC_by_lx[1, :], c='#003366', lw=2)
    ax.plot(surv_rel_A, C_by_lx[1, :],   c='cornflowerblue', lw=2)
    ax.plot(surv_rel_A, rB_by_lx[1, :],  c='#add8e6', lw=2)
    ax.plot(surv_rel_A_m0, np.sum(rBC, axis=0)[onset],
            c='#003366', marker='o')
    ax.axhline(0, c='k', lw=1)
    ax.tick_params(axis='both', which='major', labelsize=18)
    ax.set_ylim(-0.0075, 0.016)
    ax.set_ylabel(r'$\lambda$', fontsize=22)
    ax.set_xlabel('Relative survival benefit (1-ratio)', fontsize=18)
    add_image(ax, male_symbol, (0.025, 0.0135), zoom=0.0152)

    # D: males – fecundity
    ax = axes[1, 1]
    ax.plot(fec_rel_A, rBC_by_mx[1, :], c='#003366', lw=2)
    ax.plot(fec_rel_A, C_by_mx[1, :],   c='cornflowerblue', lw=2)
    ax.plot(fec_rel_A, rB_by_mx[1, :],  c='#add8e6', lw=2)
    ax.plot(fec_rel_A_m0, np.sum(rBC, axis=0)[onset],
            c='#003366', marker='o')
    ax.axhline(0, c='k', lw=1)
    ax.tick_params(axis='both', which='major', labelsize=18)
    ax.set_ylim(-0.0075, 0.016)
    ax.set_xlabel('Relative fecundity benefit (1-ratio)', fontsize=18)
    add_image(ax, male_symbol, (0.025, 0.0135), zoom=0.0152)

                    
    fig_panel_labels(axes, letters=letters, uppercase=uppercase, xcoord=xcoord, ycoord=ycoord, panel_label_size=panel_label_size)
    plt.show()


# %% [code]
λ_by_help_A(L_no_effects, r_to_pod_10yr_original, Δmaternal_mortality=0.97, mx_contribution=[0.95, 0.75], lx_contribution=[0.8, 0.5], sex_ratio=[0.6,0.4], ε=10**(-50), onset=5, plot_age_dist=True)#, age_dist=age_distribution(L_mp, ρ), ρ=ρ)


# %% [code]
def λ_by_help_merged(L_no_effects, r_to_pod, Δmaternal_mortality=1,
                    mx_contribution=np.array([0.95, 0.95]),
                    lx_contribution=np.array([0.95, 0.95]),
                    sex_ratio=[0.6, 0.4], ε=10**(-5),
                    age_dist=None, ρ=None,
                    onset='auto', plot_age_dist=False,
                    view_plot=False):

    # 1. Initial calculation to find baseline lambda and onset age
    # This provides the single "starting point" for our markers
    rBC_baseline, _, _, _ = different_onset_ages(
        L_no_effects, r_to_pod,
        Δmaternal_mortality=Δmaternal_mortality,
        mx_contribution=mx_contribution,
        lx_contribution=lx_contribution,
        sex_ratio=sex_ratio, return_all=True, ε=ε,
        age_dist=age_dist, ρ=ρ, plot_age_dist=False
    )

    if onset == 'auto':
        try:
            onset = np.where(np.sum(rBC_baseline, axis=0) > ε)[0][0]
        except IndexError:
            onset = 4

    # The baseline value at the onset age
    baseline_y = np.sum(rBC_baseline, axis=0)[onset]

    # 2. Define the range for sensitivity
    ratios = np.linspace(0.005, 1, 150)
    rel_benefit_x = 1.0 - ratios

    rBC_by_mx = np.empty((2, ratios.shape[0]))
    rBC_by_lx = np.empty((2, ratios.shape[0]))

    # 3. Data Generation Loop
    for i in range(ratios.shape[0]):
        # Female-specific variations (holding male constant)
        res_lx_f, _, _, _ = different_onset_ages(
            L_no_effects, r_to_pod, Δmaternal_mortality=Δmaternal_mortality,
            mx_contribution=mx_contribution,
            lx_contribution=[ratios[i], lx_contribution[1]],
            sex_ratio=sex_ratio, return_all=True, ε=ε
        )
        res_mx_f, _, _, _= different_onset_ages(
            L_no_effects, r_to_pod, Δmaternal_mortality=Δmaternal_mortality,
            mx_contribution=[ratios[i], mx_contribution[1]],
            lx_contribution=lx_contribution,
            sex_ratio=sex_ratio, return_all=True, ε=ε
        )
        
        # Male-specific variations (holding female constant)
        res_lx_m, _, _, _ = different_onset_ages(
            L_no_effects, r_to_pod, Δmaternal_mortality=Δmaternal_mortality,
            mx_contribution=mx_contribution,
            lx_contribution=[lx_contribution[0], ratios[i]],
            sex_ratio=sex_ratio, return_all=True, ε=ε
        )
        res_mx_m, _, _, _ = different_onset_ages(
            L_no_effects, r_to_pod, Δmaternal_mortality=Δmaternal_mortality,
            mx_contribution=[mx_contribution[0], ratios[i]],
            lx_contribution=lx_contribution,
            sex_ratio=sex_ratio, return_all=True, ε=ε
        )

        rBC_by_lx[0, i] = np.sum(res_lx_f[:, onset], axis=0)
        rBC_by_lx[1, i] = np.sum(res_lx_m[:, onset], axis=0)
        rBC_by_mx[0, i] = np.sum(res_mx_f[:, onset], axis=0)
        rBC_by_mx[1, i] = np.sum(res_mx_m[:, onset], axis=0)

    # 4. FIGURE: Consolidated Single Plot
    fig, ax = plt.subplots(1, 1, figsize=(12, 8))

    # --- Female Curves (Pink) ---
    ax.plot(rel_benefit_x, rBC_by_lx[0, :], c='hotpink', lw=3, label='Female Survival ($l_x$)')
    ax.plot(rel_benefit_x, rBC_by_mx[0, :], c='hotpink', lw=3, ls='--', label='Female Fecundity ($m_x$)')
    
    # --- Male Curves (Blue) ---
    ax.plot(rel_benefit_x, rBC_by_lx[1, :], c='cornflowerblue', lw=3, label='Male Survival ($l_x$)')
    ax.plot(rel_benefit_x, rBC_by_mx[1, :], c='cornflowerblue', lw=3, ls='--', label='Male Fecundity ($m_x$)')

    # --- The Two Baselines ---
    # Female baseline point: at (1 - current female contribution ratio)
    ax.plot(1.0 - lx_contribution[0], baseline_y, c='hotpink', 
            marker='*', markersize=22, markeredgecolor='k')
    ax.plot(1.0 - mx_contribution[0], baseline_y, c='hotpink', 
            marker='*', markersize=22, markeredgecolor='k')
    
    
    # Male baseline point: at (1 - current male contribution ratio)
    ax.plot(1.0 - lx_contribution[1], baseline_y, c='cornflowerblue', 
            marker='*', markersize=22, markeredgecolor='k')
    ax.plot(1.0 - mx_contribution[1], baseline_y, c='cornflowerblue', 
            marker='*', markersize=22, markeredgecolor='k')

    # Formatting
    ax.axhline(0, c='k', lw=1.5, alpha=0.6)
    ax.set_ylim(-0.0075, 0.016)
    ax.set_xlim(0, 1)
    ax.set_xlabel('Relative benefit', fontsize=25)
    ax.set_ylabel(r'$\lambda$', fontsize=25)
    ax.tick_params(axis='both', which='major', labelsize=23)
    
    ax.legend(fontsize=16, loc='upper left', frameon=True, ncol=2)
    # ax.grid(True, linestyle=':', alpha=0.5)

    plt.tight_layout()
    plt.show()

# %% [code]
λ_by_help_merged(L_no_effects, r_to_pod_10yr_original, Δmaternal_mortality=0.97, mx_contribution=[0.95, 0.75], lx_contribution=[0.8, 0.5], sex_ratio=[0.6,0.4], ε=10**(-50), onset=5, plot_age_dist=True)#, age_dist=age_distribution(L_mp, ρ), ρ=ρ)


# %% [code]
def fig_panel_labels(axes, letters=None, uppercase=True,
                     xcoord=-0.17, ycoord=0.92, panel_label_size=None):
    import string
    from matplotlib.font_manager import FontProperties
    import numpy as np

    # Convert rcParam to a numeric size
    if panel_label_size is None:
        fp = FontProperties(size=plt.rcParams['axes.titlesize'])
        panel_label_size = fp.get_size() * 1.3

    # Generate default letters if needed
    if letters is None:
        letters = string.ascii_uppercase if uppercase else string.ascii_lowercase

    axes_arr = np.array(axes, dtype=object)
    flat_axes = list(axes_arr.flat)          # make a list so we can iterate safely
    n_axes = len(flat_axes)

    # Truncate / slice letters to the number of axes
    letters = list(letters[:n_axes])

    # --- handle xcoord ---
    # scalar: same for all panels
    # sequence length == ncols: per-column xcoord
    # sequence length == n_axes: per-panel xcoord
    if hasattr(xcoord, "__len__") and not isinstance(xcoord, str):
        xcoord_list = list(xcoord)
        if axes_arr.ndim == 2 and len(xcoord_list) == axes_arr.shape[1]:
            # per-column values
            nrows, ncols = axes_arr.shape
            xs = []
            for i in range(n_axes):
                col = i % ncols
                xs.append(xcoord_list[col])
        elif len(xcoord_list) == n_axes:
            xs = xcoord_list
        else:
            raise ValueError("xcoord must be a scalar, length ncols, or length n_axes")
    else:
        xs = [xcoord] * n_axes

    ys = [ycoord] * n_axes  # keep ycoord simple (one value for all)

    # --- annotate ---
    annots = []
    for ax, letter, xc, yc in zip(flat_axes, letters, xs, ys):
        annots.append(
            ax.annotate(
                letter, (xc, yc),
                xycoords='axes fraction',
                fontsize=panel_label_size
            )
        )
    return annots


# %% [markdown]
# Bibliography
> Johnstone R.A. & Cant M.A. (2010). The evolution of menopause in cetaceans and humans: the role of demography. Proc. R. Soc. B.2773765–3771
> 
> Nielsen M.L.K., Ellis S., Towers J.R., Doniol-Valcroze T., Franks D.W., Cant M.A., Weiss M.N., Johnstone R.A., Balcomb K.C. 3rd, Ellifrit D.K., Croft D.P. (2021). A long postreproductive life span is a shared trait among genetically distinct killer whale populations. Ecol Evol. Jun 16;11(13):9123-9136. doi: 10.1002/ece3.7756. PMID: 34257948; PMCID: PMC8258204.
>
> Olesiuk, P. F., Graeme M. E., John K. B. F. Life History and Population Dynamics of Northern Resident Killer 
Whales (Orcinus Orca) in British Columbia. Canadian Science Advisory Secretariat (CSAC). (2005)
>> 
> Rogers, A. (1993).R. Why menopause?. Evol Ecol 7, 406–43)72

# %% [code]
```
